# Predicting cortical signal mixing in coarse-resolution fMRI: a forward-operator analysis of surface mapping and the conditional benefit of correction

**DOI:** 10.64898/2026.10.27.762317

**Authors:** Halil Aziz Velioğlu

**Author notes:** Correspondence: Halil Aziz Velioğlu, Department of Physiology, School of Medicine, Istanbul Medipol University, Istanbul, Türkiye.

## Abstract

Surface-based fMRI projection can mix signals from cortical locations that are close in the volume but distant along the folded cortical sheet. We constructed an anatomy- and acquisition-informed linear forward operator to estimate this mixing and evaluated a regularised surface reconstruction that inverts it. In HCP 7T geometry, the operator-predicted net far-cortex contribution to a standard surface value exceeded 5% in 4%, 10% and 22% of the group-average cortex at 1.6 mm and at emulated 2.4 and 3.2 mm resolution. Including the modelled far component improved predictions of emulated coarser maps and of real coarser StudyForrest sessions in high-leakage regions. Against finer-resolution, cross-run fMRI references, the regularised reconstruction reduced the residual error in HCP development data (n = 15) and in a pre-specified, family-independent confirmatory analysis (158 participants in 86 families). The improvement increased with a response-weighted contamination measure in HCP and in an independent 3T dataset (35 test participants), although a coarse-data-only version of the measure did not generalise to 3T, where an anatomical atlas predicted the gain at twofold coarsening. Overall performance improved at twofold, but not 1.5-fold, coarsening in the 3T sample. Ablation and data-reduction analyses were consistent with an important contribution from local deconvolution but did not uniquely identify the mechanism. Benefits were smaller or absent for multi-run averages, and no robust improvement across separately acquired sessions was established. The leakage maps characterise model-predicted, signed mixing rather than directly measured signal origins, and every gain is measured against an fMRI reference rather than against the true cortical response. Code, analysis plans and group leakage maps (7T and 3T) are provided in a public repository.

## 1. Introduction

Mapping volumetric fMRI onto the cortical surface is widely used for analyses of topographic maps, laminar-adjacent questions and group comparisons in a common surface space. The standard approach is ribbon-constrained mapping, as in the HCP minimal preprocessing pipelines (Van Essen et al., 2013; Glasser et al., 2013): it averages the voxels that intersect the cortical ribbon around each vertex, weighted by their overlap. This is a weighted projection, not an inversion of the acquisition, so the mixing produced by voxel sampling and the point-spread function (PSF) remains in the surface data. Volumetric processing is known to damage localisation (Coalson et al., 2018), and blur accumulates at every resampling step (Polimeni et al., 2018; Wang et al., 2022). For an individual anatomy and an assumed acquisition model, however, the contribution of surface-distant cortex to each surface value, and its relationship to the performance of a reconstruction that models it, have not been quantified vertex by vertex.

Inverting an explicit cortex-to-voxel operator under an anatomical prior has precedents: anatomically informed basis functions (Kiebel et al., 2000), an inverse formulation of volume-to-surface projection (Thiébaut Lonjaret et al., 2017) and a unified surface/volume inference for arterial spin labelling (Kirk et al., 2023) pose the problem this way, and anatomically informed convolution kernels (Operto et al., 2008) addressed the forward direction, how cortical geometry should shape the projection of voxel data onto the surface. Such methods have mostly been assessed with simulated or emulated data, in which the mismatch between the forward model and the data can be smaller than in an independent acquisition.

We examine three propositions, each in more than one dataset: 1. **Mixing can be predicted.** An acquisition-specific leakage map estimates, per vertex, the signed net contribution of surface-distant cortex to the predicted standard surface value. Its far part predicts how maps change under emulated coarsening and in real coarser sessions of the same subjects. 2. **Contamination predicts where correction pays.** The gain of inverting the operator over strong simple competitors grows with contamination, the leak weighted by how differently the mixed cortex responds: in HCP development data, in a pre-specified confirmatory cohort of 158 subjects in 86 families and in an independent 3T dataset. The leak alone predicts it less reliably. Estimated from the coarse data alone, contamination predicted the gain in HCP but not at 3T, where an anatomical atlas did at twofold coarsening. 3. **Correction helps in a definable regime.** The tested reconstruction performed best against noise-limited reference maps or under stronger coarsening, provided the anatomical registration was adequate. With many-run averages a well-tuned upsampling is as good except where differently responding cortex mixes strongly, and we found no robust gain across sessions.

We do not claim super-resolution. The claim is narrower: under an explicitly specified forward model, partial-volume mixing between surface-distant cortex can be estimated for a given acquisition and anatomy, and a regularised inversion of that model improves selected retinotopic comparisons where the mixing is high.

## 2. Theory

### 2.1 Forward model

Every surface map passes through two steps that mix cortex. The acquisition records each voxel as a blurred average of the tissue around it, and the mapping gives each vertex an average of the voxels around it. Because the cortex is folded, the tissue around a voxel can include cortex that is far away along the sheet: the opposite bank of a sulcus can lie a few millimetres away in the volume but much farther along the surface, down one bank and up the other (Fig. 1A, left). The forward model writes both steps as one linear operator. Given the acquisition parameters, a reconstructed cortical surface and a registration, it predicts the spatial sensitivity of each mapped value before the functional data are analysed, and it is the model that the estimator of Section ^7^2^2^.4 inverts.

**Figure 1.**
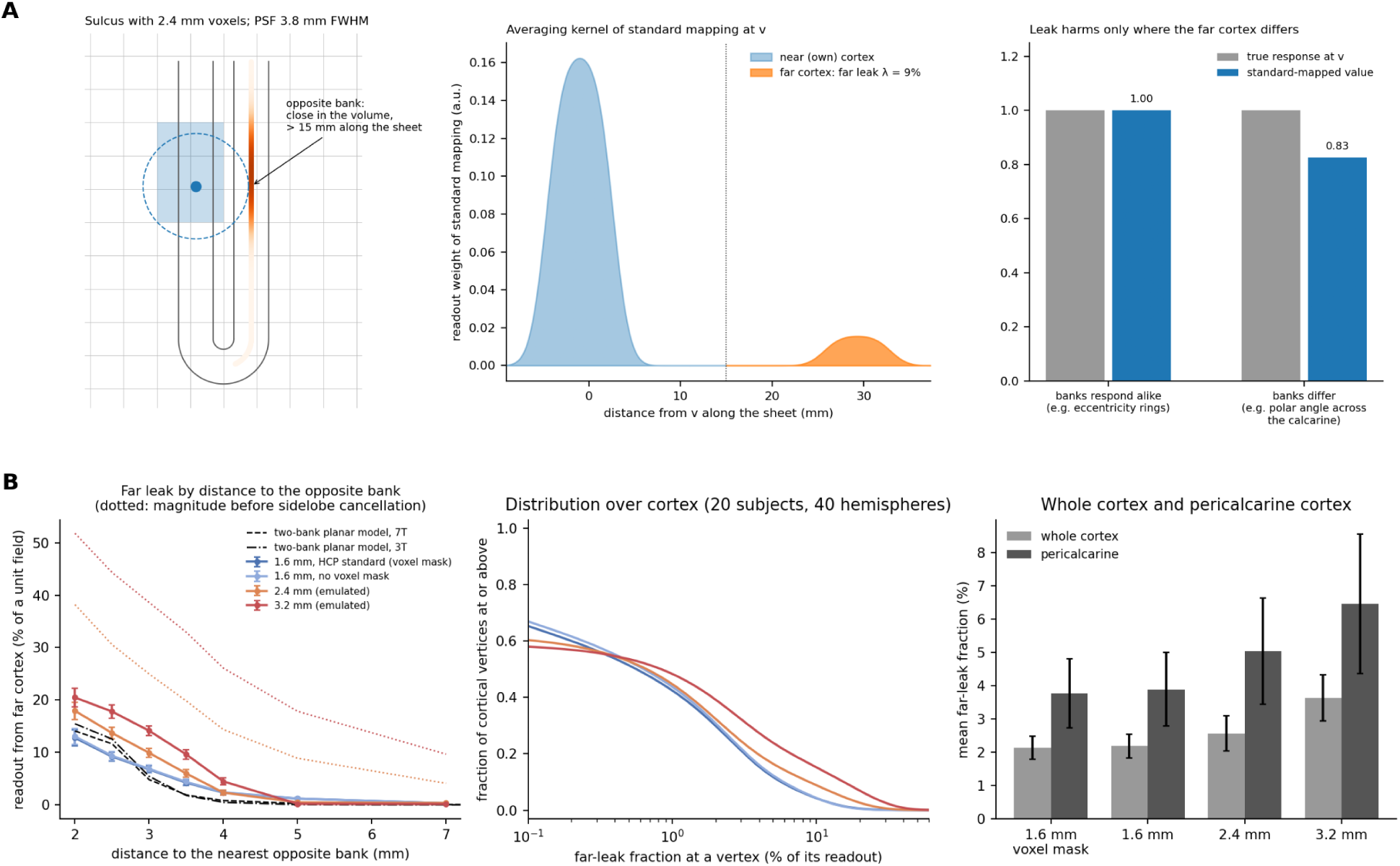
Partial-volume leakage of standard surface mapping. (A) Schematic, three panels. Left: a sulcus with a 2.4 mm grid; the voxels of standard mapping at vertex v (blue) and the foot- print of the effective PSF (dashed). Middle: the averaging kernel of standard mapping at v along the sheet. Its far part (orange; > 15 mm along the sheet) comes from the opposite bank, which is close in the volume. Right: the far part biases the mapped value only when the far cortex responds differently. (B) Real HCP geometry (20 development subjects, mean ± SD over 40 hemispheres), at 1.6 mm with and without the HCP voxel mask and emulated 2.4 / 3.2 mm: net far readout of a unit field against the distance to the opposite bank (left; dotted: the magnitude of the far readout before cancellation between positive and negative kernel lobes; black dashed and dash-dotted: a two-bank planar model at 7T and 3T), the distribution of the far-leak fraction over cortex (middle), and whole cortex against pericalcarine cortex (right).

We write f for a latent BOLD response parameterised on the cortical sheet: the response assigned to each point of cortex before the modelled acquisition blur and voxel sampling (here, the polar-angle response, a complex number whose phase gives the polar angle). In practice f has one value per surface vertex, the response of the column of cortex under it, taken to be the same at all depths between the white and pial surfaces. Because f is the BOLD response and not neural activ ity, the spread of the haemodynamic response along the cortex is part of f and is not undone.

The scanner does not record f directly. The acquired volume is y = K f + ε, where ε is noise and K is a matrix with one row per voxel and one column per vertex. For each vertex u, K describes how the response of the cortical column there is spread over the voxels: the column’s volume is integrated at five points between the white and pial surfaces, placed in the voxel grid through the subject’s registration, and spread by the effective point-spread function (PSF) of the acquisition. The effective PSF has four parts, each parameterised from the acquisition and the assumed pre processing rather than estimated from each subject’s functional signal (parameters: Supplementary S1, S2 and S15.1). Along the phase-encoding axis, the signal decays with T2* while the lines of k-space are acquired, which blurs that axis. In plane, k-space is sampled only up to the resolution limit and tapered at its edge, which gives a sinc-like kernel with small negative side lobes. Across slices, the slice profile has soft edges. On every axis, interpolation in pre-processing adds a Gaussian blur. Under this parameterisation the effective PSF is wider than the nominal voxel, so that each voxel receives contributions from tissue beyond its borders; for the emulated coarser grids it is the native PSF band-limited by the emulation window (Section 3.3).

Standard ribbon-constrained mapping then gives each vertex a weighted average of the voxels that intersect the cortical ribbon around it, each voxel weighted by its overlap with the ribbon there. We write it as a matrix W with one row per vertex and one column per voxel (the Connectome Workbench ribbon-constrained weights; Marcus et al., 2011; Glasser et al., 2013). The standard surface map is therefore z = W y = A f + W ε, with A = W K. The matrix A is the whole path from cortex to surface map, the acquisition followed by the mapping, and it is fixed by the acquisition and the anatomy alone. Its row a_v_, the averaging kernel of vertex v, lists the weight a_vu_ with which the response at each vertex u enters the value mapped at v (Fig. 1A, middle). An ideal mapping would put all of this weight at v. In practice the kernel differs from this ideal in three ways. It spreads over the cortex around v along the sheet, a loss of resolution much like surface smoothing. It also reaches cortex that the folding brings close to v in the volume, the subject of Section 2.2. Finally, within this cortex-only source model its weights sum to less than one, because the voxels at the edges of the ribbon also contain white matter and cerebrospinal fluid, which are not modelled as responding sources (the predicted attenuation, called dilution in Table 1). Some weights are negative, from the side lobes of the PSF.

**Table 1.**
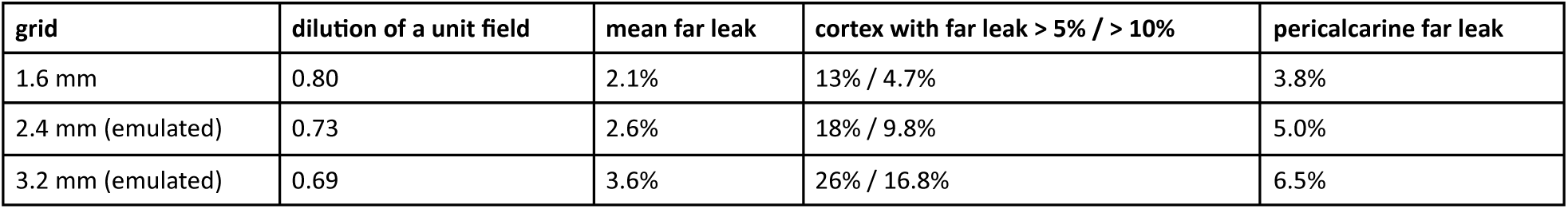
Leakage of standard ribbon mapping in HCP 7T geometry (20 subjects, individual maps).

The model is linear because acquisition, interpolation and mapping are all linear operations for a fixed registration. It assumes that the response is the same at all depths of a column, that the registration places the surfaces correctly in the volume and that the PSF follows from the acquisition parameters. Sections 4.3 and 4.4 give limited sensitivity analyses of the last two, an imposed 1 mm registration displacement and a 15% perturbation of the PSF width, and Section 4.2 shows how a narrower PSF changes the leakage figures.

### 2.2 Far leak

Not all mixing matters equally. Mixing with cortex near v along the sheet blurs the map but keeps its layout, as surface smoothing does, and analyses at a given resolution expect it. Mixing with cortex far from v along the sheet is different. In this geometric model, such cortex contributes to v only because the folding brings it close in three-dimensional space, and it can respond quite differently, as the two banks of the calcarine sulcus do for polar angle (one represents the upper visual field, the other the lower). This mixing does not just lower the resolution: it moves signal from one part of the map to another. Smoothing along the surface cannot remove it, because along the surface the two sources are not neighbours.

We call a point of cortex *far* from vertex v if it lies in the other hemisphere or more than 15 mm from v along the surface. Distance along the folded surface is approximated by distance on the subject’s spherical surface (FreeSurfer’s inflation of the cortex to a sphere), scaled so that its area equals that of the hemisphere’s cortex on the midthickness surface (radius √(S/4π) for area S). This proxy is least accurate where the inflation distorts the surface most; true geodesic distances give similar maps (Section 4.2). Writing F(v) for the set of far points of v, the far leak λ_v_ = Σ_u∈F(v)_ a_vu_ / Σ_u_ a_vu_ is the signed sum of the far-cortex kernel weights, normalised by the signed sum of all cortical kernel weights (Fig. 1A, middle). For a spatially uniform cortical response, λ_v_ is the ratio of the net far-cortex contribution to the net predicted cortical readout at v. It is a net, signed quantity: because the PSF has negative lobes, a far leak of 10% should not be read as a measured, non-negative fraction of physiological signal originating more than 15 mm away, but as the net weight the model assigns to far cortex after cancellation between positive and negative weights. Both sums run over the modelled cortical sources only, so λ_v_ is conditional on excluding non-cortical sources and does not depend on how much white matter or fluid the voxels contain. Summed without their signs, the far weights give a different, unsigned measure of mixing, which is larger (Fig. 1B; Section 4.1), a point we return to in Section 4.6.

The far leak depends only on the acquisition and the anatomy, so it can be computed from the anatomy, the registration (or a planned grid) and the acquisition parameters, before the functional data are analysed, and shown over the cortex as a leakage map (Section 4.2). It is high where the cortex folds tightly, so that the opposite bank of a sulcus or a neighbouring gyrus lies within a few millimetres in the volume (Fig. 1B), and it grows with the voxel size and the width of the PSF. Averaged over the cortex it is only a few percent, but it exceeds 10% in a sizeable part of the cortex, and a larger part as the grid coarsens (Table 1). In the acquisitions studied here it is almost entirely mixing within a hemisphere, across sulci and between neighbouring gyri; the other hemisphere contributes little (Section 4.2). The 15 mm threshold is a convention. It is more than twice the width (FWHM) of the widest PSF studied here (Section 3.3), so cortex counted as far contributes mainly where the folding brings it close in the volume; Section 4.2 shows that the figures change smoothly with the threshold.

### 2.3 Contamination-weighted leak

The signed far leak is a predicted net contribution, not the error of a functional map. Far mixing biases a mapped response relative to a local reference only when the far and local responses differ: if far cortex has the same response as v, mixing it in leaves the mapped value unchanged, however large λ_v_ is (Fig. 1A, right). The contamination-weighted leak (CWL) therefore weights the far leak by the response. Given an estimate x of the response at each vertex, we define the estimated far-source response contrast at v as c_v_ = Σ_u∈F(v)_ a_vu_ (x_u_ − x_v_) / Σ_u_ a_vu_, the far part of the mapped value minus what it would be if all far cortex responded like v. In other words, c_v_ is the far leak λ_v_ times the difference between the mean response of the far cortex, weighted by the kernel, and the response at v. Its magnitude depends jointly on the signed kernel and on the source– target response differences, and it too can be reduced by cancellation between contributions; it is large only where much of the kernel lies on far cortex and that cortex responds differently.

The same anatomy can therefore be contaminated for one stimulus and not for another. For polar angle, the two banks of the calcarine sulcus respond differently (Section 2.2), and far mixing across the sulcus shifts the mapped angle. For eccentricity rings, points facing each other across the sulcus represent the same eccentricity and respond alike, so mixing directly across these banks changes little, although more distant eccentricity representations can still differ (Fig. 1A, right). The CWL is the size of the error, ||c_v_|| (for a complex response, its magnitude), divided by the root-mean-square response over the scored vertices (Section 3.4). It is thus a fraction of the typical response: a CWL of 10% corresponds to a model-predicted far-source contrast whose magnitude is a tenth of the root-mean-square response. This makes the CWL comparable across subjects, stimuli and datasets, and lets the same bin edges (2, 5, 10, 20%) be used throughout.

The response estimate x can come from three sources, which trade accuracy against availability. CWL-M takes it from a native-resolution map of the source data, the finer data from which the coarse data were made. It is the most direct estimate, but it needs finer data of the same subject, which a study at the coarse resolution would not normally have. CWL-P takes it from the coarse data themselves, upsampled in k-space to the native grid and mapped as the native data are. It needs nothing beyond the coarse acquisition and the anatomy, but its response estimate is itself blurred and mixed. CWL-A takes it from an anatomical retinotopy atlas, the template of Benson et al. (2014; Benson & Winawer, 2018), version 4.0, carried to the subject’s surface through the subject’s registration to fsaverage: x = exp(iφ), with φ the template’s polar angle, inside its 12 visual areas and the stimulus radius (12.4°; Himmelberg et al., 2021), and x = 0 elsewhere. Given the anatomical reconstruction and its registration to the atlas, it needs no functional data of the participant, but it gives the typical layout of the maps, not the subject’s own.

In the analyses, vertices are grouped by CWL into bins, and the dose contrast D compares the gain of correction in the most contaminated bin with that in the least (Section 3.5). If response-weighted contamination is informative about the benefit of reconstruction, the model − competitor difference should become more negative across increasing contamination strata, so the prediction is a negative D, which Section 4.4 tests for each version (further versions: Supplementary S8). The CWL is meant to predict where correction pays, not to describe how the estimator achieves the gain; Section 4.6 examines what the gain consists of.

### 2.4 Estimator

To undo the mixing, we estimate f from the coarse data by inverting the forward model. The estimate is the posterior mean of f given the data, under a Gaussian noise model with a separate noise variance for each voxel and a prior that expects the response to vary smoothly along the cortical surface. The prior is a Matérn field (ν = 1) on each hemisphere, handled efficiently on the subject’s surface mesh through its link to a sparse Gaussian Markov random field (the SPDE approach; Lindgren et al., 2011). To compare the estimate f̂ with a finer target map, we pass it through the target acquisition’s own operator, W_t_ K_t_ f̂, so that it is mapped exactly as the target was. The prior has two settings, its spatial range and its amplitude, chosen on separate tuning subjects with the same rule as every competitor’s settings (Section 3.4).

## 3. Methods

### 3.1 Data

We used four public datasets. The HCP 7T retinotopy data (Benson et al., 2018) provided 20 subjects for development, split by family in advance into 5 tuning and 15 development subjects, and 158 further subjects in 86 families as a confirmatory cohort, with polar-angle wedges and, in the cohort, eccentricity rings at 7T 1.6 mm and 2.4 / 3.2 mm emulated in k-space; they were used for operator validation, emulation, the leakage maps, the dose-response and the confirmatory tests. StudyForrest multi-resolution data (Sengupta et al., 2017a, 2017b, 2018) provided 7 subjects with 7T sessions at 0.8 / 1.4 / 2.0 / 3.0 mm, 5 of them also with 3T sessions at 1.4 / 2.0 / 3.0 mm, and a hemifield orientation stimulus, for real coarser sessions, within-session emulation and a cross-session test. OpenNeuro ds002702 (Kay et al., 2019, 2020a, 2020b; Markiewicz et al., 2021) provided 5 subjects with six eccentricity rings at 7T GE 0.8 mm, two of them also at 3T GE 2.4 mm and two at 7T SE 1.05 mm, for within-session emulation and real second acquisitions. The NYU Retinotopy Dataset (OpenNeuro ds003787; Himmelberg et al., 2021) provided 44 subjects (8 tuning, 36 test; 43 included) with sweeping bars at 3T 2.0 mm (multiband 6) and 3.0 / 4.0 mm emulated in k-space, as an independent pre-specified replication. HCP restricted family information was used only to form family-wise splits and to pick one subject per family for the cohort’s primary tests; no restricted information is reported. ds002702 and ds003787 were processed with the authors’ FreeSurfer reconstructions (Fischl, 2012).

This study is a secondary analysis of four public datasets; no participants were newly recruited or scanned. In each, participants gave written informed consent, and data collection was approved by the responsible ethics board: the Washington University in St. Louis Institutional Review Board (HCP), the Ethics Committee of the Otto-von-Guericke University Magdeburg (StudyForrest), the University of Minnesota Institutional Review Board (ds002702) and the New York University committee on activities involving human participants (NYU Retinotopy Dataset). StudyForrest and NYU participants also consented to the public release of their anonymised data. HCP data were used under the WU-Minn HCP Open Access and Restricted Data Use Terms; as the latter require, HCP subject identifiers are not reported.

### 3.2 Preprocessing and response maps

For HCP, we used the released 7T retinotopy data (Benson et al., 2018) as preprocessed by the HCP minimal preprocessing pipelines (Glasser et al., 2013): corrected for gradient nonlinearity, head motion and EPI distortion, registered to the subject’s anatomy and resampled to MNINonLinear space at 1.6 mm (TR 1 s), then denoised with ICA-FIX (Salimi-Khorshidi et al., 2014; Griffanti et al., 2014). From the concatenated FIX-cleaned series we took the two rotating-wedge runs, counter-clockwise and clockwise, of 300 volumes each (22 s blank, then 8 cycles of 32 s). Each run was mapped to the subject’s native mesh with the HCP pipelines’ ribbon-constrained weights and that run’s goodvoxels mask. The response at each vertex is the complex Fourier component at the stimulus frequency, whose phase encodes polar angle; its noise variance is the mean power at the eight neighbouring frequency bins, and coherent vertices are those with coherence > 0.3 in the 1.6 mm maps. In the confirmatory cohort the expanding- and contracting-ring runs were analysed in the same way, with their own goodvoxels masks.

Opposite stimulus directions. The counter-clockwise and clockwise wedge runs, and the expanding and contracting ring runs, sweep the same positions in opposite temporal order, so that, up to the haemodynamic phase delay δ_v, the clockwise response is the complex conjugate of the counter-clockwise one: exp(i(θ_v + δ_v)) against exp(i(−θ_v + δ_v)), with θ_v the polar angle (for rings, the eccentricity phase). No temporal reversal or delay correction is applied to the time series. Every method, the model included, is applied to each run separately; in the cross-run score (Section 3.3) the prediction from one run is conjugated before it is compared with the other run’s map, the single global complex scale factor of the score (Section 3.4) absorbs the common delay and amplitude, and residuals are computed on complex values, so no phase unwrapping is involved. For polar-angle maps in degrees (Fig. 2; Section 4.3), the angle is half the difference of the two runs’ phases and the delay half their sum, resolved modulo π per vertex to the branch nearest a global delay estimate (coherence-weighted over pericalcarine cortex at 1.6 mm: 0.80 ± 0.17 rad over the 20 development subjects). As a check of the convention, one 1.6 mm wedge map predicts the other after conjugation with a residual of 0.362 (0.305 after 6 mm smoothing; Section 4.3), and the maps show the upper and lower visual field on opposite banks of the calcarine sulcus (Fig. 2). The GLM-based maps of the other three datasets involve no sweep direction.

**Figure 2.**
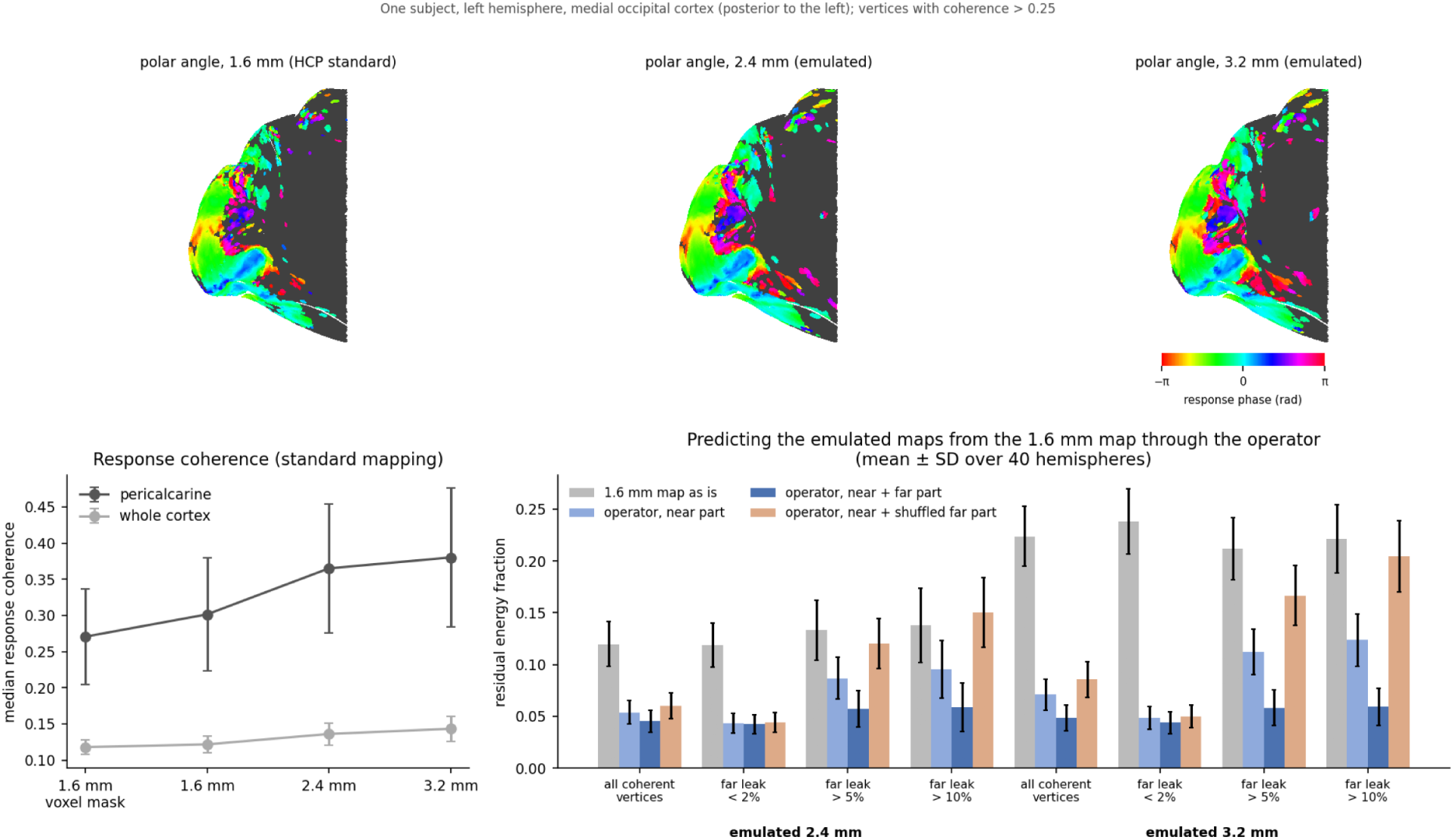
The operator predicts how the maps change under emulated coarsening. Top: polar-angle maps (response phase) of one subject’s medial occipital cortex at 1.6 mm and emulated 2.4 / 3.2 mm (vertices with coherence > 0.25 shown; the analyses use > 0.3). Bottom left: response coherence by grid (median over the vertices of a hemisphere; mean ± SD over 40 hemispheres). Bottom right: residual of predicting the emulated maps from the 1.6 mm field estimate through the operator, by vertex stratum (mean ± SD over 40 hemispheres). Predictors: the 1.6 mm map itself (grey), the operator’s near part only, near + far part, and near part + far part shuffled across vertices.

The StudyForrest 7T sessions at 0.8, 1.4, 2.0 and 3.0 mm were distributed with distortion correction applied by the data providers; the 3T sessions have no field maps. The runs of each session were motion-corrected to its first run (MCFLIRT; Jenkinson et al., 2002), and the session mean was registered to the subject’s FreeSurfer anatomy with boundary-based registration (bbregister; Greve & Fischl, 2009). A GLM per run, with a canonical HRF and its temporal derivative, Legendre drifts and the motion parameters, gave betas in percent signal change for the left- and right-hemifield conditions, which we combined into one complex map (left + i·right).

In ds002702, each session (7T gradient echo at 0.8 mm; 3T gradient echo at 2.4 mm; 7T spin echo at 1.05 mm) was motion-corrected to its first run (MCFLIRT), and the session mean was registered to the authors’ FreeSurfer anatomy with FLIRT from three starting positions followed by boundary-based registration, on canonically reoriented copies. Registration was checked by the EPI grey–white contrast across the white surface against copies shifted by 2 mm. Field-map correction (FSL fugue; Jezzard & Balaban, 1995; Jenkinson et al., 2012), with the field map closest to run 1 and the unwarping direction chosen per session by the registration cost, was kept only where it improved this check. A per-run GLM as for StudyForrest gave six real maps, one per eccentricity ring; the GLMs ran sequentially, and every session passed a per-run consistency check (Supplementary S9).

The NYU data were acquired at 3T (Siemens Prisma) at 2.0 mm isotropic, TR 1 s and multiband factor 6, with sweeping bars in 4–12 runs of 192 s and spin-echo field maps with opposite phase encoding. The field was estimated from the spin-echo pair with topup (Andersson et al., 2003). Every run was motion-corrected (MCFLIRT) to the first run’s single-band reference, itself registered rigidly to the spin-echo image (for three subjects without a single-band reference, the mean of the first run’s first 10 volumes), and distortion-corrected with applytopup with Jacobian modulation. The corrected session mean was registered to the FreeSurfer anatomy with FLIRT from three starting positions followed by boundary-based registration, keeping the lowest cost. Runs whose maximum translation exceeded 4 mm were excluded, and so were subjects left with fewer than 4 runs or a registration cost above 0.8; this removed one test subject. Per run, a GLM with the bar aperture’s coverage of the visual field weighted by 1, cos θ and sin θ (each convolved with the HRF), the temporal derivative of the first, Legendre drifts of order 0–3 and the six motion parameters gave the complex map β_cos_ + i·β_sin_ in percent signal change. This is the first angular harmonic of each voxel’s response, and its phase estimates polar angle as the wedge’s Fourier phase does in HCP; the noise variance came from the GLM residuals. Odd and even runs form two halves, and responsive vertices are those where the stimulus regressors explain R² ≥ 0.2 of the run-averaged native series.

Acquisition parameters used by the forward model are listed in Supplementary S1.

### 3.3 Operators, emulation and leakage maps

The operator’s kernels are tabulated per voxel axis from the acquisition parameters, and each cortical column is integrated at quadrature points between the white and pial surfaces (checked against brute-force integration: maximum relative error ≈ 10⁻⁷; column mass 0.99–1.01). The effective PSF FWHM at HCP 1.6 mm is 2.56 / 2.70 / 2.55 mm (x / y / z).

Coarser acquisitions are emulated by cropping k-space and applying a Tukey window (α = 0.5) that is zero at the coarse Nyquist frequency, frame by frame on the time series: HCP 1.6 → 2.4 / 3.2 mm, NYU 2.0 → 3.0 / 4.0 mm (×1.5, ×2). The coarse window itself is exact per axis and the real noise and physiology are kept, but this is an emulation of already reconstructed data: the window is a modelling choice, the native PSF and the preprocessing history are only approximated, the emulated PSF is 1.6–1.7 voxels wide at every grid, and the noise is band-limited together with the signal.

In HCP, each coarsened wedge run predicts the other run’s 1.6 mm map. In StudyForrest, ds002702 and NYU, odd and even runs form two halves, the coarsened half predicts the other half’s native map, and both directions are averaged. This cross-run design keeps the noise of the prediction and of the reference independent, but an fMRI reference map is not a ground truth for the latent cortical response: it carries its own noise, physiological variability and registration error, and agreement with it is what every score below measures.

Leakage maps give the model-predicted signed far leak of each subject at the native and emulated grids. HCP group maps were made on fs_LR 32k through each subject’s MSMAll sphere (Robin-son et al., 2014, 2018) and averaged over 19 subjects; NYU group maps were made on fsaverage.

### 3.4 Competitors, scoring and settings

All competitors received the same data and the same setting-selection rule as the model: standard mapping of the coarse data ± surface smoothing (FWHM 2–16 mm); k-space upsampling to the target grid, then the target’s mapping, ± smoothing; across sessions, resampling to the target grid ± smoothing. The score is the residual energy fraction of the finer reference map after one global (complex, for polar-angle maps) scale factor has been fitted to the prediction; lower values mean closer agreement with that reference, not necessarily greater accuracy relative to the unobserved cortical response. Vertex sets: coherent (HCP: response coherence > 0.3 in the 1.6 mm maps) or responsive (NYU: run-averaged native R² ≥ 0.2 for the stimulus regressors), visual labels (Study-Forrest, ds002702; Desikan et al., 2006), and strata by far leak or CWL at the coarse grid among vertices clean at the native grid (far leak < 2%). The coherent and responsive sets are defined from the native-resolution data of both runs (HCP: the mean coherence of the two wedge runs) or of all runs (NYU), so they depend on the reference as well as on the source data; they are the same for every method and depend on no method’s output, but they are not source-only masks. The strata use anatomy alone (far leak), the source run’s own native map (CWL-M), the coarse source data (CWL-P) or the atlas (CWL-A). Two comparators appear below: the competitor fixed on the tuning subjects, which is the pre-specified comparator of the confirmatory and NYU tests, and, in the cross-dataset summary of Section 4.5, the better of the two smoothing competitors for each subject, chosen on the test outcome itself; the latter is a demanding but retrospective comparator, not a prospective deployment comparison. Settings were chosen with one rule for every method: fixed on separate tuning subjects (HCP 5, NYU 8), leave-one-subject-out (StudyForrest and ds002702 within a session), or on the other subject of a pair type (real second acquisitions). For ds002702 and NYU the search grids were widened once, before any held-out result was summarised, because the first fits had their optimum at the grid’s edge (Supplementary S3, S16).

### 3.5 Pre-specification and statistics

The unit of analysis is the subject. The dose contrast D = gain in the highest bin with ≥ 30 vertices − gain in the 0–2% bin, with gain = residual(model) − residual(competitor), is tested with a one-sided Wilcoxon signed-rank test, with a two-sided bootstrap 95% interval of the mean (10,000 resamples). The tests, bins, vertex sets and settings reported below were fixed in analysis plans written before the corresponding analyses were run, with their SHA-1 and SHA-256 sums recorded; the plans are reproduced verbatim in Supplementary S11. This is pre-specification at-tested by the author, not a registration with a third party. The first contamination analyses were planned after the development-set results of Sections 4.1–4.3 were known, so their development-set tests are pre-specified within an explored dataset.

In the confirmatory cohort (HCP, 158 subjects in 86 families), the primary tests used one subject per family (n = 86), and all 158 subjects with a family bootstrap formed a secondary analysis. H1: the contamination dose-response replicates (D < 0 at both grids). H2: for rings, contamination predicts the gain, and more sharply than the plain leak. H3: with half the source data the gain where no far unmixing is needed shrinks (a denoising gain would grow). H4: the overall gain survives a PSF assumed 15% too wide. The plan fixed eight primary one-sided tests (P1–P8 in Table 4) with no correction across them and Holm correction within each family of secondary tests; Holm- and Bonferroni-adjusted p-values across the eight primary tests are reported in Table 4 and Supplementary S13.7 as a post hoc addition. H2 and H3 were formulated after an exploratory look at the 15 development subjects; an interim look at half of the batches changed nothing but a labelled sensitivity set.

The external replication (NYU, 35 included test subjects) tested three prospective hypotheses, fixed before any NYU fit. N1: the model beats the competitor chosen on the tuning subjects over responsive vertices; N2: D_far_ < 0; N3: D_CWL-M_ < 0; at both grids, Holm across the six tests. Two retrospective plans followed: a second plan, written after N1–N3 were known, added the coarse-data rule and the registration analysis of Section 4.5 and the 3T group maps; a third added contamination from an anatomical atlas (Section 4.4; two tests, Holm across them).

## 4. Results

### 4.1 The leakage map predicts mixing

Standard ribbon mapping on real HCP geometry (20 subjects, individual maps; Table 1):

Leak rises steeply as the opposite bank of a sulcus comes close (Fig. 1B): at 1.6 mm it is 12.8% where the opposite bank is within 2.25 mm, and 0.3% beyond 6 mm.

The far leak is a signed, net quantity (Section 2.2). Summed without their signs, the far weights of standard mapping average 3.06%, 9.93% and 18.55% of a unit field at 1.6, 2.4 and 3.2 mm, against net far weights of 1.70%, 1.95% and 2.73% (the far leak of Table 1 divides the net weight by the cortical row mass), so most of the unsigned far weight cancels between the positive and negative lobes of the PSF, and more so at the coarser grids (Fig. 1B, dotted); where the opposite bank lies within 2.25 mm the unsigned far weight is 19.7%, 38.2% and 51.8% against a net 12.8%, 17.9% and 20.5%. The denominator of the far leak, the cortical row mass, averages 0.80, 0.73 and 0.69 (Table 1, dilution; standard deviation over the 40 hemispheres 0.01–0.02). Per-vertex distributions of the row mass and the share of vertices with a negative net far leak were not compiled (Section 5).

The predicted mixing is geometrically right. Predicting the emulated 2.4 / 3.2 mm HCP maps from the 1.6 mm field estimate, adding the operator’s far part to its near part lowered the residual in **all 20 subjects (40 of 40 hemispheres) at both grids**; where the predicted leak exceeds 10%, from 0.095 to 0.059 at 2.4 mm and from 0.124 to 0.059 at 3.2 mm, whereas a far part shuffled across vertices made it worse in every subject (0.150 and 0.204) (Fig. 2). Because emulated maps and prediction share the emulation window, the same test was run on real coarser acquisitions: StudyForrest has 7T sessions of the same subjects at 0.8, 1.4, 2.0 and 3.0 mm. Through each coarser session’s own operator, adding the far part lowered the residual where the predicted leak exceeds 10% from 0.378 to 0.345 (7 of 7 subjects; primary pair 1.4 → 3.0 mm: 0.344 → 0.303); a shuffled far part made it worse (0.426, 7/7), and a near part renormalised to the full row mass did not help (0.373, 7/7) (Fig. 3B).

**Figure 3.**
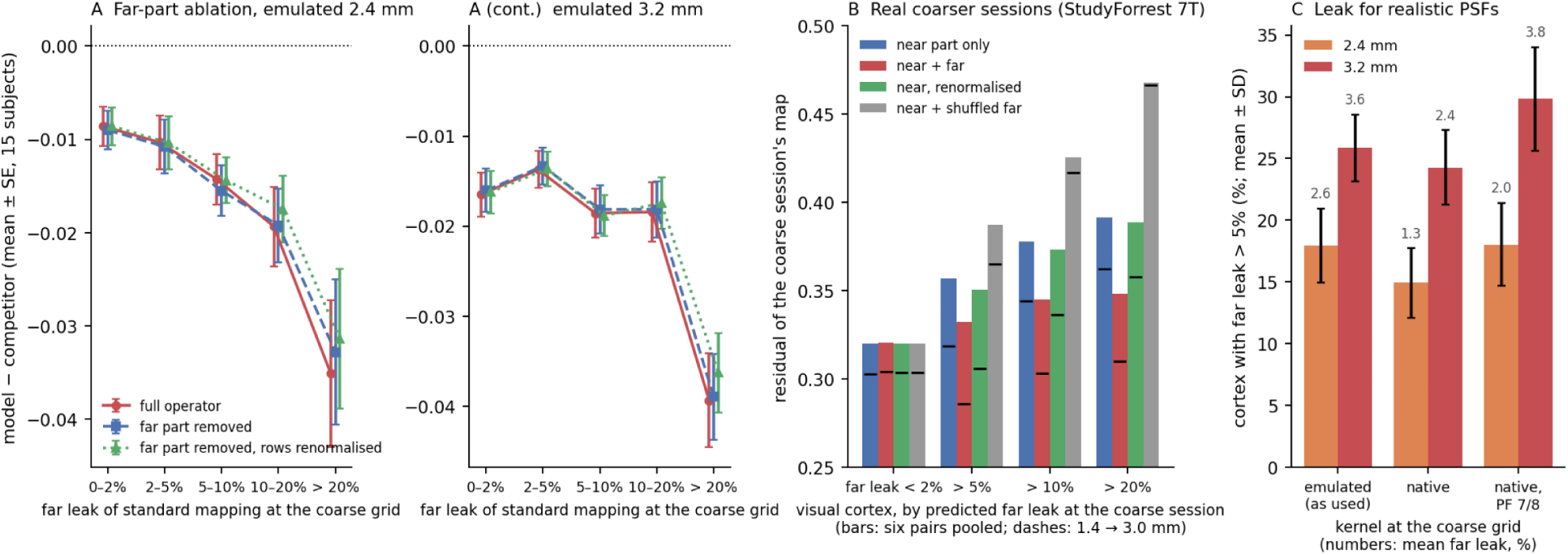
Three further tests. (A) Far-part ablation: the model’s gain over upsampling + 8 mm smoothing by far-leak bin (15 development subjects, mean ± SE) with the full operator, with the far entries of the operator removed per voxel, and with the far entries removed and the rows renormalised, at emulated 2.4 mm (left) and 3.2 mm (right). (B) Real coarser sessions (StudyForrest 7T, 7 subjects): residual of the coarser session’s standard map predicted through its operator from the finer session’s field estimate, with the near part only, near + far, the near part renormalised and near + shuffled far part, by the predicted far leak at the coarser session in visual cortex; bars pool the six session pairs per subject, dashes mark the 1.4 → 3.0 mm pair. (C) Share of cortex with far leak > 5% (mean ± SD over 20 development subjects; numbers: mean far leak in %) at 2.4 and 3.2 mm for the emulated kernel used throughout, a native-acquisition kernel at that voxel size, and the same with partial Fourier 7/8.

### 4.2 Leakage maps

The HCP group-mean signed far leak is 2.0–2.1%, 2.3–2.4% and 3.1–3.3% at 1.6, 2.4 and 3.2 mm; leak above 5% covers 4%, 10% and 22% of cortex (Fig. 4). Group maps are lower than individual maps: hot spots do not align perfectly across subjects, and resampling to an average surface changes the area weights. The highest leak at 3.2 mm, as area-weighted mean far leak (share of the region above 10%), is in the posterior cingulate (8.7%, 40%), cuneus (8.4%, 35%), banks of the superior temporal sulcus (7.0%, 33%), transverse temporal gyrus (6.5%, 32%) and the pericalcarine and lingual cortex (6.3% and 6.2%, 27% each); the central sulcus appears as a band on both banks, and the frontal and temporal poles and orbitofrontal cortex are lowest (< 1.5%).

**Figure 4.**
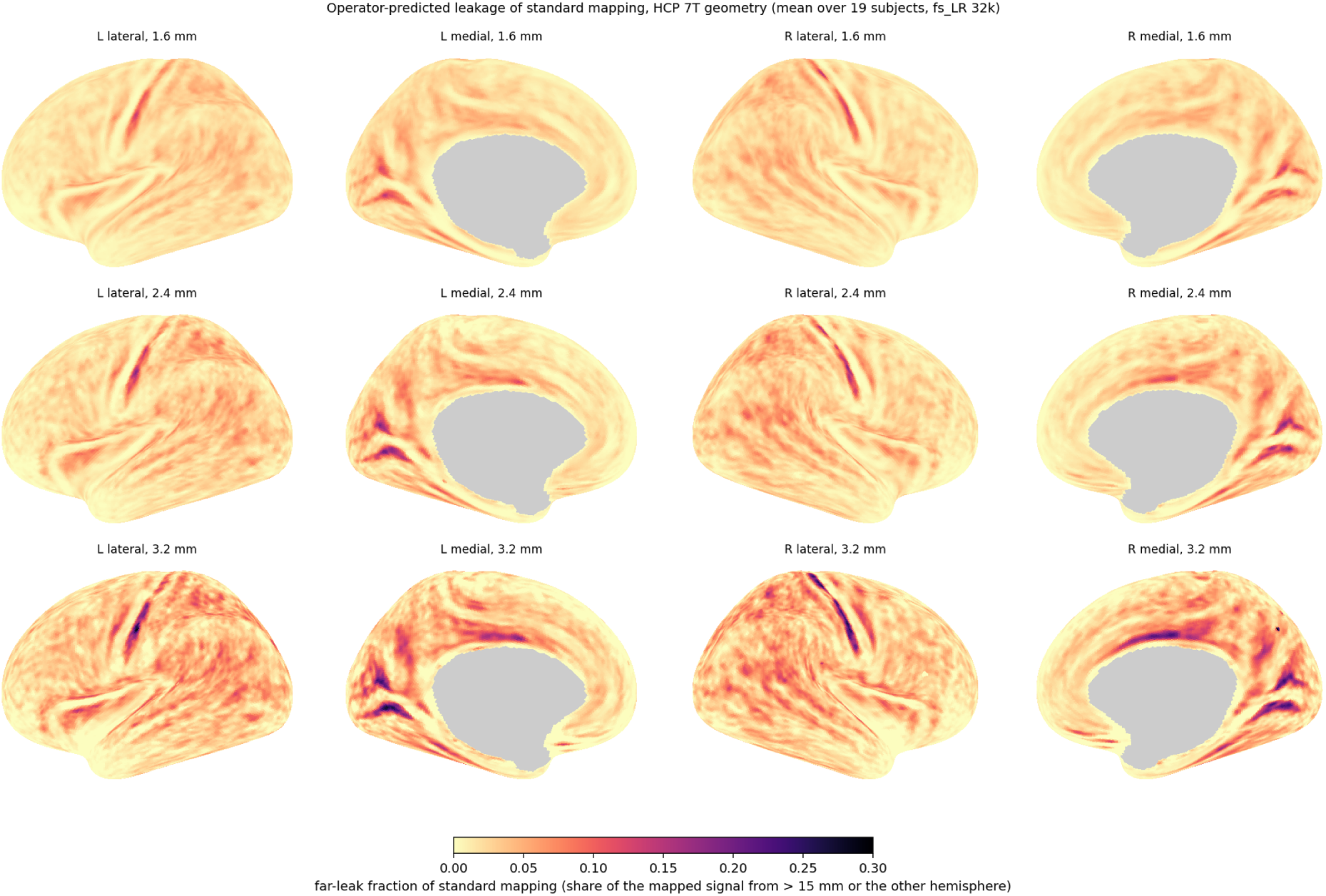
Group maps of operator-predicted far leak of standard mapping. HCP 7T geometry, mean over 19 of the 20 development subjects (one lacked an MSMAll sphere) on fs_LR 32k, at 1.6 mm (top) and emulated 2.4 and 3.2 mm (Tukey-windowed k-space crop; effective PSF 3.8– 4.1 and 5.1–5.4 mm); lateral and medial views; medial wall in grey.

A typical 3T acquisition leaks as much. For the NYU 2.0 mm acquisition the mean far leak is 3.2% and 19% of cortex lies above 5% (43 subjects, individual maps), and at the 3.0 and 4.0 mm emulations 4.2% and 27%, 5.0% and 33%. On fsaverage the 43-subject group map has a mean far leak of 3.0%, 3.8% and 4.0% at 2.0, 3.0 and 4.0 mm and exceeds 5% in 17%, 28% and 33% of cortex (Fig. S2). Its means are lower than in subject space because the average surface weights vertices differently; each subject’s own map, resampled, drops by as much. The same regions lead as in HCP: at 2.0 mm the cuneus (7.7%), pericalcarine cortex (7.5%), transverse temporal gyrus (6.4%), lingual gyrus (6.1%), banks of the superior temporal sulcus and isthmus and posterior cingulate (5.5–5.6%). Both sets of group maps are released.

The figures are robust in pattern and uncertain in scale. Native-acquisition kernels without the emulation window give similar or lower figures (net far leak 1.3–2.0% at 2.4 mm and 2.4–3.8% at 3.2 mm against 2.6% and 3.6%, the higher values with partial Fourier 7/8) (Fig. 3C). A kernel 15% narrower, the width the inversion preferred in a scan of the PSF model, lowers them by about a fifth (2.0% and 2.8%). With the far threshold at 10 or 20 mm they scale smoothly (4.4 / 2.6 / 1.6% at 2.4 mm), and the other hemisphere alone contributes 0.2–0.35%: the far leak of these acquisitions is same-hemisphere mixing across sulci and between neighbouring gyri. True geodesic distances along the midthickness surface give maps that agree with the sphere version (per-vertex r = 0.86–0.90), which slightly overstates the net far leak at the coarse grids (2.8% against 2.6% at 2.4 mm, 4.0% against 3.6% at 3.2 mm, 5 of 5 subjects) (Supplementary S14.2, S15).

### 4.3 Correction in HCP emulation

Table 2 gives the cross-run residual against the other run’s 1.6 mm map for the 15 development subjects, with settings fixed on the five tuning subjects:

**Table 2.**
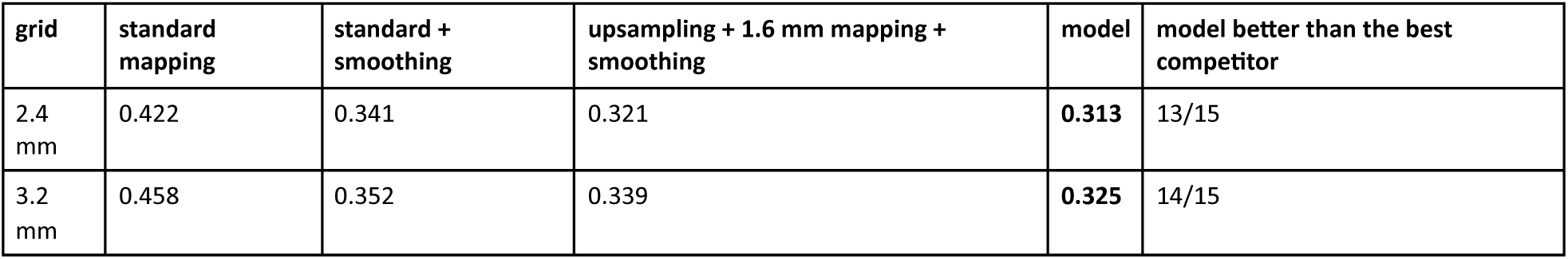
Cross-run residual in HCP emulation (15 development subjects; lower is better).

The target is a single run (Fig. 5A): the other 1.6 mm run itself, smoothed 6 mm, scores 0.305. Against this ceiling the model closes 51% (2.4 mm) and 41% (3.2 mm) of the gap left by upsampling + smoothing (from the unrounded means, 0.3212, 0.3131 and 0.3052 at 2.4 mm; the rounded values of Table 2 give 50%; per-subject medians 46% and 40%), and reaches it where leak exceeds 10% at 2.4 mm (0.343 against 0.344). In polar angle, the model’s advantage is 0.5–0.6° over all coherent vertices (13–14 of 15 subjects) and 1.9–2.1° where mixing is strong (> 20% far leak), against a single-run noise floor of about 30°.

**Figure 5.**
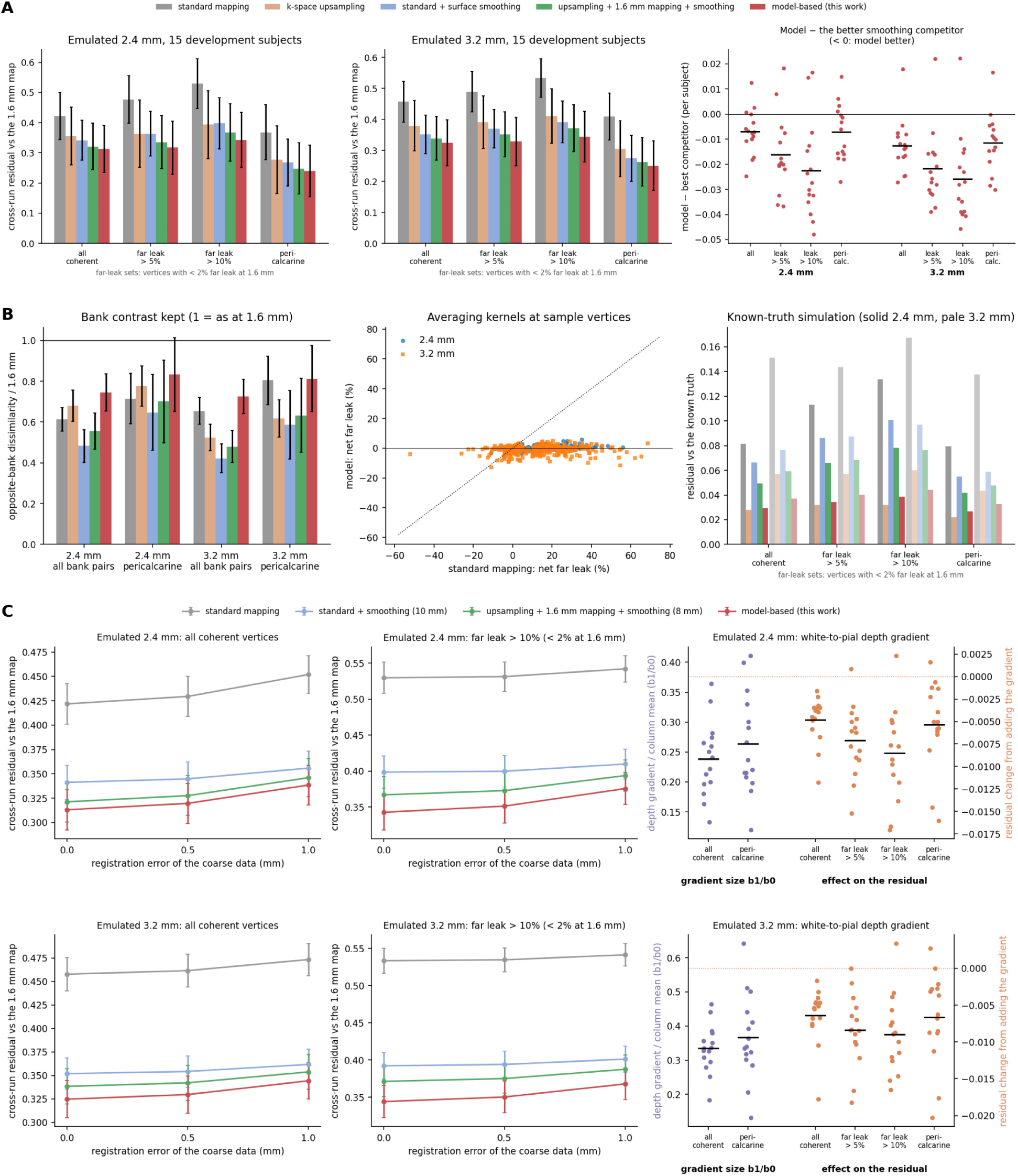
Model-based estimation in HCP emulation (15 development subjects). (A) Cross-run residual against the other run’s 1.6 mm map for every method, by vertex set (mean ± SD over subjects); the far-leak sets hold only vertices with < 2% far leak at 1.6 mm. Right: per-subject difference between the model and the better smoothing competitor. (B) Polar-angle contrast kept across sulcal banks (left), net far leak of the model’s averaging kernels against standard mapping’s (middle; 360 sample vertices per grid), and a known-truth simulation (right; solid bars 2.4 mm, pale bars 3.2 mm); colours as in A. (C) Robustness to a translational misregistration given to every method (left, middle; mean ± SE over the 15 subjects), and the depth-gradient component (right): its size relative to the column mean (b1/b0, purple) and the change in residual when it is added (orange).

Where unmixing is needed (clean at 1.6 mm, leak > 10%), the lead grows to −0.024 (2.4 mm) and −0.027 (3.2 mm), 14/15 subjects each, and the model keeps more of the polar-angle contrast across sulcal banks (dissimilarity 0.75 against 0.61 for standard mapping and 0.68 for upsampling at 2.4 mm) (Fig. 5B).

The lead survived two imposed perturbations: a 1 mm translational misregistration applied identically to every method (−0.008 / −0.009, 14/15) (Fig. 5C), which tests a common displacement and not method-specific registration error, and, in the confirmatory cohort, a PSF assumed 15% too wide (63 of 86 families; Section 4.4). A scan of the kernel width puts the optimum slightly narrower than modelled (α = 0.85 lowers the residual in 5 of 5 tuning subjects, by 0.002–0.003) without changing the dose-response. A stationary correlated-noise likelihood, its spectrum estimated from neighbouring frequencies, made every development subject worse (+0.054 and +0.026), also with its own prior re-tuned. With an exact operator and little noise (a known-truth simulation), band-limited upsampling is near-exact at 2.4 mm and edges out the model there, while the model wins clearly at 3.2 mm (0.037 against 0.057). Details: Supplementary S5, S15, S17.

### 4.4 Contamination predicts where correction pays

In the HCP development data, the model’s gain over the best competitor rises with the far leak of the coarse map, from −0.007 (clean bin) to −0.035 above 20% at 2.4 mm and from −0.016 to −0.038 at 3.2 mm (Fig. 6A), and with the contamination-weighted leak (Fig. 6B).

**Figure 6.**
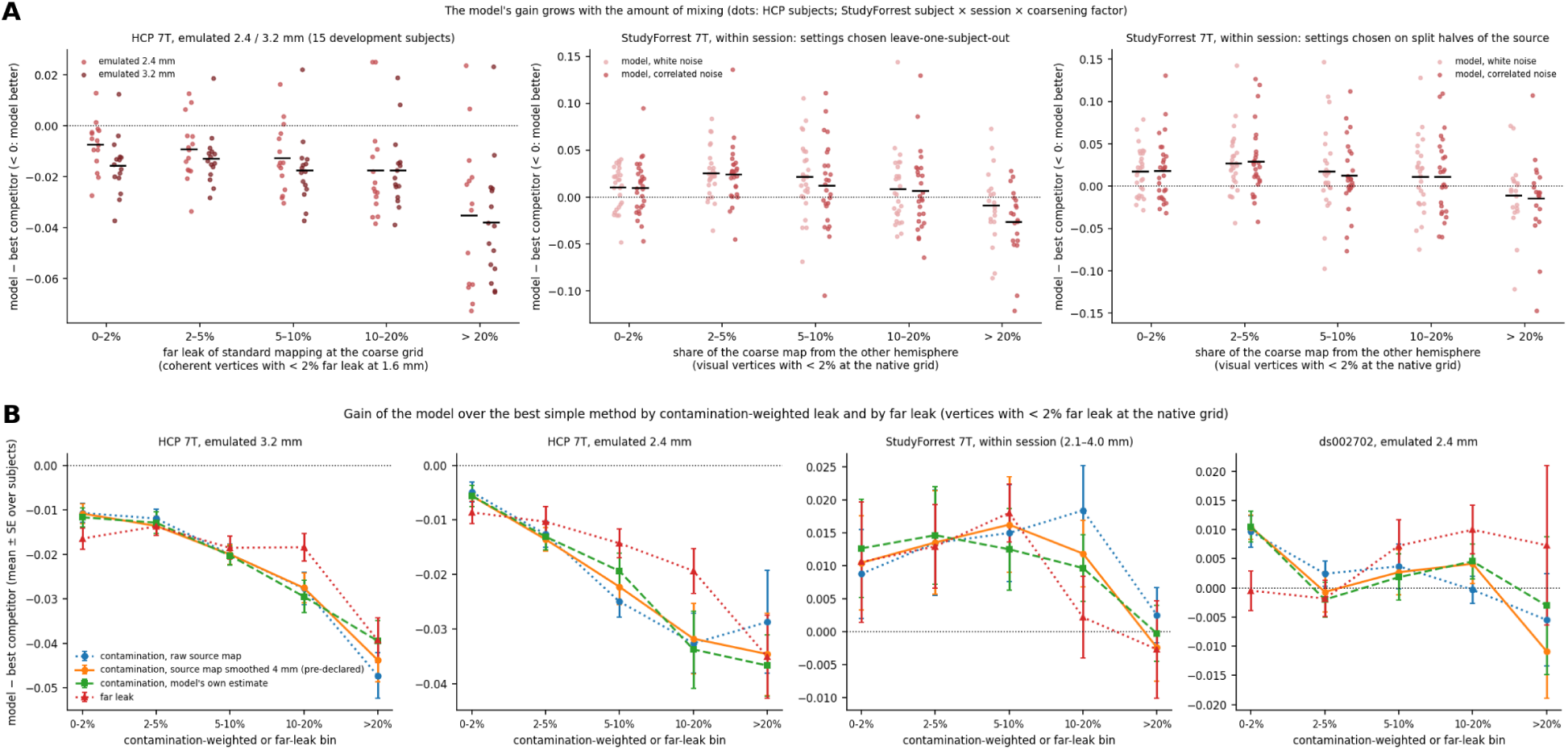
The model’s gain grows with mixing of differently responding cortex. (A) Model − best competitor by far leak in HCP (left), and by the other-hemisphere share in StudyForrest within session under two setting rules (middle: leave-one-subject-out; right: split halves of the source). Dots are subjects (HCP) or subject × session × coarsening-factor cells (StudyForrest; 20 of 28 cells reach the > 20% bin). (B) Model − best competitor by contamination-weighted leak, with the raw, smoothed (4 mm, pre-declared) and model-based response estimates, and by far leak, for HCP 3.2 / 2.4 mm, StudyForrest and ds002702 2.4 mm (mean ± SE over subjects).

A pre-declared test with the contamination-weighted leak gave the dose contrasts in Table 3:

**Table 3.**
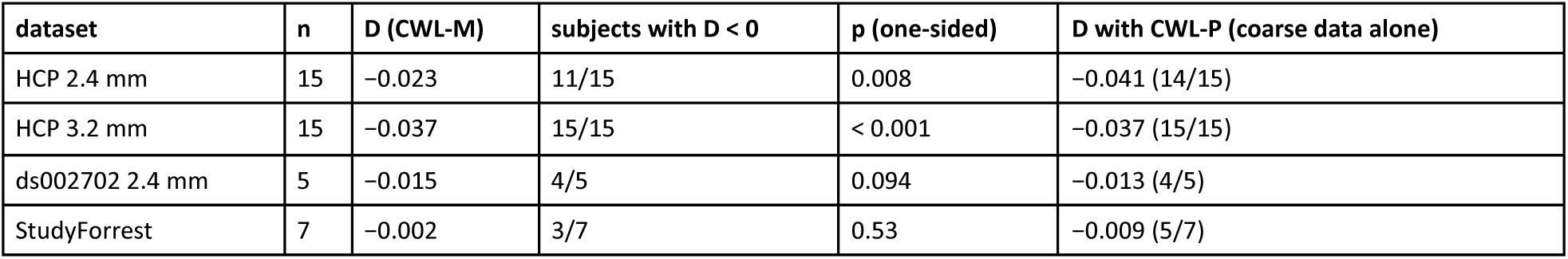
Contamination and the model’s gain: pre-declared dose contrasts in the development data.

| **dataset** | **n** | **D (CWL-M)** | **subjects with D < 0** | **p (one-sided)** | **D with CWL-P (coarse data alone)** |
|---|---|---|---|---|---|
| HCP 2.4 mm | 15 | −0.023 | 11/15 | 0.008 | −0.041 (14/15) |
| HCP 3.2 mm | 15 | −0.037 | 15/15 | < 0.001 | −0.037 (15/15) |
| ds002702 2.4 mm | 5 | −0.015 | 4/5 | 0.094 | −0.013 (4/5) |
| StudyForrest | 7 | −0.002 | 3/7 | 0.53 | −0.009 (5/7) |

The coarse-data-only measure predicts the HCP gain as well as the one from fine data. In the resolution-limited datasets the evidence stays suggestive: a denoised response estimate moved both in the predicted direction (ds002702 −0.021, p = 0.062; StudyForrest −0.010, p = 0.19), and in StudyForrest the share of the other hemisphere, where hemifield stimuli differ, predicts the gain (−0.019, 6/7; exploratory).

In the confirmatory cohort (158 HCP subjects in 86 families, all of whom passed the pipeline’s checks; primary tests on one subject per family), the pre-specified tests gave the results in Table 4 (full table: Supplementary S13; the eight primary tests with adjusted p-values: Supplementary S13.7). Roles are as the plan fixed them: primary (P1–P8), secondary (Holm within each hypothesis’ family) or descriptive; the last column adjusts the primary tests across their family of eight, post hoc:

**Table 4.** Confirmatory cohort: pre-specified tests (158 HCP subjects; primary analyses on one subject per family, n = 86).

| hypothesis | role | test | grid | n | mean | in direction | bootstrap 95% | p | p (Holm, P1–P8) |
|---|---|---|---|---|---|---|---|---|---|
| H1 | P1, P2 | D, contamination (CWL-M) | 2.4 / 3.2 mm | 86 | −0.0264 / −0.0219 | 64/86, 77/86 | [−0.0332, −0.0197], [−0.0262, −0.0177] | < 0.001 | < 0.001 |
| H1 | secondary | D, contamination (CWL-P, coarse data only) | 2.4 / 3.2 mm | 86 | −0.0339 / −0.0211 | 70/86, 71/86 | – | < 0.001 | – |
| H1 | secondary | D, plain far leak | 2.4 / 3.2 mm | 86 | −0.0201 / −0.0049 | 57/86, 48/86 | – | < 0.001, 0.015 | – |
| H1 | secondary | overall gain, all coherent vertices | 2.4 / 3.2 mm | 86 | −0.0087 / −0.0128 | 82/86, 85/86 | – | < 0.001 | – |
| H2a | P3 | D<sub>far</sub>(wedges) − D<sub>far</sub>(rings) | 3.2 mm | 86 | −0.0047 | 46/86 | [−0.0087, −0.0006] | 0.010 | 0.013 |
| H2b | P4, P5 | D, contamination, rings | 2.4 / 3.2 mm | 86 | −0.0263 / −0.0209 | 71/86, 79/86 | – | < 0.001 | < 0.001 |
| H2c | secondary | D<sub>CWL</sub>(rings) − D<sub>far</sub>(rings) | 3.2 mm | 86 | −0.0207 | 78/86 | [−0.0243, −0.0170] | < 0.001 | – |
| H3a | P6, P7 | Δ_clean(half) − Δ_clean(full) > 0 | 2.4 / 3.2 mm | 86 | +0.0032 / +0.0010 | 71/86, 53/86 | [+0.0023, +0.0041], [+0.0002, +0.0018] | < 0.001, 0.007 | < 0.001, 0.013 |
| H4 | P8 | overall gain, PSF 15% too wide | 2.4 mm | 86 | −0.0038 | 63/86 | [−0.0051, −0.0025] | < 0.001 | < 0.001 |

The contamination dose-response replicates at both grids, also from the coarse data alone, and for eccentricity rings as for polar angle (Fig. 7). The plain far-leak contrast is clear at 2.4 mm but weak at 3.2 mm; for rings at 3.2 mm it is absent (−0.0002 [−0.0036, +0.0031]) while contamination predicts the ring gain in 79 of 86 families. The development set was optimistic in the highest-contamination bin (−0.022 against −0.037 at 3.2 mm). Holm-adjusted across the eight primary tests, every one remains below 0.05 (largest adjusted p = 0.013, for P3 and P7); under a Bonferroni correction these two, the smallest effects, would not (0.053 and 0.077) (Supplementary S13.7). The plain far-leak contrast at 3.2 mm (secondary, p = 0.015) is the weakest result of the cohort.

**Figure 7.**
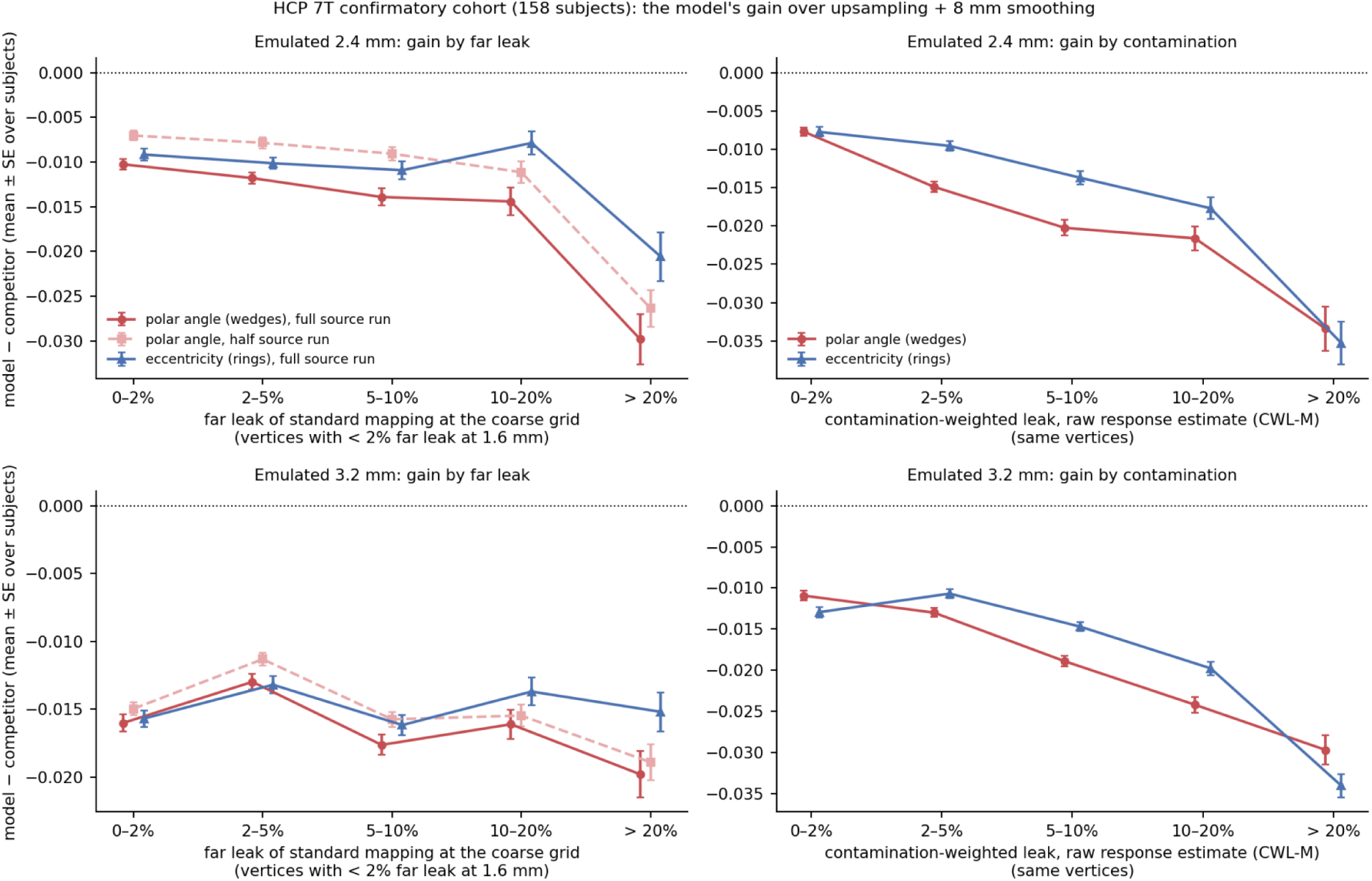
Confirmatory cohort (158 further HCP subjects). Model − competitor (upsampling + 1.6 mm mapping + 8 mm smoothing) at emulated 2.4 mm (top) and 3.2 mm (bottom), mean ± SE over all 158 subjects (the tests of Section 4.4 use one subject per family; the SE bars ignore family structure). Left: by the far leak of standard mapping at the coarse grid, for polar-angle maps from full and from half source runs and for eccentricity maps (rings); the nearly flat 3.2 mm curves are the D_far_ = −0.005 result. Right: by the contamination-weighted leak with the source run’s own native map as the response estimate (CWL-M), for wedges and rings. Vertices with < 2% far leak at 1.6 mm.

The independent 3T data (NYU, 35 subjects) showed the same dissociation, in data from another site, scanner, field strength and stimulus (Table 5):

**Table 5.** External replication at 3T (NYU, 35 test subjects): dose contrasts with Holm-adjusted p values.

| **test (prospective, plan 1; Holm over the six)** | **3.0 mm (×1.5)** | **4.0 mm (×2.0)** |
|---|---|---|
| N1: model − fixed competitor, responsive vertices | +0.005 [+0.001, +0.008], 13/35, Holm p = 1.0 | −0.007 [−0.011, −0.004], 27/35, Holm p < 0.001 |
| N2: D<sub>far</sub> | −0.002 [−0.011, +0.006], 19/35, Holm p = 0.93 | +0.003 [−0.004, +0.011], 18/35, Holm p = 1.0 |
| N3: D<sub>CWL-M</sub> | −0.015 [−0.026, −0.005], 25/35, Holm p = 0.018 | −0.011 [−0.019, −0.003], 25/35, Holm p = 0.029 |

At 3.0 mm the gain moves from +0.010 in the lowest contamination bin, where the model loses, to −0.005 in the highest, where it wins; by the plain far leak it shows no trend (Fig. 8B, C). Contamination estimated from the coarse data alone, which predicted the HCP gain, did not predict it at 3T (D_CWL-P_ +0.0001 and +0.0026, 21 and 19 of 35 subjects negative; descriptive, added after N1–N3). Contamination from an anatomical retinotopy atlas (Benson et al., 2014; Benson & Winawer, 2018), which uses no functional data of the subject, predicted the gain at twofold coarsening (D_CWL-A_ −0.0088 [−0.0170, −0.0007], 24/35, Holm p = 0.029) but not significantly at 1.5- fold (−0.0056 [−0.0131, +0.0017], 20/35, Holm p = 0.12; pre-specified, added after the results above) (Supplementary S16.6). These results support response-weighted contamination as a more informative retrospective stratifier of the gain than the plain signed leak in the tested 3T data. Its prospective utility is uncertain: the coarse-data-only version did not generalise, and the atlas version held only at twofold coarsening.

**Figure 8.**
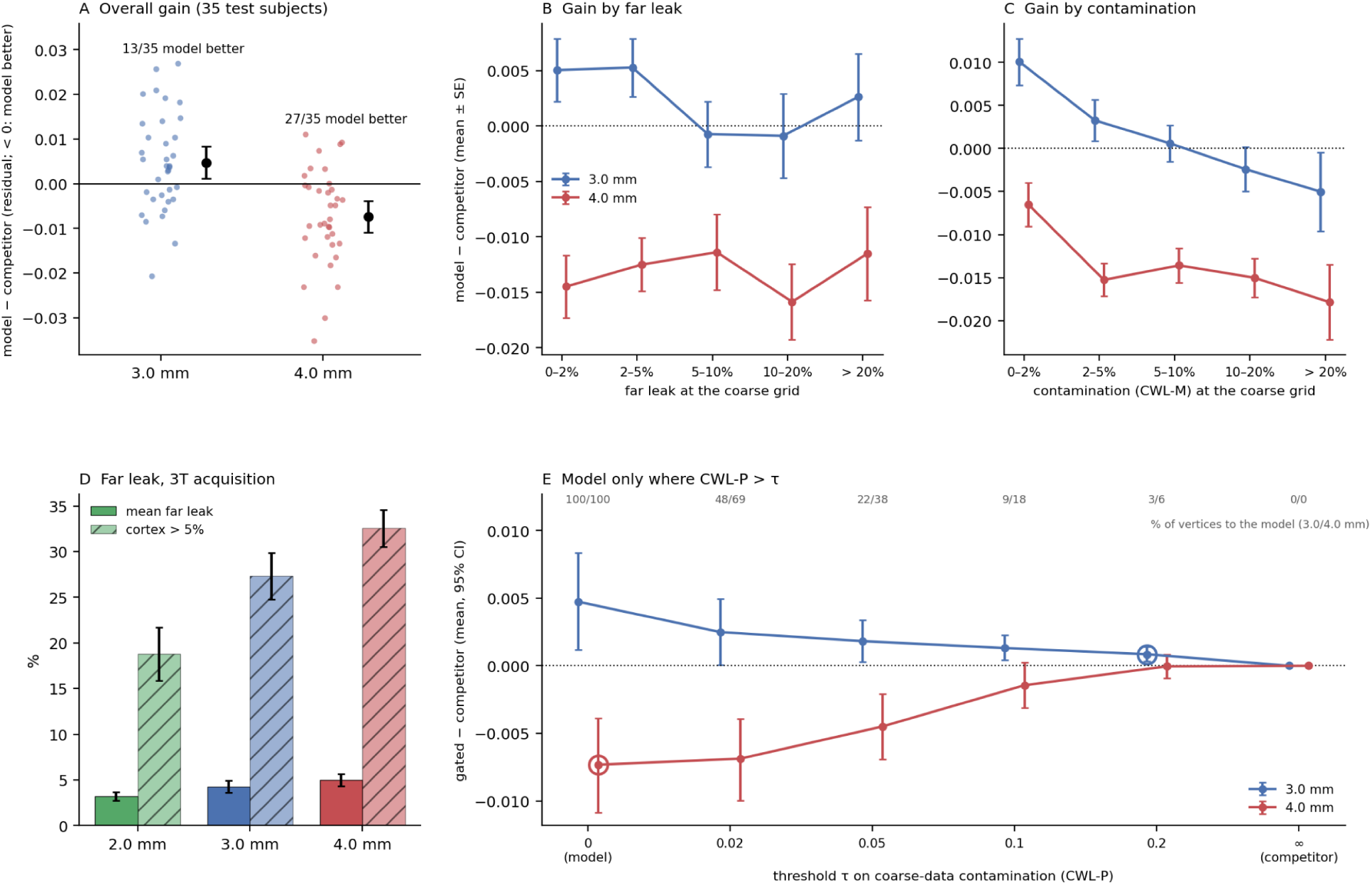
External replication in the NYU Retinotopy Dataset (3T, 2.0 mm, sweeping bars; 35 test subjects). (A) Model − competitor (upsampling + 2 mm smoothing, chosen on 8 tuning subjects) over responsive vertices, per subject, at emulated 3.0 and 4.0 mm; black: mean and bootstrap 95% interval; numbers: subjects in which the model is better. (B) The same difference by bin of the far leak of standard mapping at the coarse grid (responsive vertices with < 2% far leak at 2.0 mm; mean ± SE over subjects). (C) The same by bin of the contamination-weighted leak (CWL-M). (D) Far leak of standard mapping for the 3T 2.0 mm acquisition and its 3.0 / 4.0 mm emulations: mean far leak and the share of cortex above 5% (mean ± SD over 43 subjects). (E) A rule from the coarse data alone (added after A–C were known): the model’s estimate where the coarse-data contamination CWL-P exceeds τ and the competitor’s elsewhere, minus the competitor (mean and bootstrap 95% interval over the test subjects), for each threshold; ringed: the threshold chosen on the tuning subjects; numbers at the top: the share of responsive vertices sent to the model at 3.0 / 4.0 mm.

### 4.5 When correction pays

Across datasets, model − best competitor (per subject the better of the smoothing competitors, a descriptive, reference-dependent choice; NYU rows: the competitor fixed on the tuning subjects; mean [bootstrap 95%], subjects with the model better, one-sided p) over all scored vertices and where mixing is strong (far leak > 10% at the coarse grid, < 2% at the native grid), with the status of each comparison (Table 6):

**Table 6.** When correction pays: model − best competitor across datasets and regimes.

| comparison | regime | status | all vertices | better | p | strong mixing | better |
|---|---|---|---|---|---|---|---|
| HCP 7T, emulated 2.4 mm | single-run targets | pre-specified table; competitor chosen per subject | −0.007 [−0.011, −0.002] | 12/15 | 0.006 | −0.022 [−0.031, −0.012] | 13/15 |
| HCP 7T, emulated 3.2 mm | single-run targets | pre-specified table; competitor chosen per subject | −0.013 [−0.017, −0.007] | 14/15 | 0.002 | −0.026 [−0.033, −0.017] | 14/15 |
| NYU 3T, emulated 3.0 mm | half-session averages | pre-specified (N1); competitor fixed on tuning subjects | +0.005 [+0.001, +0.008] | 13/35 | 0.99 | +0.001 [−0.005, +0.006] | 15/35 |
| NYU 3T, emulated 4.0 mm | half-session averages | pre-specified (N1); competitor fixed on tuning subjects | −0.007 [−0.011, −0.004] | 27/35 | < 0.001 | −0.014 [−0.020, −0.007] | 28/35 |
| StudyForrest 7T, emulated 2.1–4.0 mm | many-run averages | pre-specified table; leave-one-subject-out settings | +0.014 [+0.001, +0.029] | 2/7 | 0.961 | −0.010 [−0.023, +0.005] | 5/7 |
| ds002702 7T, emulated 1.6 mm | many-run averages | pre-specified table; leave-one-subject-out settings | +0.029 [+0.023, +0.034] | 0/5 | 1.000 | −0.002 [−0.022, +0.016] | 3/5 |
| ds002702 7T, emulated 2.4 mm | many-run averages | pre-specified table; leave-one-subject-out settings | +0.003 [−0.004, +0.009] | 2/5 | 0.781 | +0.009 [−0.008, +0.026] | 2/5 |
| StudyForrest 3T → 7T, voxel ratio ≥ 2.5 | real second acquisition | pre-specified table; n = 5 | +0.014 [−0.004, +0.040] | 2/5 | 0.844 | – | – |
| ds002702 3T 2.4 mm → 7T 0.8 mm | real second acquisition | pre-specified table; n = 2 | −0.024 [−0.028, −0.021] | 2/2 | 0.250 | −0.029 [−0.049, −0.008] | 2/2 |
| ds002702 7T SE 1.05 mm → GE 0.8 mm | real second acquisition | pre-specified table; n = 2 | +0.086 [+0.081, +0.090] | 0/2 | 1.000 | – | – |

(HCP and StudyForrest/ds002702 rows: the pre-specified summary table, Supplementary S11.3; NYU rows: the competitor chosen on the tuning subjects. Dashes: fewer than 30 strongly mixed vertices.) Against the competitor fixed on the five tuning subjects (upsampling + 1.6 mm mapping + 8 mm smoothing), the HCP development residuals are 0.313 against 0.321 (13/15) at 2.4 mm and 0.325 against 0.339 (14/15) at 3.2 mm over all coherent vertices (Table 2), and 0.343 against 0.367 and 0.344 against 0.371 (14/15 each) where mixing is strong; in degrees of polar angle, a map-specific error, the model’s mean absolute error is lower than that competitor’s by 0.6° and 0.5° overall and by 1.3° and 1.5° where the far leak exceeds 10% (Supplementary S5.7).

With the single-run HCP reference maps the model improved the mean score in every scored region. In HCP the targets are single runs. The lead holds in 12–14 of 15 development subjects and in 82–85 of 86 cohort families (Section 4.4).

With averaged targets the result depends on the coarsening. NYU’s half-maps average two to six runs, and smoothing barely helps any method (the chosen kernels are 2 mm, against 8–10 mm in HCP). At twofold emulated coarsening the model’s overall lead was observed in this independent cohort (27 of 35 subjects; 28 of 35 where mixing is strong); at 1.5-fold it was not (13 of 35), and plain upsampling is better still (+0.009). With many-run 7T averages (StudyForrest, ds002702 within a session) the model trails overall and is close to the competitor where mixing is strong (Fig. 9).

**Figure 9.**
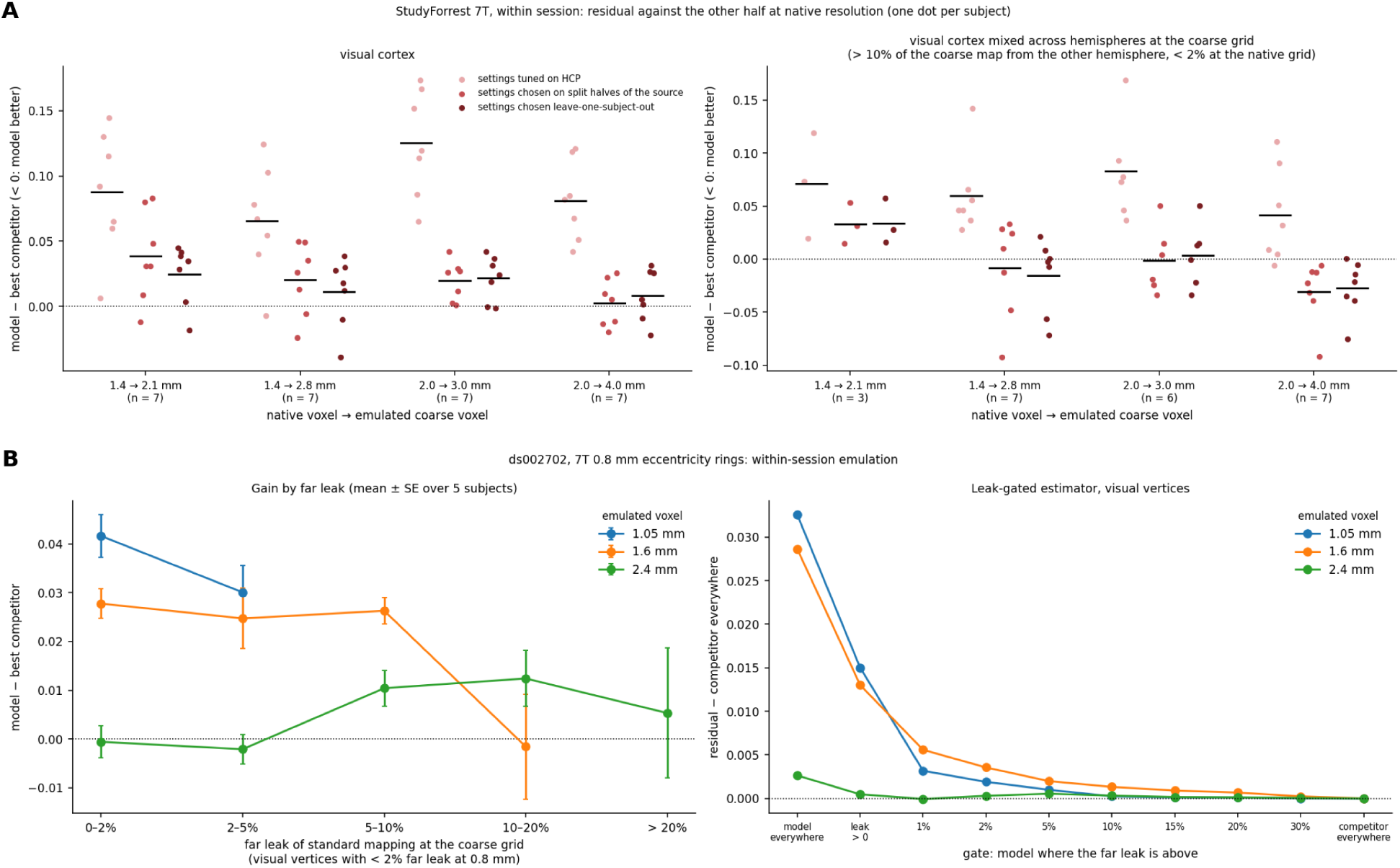
Resolution-limited real data within one session. (A) StudyForrest: model − best competitor in visual cortex (left) and where the coarse map mixes the hemispheres (right; cells with ≥ 30 such vertices: 3, 7, 6 and 7 of 7), under three setting rules. (B) ds002702: model − best competitor by far leak (left), and the leak-gated estimator (right; residual of the gated estimate minus that of the competitor used everywhere), for three emulated voxel sizes.

The real cross-session comparisons, with two to five subjects each, did not establish a reliable overall gain. Field-map correction mattered more than the estimator (up to 0.12). The two-subject gain for ds002702 3T → 7T did not replicate in StudyForrest, which lacks 3T field maps, and a spin-echo source penalised the model, whose forward model has no term for the different vascular weighting of SE and GE BOLD (Fig. 10).

**Figure 10.**
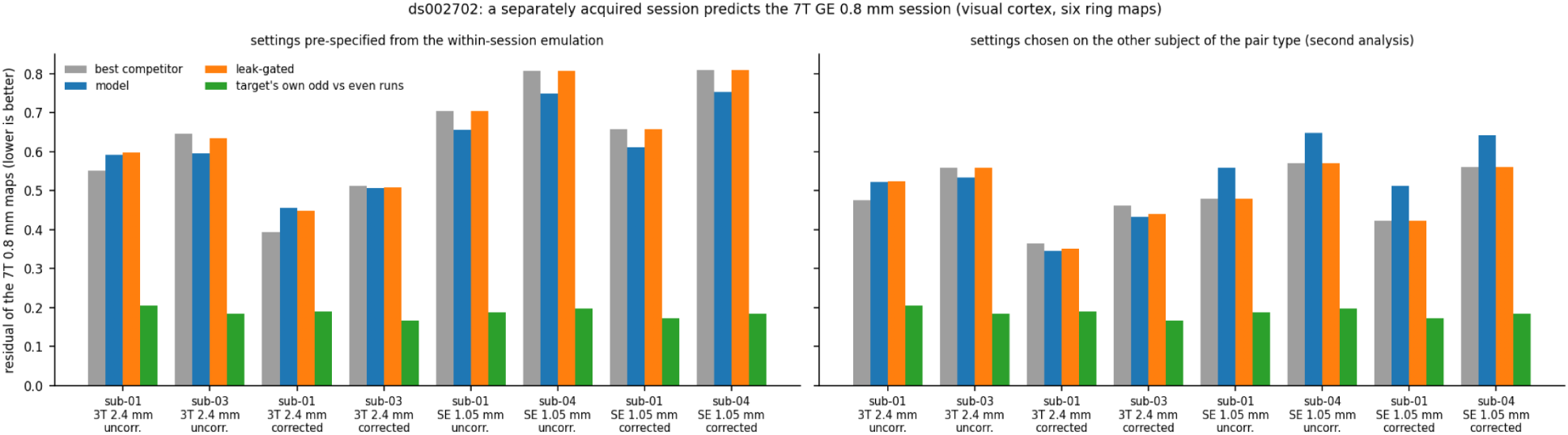
Real second acquisitions (ds002702). Residual of the 7T GE 0.8 mm maps predicted from a separately acquired 3T 2.4 mm or 7T SE 1.05 mm session, uncorrected or field-map corrected. Left: settings pre-specified from the within-session emulation; both methods carry within-session settings that are wrong across sessions (the competitor needs 6–8 mm of smoothing there), so this comparison is uninformative although the model’s bar is the lower one in 6 of 8 pairs. Right: settings chosen on the other subject of the pair type. “Best competitor” is the competitor chosen by the rule named in the panel title. Green: the target’s own odd-run vs even-run agreement.

In this 3T evaluation, a vertex-level rule based on the coarse-data statistic did not improve on a method selected globally on the tuning subjects. A second NYU plan, written after N1–N3 were known, gave each vertex the model’s estimate where CWL-P exceeds a threshold τ and the competitor’s elsewhere, with τ chosen on the tuning subjects (Fig. 8E). The choice was τ = 0.2 at 3.0 mm, which sends 3% of responsive vertices to the model, and τ = 0, the model everywhere, at 4.0 mm. The rule therefore beat the competitor only at 4.0 mm (−0.0073, 27/35, Holm p < 0.001), and the model only before correction at 3.0 mm (−0.0039, 22/35, Holm p = 0.090); at neither grid did it beat both. On the test subjects every intermediate threshold scored between the two pure strategies, and contamination from the coarse data alone did not predict the 3T gain (Section 4.4). What worked was the whole-map choice on eight tuning subjects: the competitor at 1.5-fold and the model at twofold coarsening, the better strategy at each grid.

The available quality metrics did not identify registration error as the explanation for the pericalcarine shortfall. At 3T the model was worse than the competitor in the pericalcarine cortex at both grids (+0.0125 and +0.0092). Across the 35 test subjects, all well registered (boundary-based registration cost 0.30–0.53), this shortfall did not grow with the registration cost (Spearman ρ = +0.10 and +0.12, p ≥ 0.49) or with the field inhomogeneity at the responsive pericalcarine vertices (ρ = +0.09 and −0.19). The one poorly registered tuning subject (cost 0.68) lost clearly over all responsive vertices (+0.045 and +0.051).

### 4.6 What the gain is

Explicit far-field terms were not essential for most of the observed improvement. Inverting with the far entries of the operator removed (per voxel, the columns farther than 15 mm from its main unknown: 18% and 25% of the voxel mass at 2.4 and 3.2 mm) gave very similar residuals (no equivalence margin was pre-specified): overall residual 0.3126 against 0.3131 (2.4 mm) and 0.3250 against 0.3248 (3.2 mm), and the same far-leak contrast (D_far_(near) − D_far_(full) +0.0025, p = 0.06, and 0.0000) (Fig. 3A). The model’s averaging kernels have no net far leak, but their gross far weight falls only from 19% to 14% (2.4 mm) and from 26% to 23% (3.2 mm): the estimate shows a reduced net far contribution while substantial unsigned kernel weight remains at distant locations; the estimator cancels the mean of the far field, it does not stop averaging over far cortex.

The reduced-data analysis does not support an explanation by noise suppression alone. With half the source data the gain where no far unmixing is needed decreased rather than increased: +0.003 (2.4 mm, 71/86, p < 0.001) and +0.001 (3.2 mm, 53/86, p = 0.007) in the cohort with fixed settings, and +0.0026 (14/15) and +0.0022 (13/15) in the development set with both methods retuned for half-run sources.

Together these findings are consistent with an important contribution from local deconvolution of the PSF within about 15 mm of each vertex, from exact sampling of the cortical ribbon and from the surface regularisation; their relative contributions have not been isolated, because no matched comparison with a smoothing-only operator, a regulariser-only model or a model with an empirically estimated PSF was run (Section 5). The gain was largest where the cortex folds tightly, which is where the leakage map is high and where standard mapping and surface smoothing are worst.

## 5. Discussion

Under its stated assumptions, the operator predicts a signed measure of the contribution of far cortex to each surface value before the functional data are analysed, and its far part is geometrically right in emulated and in real coarser acquisitions (Section 4.1). The leakage map alone does not say whether correction pays; the response-weighted contamination was the more informative description of where the gain fell. In HCP it predicted the model’s gain in the development set, in 158 further subjects and for rings as well as wedges; in independent 3T data it predicted the gain at both coarsenings while the plain far leak predicted nothing (Section 4.4). Its coarse-data-only version predicted as well in HCP but not at 3T. An anatomical atlas did predict the 3T gain at twofold coarsening, the regime where correction pays at 3T; this supports a prospective evaluation of atlas-derived contamination maps for deciding where correction might be worthwhile under strong coarsening, but it does not yet establish a general rule.

The anatomical mixing measure alone is not a sufficient predictor of the benefit. We expected eccentricity rings to be a negative control for the plain leak, because they drive both banks of the calcarine alike. They were so only in ds002702 at 2.4 mm; elsewhere rings carried far-leak dose-responses of their own, because foveal and peripheral representations are far apart on the sheet and the far part of the kernel mixes different eccentricities too. The response-weighted measure showed the more consistent association with the gain across the pre-specified HCP and NYU comparisons; the evidence from the smaller real-resolution datasets (five and seven subjects) remains limited.

In practice, the model merits consideration where the reference data are noise-limited or the coarsening is strong, and most of its gain can be expected where contamination is high: single 7T retinotopy runs (82–85 of 86 families) or a twofold coarsening at 3T (27 of 35 subjects). With averaged targets and a mild coarsening, k-space upsampling with light smoothing is as good, and better outside the contaminated cortex. In the 3T test, choosing one method per coarsening on eight tuning subjects picked the better one at both, and a vertex-wise switch driven by coarse-data contamination added nothing to that choice (Section 4.5). Good registration is a precondition: the model uses the geometry to undo mixing, and a misplaced geometry undoes the wrong mixing (the poorly registered NYU tuning subject lost clearly at both grids), but among well-registered subjects the 3T pericalcarine shortfall did not depend on registration quality, and its cause is open. The operator costs 30–50 min per subject to build and a fit 1–25 min on 28 cores.

In residual energy the differences in the favourable comparisons are 0.01–0.03 of the target’s energy; per vertex they are half a degree of polar angle overall and about two degrees where mixing is strong, against a single-run noise of about 30° (Section 4.3). The practical significance is therefore a modest, spatially patterned difference rather than a large change at individual vertices, concentrated where the contamination map is high: for group maps, and for analyses that compare responses across sulcal banks, where the model keeps more of the contrast.

Emulation may overstate the gains that an independent coarse acquisition would show. The emulation window is known exactly and is shared by the forward model and by band-limited upsampling, whereas the native PSF and the processing history are only approximated; hyperparameters tuned in one regime did not transfer to another, and real second acquisitions showed no robust gain (Section 4.5), with field-map correction mattering more than the estimator. A convincing test needs a stimulus with contrast across sulcal banks (polar angle) at several resolutions in the same people, with field maps. One multi-sequence visual-fMRI dataset may be suitable (Schellekens et al., 2025); whether it provides a matched-contrast resolution benchmark would have to be checked before it is used for this purpose.

All gains are measured against finer fMRI reference maps, not against the true cortical response. A cross-run reference carries its own noise, physiological variability and registration error, so a lower residual shows closer agreement between two fMRI measurements made at different resolutions. The latent field that the estimator recovers was not validated independently, the PSF was not measured in these subjects, and the only known-truth evidence is the simulation of Section 4.3.

The study has limitations. The HCP and NYU coarse data are emulated: the window is a modelling choice, the emulated PSF is 1.6–1.7 voxels wide, and the noise is band-limited with the signal. The far part was tested against real coarser sessions in one dataset. The PSF model ignores partial Fourier and approximates spin-echo decay, and it has no term for motion, residual B0 distortion, receive-coil weighting, magnitude reconstruction or the vascular origin of the signal; a kernel-width scan suggests it is 15–30% too wide, which would lower the leak figures by about a fifth, and its parameters were not estimated per subject. “Far” is a 15 mm convention on an area-scaled sphere; Section 4.2 gives the figures at other thresholds and with true geodesics. The likelihood treats the noise as spatially white (a stationary correlated alternative made the estimates worse), the posterior uncertainty is not calibrated without an empirical factor, the model has no compartment for pial veins or CSF, and it takes the response to be the same at all cortical depths. Contamination depends on a response estimate, which is noisy where the true contrast is confined to few vertices; estimated from the coarse data alone, it predicted the gain in HCP but not at 3T, for reasons we do not know. The atlas-based version was tested only at 3T, where it held at twofold but not at 1.5-fold coarsening. The confirmatory tests are one-sided; the plan set no correction across its eight primary tests, under Holm all eight remain below 0.05, and the two smallest would not survive Bonferroni (Section 4.4). Two cohort hypotheses and the later NYU analyses were formulated after earlier results were known, and pre-specification rests on the author’s hashed plans, not on a public timestamp. The coherent and responsive vertex sets depend on the reference as well as the source data, and the per-subject best competitor of Section 4.5 is chosen on the test outcome (Section 3.4). The mechanistic ablations lack a matched smoothing-only or regulariser-only comparator, and the sign diagnostics of the leak are limited to the hemisphere-level summaries of Section 4.1. The real cross-session comparisons have two to five subjects and cannot support generalisation. Only visual-field mapping paradigms were tested (rotating wedges, expanding rings, sweeping bars), and real two-resolution data are few.

## Declaration of the use of AI

The research question, hypotheses, study design and interpretation of the results are the author’s; an AI assistant worked under the author’s direction throughout. The assistant was Claude Code (Anthropic; version 2.1.280) with the models Claude Opus 5.5 (claude-opus-5-5) and, from 2 to 5 October 2026, Claude Fable 5.1 (claude-fable-5-1). In the research, it wrote and debugged the analysis code, drafted the pre-specified analysis plans for the author’s approval, ran the analyses on the computing servers, compiled the results and checked every number in the manuscript against the result files by script. In preparing this paper, we used AI tools for language editing and manuscript restructuring. All AI-assisted content was subsequently verified for accuracy. The AI assistant drafted text under the author’s direction, which the author then revised and edited. All figures were generated using plotting code written by the AI assistant; no AI image generation was used. References were checked against Crossref and DataCite records by script, and each was verified by the author. The author takes full responsibility for the content of this article.

## Data and Code Availability

HCP 7T retinotopy: ConnectomeDB (db.humanconnectome.org), under the WU-Minn HCP Open Access and Restricted Data Use Terms. Restricted family information was used only to assign subjects to the tuning, development and cohort sets and to pick one subject per family for the cohort’s primary tests; it is not shared. As the Restricted Data Use Terms require, HCP subject IDs are not published: the release identifies subjects by study codes, and the key linking the codes to HCP IDs will be submitted to the HCP on acceptance. A script in the repository rebuilds every subject set from the HCP restricted table and the key, and checks the larger sets against SHA-256 sums given in the repository.

StudyForrest multi-resolution data: studyforrest.org.

ds002702: OpenNeuro, doi:10.18112/openneuro.ds002702.v1.0.1 (CC0), with the authors’ FreeSurfer reconstructions at osf.io/j2wsc.

NYU Retinotopy Dataset: OpenNeuro ds003787, version 1.0.0 (doi:10.18112/openneuro.d-s003787.v1.0.0, CC0), the snapshot downloaded and analysed here; the later snapshot 1.0.1, created the same day, lists the same files and subjects. The authors’ FreeSurfer reconstructions and stimulus apertures are in its derivatives.

Code, analysis plans and group leakage maps (GIFTI: HCP 7T geometry on fs_LR 32k, the NYU 3T acquisition on fsaverage): https://github.com/halilaziz/fmri-surface-leakage (a persistent identifier for the archived release will be added); the repository documents how to rerun each analysis, and its README lists the analysis stages and their folders.

## Author Contributions

Halil Aziz Velioğlu: Conceptualization, Methodology, Software, Validation, Formal analysis, Investigation, Data curation, Visualization, Writing – original draft, Writing – review & editing, Project administration.

## Acknowledgements

Data were provided in part by the Human Connectome Project, WU-Minn Consortium (Principal Investigators: David Van Essen and Kamil Ugurbil; 1U54MH091657) funded by the 16 NIH Institutes and Centers that support the NIH Blueprint for Neuroscience Research; and by the McDonnell Center for Systems Neuroscience at Washington University. We thank the StudyForrest, ds002702 and NYU Retinotopy Dataset authors for sharing their data. The numerical calculations reported in this paper were partially performed at TUBITAK ULAKBIM, High Performance and Grid Computing Center (TRUBA resources).

## Supplementary Material

Supplementary material (Supplementary S1–S17 and Figures S1–S2): data and acquisition, operator and estimator details, results in full, the analysis plans verbatim with their hashes, quality control, software and compute, and the full-length text of the analyses (S17).

