## Supplementary Material (S1-S17, Figures S1-S2) for "Predicting cortical signal mixing in coarse-resolution fMRI: a forward-operator analysis of surface mapping and the conditional benefit of correction"

Tables in this supplement are copied from the study's analysis reports, not retyped, so every number matches its source. The code repository (https://github.com/halilaziz/fmri-surface-leakage; a persistent identifier for the archived release will be added) holds the code and documents how to rerun each analysis.

Contents: S1 data and acquisition · S2 forward operator and emulation · S3 estimator · S4 competitors, scoring and setting selection · S5 HCP results in full · S6 StudyForrest · S7 ds002702 · S8 contamination-weighted leak and pre-declared tests · S9 quality control, including a processing bug and its check · S10 resolvability theory · S11 analysis plans · S12 software and compute · S13 confirmatory cohort (158 further HCP subjects) · S14 reviewer-requested tests (far-part ablation, realistic kernels, real coarser sessions) · S15 reviewer-requested tests, second set (kernel-width scan, re-tuned half-run settings, correlated noise, the far criterion) · S16 external replication in the NYU Retinotopy Dataset (3T), with the coarse-data-only rule, registration, 3T group leakage maps and contamination from an anatomical atlas · S17 the full-length text of draft v1.5 (Sections 2–4) · Figures S1–S2.

The main text is the condensed version 2. The copied reports and plans cite sections and figures of the full-length text, which is reproduced as S17 with its own section numbering. Its Figures 1–11 are, in order, the main text's Figures 1, 2, 4, 5, 6, 9, 10, 7 and 3, Figure S1, and the main text's Figure 8; S17 itself cites figures in the main text's numbering.

##### S1. Data and acquisition

**HCP 7T retinotopy.** 20 subjects of the 7T release (the development set): 5 for tuning and 15 held out, assigned by family (restricted family information was used only for this split and is not reported). Released MNINonLinear time series, 1.6 mm isotropic. The forward model used an effective echo spacing of 0.32 ms, a phase-encoding field of view of 208 mm along y and T2* = 28 ms, and in-plane Tukey taper 0.3. It also used a slice-profile edge of 0.15 voxel, 0.4 mm interpolation blur, and MNI/ACPC axis scaling from each subject's warps. Wedge runs (counter-clockwise and clockwise; 22 s blank, then 8 cycles of 32 s at TR 1 s); response = the Fourier component at the stimulus frequency; coherent vertices: coherence > 0.3 at 1.6 mm. HCP's own good-voxels masks and ribbon weights define standard mapping. The 20 development subjects (5 tuning, 15 development) belong to 20 different families, and no cohort subject shares a family with a tuning subject (checked by `src/14_robustness_checks/verify_splits.py` against the restricted table).

**HCP confirmatory cohort** (S13; S17, Sections 3.9 and 4.9; main text Section 4.4). The 158 remaining 7T retinotopy subjects (86 families) and a sensitivity set of 5, with the same preprocessing and settings. Besides the wedge runs, the eccentricity runs were analysed (expanding and contracting rings; the same 22 s blank and 8 cycles of 32 s, confirmed on the pericalcarine power spectrum), with ribbon weights from the ring runs' own goodvoxels masks, and wedge fits were repeated from the first and from the second half of each run (4 cycles; the reference map always comes from the full other run). Raw volumes (2.8 GB per subject) were deleted after the fits, each with a SHA-1 manifest; the per-subject maps, weights, contamination maps, scores and logs are kept.

**StudyForrest multi-resolution** (Sengupta et al., 2017, 2018). 7 subjects with 7T sessions at 0.8 / 1.4 / 2.0 / 3.0 mm (10 runs each; distortion-corrected by the providers), 5 of them also with 3T sessions at 1.4 / 2.0 / 3.0 mm; left/right hemifield orientation stimulus. The 7T echo spacings are not published and were assumed; there are no field maps for the 3T sessions.

**ds002702** (Kay et al., 2019, 2020; OpenNeuro):

| session | subjects | voxel | runs | TR | readout (effective echo spacing × PE lines) | other |
|---|---|---|---|---|---|---|
| 7T GE (ring7T) | 5 | 0.8 mm | 9 | 2.2 s | 0.333 ms × 162 | in-plane R 3, MB 2, partial Fourier 6/8, TE 22.6 ms |
| 3T GE (ring3T) | sub-01, sub-03 | 2.4 mm | 6 / 5 | 1.1 s | 0.55 ms × 80 | MB 2, full Fourier, TE 30 ms |
| 7T SE (ringSE) | sub-01, sub-04 | 1.05 mm | 9 | 2.2 s | 0.5 ms × 106 | R 2, MB 2, partial Fourier 6/8, TE 39 ms |

Known issue reported with the dataset: sub-04 ring7T misses a small amount of brain at the right occipital pole. sub-03 ring3T has 5 runs, sub-01 ring3T 6 (one dropped by the pre-declared run-quality rule, S9).

##### S2. Forward operator and emulation

The effective PSF is evaluated per axis of the acquisition voxel frame, as described below.

- **Frequency-encoding and phase-encoding axes:** a Tukey-tapered sinc. Along phase encoding it is multiplied by the T2* decay over the echo train (effective echo spacing × lines from the k-space centre).
- **Slice axis:** an erf-edged slice profile.
- **All axes:** a Gaussian interpolation blur.

Kernels are tabulated, truncated with a smooth roll-off at (4, 4, 3) voxels for native grids and (5, 5, 5) voxels for emulated coarse grids, and renormalised to unit integral. Cortical columns are integrated at quadrature points between the white and pial surfaces (column mean b0; optionally a depth gradient b1).

Far cortex, for the far leak and the contamination measures (main text, Section 2.2), is the other hemisphere, or cortex more than 15 mm away on the subject's own spherical surface (FreeSurfer `?h.sphere`; in HCP `sphere.native`), the inflation of the cortex to a sphere from which registration starts, scaled to the cortical area of the hemisphere's midthickness surface. The analysis plans

(S11) and the full-length text (S17) call this the registration sphere; registered spheres (`?h.-sphere.reg`, HCP MSMAll) were used only to carry maps between subjects and templates.

**Emulation.** Coarse grids are emulated as the central k-space block times a separable Tukey-0.5 window, which is zero at the coarse Nyquist. A sharp crop would leave sinc ringing that no compact operator represents. The emulated kernel is the native kernel's spectrum times the same window. On oblique grids the operator is applied per axis with exact matrices.

Numerical checks (HCP):

| grid | effective PSF FWHM x / y / z (mm) | nonzeros in K (millions) | median voxels per vertex (HCP weights) |
|---|---|---|---|
| 1.6 mm | 2.56 / 2.70 / 2.55 | 194 | 9 |
| 2.4 mm | 3.83 / 4.08 / 3.96 | 399 | 4 |
| 3.2 mm | 5.07 / 5.36 / 5.11 | 386 | 2 |

MNI/ACPC scale x/y/z [1.096, 1.041, 1.156]; HCP goodvoxels (RETCCW) leave 0.25 ± 0.12% of cortex without usable voxels (excluded from the averages); run time per subject 30.0 ± 6.4 min.

**Checks (test subjects)**

| subject | grid | brute-force max rel. error | column mass p1 / p50 / p99 |
|---|---|---|---|
| T01 | 1.6 | 1.1e-07 | 0.990 / 1.000 / 1.010 |
| T01 | 2.4 | 1.2e-07 | 0.997 / 1.000 / 1.003 |
| T01 | 3.2 | 1.0e-07 | 0.997 / 1.000 / 1.004 |
| T02 | 1.6 | 1.1e-07 | 0.989 / 1.000 / 1.011 |
| T02 | 2.4 | 1.1e-07 | 0.997 / 1.000 / 1.003 |
| T02 | 3.2 | 1.1e-07 | 0.997 / 1.000 / 1.004 |

Emulation consistency, S(K_1.6 f) vs K_g f (relative L2 error on the operator support / after standard mapping):

| subject | field | 2.4 mm | 3.2 mm |
|---|---|---|---|
| T01 | uniform | 0.004 / 0.003 | 0.004 / 0.002 |
| T01 | smooth_8-30mm | 0.005 / 0.004 | 0.005 / 0.004 |
| T01 | rough_white | 0.012 / 0.009 | 0.011 / 0.009 |
| T01 | single columns (median of 5) | 0.011 | 0.010 |
| T02 | uniform | 0.004 / 0.003 | 0.004 / 0.002 |
| T02 | smooth_8-30mm | 0.005 / 0.004 | 0.005 / 0.003 |
| T02 | rough_white | 0.012 / 0.009 | 0.011 / 0.008 |
| T02 | single columns (median of 5) | 0.010 | 0.008 |

Depth-gradient component b1 (native grid), median |net| / gross column mass: T01 0.122, T02 0.126

Standard mapping on real geometry (unit field):

| | dilution (own + leak) | leak (% of unit field) | of which other hemisphere | leak magnitude | leak fraction (%) | cortex with leak fraction > 5% / > 10% | pericalcarine leak fraction (%) |
|---|---|---|---|---|---|---|---|
| 1.6 mm (HCP standard, goodvoxels) | 0.800 ± 0.010 | 1.70 ± 0.29 | 0.17 ± 0.03 | 3.06 ± 0.43 | 2.13 ± 0.34 | 13.0 ± 2.7% / 4.7 ± 1.5% | 3.77 ± 1.04 |
| 1.6 mm (no voxel mask) | 0.791 ± 0.011 | 1.74 ± 0.30 | 0.18 ± 0.03 | 2.93 ± 0.40 | 2.18 ± 0.35 | 13.7 ± 2.9% / 4.7 ± 1.6% | 3.88 ± 1.10 |
| 2.4 mm emulated | 0.733 ± 0.015 | 1.95 ± 0.43 | 0.19 ± 0.05 | 9.93 ± 0.87 | 2.56 ± 0.53 | 17.9 ± 3.0% / 9.8 ± 2.3% | 5.03 ± 1.60 |
| 3.2 mm emulated | 0.685 ± 0.017 | 2.73 ± 0.55 | 0.27 ± 0.08 | 18.55 ± 1.37 | 3.62 ± 0.69 | 25.8 ± 2.7% / 16.8 ± 2.5% | 6.45 ± 2.10 |

Leak (% of a unit far field) by opposite-bank distance, and the two-bank planar model:

| d_opp bin (mm) | area share | 1.6 mm (HCP standard, goodvoxels) | 1.6 mm (no voxel mask) | 2.4 mm emulated | 3.2 mm emulated | planar model 7T | planar model 3T |
|---|---|---|---|---|---|---|---|
| 0-2.25 | 0.4% | 12.77 ± 1.55 | 13.04 ± 1.56 | 17.93 ± 1.68 | 20.47 ± 1.77 | 14.09 | 15.46 |
| 2.25-2.75 | 1.7% | 9.19 ± 0.89 | 9.35 ± 0.86 | 13.69 ± 1.05 | 17.79 ± 1.25 | 11.63 | 12.54 |
| 2.75-3.25 | 5.1% | 6.74 ± 0.66 | 6.85 ± 0.68 | 9.90 ± 0.81 | 14.12 ± 0.87 | 4.80 | 5.42 |
| 3.25-3.75 | 8.4% | 4.21 ± 0.57 | 4.35 ± 0.59 | 5.97 ± 0.75 | 9.61 ± 0.92 | 1.87 | 1.78 |
| 3.75-4.5 | 15.0% | 2.30 ± 0.32 | 2.37 ± 0.33 | 2.32 ± 0.46 | 4.45 ± 0.64 | 0.81 | 0.45 |
| 4.5-6 | 26.4% | 1.19 ± 0.14 | 1.19 ± 0.14 | 0.46 ± 0.20 | 0.19 ± 0.24 | 0.25 | 0.05 |
| 6-inf | 42.5% | 0.27 ± 0.06 | 0.27 ± 0.05 | 0.34 ± 0.06 | 0.14 ± 0.12 | – | – |

Leak magnitude (sum of |A| over far cortex, % of a unit field; what a patterned far field can carry before positive and negative sidelobe contributions cancel) by the same bins:

| d_opp bin (mm) | 1.6 mm (HCP standard, goodvoxels) | 1.6 mm (no voxel mask) | 2.4 mm emulated | 3.2 mm emulated |
|---|---|---|---|---|
| 0-2.25 | 19.66 ± 2.19 | 19.36 ± 2.17 | 38.20 ± 3.54 | 51.81 ± 4.69 |
| 2.25-2.75 | 14.50 ± 1.18 | 14.12 ± 1.11 | 30.56 ± 1.97 | 44.33 ± 2.63 |
| 2.75-3.25 | 11.09 ± 0.72 | 10.71 ± 0.70 | 24.96 ± 1.30 | 38.61 ± 1.81 |
| 3.25-3.75 | 7.75 ± 0.55 | 7.46 ± 0.54 | 19.81 ± 1.03 | 32.93 ± 1.56 |
| 3.75-4.5 | 4.72 ± 0.30 | 4.50 ± 0.30 | 14.35 ± 0.72 | 26.08 ± 1.36 |
| 4.5-6 | 2.17 ± 0.11 | 2.04 ± 0.11 | 8.87 ± 0.41 | 17.84 ± 0.96 |
| 6-inf | 0.46 ± 0.05 | 0.44 ± 0.05 | 4.10 ± 0.22 | 9.66 ± 0.54 |

Per-vertex correlation with the 7T planar-model map (1.6 mm, no mask): 0.699 ± 0.037; with goodvoxels: 0.660 ± 0.037.

Cross-resolution prediction through the operator:

Residual energy fraction of the emulated map after the best complex scaling of each predictor (both wedge runs pooled); mean ± SD over hemispheres, and the number of hemispheres in which adding the far part (or a shuffled far part) beats the near part alone.

| grid | vertices | n per hemisphere | 1.6 mm map as is | operator, near part | operator, near + far | near + shuffled far | near + far better | near + shuffled better |
|---|---|---|---|---|---|---|---|---|
| 2.4 mm | all coherent vertices | 13236 ± 2653 | 0.1196 ± 0.0216 | 0.0538 ± 0.0115 | 0.0451 ± 0.0106 | 0.0601 ± 0.0123 | 40/40 | 0/40 |
| 2.4 mm | model leak fraction < 2% | 8927 ± 1755 | 0.1186 ± 0.0214 | 0.0432 ± 0.0096 | 0.0422 ± 0.0094 | 0.0438 ± 0.0096 | 39/40 | 0/40 |
| 2.4 mm | model leak fraction > 5% | 2237 ± 635 | 0.1331 ± 0.0288 | 0.0866 ± 0.0201 | 0.0574 ± 0.0176 | 0.1203 ± 0.0241 | 40/40 | 0/40 |

| grid | vertices | n per hemisphere | 1.6 mm map as is | operator, near part | operator, near + far | near + shuffled far | near + far better | near + shuffled better |
|---|---|---|---|---|---|---|---|---|
| 2.4 mm | model leak fraction > 10% | 1335 ± 419 | 0.1379 ± 0.0358 | 0.0953 ± 0.0276 | 0.0587 ± 0.0233 | 0.1500 ± 0.0336 | 40/40 | 0/40 |
| 2.4 mm | coherent pericalcarine | 1200 ± 304 | 0.1208 ± 0.0345 | 0.0475 ± 0.0140 | 0.0388 ± 0.0133 | 0.0541 ± 0.0153 | 40/40 | 0/40 |
| 3.2 mm | all coherent vertices | 13236 ± 2653 | 0.2237 ± 0.0289 | 0.0709 ± 0.0151 | 0.0484 ± 0.0125 | 0.0854 ± 0.0173 | 40/40 | 0/40 |
| 3.2 mm | model leak fraction < 2% | 8325 ± 1635 | 0.2380 ± 0.0314 | 0.0482 ± 0.0108 | 0.0439 ± 0.0106 | 0.0501 ± 0.0110 | 40/40 | 0/40 |
| 3.2 mm | model leak fraction > 5% | 3047 ± 779 | 0.2119 ± 0.0300 | 0.1121 ± 0.0223 | 0.0581 ± 0.0174 | 0.1665 ± 0.0290 | 40/40 | 0/40 |
| 3.2 mm | model leak fraction > 10% | 2075 ± 560 | 0.2212 ± 0.0332 | 0.1236 ± 0.0255 | 0.0590 ± 0.0181 | 0.2044 ± 0.0345 | 40/40 | 0/40 |
| 3.2 mm | coherent pericalcarine | 1200 ± 304 | 0.2259 ± 0.0478 | 0.0626 ± 0.0197 | 0.0394 ± 0.0150 | 0.0774 ± 0.0242 | 40/40 | 0/40 |

Model: effective PSF = native 7T PSF (Tukey 0.3 in-plane, T2* along PE (y), slice profile, 0.4 mm resampling blur; MNI/ACPC-scaled), truncated at 4/4/3 voxels; emulated grids = central k-space block x Tukey 0.5 window (zero at the coarse Nyquist), operators at 5 voxels support. Signal only from grey matter (column mean, b0). Far cortex = other hemisphere or > 15 mm geodesic on the sphere. Emulated acquisitions keep the 7T noise and physiological structure (k-space cropping of the acquired data), so their SNR gain is at most sqrt(voxel-volume ratio), less than a native acquisition at that voxel size would give.

#### S3. Estimator

Posterior mean of the field f on each hemisphere's cortex under a Matérn ($\nu = 1$) prior. The prior precision is $Q = \tau^2 A C^{-1} A$ with $A = \kappa^2 C + G$ on the subject's mesh (lumped mass C, stiffness G; SPDE–GMRF link, Lindgren et al., 2011), $\kappa^2 = 8 / \rho^2$ for range $\rho$, and $\tau^2 = 1 / (4\pi \kappa^2 \sigma^2)$ for marginal s.d. $\sigma$. $\sigma$ is set relative to the data ($\sigma 0$ = median responsive amplitude / dilution, times the amplitude scale of the grid). The normal equations $(K^T D K + Q) f = K^T D y$ are solved by block preconditioned conjugate gradients, all maps at once, with relative tolerance 1e-4. The preconditioner is $Q + \mathrm{diag}(K^T D K \cdot 1)$, factorised by SuperLU with COLAMD ordering; with correlated noise, D is replaced by $D^{1/2} C_\varepsilon^{-1} D^{1/2}$.

**Noise models.**

- **White.** A noise power per voxel and run (HCP: the mean power at the eight frequency bins neighbouring the stimulus frequency, harmonics excluded; StudyForrest and ds002702: the run-to-run variance of the betas), propagated exactly through any emulation, times one calibration factor: observed / propagated variance of the coarse runs, which absorbs spatial noise correlation in the marginal variance only.
- **Correlated.** The same marginal variance times a stationary correlation. Its spectrum comes from the source runs' unit residuals, radially averaged and floored at 0.1 of white, and is applied with FFTs on the full grid. Voxels without cortex are kept as noise references.

**Prior grids (range in mm / amplitude scale).**

- HCP: tuned on 5 subjects. 2.4 mm: 12 / 0.125; 3.2 mm: 24 / 0.25; competitors smoothing 10 mm, upsampling + 8 mm.
- StudyForrest cross-session: 3–24 × 0.125–2.
- StudyForrest within session: white noise on the cross-session grid plus two weaker priors (3 and 6 mm × 4); correlated noise 3–12 × 0.25–4.
- ds002702 ×2 / ×3: correlated 6–24 × 0.5–4, white 12 × 1–2.
- ds002702 ×1.3125: white 3–12 × 0.5–2, correlated 6–12 × 1–4.

The ds002702 grids were widened before summarising, after the first fits had their optimum at the edge of the initial grid.

#### S4. Competitors, scoring and setting selection

##### Competitors.

- Standard mapping of the source, ± surface smoothing (implicit heat equation, FWHM 2–16 mm).
- Within a session: band-limited upsampling (zero-padded k-space) to the native grid, then the native mapping, ± smoothing.
- Across sessions: resampling to the target grid (cubic B-spline, or trilinear), then the target mapping, ± smoothing.

**Score.** Residual energy fraction of the target after the best single scaling of the prediction:

- complex wedge maps: conjugated across directions;
- StudyForrest hemifield maps: real and imaginary parts, each with its own scaling;
- ds002702: each ring map with its own scaling.

**Setting selection.** Every method's setting is chosen by the same rule:

- fixed on separate subjects (HCP);
- leave-one-subject-out (StudyForrest, ds002702 within session);
- split-half on the source alone (StudyForrest, within and across sessions);
- on the other subject of the pair type (ds002702 cross-session, second analysis), or on the other subjects of the pair type (StudyForrest cross-session, first pre-declared analysis).

#### S5. HCP results in full

##### S5.1 Held-out test

| grid | vertices | n per subject | standard mapping | k-space upsampling | standard + surface smoothing | upsampling + 1.6 mm mapping + smoothing | model-based (this work) | model better than smoothing / than upsampling+smoothing (subjects) |
|---|---|---|---|---|---|---|---|---|
| 2.4 mm | all coherent | 26563 | 0.422 ± 0.078 | 0.356 ± 0.095 | 0.341 ± 0.066 | 0.321 ± 0.077 | 0.313 ± 0.078 | 14/15 / 13/15 |

| grid | vertices | n per subject | standard mapping | k-space upsampling | standard + surface smoothing | upsampling + 1.6 mm mapping + smoothing | model-based (this work) | model better than smoothing / than upsampling+smoothing (subjects) |
|---|---|---|---|---|---|---|---|---|
| 2.4 mm | clean at 1.6, leak > 5% | 1233 | 0.477 ± 0.078 | 0.363 ± 0.111 | 0.363 ± 0.074 | 0.335 ± 0.088 | 0.317 ± 0.088 | 14/15 / 14/15 |
| 2.4 mm | clean at 1.6, leak > 10% | 420 | 0.530 ± 0.082 | 0.393 ± 0.113 | 0.399 ± 0.085 | 0.367 ± 0.095 | 0.343 ± 0.092 | 14/15 / 14/15 |
| 2.4 mm | pericalcarine | 2354 | 0.368 ± 0.091 | 0.277 ± 0.112 | 0.269 ± 0.078 | 0.248 ± 0.085 | 0.240 ± 0.085 | 14/15 / 12/15 |
| 3.2 mm | all coherent | 26563 | 0.458 ± 0.066 | 0.380 ± 0.081 | 0.352 ± 0.062 | 0.339 ± 0.071 | 0.325 ± 0.074 | 14/15 / 14/15 |
| 3.2 mm | clean at 1.6, leak > 5% | 2339 | 0.490 ± 0.065 | 0.391 ± 0.085 | 0.370 ± 0.062 | 0.352 ± 0.073 | 0.328 ± 0.078 | 14/15 / 14/15 |
| 3.2 mm | clean at 1.6, leak > 10% | 1121 | 0.534 ± 0.062 | 0.411 ± 0.088 | 0.392 ± 0.067 | 0.371 ± 0.076 | 0.344 ± 0.082 | 14/15 / 14/15 |
| 3.2 mm | pericalcarine | 2354 | 0.409 ± 0.076 | 0.305 ± 0.090 | 0.275 ± 0.074 | 0.263 ± 0.078 | 0.251 ± 0.080 | 14/15 / 14/15 |

Secondary (settings tuned where unmixing is needed, clean at 1.6 mm and leak > 10%):

| grid | vertices | smoothing (high-leak tuned) | model (high-leak tuned) | model better (subjects) |
|---|---|---|---|---|
| 2.4 mm | all coherent | 0.341 ± 0.066 | 0.314 ± 0.080 | 14/15 |
| 2.4 mm | clean at 1.6, leak > 5% | 0.363 ± 0.074 | 0.318 ± 0.091 | 14/15 |
| 2.4 mm | clean at 1.6, leak > 10% | 0.399 ± 0.085 | 0.343 ± 0.096 | 13/15 |
| 2.4 mm | pericalcarine | 0.269 ± 0.078 | 0.240 ± 0.087 | 14/15 |
| 3.2 mm | all coherent | 0.352 ± 0.062 | 0.325 ± 0.074 | 14/15 |
| 3.2 mm | clean at 1.6, leak > 5% | 0.370 ± 0.062 | 0.328 ± 0.078 | 14/15 |
| 3.2 mm | clean at 1.6, leak > 10% | 0.392 ± 0.067 | 0.344 ± 0.082 | 14/15 |
| 3.2 mm | pericalcarine | 0.275 ± 0.074 | 0.251 ± 0.080 | 14/15 |

#### S5.2 Mis-specified PSF

| grid | vertices | model, PSF -15% | model, exact PSF | model, PSF +15% | best competitor (smoothing / upsampling+smoothing) |
|---|---|---|---|---|---|
| 2.4 mm | all coherent | 0.311 ± 0.078 | 0.313 ± 0.078 | 0.318 ± 0.077 | 0.320 ± 0.073 |
| 2.4 mm | clean at 1.6, leak > 5% | 0.315 ± 0.088 | 0.317 ± 0.088 | 0.322 ± 0.086 | 0.334 ± 0.083 |
| 2.4 mm | clean at 1.6, leak > 10% | 0.341 ± 0.092 | 0.343 ± 0.092 | 0.347 ± 0.091 | 0.365 ± 0.090 |
| 2.4 mm | pericalcarine | 0.238 ± 0.086 | 0.240 ± 0.085 | 0.244 ± 0.084 | 0.247 ± 0.081 |
| 3.2 mm | all coherent | 0.322 ± 0.074 | 0.325 ± 0.074 | 0.333 ± 0.074 | 0.337 ± 0.068 |
| 3.2 mm | clean at 1.6, leak > 5% | 0.326 ± 0.077 | 0.328 ± 0.078 | 0.336 ± 0.077 | 0.350 ± 0.069 |
| 3.2 mm | clean at 1.6, leak > 10% | 0.341 ± 0.081 | 0.344 ± 0.082 | 0.352 ± 0.082 | 0.370 ± 0.073 |
| 3.2 mm | pericalcarine | 0.249 ± 0.079 | 0.251 ± 0.080 | 0.257 ± 0.080 | 0.262 ± 0.075 |

#### S5.3 Opposite-bank contrast, averaging kernels, known-truth simulation

**Construction.** Opposite-bank pairs: for each coherent vertex, the nearest coherent vertex (Euclidean distance on the midthickness, ≤ 4 mm) that is far along the sheet (> 15 mm on the scaled sphere); the dissimilarity of a pair is 1 − cos of the polar-angle difference, and the index reported is the dissimilarity of a method's maps divided by that of the 1.6 mm maps. Averaging kernels: 24 sample vertices per subject and grid, a stratified random sample of high-leak and other coherent vertices (`run_subject.py`); the far weight of a kernel is the share of its weights on far cortex (net:

signed; gross: absolute values), its spread the weight-RMS distance of the near weights along the sheet. Known-truth simulation: the truth per run is the two runs' 1.6 mm standard maps averaged (the other run conjugated and phase-aligned), surface-smoothed 3 mm and divided by the dilution of a unit field (floor 0.3), so that it carries little single-run noise; coarse data y = K_g f_true + complex Gaussian noise with the empirical per-voxel noise power; every method is applied to y with its real-data settings (no re-tuning) and scored in the reference domain against the noise-free rendering W_1.6 K_1.6 f_true.

| grid | pairs | standard mapping | k-space upsampling | standard + surface smoothing | upsampling + 1.6 mm mapping + smoothing | model-based (this work) |
|---|---|---|---|---|---|---|
| 2.4 mm | all coherent pairs | 0.61 ± 0.06 | 0.68 ± 0.08 | 0.48 ± 0.08 | 0.55 ± 0.09 | 0.75 ± 0.09 |
| 2.4 mm | pericalcarine pairs | 0.71 ± 0.12 | 0.78 ± 0.10 | 0.65 ± 0.19 | 0.70 ± 0.20 | 0.83 ± 0.18 |
| 3.2 mm | all coherent pairs | 0.65 ± 0.07 | 0.52 ± 0.07 | 0.42 ± 0.07 | 0.48 ± 0.08 | 0.72 ± 0.08 |
| 3.2 mm | pericalcarine pairs | 0.80 ± 0.12 | 0.62 ± 0.09 | 0.59 ± 0.17 | 0.63 ± 0.18 | 0.81 ± 0.16 |

Pairs per subject: 4067 (pericalcarine 598).

| grid | vertices | net leak, standard | net leak, model | gross far weight, standard | gross far weight, model | spread along the sheet (mm), standard | spread, model |
|---|---|---|---|---|---|---|---|
| 2.4 mm | 360 | 11.3 ± 11.5% | 0.1 ± 1.2% | 18.8 ± 12.0% | 13.6 ± 6.9% | 3.74 ± 0.32 | 4.62 ± 0.48 |
| 3.2 mm | 360 | 12.8 ± 16.2% | -1.1 ± 2.8% | 26.0 ± 12.8% | 22.9 ± 8.1% | 4.64 ± 0.39 | 5.49 ± 0.54 |

| grid | vertices | standard mapping | k-space upsampling | standard + surface smoothing | upsampling + 1.6 mm mapping + smoothing | model-based (this work) |
|---|---|---|---|---|---|---|
| 2.4 mm | all coherent | 0.0814 ± 0.0117 | 0.0278 ± 0.0066 | 0.0665 ± 0.0043 | 0.0492 ± 0.0033 | 0.0293 ± 0.0081 |
| 2.4 mm | clean at 1.6, leak > 5% | 0.1131 ± 0.0344 | 0.0320 ± 0.0099 | 0.0863 ± 0.0186 | 0.0660 ± 0.0137 | 0.0344 ± 0.0106 |
| 2.4 mm | clean at 1.6, leak > 10% | 0.1337 ± 0.0536 | 0.0320 ± 0.0108 | 0.1010 ± 0.0292 | 0.0784 ± 0.0194 | 0.0387 ± 0.0127 |
| 2.4 mm | pericalcarine | 0.0795 ± 0.0197 | 0.0219 ± 0.0060 | 0.0548 ± 0.0131 | 0.0417 ± 0.0098 | 0.0269 ± 0.0123 |
| 3.2 mm | all coherent | 0.1513 ± 0.0191 | 0.0569 ± 0.0099 | 0.0761 ± 0.0055 | 0.0591 ± 0.0044 | 0.0371 ± 0.0088 |
| 3.2 mm | clean at 1.6, leak > 5% | 0.1438 ± 0.0264 | 0.0570 ± 0.0124 | 0.0875 ± 0.0159 | 0.0684 ± 0.0132 | 0.0402 ± 0.0105 |
| 3.2 mm | clean at 1.6, leak > 10% | 0.1676 ± 0.0366 | 0.0601 ± 0.0149 | 0.0971 ± 0.0223 | 0.0764 ± 0.0181 | 0.0443 ± 0.0115 |
| 3.2 mm | pericalcarine | 0.1376 ± 0.0231 | 0.0433 ± 0.0097 | 0.0588 ± 0.0130 | 0.0476 ± 0.0104 | 0.0325 ± 0.0118 |

#### S5.4 Misregistration, depth, uncertainty, theory link

##### (b) Misregistration

The coarse data were given a rigid registration error of 0.5 or 1.0 mm (one random direction per subject). The model inverted with surfaces shifted by that amount; the competitors received the same error (the coarse data resampled at x + d with a band-limited Fourier shift, which is exactly what an inversion with geometry P + d sees). Rendering and scoring stay nominal. Cross-run residual against the other run's 1.6 mm map, all coherent vertices (lower is better):

| grid | error | standard | standard + smoothing (10 mm) | upsampling + 1.6 mm mapping + smoothing (8 mm) | model | model − upsampling+smoothing, same error (subjects where model better) |
|---|---|---|---|---|---|---|
| 2.4 mm | 0 | 0.422 | 0.341 | 0.321 | 0.313 | −0.008 (13/15) |
| 2.4 mm | 0.5 mm | 0.429 | 0.345 | 0.328 | 0.320 | −0.008 (14/15) |
| 2.4 mm | 1 mm | 0.452 | 0.356 | 0.346 | 0.338 | −0.008 (14/15) |
| 3.2 mm | 0 | 0.458 | 0.352 | 0.339 | 0.325 | −0.014 (14/15) |
| 3.2 mm | 0.5 mm | 0.462 | 0.354 | 0.342 | 0.330 | −0.013 (14/15) |
| 3.2 mm | 1 mm | 0.473 | 0.362 | 0.354 | 0.344 | −0.009 (14/15) |

Where unmixing is needed (clean at 1.6 mm, leak > 10%) the model's lead over upsampling + smoothing is −0.024 / −0.022 / −0.018 at 2.4 mm and −0.027 / −0.025 / −0.020 at 3.2 mm for 0 / 0.5 / 1 mm (13–14/15 subjects each). In the pericalcarine stratum it shrinks from −0.008 to −0.004 (9/15) at 2.4 mm and 1 mm error. Full tables with standard errors: `hcp_report_section.md`.

**(b) Depth-varying signal**

A second field b1 (white-to-pial gradient $\sqrt{3}(2\zeta - 1)$, orthogonalised to the column mean under each column's volume measure) with its own Matérn prior (s.d. 0.5 × that of b0):

| grid | b1/b0, coherent | positive fraction | b1/b0, pericalcarine | pial / white amplitude implied | residual change vs b0-only model: all coherent / leak > 10% (subjects better) |
|---|---|---|---|---|---|
| 2.4 mm | 0.239 ± 0.060 | 0.93 | 0.263 ± 0.080 | 2.4 | −0.0048 (15/15) / −0.0085 (14/15) |
| 3.2 mm | 0.335 ± 0.067 | 0.92 | 0.367 ± 0.121 | 3.8 | −0.0064 (15/15) / −0.0090 (14/15) |

The per-subject b1/b0 ratios at the two grids correlate r = 0.86 (15 subjects), so the estimate is subject-specific and reproducible, but its size depends on the grid.

**(c) Uncertainty calibration**

Noise part: run-to-run differences of the estimates (CW run conjugated and scaled onto CCW) against the predicted noise variance, in noise-dominated vertices (coherence < 0.1). Posterior part: coverage of the known truth in the simulation of S5.3.

| grid | quantity (nominal) | model | standard mapping |
|---|---|---|---|
| 2.4 mm | observed / predicted variance (1) | 5.17 ± 0.66 (MC-corrected) | 1.74 ± 0.14 |
| 2.4 mm | beyond the 95% bound (0.05) | 0.556 | 0.161 |
| 2.4 mm | same, after one variance factor estimated on the other hemisphere (0.05) | 0.060 | 0.052 |

| grid | quantity (nominal) | model | standard mapping |
|---|---|---|---|
| 3.2 mm | observed / predicted variance (1) | 4.40 ± 0.62 (MC-corrected) | 1.85 ± 0.19 |
| 3.2 mm | beyond the 95% bound (0.05) | 0.500 | 0.160 |
| 3.2 mm | same, after one variance factor estimated on the other hemisphere (0.05) | 0.061 | 0.052 |
| 2.4 mm | simulation: posterior coverage 50 / 80 / 95%, all vertices | 0.47 / 0.71 / 0.84 | – |
| 2.4 mm | simulation: posterior coverage 50 / 80 / 95%, coherent vertices | 0.08 / 0.16 / 0.26 | – |
| 3.2 mm | simulation: posterior coverage 50 / 80 / 95%, all vertices | 0.48 / 0.72 / 0.85 | – |
| 3.2 mm | simulation: posterior coverage 50 / 80 / 95%, coherent vertices | 0.09 / 0.18 / 0.28 | – |

In coherent vertices the run-to-run differences are dominated by direction-dependent signal (the haemodynamic delay varies across cortex, so CW is not an exact conjugate of CCW) and cannot test the noise model (observed/predicted 31–91).

###### (d) Averaging-kernel spread vs the resolvable wavelength (S10)

Spearman correlation, per subject, between the kernel spread along the sheet at the 24 kernel sample vertices of S5.3 and the orientation-dependent resolvable wavelength of S10 at the same vertices (3.5–4.8 mm at 2.4 mm; 4.6–6.3 mm at 3.2 mm): model $\rho = +0.03 \pm 0.21$ (2.4 mm) and $-0.03 \pm 0.14$ (3.2 mm), mean ± SD over 15 subjects (SE 0.06 / 0.04); standard-mapping kernels −0.04 / +0.02. No relation.

Figures: `figW4-1_misregistration_depth.png`, `figW4-2_uncertainty.png`.

##### S5.5 Dose-response on far leak

| mixing at the coarse grid | HCP 2.4 mm (far leak) | HCP 3.2 mm (far leak) | StudyForrest, white noise (other hemisphere) | StudyForrest, correlated noise (other hemisphere) |
|---|---|---|---|---|
| 0–2% | −0.007 (13/15) | −0.016 (14/15) | +0.011 (9/28) | +0.010 (10/28) |
| 2–5% | −0.009 (12/15) | −0.013 (14/15) | +0.026 (4/28) | +0.025 (3/28) |
| 5–10% | −0.013 (12/15) | −0.017 (14/15) | +0.022 (5/26) | +0.013 (11/26) |
| 10–20% | −0.017 (13/15) | −0.017 (13/15) | +0.009 (12/26) | +0.007 (13/26) |
| > 20% | −0.035 (12/14) | −0.038 (14/15) | −0.009 (12/20) | −0.026 (16/20) |

##### S5.6 Leak-gated estimator

| grid | vertices | competitor everywhere | model everywhere | gated (τ LOSO) | gated − competitor (better) | gated − model everywhere (better) |
|---|---|---|---|---|---|---|
| 2.4 mm | all coherent | 0.3212 | 0.3131 | 0.3172 | −0.0041 (12/15) | +0.0041 (1/15) |
| 2.4 mm | clean at 1.6, leak > 10% | 0.3671 | 0.3428 | 0.3428 | −0.0243 (14/15) | −0.0000 (2/15) |
| 3.2 mm | all coherent | 0.3385 | 0.3248 | 0.3326 | −0.0059 (13/15) | +0.0078 (0/15) |
| 3.2 mm | clean at 1.6, leak > 10% | 0.3712 | 0.3439 | 0.3439 | −0.0272 (14/15) | −0.0000 (2/15) |

##### S5.7 The gain in degrees of polar angle (15 development subjects)

Mean over subjects of the per-subject mean absolute polar-angle error (degrees) of the cross-run prediction against the other run's 1.6 mm map, coherent vertices weighted equally, averaged over the two wedge directions; ± SE over subjects. Strata as in the far-leak dose-response (coherent, clean at 1.6 mm, binned by the far leak at the coarse grid). The last column is the run-to-run disagreement of the two 1.6 mm maps themselves. Δ = model − upsampling + 8 mm smoothing, with the subjects in which the model has the smaller error. Source: `src/14_robustness_checks/angle_error.py`; per-subject files in `angle_error/`.

##### Emulated 2.4 mm

| stratum | n vertices (mean) | standard mapping | standard + smoothing 10 mm | upsampling + 1.6 mm mapping | upsampling + 1.6 mm mapping + smoothing 8 mm | model | 1.6 mm maps, run against run | Δ model − upsample+smooth | model better |
|---|---|---|---|---|---|---|---|---|---|
| all coherent vertices (coherence > 0.3) | 26563 | 31.5 ± 1.4 | 31.2 ± 1.3 | 30.6 ± 1.5 | 30.3 ± 1.4 | 29.7 ± 1.4 | 31.6 ± 1.5 | −0.6 ± 0.1 | 14/15 |
| pericalcarine | 2354 | 28.6 ± 2.2 | 29.8 ± 2.1 | 27.7 ± 2.2 | 28.5 ± 2.2 | 27.6 ± 2.2 | 29.0 ± 2.2 | −1.0 ± 0.3 | 11/15 |
| far leak 0–2% | 6903 | 30.7 ± 1.4 | 30.3 ± 1.3 | 29.9 ± 1.4 | 29.4 ± 1.3 | 28.9 ± 1.4 | 30.9 ± 1.4 | −0.5 ± 0.1 | 13/15 |
| 2–5% | 3072 | 29.9 ± 1.3 | 30.2 ± 1.3 | 29.1 ± 1.4 | 29.2 ± 1.3 | 28.5 ± 1.3 | 29.8 ± 1.3 | −0.7 ± 0.1 | 14/15 |
| 5–10% | 812 | 31.4 ± 1.6 | 31.9 ± 1.6 | 30.1 ± 1.7 | 30.6 ± 1.7 | 29.5 ± 1.6 | 31.4 ± 1.8 | −1.1 ± 0.3 | 13/15 |
| 10–20% | 304 | 35.1 ± 1.5 | 34.5 ± 1.5 | 33.0 ± 1.4 | 33.4 ± 1.5 | 32.5 ± 1.4 | 33.5 ± 1.6 | −0.9 ± 0.5 | 11/15 |
| > 20% | 117 | 36.6 ± 2.0 | 35.2 ± 2.3 | 32.4 ± 2.2 | 33.8 ± 2.3 | 31.9 ± 2.2 | 35.2 ± 2.4 | −1.9 ± 0.7 | 11/14 |
| > 10% | 420 | 35.6 ± 1.5 | 34.9 ± 1.6 | 32.9 ± 1.5 | 33.7 ± 1.6 | 32.5 ± 1.5 | 34.0 ± 1.7 | −1.3 ± 0.4 | 13/15 |
| strongly coherent vertices (coherence > 0.5) | 16055 | 22.6 ± 1.0 | 22.0 ± 0.9 | 21.7 ± 1.1 | 21.3 ± 0.9 | 20.8 ± 1.0 | 22.8 ± 1.1 | −0.5 ± 0.1 | 15/15 |
| coherence > 0.5, clean at 1.6 mm, far leak > 10% | 271 | 27.5 ± 1.1 | 25.6 ± 1.2 | 24.5 ± 1.1 | 24.9 ± 1.2 | 23.3 ± 1.1 | 26.2 ± 1.2 | −1.6 ± 0.3 | 12/15 |

##### Emulated 3.2 mm

| stratum | n vertices (mean) | standard mapping | standard + smoothing 10 mm | upsampling + 1.6 mm mapping | upsampling + 1.6 mm mapping + smoothing 8 mm | model | 1.6 mm maps, run against run | Δ model − upsample+smooth | model better |
|---|---|---|---|---|---|---|---|---|---|
| all coherent vertices (coherence > 0.3) | 26563 | 32.7 ± 1.4 | 31.6 ± 1.3 | 30.9 ± 1.4 | 30.8 ± 1.4 | 30.3 ± 1.4 | 31.6 ± 1.5 | −0.5 ± 0.1 | 13/15 |
| pericalcarine | 2354 | 30.5 ± 2.1 | 30.0 ± 2.1 | 27.8 ± 2.1 | 28.7 ± 2.1 | 27.8 ± 2.1 | 29.0 ± 2.2 | −0.9 ± 0.3 | 10/15 |
| far leak 0–2% | 5219 | 32.8 ± 1.3 | 30.7 ± 1.2 | 30.2 ± 1.4 | 29.9 ± 1.3 | 29.5 ± 1.3 | 31.1 ± 1.4 | −0.5 ± 0.1 | 14/15 |
| 2–5% | 3242 | 30.7 ± 1.3 | 30.2 ± 1.3 | 29.7 ± 1.4 | 29.3 ± 1.3 | 29.0 ± 1.3 | 30.2 ± 1.3 | −0.3 ± 0.1 | 12/15 |
| 5–10% | 1218 | 32.2 ± 1.6 | 31.6 ± 1.6 | 30.7 ± 1.6 | 30.7 ± 1.6 | 30.2 ± 1.6 | 31.3 ± 1.7 | −0.6 ± 0.1 | 13/15 |
| 10–20% | 676 | 34.8 ± 1.4 | 34.4 ± 1.4 | 32.8 ± 1.5 | 33.3 ± 1.4 | 32.3 ± 1.4 | 33.8 ± 1.7 | −1.1 ± 0.2 | 15/15 |
| > 20% | 445 | 38.9 ± 1.2 | 36.4 ± 1.5 | 34.5 ± 1.5 | 35.2 ± 1.5 | 33.1 ± 1.6 | 34.7 ± 1.6 | −2.1 ± 0.5 | 14/15 |

| stratum | n vertices (mean) | standard mapping | standard + smoothing 10 mm | upsampling + 1.6 mm mapping | upsampling + 1.6 mm mapping + smoothing 8 mm | model | 1.6 mm maps, run against run | Δ model − upsample+smooth | model better |
|---|---|---|---|---|---|---|---|---|---|
| > 10% | 1121 | 36.4 ± 1.3 | 35.2 ± 1.3 | 33.5 ± 1.4 | 34.1 ± 1.4 | 32.6 ± 1.4 | 34.2 ± 1.6 | −1.5 ± 0.3 | 15/15 |
| strongly coherent vertices (coherence > 0.5) | 16055 | 24.0 ± 1.0 | 22.4 ± 0.9 | 22.0 ± 1.0 | 21.7 ± 0.9 | 21.2 ± 0.9 | 22.8 ± 1.1 | −0.5 ± 0.1 | 15/15 |
| coherence > 0.5, clean at 1.6 mm, far leak > 10% | 686 | 27.4 ± 0.9 | 24.8 ± 0.9 | 24.3 ± 1.0 | 24.1 ± 0.9 | 22.9 ± 1.0 | 25.5 ± 1.1 | −1.2 ± 0.2 | 14/15 |

#### Medians and the share of vertices within 30° (all coherent vertices)

| grid | method | median error (°) | within 30° |
|---|---|---|---|
| 2.4 mm | standard mapping | 21.4 | 64% |
| 2.4 mm | standard + smoothing 10 mm | 20.9 | 65% |
| 2.4 mm | upsampling + 1.6 mm mapping | 21.0 | 65% |
| 2.4 mm | upsampling + 1.6 mm mapping + smoothing 8 mm | 20.3 | 66% |
| 2.4 mm | model | 20.1 | 66% |
| 2.4 mm | 1.6 mm maps, run against run | 21.2 | 64% |
| 3.2 mm | standard mapping | 21.8 | 63% |
| 3.2 mm | standard + smoothing 10 mm | 21.2 | 64% |
| 3.2 mm | upsampling + 1.6 mm mapping | 21.3 | 64% |
| 3.2 mm | upsampling + 1.6 mm mapping + smoothing 8 mm | 20.6 | 65% |
| 3.2 mm | model | 20.5 | 65% |
| 3.2 mm | 1.6 mm maps, run against run | 21.2 | 64% |

#### S5.8 Noise ceiling: the other 1.6 mm run as the predictor, and the gap the model closes

Cross-run residual against the other run's 1.6 mm map (mean over subjects). "Ceiling": the other 1.6 mm run itself as the predictor, raw or surface-smoothed. Competitors and model predict from the coarsened run (Section 4.3). Gap closed = (competitor − model) / (competitor − ceiling) from the means; in brackets the median over subjects of the per-subject ratio. Source: `src/14_robustness_checks/ceiling.py`.

| grid | vertex set | ceiling raw | ceiling 4 mm | ceiling 6 mm | upsampling + smoothing 8 mm | per-subject best competitor | model | gap closed vs 6 mm ceiling (competitor = upsampling + smoothing) | gap closed (per-subject best competitor) |
|---|---|---|---|---|---|---|---|---|---|
| 2.4 mm | all coherent vertices | 0.362 | 0.311 | 0.305 | 0.321 | 0.320 | 0.313 | 51% [46%] | 47% [46%] |
| 2.4 mm | clean at 1.6 mm, far leak > 5% | 0.369 | 0.315 | 0.312 | 0.335 | 0.334 | 0.317 | 76% [58%] | 74% [58%] |
| 2.4 mm | clean at 1.6 mm, far leak > 10% | 0.408 | 0.348 | 0.344 | 0.367 | 0.365 | 0.343 | 107% [82%] | 107% [82%] |
| 2.4 mm | pericalcarine | 0.281 | 0.235 | 0.231 | 0.248 | 0.247 | 0.240 | 48% [35%] | 44% [38%] |
| 3.2 mm | all coherent vertices | 0.362 | 0.311 | 0.305 | 0.339 | 0.337 | 0.325 | 41% [40%] | 39% [41%] |
| 3.2 mm | clean at 1.6 mm, far leak > 5% | 0.377 | 0.318 | 0.312 | 0.352 | 0.350 | 0.328 | 59% [52%] | 57% [57%] |
| 3.2 mm | clean at 1.6 mm, far leak > 10% | 0.402 | 0.337 | 0.331 | 0.371 | 0.370 | 0.344 | 68% [61%] | 67% [62%] |

| grid | vertex set | ceiling raw | ceiling 4 mm | ceiling 6 mm | upsampling + smoothing 8 mm | per-subject best competitor | model | gap closed vs 6 mm ceiling (competitor = upsampling + smoothing) | gap closed (per-subject best competitor) |
|---|---|---|---|---|---|---|---|---|---|
| 3.2 mm | pericalcarine | 0.281 | 0.235 | 0.231 | 0.263 | 0.262 | 0.251 | 38% [36%] | 37% [36%] |

#### S6. StudyForrest

##### S6.1 Cross-session, settings fixed from HCP and re-tuned

**Data.** StudyForrest multi-resolution visual-orientation study (7T: 0.8 / 1.4 / 2.0 / 3.0 mm sessions, 10 runs each; oriented gratings in the left and/or right visual hemifield, 3-s trials; 3T: 1.4 / 2.0 / 3.0 mm in 5 of the 7 subjects), the subjects' FreeSurfer surfaces; same people and stimulus. Left- and right-hemifield GLM betas (% signal change) per session; run-to-run variance of the mean as the voxel noise. Every session is modelled on its own native grid.

**Processing.** Per session: motion correction to run 1, session mean, bbregister (6 dof) to the FreeSurfer anatomy, FLIRT-initialised; sessions still above cost 0.6 are re-initialised from the subject's best-registered session (same-subject EPI-to-EPI, then bbregister), and from the best session of the same scanner if still above 0.6. Slab-edge voxels whose mean in any run falls below half the session mean are masked (zero-filled after motion correction; they made the percent-change betas explode). Pairs involving a session whose final cost stays above 0.6 would be excluded; after the anchoring passes none does (final costs 0.07–0.58, listed below).

**Methods compared** (each source predicts the target session's standard ribbon-mapped map; residual energy fraction after the best real scaling, pooled over the two hemifield maps; lower is better):

- standard: the source's own standard map;
- standard + smoothing: surface smoothing, FWHM 10 mm (HCP choice);
- resample + smoothing: source volume resampled onto the target grid, target mapping, smoothing 8 mm (HCP choice);
- model: posterior mean from the source with the source's acquisition PSF (echo spacing / acceleration from the sidecars; 7T echo spacings not published, assumed), prior fixed from HCP by source resolution (1.4 / 2.0 mm: 12 mm, 0.125; 3.0 mm: 24 mm, 0.25), rendered through the target's operator and mapping ($W_t K_t f$).

Vertex sets: covered by both sessions (partial-volume dilution > 0.5), and its visual-cortex subset (pericalcarine, cuneus, lingual, lateral occipital; the hemifield stimulus drives these). Reference point: the target's odd-run map predicting its even-run map (half the data on both sides, so this is a noisy reference, not a ceiling).

**Pre-specified comparison (settings fixed from HCP), visual vertices**

| pair | subjects | vertices | standard mapping (source) | standard + smoothing (10 mm) | resample to target + smoothing (8 mm) | model-based | target's odd vs even runs | model − smoothing (better) | model − resample+smoothing (better) |
|---|---|---|---|---|---|---|---|---|---|
| 7T-r30 → 7T-r08 | 7 | 19186 | 0.545 ± 0.109 | 0.487 ± 0.117 | 0.484 ± 0.116 | 0.499 ± 0.120 | 0.526 ± 0.197 | +0.012 (3/7) | +0.015 (3/7) |
| 7T-r30 → 7T-r14 | 7 | 23770 | 0.383 ± 0.096 | 0.355 ± 0.087 | 0.353 ± 0.086 | 0.395 ± 0.071 | 0.168 ± 0.112 | +0.040 (0/7) | +0.042 (0/7) |
| 7T-r30 → 7T-r20 | 7 | 24101 | 0.336 ± 0.104 | 0.320 ± 0.107 | 0.318 ± 0.106 | 0.374 ± 0.121 | 0.151 ± 0.078 | +0.054 (0/7) | +0.056 (0/7) |
| 7T-r20 → 7T-r08 | 7 | 25015 | 0.496 ± 0.140 | 0.457 ± 0.157 | 0.452 ± 0.158 | 0.479 ± 0.161 | 0.520 ± 0.202 | +0.022 (1/7) | +0.027 (1/7) |
| 7T-r20 → 7T-r14 | 7 | 30855 | 0.259 ± 0.034 | 0.278 ± 0.036 | 0.269 ± 0.037 | 0.349 ± 0.048 | 0.171 ± 0.113 | +0.071 (0/7) | +0.080 (0/7) |
| 7T-r14 → 7T-r08 | 7 | 26564 | 0.491 ± 0.127 | 0.456 ± 0.146 | 0.450 ± 0.146 | 0.467 ± 0.146 | 0.518 ± 0.203 | +0.011 (2/7) | +0.017 (1/7) |
| 3T-r30 → 7T-r08 | 5 | 20109 | 0.617 ± 0.146 | 0.526 ± 0.181 | 0.521 ± 0.182 | 0.535 ± 0.173 | 0.554 ± 0.231 | +0.009 (3/5) | +0.014 (2/5) |
| 3T-r30 → 7T-r14 | 5 | 25407 | 0.483 ± 0.055 | 0.384 ± 0.038 | 0.383 ± 0.037 | 0.418 ± 0.049 | 0.179 ± 0.138 | +0.034 (0/5) | +0.035 (1/5) |
| 3T-r30 → 7T-r20 | 5 | 25763 | 0.447 ± 0.019 | 0.356 ± 0.060 | 0.356 ± 0.053 | 0.409 ± 0.068 | 0.135 ± 0.078 | +0.053 (0/5) | +0.053 (0/5) |
| 3T-r30 → 7T-r30 | 5 | 21822 | 0.480 ± 0.123 | 0.384 ± 0.145 | 0.376 ± 0.145 | 0.455 ± 0.141 | 0.225 ± 0.178 | +0.071 (0/5) | +0.079 (0/5) |
| 3T-r20 → 7T-r08 | 5 | 25463 | 0.589 ± 0.154 | 0.502 ± 0.184 | 0.497 ± 0.183 | 0.519 ± 0.180 | 0.551 ± 0.234 | +0.016 (2/5) | +0.022 (1/5) |
| 3T-r20 → 7T-r14 | 5 | 32060 | 0.473 ± 0.072 | 0.378 ± 0.054 | 0.373 ± 0.051 | 0.417 ± 0.068 | 0.181 ± 0.141 | +0.039 (0/5) | +0.045 (0/5) |
| 3T-r20 → 7T-r20 | 5 | 31949 | 0.447 ± 0.050 | 0.356 ± 0.078 | 0.347 ± 0.071 | 0.417 ± 0.075 | 0.137 ± 0.078 | +0.061 (0/5) | +0.071 (0/5) |
| 3T-r20 → 7T-r30 | 5 | 25163 | 0.512 ± 0.106 | 0.400 ± 0.128 | 0.382 ± 0.127 | 0.462 ± 0.121 | 0.225 ± 0.177 | +0.062 (0/5) | +0.080 (0/5) |
| 3T-r14 → 7T-r08 | 5 | 26798 | 0.650 ± 0.145 | 0.503 ± 0.194 | 0.504 ± 0.192 | 0.515 ± 0.194 | 0.546 ± 0.237 | +0.012 (0/5) | +0.011 (2/5) |
| 3T-r14 → 7T-r14 | 5 | 32861 | 0.584 ± 0.062 | 0.391 ± 0.060 | 0.388 ± 0.058 | 0.418 ± 0.071 | 0.174 ± 0.138 | +0.027 (0/5) | +0.031 (0/5) |
| 3T-r14 → 7T-r20 | 5 | 31917 | 0.579 ± 0.071 | 0.387 ± 0.087 | 0.380 ± 0.085 | 0.421 ± 0.091 | 0.134 ± 0.079 | +0.034 (0/5) | +0.041 (0/5) |
| 3T-r14 → 7T-r30 | 5 | 24712 | 0.651 ± 0.101 | 0.439 ± 0.136 | 0.451 ± 0.132 | 0.462 ± 0.136 | 0.222 ± 0.176 | +0.023 (0/5) | +0.011 (1/5) |

Pooled over pairs and subjects (pre-specified settings): model − smoothing / model − resample+smoothing, mean (subject-pairs where the model is better):

- 7T→7T, covered: +0.027 (7/42) / +0.031 (7/42)

- 3T→7T, covered: +0.034 (5/60) / +0.037 (5/60)
- all, covered: +0.031 (12/102) / +0.034 (12/102)
- 7T→7T, visual: +0.035 (6/42) / +0.039 (5/42)
- 3T→7T, visual: +0.037 (5/60) / +0.041 (7/60)
- all, visual: +0.036 (11/102) / +0.040 (12/102)

Covered-vertex and pericalcarine tables: `sf/sf_report_section.md`. Figure: `sf/figW4-3_sf_prediction.png` (differences to standard + smoothing, one dot per subject); example maps `sf/figW4-6_sf_example_maps.png` (sub-18, 3.0 → 1.4 mm: both HCP-tuned predictions lose the fine structure the 1.4 mm target shows).

###### How much of the residual is resolution?

Best competitor (smoothing or resample + smoothing), visual vertices: 3T source predicting the 7T session at the same voxel size vs predicting finer 7T sessions (the 0.8 mm target is left out: its own noise dominates).

| 3T source | → 7T same voxel size | → finer 7T targets | target odd vs even (same-size target) |
|---|---|---|---|
| 3T 3.0 mm | 0.376 | 7T 1.4 mm: 0.383, 7T 2.0 mm: 0.356 | 0.225 |
| 3T 2.0 mm | 0.347 | 7T 1.4 mm: 0.373 | 0.137 |
| 3T 1.4 mm | 0.388 | – | 0.174 |

###### Exploratory: settings re-tuned on StudyForrest (leave-one-subject-out), visual vertices

Smoothing FWHM 2–16 mm; model prior range 3 / 6 / 12 / 24 mm × s.d. scale 0.125–2. Each held-out subject is scored with the setting that is best on the other subjects (same pair and vertex set).

| pair | vertices | standard + smoothing | resample + smoothing | model | model better than smoothing / resample+smoothing | settings chosen most often (smoothing; resample; model) |
|---|---|---|---|---|---|---|
| 7T-r30 → 7T-r08 | visual | 0.488 ± 0.117 | 0.484 ± 0.117 | 0.490 ± 0.118 | 5/7 / 4/7 | smooth10; resample_smooth6; r12_s0.5 |
| 7T-r30 → 7T-r14 | visual | 0.338 ± 0.094 | 0.334 ± 0.092 | 0.345 ± 0.071 | 2/7 / 2/7 | smooth4; resample_smooth2; r6_s1 |
| 7T-r30 → 7T-r20 | visual | 0.294 ± 0.106 | 0.287 ± 0.105 | 0.303 ± 0.101 | 4/7 / 3/7 | smooth4; resample_smooth2; r6_s1 |
| 7T-r20 → 7T-r08 | visual | 0.450 ± 0.155 | 0.445 ± 0.156 | 0.445 ± 0.163 | 4/7 / 4/7 | smooth6; resample_smooth4; r6_s0.5 |
| 7T-r20 → 7T-r14 | visual | 0.222 ± 0.032 | 0.218 ± 0.035 | 0.244 ± 0.039 | 1/7 / 1/7 | smooth3; resample_smooth2; r3_s1 |
| 7T-r14 → 7T-r08 | visual | 0.448 ± 0.143 | 0.442 ± 0.144 | 0.445 ± 0.146 | 4/7 / 3/7 | smooth6; resample_smooth4; r6_s0.5 |
| 3T-r30 → 7T-r08 | visual | 0.526 ± 0.182 | 0.521 ± 0.182 | 0.532 ± 0.175 | 3/5 / 2/5 | smooth12; resample_smooth8; r12_s0.25 |
| 3T-r30 → 7T-r14 | visual | 0.382 ± 0.039 | 0.382 ± 0.040 | 0.403 ± 0.052 | 1/5 / 1/5 | smooth8; resample_smooth6; r12_s0.5 |
| 3T-r30 → 7T-r20 | visual | 0.353 ± 0.050 | 0.348 ± 0.034 | 0.381 ± 0.059 | 0/5 / 1/5 | smooth6; resample_smooth4; r12_s1 |
| 3T-r30 → 7T-r30 | visual | 0.380 ± 0.144 | 0.361 ± 0.142 | 0.411 ± 0.148 | 0/5 / 0/5 | smooth6; resample_smooth3; r3_s2 |

| pair | vertices | standard + smoothing | resample + smoothing | model | model better than smoothing / resample+smoothing | settings chosen most often (smoothing; resample; model) |
|---|---|---|---|---|---|---|
| 3T-r20 → 7T-r08 | visual | 0.503 ± 0.184 | 0.497 ± 0.183 | 0.514 ± 0.180 | 2/5 / 2/5 | smooth10; resample_smooth8; r12_s0.25 |
| 3T-r20 → 7T-r14 | visual | 0.377 ± 0.056 | 0.370 ± 0.054 | 0.399 ± 0.072 | 1/5 / 1/5 | smooth6; resample_smooth6; r12_s0.5 |
| 3T-r20 → 7T-r20 | visual | 0.348 ± 0.068 | 0.337 ± 0.058 | 0.392 ± 0.071 | 0/5 / 0/5 | smooth6; resample_smooth4; r12_s1 |
| 3T-r20 → 7T-r30 | visual | 0.397 ± 0.126 | 0.378 ± 0.127 | 0.422 ± 0.132 | 0/5 / 0/5 | smooth8; resample_smooth6; r3_s2 |
| 3T-r14 → 7T-r08 | visual | 0.504 ± 0.194 | 0.504 ± 0.193 | 0.517 ± 0.192 | 0/5 / 2/5 | smooth10; resample_smooth10; r24_s0.25 |
| 3T-r14 → 7T-r14 | visual | 0.391 ± 0.060 | 0.388 ± 0.058 | 0.416 ± 0.072 | 0/5 / 0/5 | smooth10; resample_smooth8; r12_s0.25 |
| 3T-r14 → 7T-r20 | visual | 0.389 ± 0.084 | 0.382 ± 0.082 | 0.415 ± 0.093 | 0/5 / 0/5 | smooth10; resample_smooth8; r12_s0.25 |
| 3T-r14 → 7T-r30 | visual | 0.438 ± 0.137 | 0.442 ± 0.133 | 0.460 ± 0.138 | 0/5 / 0/5 | smooth12; resample_smooth12; r12_s0.25 |

Pooled over pairs and subjects (re-tuned, leave-one-subject-out): model − smoothing / model − resample+smoothing, mean (subject-pairs where the model is better):

- 3T→7T, covered: +0.031 (3/60) / +0.036 (2/60)
- 3T→7T, visual: +0.023 (7/60) / +0.029 (9/60)
- 7T→7T, covered: +0.008 (17/42) / +0.012 (15/42)
- 7T→7T, visual: +0.006 (20/42) / +0.010 (17/42)
- all, covered: +0.022 (20/102) / +0.026 (17/102)
- all, visual: +0.016 (27/102) / +0.021 (26/102)

Smoothing-width curves (each method at every setting, mean over subjects): `sf/sf_report_section.md`; figures `sf/figW4-4_sf_fwhm_curves_exploratory.png`, `sf/figW4-5_sf_retuned_exploratory.png`. Post-smoothing the model never helps (best 0–2 mm), so its HCP estimate was already too smooth rather than too noisy.

**Registration (bbregister cost after within-subject anchoring; lower is better)**

| subject | 7T-r08 | 7T-r14 | 7T-r20 | 7T-r30 | 3T-r14 | 3T-r20 | 3T-r30 |
|---|---|---|---|---|---|---|---|
| sub-04 | 0.25 (a) | 0.26 (a) | 0.27 | 0.51 | – | – | – |
| sub-06 | 0.24 (a) | 0.21 (a) | 0.23 (a) | 0.40 | 0.07 (a) | 0.13 (a) | 0.29 |
| sub-16 | 0.36 (a) | 0.47 (a) | 0.35 (a) | 0.50 | 0.23 | 0.30 | 0.46 |
| sub-17 | 0.32 (a) | 0.28 | 0.33 | 0.53 | 0.19 (a) | 0.25 (a) | 0.47 |
| sub-18 | 0.38 (a) | 0.31 (a) | 0.38 (a) | 0.54 | – | – | – |
| sub-20 | 0.39 (a) | 0.52 (a) | 0.45 | 0.58 | 0.20 | 0.24 | 0.43 |
| sub-21 | 0.45 | 0.32 (a) | 0.38 (a) | 0.58 | 0.22 | 0.26 | 0.49 |

(a) = improved by anchoring to the subject's best-registered session (sub-16 7T 0.8 mm: to its best 7T session, 0.91 → 0.36).

##### S6.2 Within session

1. **Within one session the model still does not beat simple methods overall.** There are no registration, distortion, day or scanner differences, only the loss of resolution. In visual cortex the model's residual is +0.016 (settings tuned on other subjects) or +0.020 (self-tuned) above the best competitor, pooled over 28 subject-cells, and the model is better in 6–7 of them. The best competitor is band-limited upsampling with little or no smoothing. The gap shrinks as the grid gets coarser, to near parity at 2.0 → 4.0 mm. So the cross-session failure (S6.1) was not only between-session differences.
2. **Where signals mix across a boundary, the model's unmixing works on real data.** In visual vertices whose coarse map draws

    10% from the other hemisphere, the two hemispheres respond to opposite hemifields. There the model beats the best competitor: pooled −0.008 (13 of 23 subject-cells), and at 2.0 → 4.0 mm −0.031 in 7 of 7 subjects. This reproduces the mechanism of the HCP emulation on real 7T data, but only for a few hundred vertices per subject.

3. **Coarsening loses information that no method recovers.** The source half at native resolution predicts the other half with residual 0.15–0.18. The best method from the coarse data reaches 0.24–0.30.
4. **The noise is spatially coherent.** Coarse runs vary 4.5× (×1.5) and 8.4× (×2) more than independent voxel noise predicts. That is close to the number of voxels averaged, so run-to-run noise hardly averages out over several millimetres. This is the model's largest misspecification on these data; the HCP uncertainty calibration (S5.4) points the same way.
5. **Settings from the data's own split halves** pick the weakest regularization on offer. Within a session they do almost as well as tuning on other subjects (+0.020 vs +0.016). Across sessions (Part B) they help for 7T sources and hurt for 3T sources, and no rule makes the model beat simple methods in general: +0.023 vs self-tuned smoothing, better in 27 of 102 subject-pairs.

Run-to-run residuals of the StudyForrest 7T betas (sub-18; unit variance per voxel; both hemifield maps), radially averaged power spectrum relative to white noise:

| grid (sub-18) | correlation at 1 voxel | correlation at ~6–8 mm | power at λ > 20 mm (× white) | power near 4 mm (× white) |
|---|---|---|---|---|
| 1.4 mm native | 0.25 | 0.12–0.14 | 69 | ~1 |
| 2.0 mm native | 0.29 | 0.15 | 35 | ~1 |
| 1.4 → 2.1 mm emulated | 0.59 | 0.25–0.27 | 47 | ~1 |
| 1.4 → 2.8 mm emulated | 0.62 | 0.28–0.30 | 26 | – |

Spectra fall roughly as $k^{-2}$ at low frequency. The emulated grids' lag-1 correlation includes the emulation window. Above (point 4), the observed / propagated variance of the coarse runs was

4.5× (×1.5) and 8.4× (×2), close to the number of native voxels averaged. Both findings describe the same coherent noise.

**2. Correlated-noise likelihood (within-session emulation, as above)**

Same data, splits, grids, targets and competitors as the within-session emulation above. The model was fitted twice: with white noise (calibrated marginal variance; a small grid around the earlier optimum) and with correlated noise (the same marginal variance times a stationary correlation whose spectrum comes from the source half's own coarse run residuals, radially averaged and floored at 0.1 of white). Settings were chosen by LOSO or by the source-only split-half rule. Visual vertices; residual vs the other half at native resolution:

| native → coarse | best competitor (LOSO) | model, white (LOSO / self) | model, correlated (LOSO / self) | correlated − white, LOSO (better) | correlated − best competitor, LOSO (better) |
|---|---|---|---|---|---|
| 1.4 → 2.1 mm | 0.242 | 0.265 / 0.283 | 0.276 / 0.292 | +0.011 (4/7) | +0.035 (1/7) |
| 1.4 → 2.8 mm | 0.287 | 0.297 / 0.307 | 0.297 / 0.319 | −0.000 (4/7) | +0.011 (2/7) |
| 2.0 → 3.0 mm | 0.248 | 0.270 / 0.269 | 0.260 / 0.265 | −0.010 (6/7) | +0.012 (2/7) |
| 2.0 → 4.0 mm | 0.299 | 0.307 / 0.301 | 0.297 / 0.296 | −0.010 (6/7) | −0.001 (4/7) |

Pooled model − best competitor (LOSO / self): visual, white +0.016 (7/28) / +0.017 (8/28); correlated +0.014 (9/28) / +0.021 (9/28). Covered vertices: white +0.020 / +0.027, correlated +0.018 / +0.027. Source half at native resolution: 0.15–0.18. Tables `w6_tables.md`.

##### S6.3 Leak-gated estimator

| native → coarse | competitor everywhere | model everywhere | gated, far-leak gate | gated, other-hemisphere gate |
|---|---|---|---|---|
| 1.4 → 2.1 mm | 0.2416 | 0.2653 | 0.2416 | 0.2418 |
| 1.4 → 2.8 mm | 0.2865 | 0.3019 | 0.2865 | 0.2865 |
| 2.0 → 3.0 mm | 0.2480 | 0.2599 | 0.2480 | 0.2478 |
| 2.0 → 4.0 mm | 0.2986 | 0.2972 | 0.3004 | 0.2972 |

##### S6.4 Cross-session with settings chosen on the other subjects (pre-declared in the first plan)

Plan SHA-1 e911a51fd3a7c32d3f8cf8fbfea01dcafc946865. Residual of the target map (lower is better); Δ = model − competitor per subject, mean over the pair types in the set. No distortion correction of the 3T sessions (no field maps).

| test | n | mean Δ | median Δ | model better | bootstrap 95% | p (one-sided) | p (Holm) |
|---|---|---|---|---|---|---|---|
| primary: 3T->7T, ratio >= 2.5, visual | 5 | +0.0141 | +0.0070 | 2/5 | [-0.0041, +0.0396] | 0.844 | nan |
| secondary: all 3T->7T, visual | 5 | +0.0291 | +0.0297 | 0/5 | [+0.0172, +0.0397] | 1.000 | 1.000 |
| secondary: all 7T->7T, visual | 7 | +0.0104 | +0.0061 | 2/7 | [-0.0005, +0.0220] | 0.945 | 1.000 |
| secondary: 3T->7T, ratio >= 2.5, covered | 5 | +0.0178 | +0.0048 | 1/5 | [+0.0010, +0.0405] | 0.969 | 1.000 |
| secondary: all 3T->7T, covered | 5 | +0.0359 | +0.0382 | 0/5 | [+0.0280, +0.0430] | 1.000 | 1.000 |

#### Per pair type (visual)

| pair | ratio | subjects | mean competitor | mean model | mean Δ | model better | target split-half | settings chosen (most often) |
|---|---|---|---|---|---|---|---|---|
| 3T-r30->7T-r08 | 3.75 | 5 | 0.5206 | 0.5316 | +0.0110 | 2/5 | 0.5542 | resample_smoothing; x_model_r12_s0.25 |
| 7T-r30->7T-r08 | 3.75 | 7 | 0.4845 | 0.4903 | +0.0059 | 4/7 | 0.5258 | x_resample_smooth6; x_model_r12_s0.5 |
| 3T-r20->7T-r08 | 2.50 | 5 | 0.4970 | 0.5142 | +0.0171 | 2/5 | 0.5506 | resample_smoothing; x_model_r12_s0.25 |
| 7T-r20->7T-r08 | 2.50 | 7 | 0.4454 | 0.4450 | -0.0005 | 4/7 | 0.5200 | x_resample_smooth4; x_model_r6_s0.5 |
| 3T-r30->7T-r14 | 2.14 | 5 | 0.3858 | 0.4032 | +0.0173 | 1/5 | 0.1791 | x_resample_smooth6; x_model_r12_s0.5 |
| 7T-r30->7T-r14 | 2.14 | 7 | 0.3342 | 0.3454 | +0.0112 | 2/7 | 0.1683 | x_resample_smooth2; x_model_r6_s1 |
| 3T-r14->7T-r08 | 1.75 | 5 | 0.5064 | 0.5167 | +0.0103 | 2/5 | 0.5461 | smoothing; x_model_r24_s0.25 |
| 7T-r14->7T-r08 | 1.75 | 7 | 0.4416 | 0.4449 | +0.0033 | 3/7 | 0.5175 | x_resample_smooth4; x_model_r6_s0.5 |
| 3T-r30->7T-r20 | 1.50 | 5 | 0.3479 | 0.3813 | +0.0333 | 1/5 | 0.1349 | x_resample_smooth4; x_model_r12_s1 |
| 7T-r30->7T-r20 | 1.50 | 7 | 0.2873 | 0.3033 | +0.0160 | 3/7 | 0.1510 | x_resample_smooth2; x_model_r6_s1 |
| 3T-r20->7T-r14 | 1.43 | 5 | 0.3697 | 0.3992 | +0.0295 | 1/5 | 0.1806 | x_resample_smooth6; x_model_r12_s0.5 |
| 7T-r20->7T-r14 | 1.43 | 7 | 0.2178 | 0.2442 | +0.0264 | 1/7 | 0.1705 | x_resample_smooth2; x_model_r3_s1 |
| 3T-r14->7T-r14 | 1.00 | 5 | 0.3876 | 0.4161 | +0.0284 | 0/5 | 0.1740 | resample_smoothing; x_model_r12_s0.25 |
| 3T-r20->7T-r20 | 1.00 | 5 | 0.3366 | 0.3915 | +0.0549 | 0/5 | 0.1373 | x_resample_smooth4; x_model_r24_s1 |
| 3T-r30->7T-r30 | 1.00 | 5 | 0.3615 | 0.4109 | +0.0494 | 0/5 | 0.2252 | x_resample_smooth3; x_model_r3_s2 |
| 3T-r14->7T-r20 | 0.70 | 5 | 0.3823 | 0.4148 | +0.0326 | 0/5 | 0.1336 | resample_smoothing; x_model_r12_s0.25 |
| 3T-r20->7T-r30 | 0.67 | 5 | 0.3779 | 0.4216 | +0.0437 | 0/5 | 0.2255 | x_resample_smooth6; x_model_r3_s2 |
| 3T-r14->7T-r30 | 0.47 | 5 | 0.4379 | 0.4598 | +0.0219 | 0/5 | 0.2217 | x_smooth12; x_model_r12_s0.25 |

## S7. ds002702

Odd → even runs and back; the source half is coarsened in k-space by ×1.3125 (1.05 mm, the SE voxel), ×2 (1.6 mm) and ×3 (2.4 mm, the 3T voxel). The target is the other half's six ring maps at 0.8 mm. Settings are chosen leave-one-subject-out. Competitors: band-limited upsampling ± 2 / 4 mm, and the coarse map + 4 / 8 mm. Model: white or correlated noise over a prior grid. The first fits had their optimum at the edge of the initial grid, so both grids were widened before any subject's results were summarised. Noise calibration (observed / propagated coarse-run variance): about 2, 3 and 4 at ×1.3125, ×2 and ×3.

| 0.8 mm → | vertices | best competitor | model | gated | source half at 0.8 mm | model − competitor (better) | chosen (most often) |
|---|---|---|---|---|---|---|---|
| 1.05 mm | visual (58044) | 0.2208 | 0.2534 | 0.2211 | 0.2188 | +0.033 (0/5) | white 3 mm / 1; upsampling |
| 1.6 mm | visual (57255) | 0.2505 | 0.2791 | 0.2505 | 0.2187 | +0.029 (0/5) | white 12 mm / 2; upsampling |
| 2.4 mm | visual (50872) | 0.3250 | 0.3276 | 0.3259 | 0.2236 | +0.003 (2/5) | correlated 12 mm / 2; upsampling |

| 0.8 mm → | vertices | best competitor | model | gated | source half at 0.8 mm | model − competitor (better) | chosen (most often) |
|---|---|---|---|---|---|---|---|
| 2.4 mm | covered (136202) | 0.4792 | 0.4805 | 0.4799 | 0.4622 | +0.001 (2/5) | correlated 6 mm / 1; upsampling + 2 mm |

The model loses at the two finer grids and ties at 2.4 mm. There is no dose-response on these data. At 2.4 mm, model − competitor by far leak at the coarse grid is −0.001, −0.002, +0.010, +0.012 and +0.005 for 0–2, 2–5, 5–10, 10–20 and > 20% (visual cortex clean at 0.8 mm; 12001 down to 303 vertices). That is expected for this stimulus. Rings drive the two banks of the calcarine alike, so mixing them changes the standard map little and there is little to unmix. The far leak measures mixing, not harm. Harm needs mixing of different responses, as with StudyForrest's hemifields and HCP's polar angle. Gating reproduces the competitor (±0.001). Tables `ds002702/summary/kay_tables.md`; figure `ds002702/summary/figW7-4_kay_dose_gate.png`.

**3b. Real second acquisition, pre-specified settings (primary analysis)**

3T 2.4 mm → 7T 0.8 mm (sub-01, sub-03) and 7T SE 1.05 mm → 7T GE 0.8 mm (sub-01, sub-04), same people, separate sessions. The target is the 7T GE session's map from all runs. Settings come from the emulation at the matching factor (`kay_prespec.json`, written before any cross-session score was read). Competitor: resampling to the target grid without smoothing (the cross-session analogue of upsampling). Model: correlated 12 mm / 2 (3T source) and white 3 mm / 1 (SE source). Two stages: uncorrected, and field-map corrected (per session only where the correction improved the registration check; it did everywhere except sub-01 SE). Visual cortex (45–63 thousand vertices):

| pair | stage | competitor (pre-spec.) | model (pre-spec.) | model − competitor | best competitor in hindsight | best model in hindsight | target odd vs even |
|---|---|---|---|---|---|---|---|
| sub-01 3T → 7T | uncorrected | 0.550 | 0.591 | +0.041 | 0.474 (8 mm) | 0.475 | 0.206 |
| sub-01 3T → 7T | field-map corrected | 0.393 | 0.455 | +0.062 | 0.353 (resample + 4 mm) | 0.344 | 0.189 |
| sub-03 3T → 7T | uncorrected | 0.645 | 0.596 | −0.049 | 0.558 (8 mm) | 0.522 | 0.184 |
| sub-03 3T → 7T | field-map corrected | 0.512 | 0.507 | −0.005 | 0.456 (6 mm) | 0.433 | 0.166 |
| sub-01 SE → GE | uncorrected | 0.704 | 0.655 | −0.049 | 0.479 (8 mm) | 0.559 | 0.187 |
| sub-01 SE → GE | field-map corrected | 0.658 | 0.611 | −0.047 | 0.422 (8 mm) | 0.512 | 0.173 |
| sub-04 SE → GE | uncorrected | 0.807 | 0.749 | −0.058 | 0.570 (8 mm) | 0.647 | 0.196 |
| sub-04 SE → GE | field-map corrected | 0.809 | 0.752 | −0.057 | 0.561 (8 mm) | 0.642 | 0.185 |

The pre-specified model beats the pre-specified competitor in 6 of 8 pair × stage cells, but the comparison is not informative: **settings chosen within a session do not transfer across sessions.** Within a session, the best simple method uses no smoothing. Across sessions it needs 6–8 mm, which absorbs registration, distortion and contrast differences. The model similarly needs a stronger prior (white 12 mm / 0.5–1). Both pre-specified methods are 0.04–0.25 worse than the same methods tuned in hindsight. This repeats the lesson of S6.1 with a new dataset. A setting-selection rule has to be validated in the regime where it is used.

###### 3c. Real second acquisition, settings chosen on the other subject (second analysis)

Added after 3b. Each method's setting and the gate threshold are chosen on the other subject of the same pair type and stage (3T: sub-01 ↔ sub-03; SE: sub-01 ↔ sub-04). No target data from the scored subject are used. Visual cortex:

| pair | stage | competitor | model | model – competitor |
|---|---|---|---|---|
| sub-01 3T → 7T | field-map corrected | 0.365 (6 mm) | 0.344 (white 12 mm / 1) | **−0.021** |
| sub-03 3T → 7T | field-map corrected | 0.461 (resample + 4 mm) | 0.433 (white 12 mm / 1) | **−0.028** |
| sub-01 3T → 7T | uncorrected | 0.474 (8 mm) | 0.521 (correlated 24 mm / 0.5) | +0.047 |
| sub-03 3T → 7T | uncorrected | 0.558 (8 mm) | 0.533 (white 12 mm / 1) | −0.025 |
| sub-01 SE → GE | field-map corrected | 0.422 (8 mm) | 0.512 (white 12 mm / 0.5) | +0.090 |
| sub-04 SE → GE | field-map corrected | 0.561 (8 mm) | 0.642 (white 12 mm / 0.5) | +0.081 |
| sub-01 SE → GE | uncorrected | 0.479 (8 mm) | 0.559 | +0.081 |
| sub-04 SE → GE | uncorrected | 0.570 (8 mm) | 0.647 | +0.078 |

- **3T 2.4 mm → 7T 0.8 mm, after field-map correction: the model beats the best simple method in both subjects** (−0.021, −0.028; covered cortex −0.013, −0.016). The gain holds in every far-leak bin (sub-01 −0.009 to −0.025, sub-03 −0.019 to −0.049). For sub-03 it is larger in the leakier bins. This is the first real, separately acquired, three-fold resolution gap on which the model wins. It rests on two subjects and a selection rule added after the primary analysis, so it is suggestive, not confirmatory.
- Without field-map correction the result splits (sub-01 +0.047, sub-03 −0.025). The setting chosen on sub-03 transfers badly to sub-01.
- **SE 1.05 mm → GE 0.8 mm: the model loses clearly** (+0.08 to +0.09, both subjects, both stages); 8 mm smoothing is best. The gap in voxel size is small (×1.3), and the two contrasts differ in fine-scale structure. SE-BOLD is weighted towards microvasculature, GE-BOLD towards pial veins. A method that preserves the source's fine structure is penalised when the target's fine structure is different. The forward model has no term for this.
- The gate never beats the better of its two parts. With the 3T source it mostly picks $\tau = 0$ (model wherever leak > 0); with the SE source it switches the model off.

**Field-map correction matters more than the estimator.** It lowers the best simple method's residual by 0.12 (sub-01 3T), 0.10 (sub-03 3T), 0.06 (sub-01 SE; target only) and 0.01 (sub-04 SE). All predictions remain far from the target's own odd/even agreement (0.17–0.21), so most of the cross-session error is still not resolution. Tables `ds002702/summary/kay_cross_tables.md` (3b) and `kay_cross_cv_tables.md` (3c); figure `figW7-3_kay_cross.png`.

#### S8. Contamination-weighted leak: pre-declared tests

##### S8.1 First pre-declared analysis (raw response estimate)

Plan SHA-1 e911a51fd3a7c32d3f8cf8fbfea01dcafc946865 (`STATS_PLAN.md`). D = gain in the highest CWL bin − gain in the 0–2% bin, per subject (negative = the model gains more where far cortex with different responses mixes in). One-sided Wilcoxon signed-rank (H1: < 0), bootstrap 95% interval of the mean.

##### Primary (CWL-M)

| dataset | n | mean D | median D | D < 0 | bootstrap 95% | p | p (Holm, HCP grids) |
|---|---|---|---|---|---|---|---|
| HCP 2.4 mm | 15 | -0.0234 | -0.0280 | 11/15 | [-0.0384, -0.0077] | 0.008 | 0.008 |
| HCP 3.2 mm | 15 | -0.0366 | -0.0347 | 15/15 | [-0.0450, -0.0286] | 0.000 | 0.000 |
| StudyForrest (4 cells) | 7 | -0.0022 | +0.0023 | 3/7 | [-0.0128, +0.0062] | 0.531 | – |
| ds002702 2.4 mm | 5 | -0.0151 | -0.0184 | 4/5 | [-0.0286, +0.0014] | 0.094 | – |

##### Secondary (Holm over all rows)

| test | n | mean | median | < 0 | bootstrap 95% | p | p (Holm) |
|---|---|---|---|---|---|---|---|
| HCP 2.4 mm: far-leak contrast | 15 | -0.0244 | -0.0255 | 12/15 | [-0.0371, -0.0114] | 0.003 | 0.023 |
| HCP 2.4 mm: CWL-P contrast | 15 | -0.0414 | -0.0377 | 14/15 | [-0.0566, -0.0275] | 0.000 | 0.001 |
| HCP 2.4 mm: CWL-P gated - competitor | 15 | -0.0076 | -0.0067 | 12/15 | [-0.0113, -0.0043] | 0.000 | 0.003 |
| HCP 3.2 mm: far-leak contrast | 15 | -0.0229 | -0.0244 | 13/15 | [-0.0310, -0.0147] | 0.000 | 0.002 |
| HCP 3.2 mm: CWL-P contrast | 15 | -0.0365 | -0.0420 | 15/15 | [-0.0464, -0.0264] | 0.000 | 0.000 |
| HCP 3.2 mm: CWL-P gated - competitor | 15 | -0.0117 | -0.0115 | 14/15 | [-0.0154, -0.0081] | 0.000 | 0.001 |
| StudyForrest (4 cells): far-leak contrast | 7 | -0.0133 | -0.0171 | 5/7 | [-0.0240, -0.0030] | 0.055 | 0.328 |
| StudyForrest (4 cells): CWL-P contrast | 7 | -0.0088 | -0.0026 | 5/7 | [-0.0228, +0.0038] | 0.188 | 0.781 |
| StudyForrest (4 cells): CWL-P gated - competitor | 7 | +0.0017 | -0.0000 | 4/7 | [+0.0005, +0.0033] | 0.766 | 1.000 |
| ds002702 2.4 mm: far-leak contrast | 5 | +0.0078 | +0.0141 | 2/5 | [-0.0156, +0.0282] | 0.781 | 1.000 |
| ds002702 2.4 mm: CWL-P contrast | 5 | -0.0133 | -0.0106 | 4/5 | [-0.0334, +0.0035] | 0.156 | 0.781 |
| ds002702 2.4 mm: CWL-P gated - competitor | 5 | +0.0024 | +0.0000 | 2/5 | [-0.0000, +0.0050] | 0.844 | 1.000 |

##### Descriptive

| quantity | n | mean | median | < 0 | bootstrap 95% | p |
|---|---|---|---|---|---|---|
| ds002702 1.6 mm: CWL-M contrast | 5 | +0.0031 | -0.0068 | 3/5 | [-0.0150, +0.0268] | 0.500 |
| ds002702 1.6 mm: far-leak contrast | 5 | -0.0235 | -0.0266 | 3/5 | [-0.0443, -0.0031] | 0.156 |
| ds002702 1.05 mm: CWL-M contrast | 5 | +0.0006 | -0.0043 | 3/5 | [-0.0184, +0.0200] | 0.500 |
| ds002702 1.05 mm: far-leak contrast | 5 | -0.0382 | -0.0253 | 5/5 | [-0.0750, -0.0097] | 0.031 |
| HCP 2.4 mm: model everywhere - competitor | 15 | -0.0082 | -0.0071 | 13/15 | [-0.0118, -0.0047] | 0.000 |
| HCP 3.2 mm: model everywhere - competitor | 15 | -0.0137 | -0.0133 | 14/15 | [-0.0175, -0.0098] | 0.000 |
| StudyForrest (4 cells): model everywhere - competitor | 7 | +0.0124 | +0.0158 | 2/7 | [-0.0010, +0.0242] | 0.945 |
| ds002702 2.4 mm: model everywhere - competitor | 5 | +0.0026 | +0.0009 | 2/5 | [-0.0035, +0.0088] | 0.781 |
| ds002702 1.6 mm: model everywhere - competitor | 5 | +0.0286 | +0.0246 | 0/5 | [+0.0231, +0.0344] | 1.000 |
| ds002702 1.05 mm: model everywhere - competitor | 5 | +0.0326 | +0.0361 | 0/5 | [+0.0275, +0.0367] | 1.000 |

**Plan B (StudyForrest cross-session, settings chosen on the other subjects; from sf_cross_loso.py)**

| test | n | mean Δ | median Δ | model better | bootstrap 95% | p |
|---|---|---|---|---|---|---|
| primary: 3T->7T, ratio >= 2.5, visual | 5 | +0.0141 | +0.0070 | 2/5 | [-0.0041, +0.0396] | 0.844 |
| secondary: all 3T->7T, visual | 5 | +0.0291 | +0.0297 | 0/5 | [+0.0172, +0.0397] | 1.000 |
| secondary: all 7T->7T, visual | 7 | +0.0104 | +0.0061 | 2/7 | [-0.0005, +0.0220] | 0.945 |
| secondary: 3T->7T, ratio >= 2.5, covered | 5 | +0.0178 | +0.0048 | 1/5 | [+0.0010, +0.0405] | 0.969 |
| secondary: all 3T->7T, covered | 5 | +0.0359 | +0.0382 | 0/5 | [+0.0280, +0.0430] | 1.000 |

**Gain (model − competitor) by bin, mean over subjects (n subjects with the bin)**

| dataset | variable | 0-2% | 2-5% | 5-10% | 10-20% | >20% |
|---|---|---|---|---|---|---|
| HCP 2.4 mm | cwlM | -0.0049 (15) | -0.0126 (15) | -0.0250 (15) | -0.0326 (15) | -0.0287 (14) |
| HCP 2.4 mm | cwlP | -0.0058 (15) | -0.0114 (15) | -0.0219 (15) | -0.0293 (15) | -0.0481 (14) |
| HCP 2.4 mm | far | -0.0086 (15) | -0.0103 (15) | -0.0143 (15) | -0.0193 (15) | -0.0351 (14) |
| HCP 3.2 mm | cwlM | -0.0107 (15) | -0.0119 (15) | -0.0200 (15) | -0.0277 (15) | -0.0473 (15) |
| HCP 3.2 mm | cwlP | -0.0116 (15) | -0.0136 (15) | -0.0180 (15) | -0.0260 (15) | -0.0481 (15) |
| HCP 3.2 mm | far | -0.0165 (15) | -0.0137 (15) | -0.0186 (15) | -0.0184 (15) | -0.0394 (15) |
| StudyForrest (4 cells) | cwlM | +0.0088 (7) | +0.0135 (7) | +0.0150 (7) | +0.0184 (7) | +0.0025 (7) |
| StudyForrest (4 cells) | cwlP | +0.0102 (7) | +0.0120 (7) | +0.0122 (7) | +0.0085 (7) | -0.0001 (7) |
| StudyForrest (4 cells) | far | +0.0105 (7) | +0.0129 (7) | +0.0180 (7) | +0.0021 (7) | -0.0027 (7) |
| ds002702 2.4 mm | cwlM | +0.0096 (5) | +0.0024 (5) | +0.0037 (5) | -0.0003 (5) | -0.0055 (5) |
| ds002702 2.4 mm | cwlP | +0.0083 (5) | -0.0030 (5) | +0.0027 (5) | +0.0028 (5) | -0.0050 (5) |
| ds002702 2.4 mm | far | -0.0004 (5) | -0.0018 (5) | +0.0072 (5) | +0.0100 (5) | +0.0073 (5) |
| ds002702 1.6 mm | cwlM | +0.0278 (5) | +0.0300 (5) | +0.0314 (5) | +0.0225 (5) | +0.0342 (3) |
| ds002702 1.6 mm | cwlP | +0.0277 (5) | +0.0298 (5) | +0.0329 (5) | +0.0388 (5) | -0.0194 (1) |
| ds002702 1.6 mm | far | +0.0278 (5) | +0.0247 (5) | +0.0221 (5) | +0.0042 (5) | – |
| ds002702 1.05 mm | cwlM | +0.0314 (5) | +0.0388 (5) | +0.0320 (5) | – | – |
| ds002702 1.05 mm | cwlP | +0.0318 (5) | +0.0387 (5) | +0.0497 (5) | – | – |
| ds002702 1.05 mm | far | +0.0390 (5) | +0.0227 (5) | -0.0091 (3) | – | – |

**Exploratory addition, from the analysis report (not in the plan): StudyForrest (7 subjects): primary test negative.** D = −0.002 (3/7; p = 0.53). An exploratory check, added after the plan, bins by the other-hemisphere share instead. It does show the effect of the within-session analysis (S6.2): D = −0.019 (6/7; p = 0.023), with gains of −0.017 above 10%. CWL misses it, probably because its response estimate (a 5-run half at native resolution) is noisy. With hemifield stimuli, the contrast that matters is across the midline, a small set of vertices; elsewhere the "response differences" CWL measures are mostly noise. The smoother coarse-data version comes closer (−0.009, 5/7). 4. **Gating by CWL does not beat the better of its two parts.**

- HCP: the chosen threshold is always the lowest. The result is essentially "model everywhere" (−0.008 / −0.012 vs the competitor; model everywhere −0.008 / −0.014).
- StudyForrest and ds002702: the gate equals the competitor (+0.002).

5. **The ds002702 cross-session gain does not generalise to StudyForrest.** Settings were chosen on the other subjects of the same pair type; there is no field-map correction, because the 3T sessions have no field maps.

- Primary (3T → 7T with a voxel ratio ≥ 2.5): model − competitor +0.014 (2/5 better; p = 0.84).
- All 3T → 7T pairs: +0.029 (0/5).
- The deficit shrinks as the resolution gap grows (+0.03 to +0.05 at equal voxel size, −0.001 to +0.017 at ×2.5–3.75), but it never becomes a clear gain.

6. **Data request drafted** (`data_request_draft.md`) for the preprint by Schellekens, Nota, Groen, Piantoni, Winawer & Petridou (bioRxiv 2025.06.03.657640). It is ready for you to review and send.

##### S8.2 Second pre-declared analysis (denoised response estimates)

Plan SHA-1 6375ccdd4bb46c96027c0c7e8907962afaa3b46c (`STATS_PLAN_w9.md`). D = gain in the highest bin − gain in the 0–2% bin per subject (negative = the model gains more where contamination is high). One-sided Wilcoxon signed-rank, bootstrap 95% interval of the mean.

###### Primary (S4: source native map smoothed 4 mm)

| test | n | mean D | median D | D < 0 | bootstrap 95% | p |
|---|---|---|---|---|---|---|
| StudyForrest (4 cells): S4 | 7 | -0.0102 | -0.0068 | 4/7 | [-0.0244, +0.0025] | 0.188 |
| ds002702 2.4 mm: S4 | 5 | -0.0213 | -0.0194 | 4/5 | [-0.0361, -0.0064] | 0.062 |

###### Secondary (Holm over all rows)

| test | n | mean D | median D | D < 0 | bootstrap 95% | p | p (Holm) |
|---|---|---|---|---|---|---|---|
| StudyForrest (4 cells): S2 | 7 | -0.0067 | -0.0010 | 4/7 | [-0.0191, +0.0020] | 0.234 | 0.594 |
| StudyForrest (4 cells): S8 | 7 | -0.0136 | -0.0114 | 5/7 | [-0.0291, +0.0010] | 0.148 | 0.594 |
| StudyForrest (4 cells): Q | 7 | -0.0103 | -0.0120 | 5/7 | [-0.0208, +0.0007] | 0.078 | 0.391 |
| ds002702 2.4 mm: S2 | 5 | -0.0148 | -0.0165 | 4/5 | [-0.0287, +0.0021] | 0.156 | 0.594 |
| ds002702 2.4 mm: S8 | 5 | -0.0239 | -0.0150 | 5/5 | [-0.0454, -0.0067] | 0.031 | 0.188 |
| ds002702 2.4 mm: Q | 5 | -0.0135 | -0.0115 | 4/5 | [-0.0354, +0.0046] | 0.219 | 0.594 |
| HCP 2.4 mm: S4 | 15 | -0.0276 | -0.0301 | 12/15 | [-0.0404, -0.0153] | 0.001 | 0.007 |
| HCP 2.4 mm: Q | 15 | -0.0311 | -0.0237 | 14/15 | [-0.0415, -0.0213] | 0.000 | 0.000 |
| HCP 3.2 mm: S4 | 15 | -0.0330 | -0.0289 | 15/15 | [-0.0410, -0.0253] | 0.000 | 0.000 |
| HCP 3.2 mm: Q | 15 | -0.0278 | -0.0301 | 15/15 | [-0.0365, -0.0193] | 0.000 | 0.000 |

###### Descriptive

| quantity | n | mean | median | < 0 | bootstrap 95% | p |
|---|---|---|---|---|---|---|
| ds002702 1.6 mm: S4 | 5 | +0.0072 | +0.0098 | 1/5 | [-0.0000, +0.0138] | 0.938 |
| ds002702 1.05 mm: S4 | 5 | -0.0026 | -0.0118 | 3/5 | [-0.0216, +0.0195] | 0.500 |
| StudyForrest (4 cells): D(S4) - D(first-analysis CWL-M) | 7 | -0.0081 | -0.0052 | 4/7 | [-0.0192, +0.0022] | 0.188 |

| quantity | n | mean | median | < 0 | bootstrap 95% | p |
|---|---|---|---|---|---|---|
| ds002702 2.4 mm: D(S4) - D(first-analysis CWL-M) | 5 | -0.0062 | -0.0010 | 3/5 | [-0.0162, +0.0030] | 0.312 |
| HCP 2.4 mm: D(S4) - D(first-analysis CWL-M) | 15 | -0.0042 | +0.0012 | 7/15 | [-0.0164, +0.0067] | 0.381 |
| HCP 3.2 mm: D(S4) - D(first-analysis CWL-M) | 15 | +0.0037 | +0.0056 | 4/15 | [-0.0010, +0.0080] | 0.958 |
| ds002702 1.6 mm: D(S4) - D(first-analysis CWL-M) | 5 | +0.0041 | +0.0082 | 1/5 | [-0.0149, +0.0199] | 0.781 |
| ds002702 1.05 mm: D(S4) - D(first-analysis CWL-M) | 5 | -0.0033 | -0.0021 | 3/5 | [-0.0241, +0.0179] | 0.500 |

#### Gain (model − competitor) by bin, mean over subjects (n subjects)

| dataset | estimate | 0-2% | 2-5% | 5-10% | 10-20% | >20% |
|---|---|---|---|---|---|---|
| StudyForrest (4 cells) | cwlS4 | +0.0104 (7) | +0.0135 (7) | +0.0162 (7) | +0.0118 (7) | -0.0025 (7) |
| StudyForrest (4 cells) | cwlQ | +0.0126 (7) | +0.0146 (7) | +0.0125 (7) | +0.0096 (7) | -0.0003 (7) |
| StudyForrest (4 cells) | cwlS8 | +0.0117 (7) | +0.0140 (7) | +0.0125 (7) | +0.0124 (7) | -0.0051 (7) |
| StudyForrest (4 cells) | cwlM | +0.0088 (7) | +0.0135 (7) | +0.0150 (7) | +0.0184 (7) | +0.0025 (7) |
| StudyForrest (4 cells) | far | +0.0105 (7) | +0.0129 (7) | +0.0180 (7) | +0.0021 (7) | -0.0027 (7) |
| ds002702 2.4 mm | cwlS4 | +0.0104 (5) | -0.0007 (5) | +0.0027 (5) | +0.0041 (5) | -0.0109 (5) |
| ds002702 2.4 mm | cwlQ | +0.0105 (5) | -0.0020 (5) | +0.0018 (5) | +0.0046 (5) | -0.0030 (5) |
| ds002702 2.4 mm | cwlS8 | +0.0093 (5) | -0.0016 (5) | +0.0026 (5) | +0.0053 (5) | -0.0146 (5) |
| ds002702 2.4 mm | cwlM | +0.0096 (5) | +0.0024 (5) | +0.0037 (5) | -0.0003 (5) | -0.0055 (5) |
| ds002702 2.4 mm | far | -0.0004 (5) | -0.0018 (5) | +0.0072 (5) | +0.0100 (5) | +0.0073 (5) |
| HCP 2.4 mm | cwlS4 | -0.0056 (15) | -0.0136 (15) | -0.0223 (15) | -0.0318 (15) | -0.0346 (13) |
| HCP 2.4 mm | cwlQ | -0.0056 (15) | -0.0131 (15) | -0.0194 (15) | -0.0337 (15) | -0.0367 (15) |
| HCP 2.4 mm | cwlS8 | -0.0059 (15) | -0.0140 (15) | -0.0215 (15) | -0.0312 (15) | -0.0374 (13) |
| HCP 2.4 mm | cwlM | -0.0049 (15) | -0.0126 (15) | -0.0250 (15) | -0.0326 (15) | -0.0287 (14) |
| HCP 2.4 mm | far | -0.0086 (15) | -0.0103 (15) | -0.0143 (15) | -0.0193 (15) | -0.0351 (14) |
| HCP 3.2 mm | cwlS4 | -0.0108 (15) | -0.0136 (15) | -0.0201 (15) | -0.0275 (15) | -0.0438 (15) |
| HCP 3.2 mm | cwlQ | -0.0117 (15) | -0.0129 (15) | -0.0203 (15) | -0.0295 (15) | -0.0395 (15) |
| HCP 3.2 mm | cwlS8 | -0.0114 (15) | -0.0139 (15) | -0.0200 (15) | -0.0280 (15) | -0.0421 (15) |
| HCP 3.2 mm | cwlM | -0.0107 (15) | -0.0119 (15) | -0.0200 (15) | -0.0277 (15) | -0.0473 (15) |
| HCP 3.2 mm | far | -0.0165 (15) | -0.0137 (15) | -0.0186 (15) | -0.0184 (15) | -0.0394 (15) |
| ds002702 1.6 mm | cwlS4 | +0.0284 (5) | +0.0303 (5) | +0.0353 (5) | +0.0364 (5) | +0.0260 (2) |
| ds002702 1.6 mm | cwlQ | +0.0287 (5) | +0.0276 (5) | +0.0376 (5) | +0.0475 (5) | +0.0307 (1) |
| ds002702 1.6 mm | cwlS8 | +0.0282 (5) | +0.0320 (5) | +0.0367 (5) | +0.0283 (5) | +0.0213 (1) |
| ds002702 1.6 mm | cwlM | +0.0278 (5) | +0.0300 (5) | +0.0314 (5) | +0.0225 (5) | +0.0342 (3) |
| ds002702 1.6 mm | far | +0.0278 (5) | +0.0247 (5) | +0.0221 (5) | +0.0042 (5) | – |
| ds002702 1.05 mm | cwlS4 | +0.0324 (5) | +0.0443 (5) | +0.0297 (5) | – | – |
| ds002702 1.05 mm | cwlQ | +0.0323 (5) | +0.0493 (5) | +0.0226 (5) | – | – |
| ds002702 1.05 mm | cwlS8 | +0.0329 (5) | +0.0414 (5) | +0.0249 (5) | – | – |
| ds002702 1.05 mm | cwlM | +0.0314 (5) | +0.0388 (5) | +0.0320 (5) | – | – |
| ds002702 1.05 mm | far | +0.0390 (5) | +0.0227 (5) | -0.0091 (3) | – | – |

#### S9. Quality control, including a processing bug and its check

**Registration (ds002702).** For each session, the check compares EPI intensity 1 mm outside and 1 mm inside the white surface along the normal: grey matter is brighter than white matter in the EPI, consistently across vertices when the registration is right. The same is measured with the registration shifted by 2 mm, and with another subject's surfaces. Registered sessions reach 64–

89% of vertices with grey > white; shifted copies 56–67%. Field-map correction was used per session only where it improved this check: everywhere except sub-01 SE (79% → 70%).

**A processing bug and its check.** The first per-run GLMs on the lab server ran in parallel threads. In that container they silently corrupted whole runs: 5 runs in three 7T sessions, with betas up to $10^7$ % and halves that did not correlate. The GLM now runs sequentially. Every session passes a per-run check, counting voxels with |beta| > 50% and correlating each run's betas with the mean of the other runs (0.23–0.51 at 7T), and the affected sessions were reprocessed. The StudyForrest per-run betas of the within-session and later analyses were produced by a threaded GLM on another system. Their odd/even means reproduced the sequential pipeline of the first cross-session analysis (S6.1) to ≤ 4e-6 % in all 14 sessions, so they are unaffected. A pre-declared run rule (r < 0.1 with the other runs) dropped one 3T run (sub-01, r = 0.05).

**Rebuilt operators.** Every contamination computation of the two pre-declared analyses rebuilt the coarse-grid operator and recomputed the far leak, as a check against the saved values:

- ds002702 and StudyForrest: ≤ 2.5e-5;
- HCP 2.4 mm: r = 1.00;
- HCP 3.2 mm: r = 0.70–1.00. The operator of the HCP emulation has columns only for cortex, so leakage into the medial wall is left out.

#### S10. Resolvability theory

##### S10.1 Summary (moved from the main text)

For planar cortex the tangential bandwidth that a voxel grid can carry depends on the sheet's orientation to the grid. Computed this way, the resolvable tangential wavelength is about 3.1–3.5 mm at 3T and 2.5–2.8 mm at 7T map-level CNR (below). The median distance from a point of cortex to the opposite bank of its sulcus is 5.5 mm, and 21% of cortex has an opposite bank within 4 mm. In our data the achieved resolution of the estimator did *not* follow this sampling limit: the averaging- kernel spread was uncorrelated with the predicted resolvable wavelength (Spearman $\rho \approx 0$). The PSF and the prior, not the lattice, set the achieved resolution. We therefore keep the resolvability theory as background.

##### S10.2 Derivation and lookups

Subjects: 20 (the development set), both hemispheres; surfaces warped MNI→ACPC with each subject's HCP warps.

###### Warp sanity check

- median |pial−white| / FreeSurfer thickness: ACPC 1.142 ± 0.012 vs MNI 1.333 ± 0.046
- correlation with thickness: ACPC 0.874 ± 0.011 vs MNI 0.737 ± 0.027
- midthickness area per hemisphere (ACPC): 916 ± 70 $cm^2$

##### Resolvable isotropic tangential wavelength (area-weighted medians, mean ± SD over hemispheres)

| | idealised formula | acquisition-limited, single frame (CNR 1) | acquisition-limited, map level (CNR 10) | with BOLD blur, map level (CNR 10) |
|---|---|---|---|---|
| 3T | 3.36 ± 0.01 mm | 3.45 ± 0.01 mm | 3.12 ± 0.01 mm | 3.42 ± 0.01 mm |
| 7T | 2.69 ± 0.01 mm | 2.76 ± 0.01 mm | 2.52 ± 0.01 mm | 2.57 ± 0.01 mm |
| fsaverage5 (idealised) | 3T 3.39 mm, 7T 2.71 mm | | | |

##### Degrees of freedom for signal per hemisphere (lookup-based)

| | acquisition-limited, single frame (CNR 1) | acquisition-limited, map level (CNR 10) | with BOLD blur, map level (CNR 10) |
|---|---|---|---|
| 3T | 37,866 ± 2,902 | 62,570 ± 4,775 | 32,428 ± 2,482 |
| 7T | 59,930 ± 4,582 | 97,590 ± 7,424 | 68,249 ± 5,211 |

##### Sulcal-bank proximity and leakage

- median distance to nearest opposite bank: 5.50 ± 0.17 mm
- cortex with an opposite bank within 3 / 4 / 5 / 6 mm: 4.2% / 20.6% / 40.9% / 57.5%
- predicted mean cross-bank leakage of ribbon averaging: 3T 0.81%, 7T 0.86%
- cortex where bank-specific patterns are NOT recoverable even for the optimal estimator (transfer < 0.5): | pattern wavelength | 3T CNR 1 | 3T CNR 10 | 7T CNR 1 | 7T CNR 10 | |---|---|---|---|---| | 7.9 mm | 0.0% | 0.0% | 0.0% | 0.0% | | 5.2 mm | 0.0% | 0.0% | 0.0% | 0.0% | | 3.9 mm | 100.0% | 0.0% | 0.0% | 0.0% |

Standard mapping modelled as box-voxel overlap averaging (as in HCP ribbon mapping), averaged over 6 random lattice positions. Assumptions: acquisition axes = ACPC axes (AutoAlign); planar-patch lookups (curvature ignored); BOLD FWHM 3.0 mm (3T) / 1.5 mm (7T); T2* 45 ms (3T) / 28 ms (7T); partial Fourier ignored; opposite-bank test uses a 15 mm geodesic threshold judged on the registration sphere.

##### Notes added after review

##### S11. Analysis plans (verbatim; hashes recorded by the author before the analyses)

The plans are reproduced as written, except that HCP subject IDs are replaced by study codes or withheld, as the HCP Restricted Data Use Terms require. Their SHA-1 sums, recorded when each plan was finalised, are of the original files and are given with each plan together with the SHA-256 sum of the released file; this is pre-specification attested by the author, not a registration with a third party. Key to the internal vocabulary of the plans: "week N" names the project's analysis stages (weeks 1–2: resolvability lookups, operator and leakage maps; 3: HCP emulation fits; 4: misregistration, depth, uncertainty and StudyForrest across sessions; 5–6: StudyForrest within session and its noise; 7: ds002702 and the gated estimator; 8–9: the two pre-declared contamination analyses; 11: the summary table; 13: the confirmatory cohort; the repository names the code folder of each stage, src/weekN, descriptively, as listed in its README); "the user" denotes the principal investigator, who approved each step; "pilot" is the development set; batch and file names refer to the release.

##### S11.1 First pre-declared analysis (`STATS_PLAN.md`, SHA-1 e911a51fd3a7c32d3f8cf8fbfea01d-cafc946865; SHA-256 ca6db91d9a4426b0688704273286543ac8f7aa5c43107db7d4288df63c93d8b4)

Written 2026-09-30, before any week-8 quantity was computed: no contrast-weighted leak had been computed on any dataset, and the StudyForrest cross-session re-analysis below had not been run. The weekly reports have already shown the far-leak dose-response of HCP and StudyForrest (weeks 6–7) and the ds002702 results (week 7) descriptively. Tests on those earlier numbers are therefore confirmatory in form only, and are marked as such. The SHA-1 of this file is written into every week-8 output.

##### A. Contrast-weighted leak (CWL): does mixing of *different* responses predict the model's gain?

**Definitions.**

- Far set F(v): unknowns u in the other hemisphere, or more than 15 mm away on the registration sphere scaled to the cortical area (week 2).
- Readout row a_v = (W_c K_c)_v: standard ribbon mapping of the coarse-grid operator. tot_v = Σ_u a_vu.
- Response estimate x_u,j per vertex and map j (real maps; complex maps split into real and imaginary parts):
  - **M (mechanism):** the source's own native-resolution standard map. This is the source half (StudyForrest, ds002702) or the source run (HCP). It never uses target data.
  - **P (practical):** the source's coarse data, upsampled in k-space and mapped at the native grid. This needs no fine-resolution data.
- Contamination c_v,j = Σ_{u∈F(v)} a_vu (x_u,j − x_v,j) / tot_v: the change of the standard map at v caused by far cortex responding differently from v.
- **CWL_v = sqrt(Σ_j c_v,j²) / sqrt(mean over the scored set of Σ_j x_v,j²)**, i.e. contamination as a fraction of the typical response. The scored set is the same vertex set as the dose-response: visual (StudyForrest, ds002702) or coherent (HCP).

**Bins** (as the far-leak bins): 0–2%, 2–5%, 5–10%, 10–20%, > 20% of the typical response. Vertices must be clean at the native grid (far leak < 2%). A bin is used for a subject if it has ≥ 30 vertices in each direction.

**Gain** Δ = residual(model) − residual(best competitor) in a bin (negative = model better). Settings are those of the original analysis:

- HCP: the week-3 fixed choices (model 12 mm / 0.125 at 2.4 mm, 24 mm / 0.25 at 3.2 mm; competitor upsampling + 8 mm).
- StudyForrest: week-7 gate runs, leave-one-subject-out.
- ds002702: week-7 emulation, leave-one-subject-out.

**Primary contrast per dataset:** D = Δ(highest CWL-M bin available for the subject) − Δ(0–2% bin), one value per subject. StudyForrest averages its four session × factor cells per subject; ds002702 uses the ×3 (2.4 mm) cells. HCP has two grids.

- H1 (predicted): D < 0 in each dataset.
- Test: one-sided Wilcoxon signed-rank over subjects (HCP n = 15, StudyForrest n = 7, ds002702 n = 5; with n = 5 the smallest attainable p is 0.031), plus a bootstrap 95% interval of the mean of D (10,000 resamples of subjects, percentile).
- Correction: none across datasets (independent). Holm across HCP's two grids.
- **The key prediction is ds002702:** there the far-leak contrast was flat or positive (week 7). H1 predicts D < 0 with CWL-M there too.

**Secondary (Holm-corrected together):**

- (a) The same contrast with far-leak bins, for comparison: confirmatory in form only.
- (b) CWL-P instead of CWL-M.
- (c) A CWL-P-gated estimator: model where CWL-P > τ, competitor elsewhere, τ ∈ {0, 1, 2, 5, 10, 20%, never} chosen leave-one-subject-out, model scaled onto the competitor without target data. Test: gated < competitor, one-sided signed-rank.

##### B. StudyForrest cross-session selection at scale (the ds002702 week-7 3c rule)

Week-4 StudyForrest cross-session predictions, all settings as logged: competitor = source map ± smoothing (2–16 mm) or resampled to the target grid + smoothing (2–16 mm); model = the week-4 prior grid (3–24 mm × 0.125–2). No field maps exist for the 3T sessions, so there is no distortion correction (the 7T data are corrected by the providers).

- **Primary:** 3T → 7T pairs with a voxel ratio ≥ 2.5 (3T 2.0 mm → 7T 0.8 mm and 3T 3.0 mm → 7T 0.8 mm; 5 subjects). Each method's setting is chosen on the other subjects of the same pair type (leave-one-subject-out), visual vertices. Δ per subject = mean over the two pair types.
  - H2: Δ < 0 (the ds002702 3T → 7T gain generalises). One-sided signed-rank (n = 5) and bootstrap interval.
- **Secondary:** all 3T → 7T pairs; all 7T → 7T pairs; covered vertices.

##### C. Reporting

Every test is reported with n, the mean and median of the per-subject values, the bootstrap interval and the p-value. A non-significant primary result is reported as such, with no post-hoc subgroup rescue. Analyses added after this plan are marked "exploratory".

##### S11.2 Second pre-declared analysis (`STATS_PLAN_w9.md`, SHA-1 6375ccdd4b-b46c96027c0c7e8907962afaa3b46c; SHA-256 62c29d0a929e43f7b6a2ca8904c2305642176909c5aedf97db0c17ac8a02f418)

Written 2026-10-01, before any week-9 quantity was computed. The week-8 results it refines are known: CWL-M was negative in HCP, in the predicted direction but not significant in ds002702, and null in StudyForrest. The SHA-1 of this file is written into every week-9 output.

###### Question

Week 8's CWL used the source's own native-resolution map as the response estimate. In StudyForrest it failed, and the suspected reason is noise in that estimate: response differences between far vertices are then partly noise. Does a denoised estimate recover the dose-response?

###### Definitions (as `results/week8/STATS_PLAN.md`, part A, except the response estimate)

- **S4 (primary):** the source's native-resolution standard map, smoothed on the surface with a 4 mm FWHM. Heat-kernel smoothing on each hemisphere's submesh of the unknowns is the same operator as the competitors' smoothing. Hemispheres are never mixed.
- **S2, S8 (secondary):** the same with 2 mm and 8 mm FWHM.
- **Q (secondary, practical):** the model's estimate from the coarse data, rendered at the native grid. It uses the setting of the week-8 analysis for that subject: ds002702 and StudyForrest leave-one-subject-out, HCP the week-3 choice. It needs no fine data.
- **Unchanged:** the far set, the readout rows, the normalisation by the typical response of the scored set, the bins (0–2, 2–5, 5–10, 10–20, > 20%), the clean-at-native criterion, the ≥ 30 vertices rule, the gain Δ and the settings, all as in week 8.

###### Tests

- **Primary:** D = Δ(highest bin with ≥ 30 vertices) − Δ(0–2% bin) per subject, using S4. Two datasets:
  - StudyForrest: 7 subjects; mean of the four session × factor cells per subject.
  - ds002702: 5 subjects; the ×3 (2.4 mm) cells.
  - H1: D < 0 in each. One-sided Wilcoxon signed-rank and a bootstrap 95% interval of the mean (10,000 resamples). No correction across the two datasets (independent), as in week 8.
- **Secondary (Holm over all):**

- S2, S8 and Q in StudyForrest and ds002702 ×3;
- S4 and Q in HCP 2.4 / 3.2 mm (consistency check: week-8 CWL-M was already negative there);
- ds002702 ×2 and ×1.3125 with S4: descriptive only.

- **Paired comparison with week 8 (descriptive):** D(S4) − D(CWL-M) per subject.

A null primary result is reported as such. The exploratory other-hemisphere result of week 8 is not part of this plan.

##### S11.3 Summary table across datasets (`TABLE_SPEC.md`, SHA-1 af8b2d8be80d0eb0624-fafe332439cabf65a5543; SHA-256 cfe3e328ec4794b16083241143c7b82fd4b1f86067f5e0e386fabb8d801ed2f9)

**Purpose.** Restate every main model-versus-competitor comparison of the study with the same statistics, so the datasets and conditions can be compared at a glance. The table is descriptive. It summarises analyses that were already run; only the high-mixing sets of the ds002702 within-session rows (A4–A6) are computed for the first time.

###### Quantity

Per subject, d = residual(model) − residual(best competitor); d < 0 when the model is better. The best competitor is the better of the competitor families for that subject, under the same condition as the model:

- HCP: standard + surface smoothing, or upsampling + 1.6 mm mapping + smoothing;
- StudyForrest within session: smoothing, or upsampling (± smoothing);
- ds002702 within session: the competitor chosen leave-one-subject-out among upsampling ± 2 / 4 mm and coarse map + 4 / 8 mm;
- cross-session rows: the competitor of that analysis.

Where a subject contributes several cells (StudyForrest sessions × coarsening factors) or directions, d is first averaged within the subject with equal weights. The subject is the unit of analysis.

###### Statistics (every row, as in the pre-declared tests)

n subjects; mean and median of d; subjects with d < 0; bootstrap 95% interval of the mean (10,000 resamples, a fresh generator with seed 0 for every row); one-sided Wilcoxon signed-rank p (alternative: d < 0; scipy, exact for small n). No correction for multiplicity. With n = 5 the smallest possible p is 0.031; with n = 2 it is 0.25.

###### Vertex sets (two per row)

- **all:** coherent vertices (HCP) or visual vertices (StudyForrest, ds002702);

- **high mixing:** far leak > 10% at the coarse grid and < 2% at the native grid (HCP, ds002702); > 10% of the coarse map from the other hemisphere and < 2% at the native grid (StudyForrest within session); source far leak > 10% and < 2% at the target grid (ds002702 cross-session). Not available for StudyForrest cross-session.

Settings follow each analysis' own rule; no row uses target data to choose settings.

**Rows**

**A. Emulation within one acquisition**

- A1, A2. HCP 7T, emulated 2.4 / 3.2 mm, 15 test subjects; settings fixed on 5 tuning subjects. Source `results/week3/week3_test_scores.json`: model − fmin(smoothing, upsample_smoothing); sets coherent / clean16_leak10.
- A3. StudyForrest 7T within session, 1.4 / 2.0 mm sessions coarsened ×1.5 / ×2, 7 subjects; settings leave-one-subject-out; correlated-noise model (as in the manuscript's Section 4.5). Source `results/week6/w6_results.json`: corr_loso − min(smoothing_loso, upsample_loso) per cell; sets visual / visual_xhemi.
- A4–A6. ds002702 7T within session, 0.8 mm coarsened to 1.05 / 1.6 / 2.4 mm, 5 subjects; settings leave-one-subject-out. All: `results/week7/ds002702/summary/kay_results.json` (model − comp, visual). High mixing: computed from `results/week7/ds002702/emul/sub-*/*.kay_emul.npz` with the procedure of `src/week7/summarize_kay.py` (same scores, same model and competitor candidates, leave-one-subject-out choice on the set itself) for visual vertices with far_n < 0.02 and far_c ≥ 0.10. A grid enters only if every subject has ≥ 30 such vertices in both directions; otherwise "too few vertices".

**B. Robustness of the HCP result (15 test subjects, settings fixed)**

- B1, B2. 1 mm rigid misregistration of the coarse data, applied to every method; 2.4 / 3.2 mm. Source `results/week4/hcp_eval/week4_hcp_eval.json`: model_shift1 − fmin(smoothing_shift1, upsample_smoothing_shift1).
- B3–B6. Model PSF assumed 15% narrower / wider than the operator's; 2.4 / 3.2 mm. Source `results/week3/week3_test_scores.json`: model_psf0.85 / model_psf1.15 − fmin(smoothing, upsample_smoothing).

**C. Second acquisitions; settings chosen on the other subjects of the pair type**

- C1. StudyForrest 3T → 7T, voxel ratio ≥ 2.5, 5 subjects (the pre-declared primary of the first analysis).
- C2. StudyForrest 7T → 7T, all pairs, 7 subjects. Source `results/week8/sf_cross/sf_cross_loso.json`, per-subject values of those tests.
- C3. ds002702 3T 2.4 mm → 7T GE 0.8 mm, field-map corrected, 2 subjects.
- C4. ds002702 7T SE 1.05 mm → 7T GE 0.8 mm, field-map corrected, 2 subjects. Source `results/week7/ds002702/summary/kay_cross_cv_results.json`: model − competitor (visual);

high mixing from the record's "10-101%" bin.

##### Checks before the table is used

- A1, A2 (all) reproduce the held-out results: 2.4 mm 13/15 and 3.2 mm 14/15 subjects better.
- A4–A6 (all) equal the week-7 table (+0.033, +0.029, +0.003); the visual set recomputed from the npz files equals `kay_results.json`, which validates the high-mixing computation.
- C1 equals the pre-declared result: +0.0141, 2/5, p = 0.844.

##### Output

`results/week11/robustness_table.md` and `.json`; the table enters the manuscript (Section 4.8) and is copied there by script.

**Deviation, recorded after computing.** The check of rows A1 and A2 expected the counts of the manuscript's Section 4.3 (13 and 14 of 15 subjects), which compare with the competitor that is best on average. With the specified per-subject best competitor the same data give 12 and 14 of 15, as in Fig. 4A (right). The check was corrected; the definition was kept.

#### S11.11 Three further analyses of the NYU data (`PLAN_NYU_GATE.md`, SHA-1 6483f196225d-dd0af38dbb5942963e1ddaa3b325, recorded 2026-10-07 after the results of S11.10 and before any of these analyses; SHA-256 43f84353b6bf72077a7f0346cac6ca16e8a8bb76503390d936e5f2072ea666a5)

User's decision (2026-10-07): "do items 2 and 3" — trim the manuscript and run three analyses on the existing NYU data. The NYU results of PLAN_NYU.md are known (the overall gain replicated at 4.0 mm, not at 3.0 mm; contamination predicted the gain at both grids, plain far leak did not); these analyses are added afterwards and are labelled as such. Settings, competitor, subjects, split and scoring are those of PLAN_NYU.md (model 12 mm / 2 at 3.0 mm and 6 mm / 1 at 4.0 mm; competitor upsampling + 2 mm smoothing; 8 tuning, 35 included test subjects).

##### A. A practical rule: correct only where coarse-data contamination is high (pre-specified tests)

- **Measure.** CWL-P, the contamination-weighted leak of standard mapping with the response estimate taken from the coarse data alone (the source half's coarse map, k-space upsampled and mapped at 2.0 mm), exactly as in the paper's CWL-P.
- **Gated estimator.** Per vertex, the model's estimate where CWL-P > $\tau$ and the competitor's elsewhere, with $\tau \in \{0, 0.02, 0.05, 0.10, 0.20, \infty\}$ ($\tau = 0$: the model everywhere; $\tau = \infty$: the competitor everywhere).
- **Choice.** Per grid, $\tau$ with the lowest mean residual over responsive vertices of the 8 tuning subjects.

- **Tests (35 test subjects, Δ over responsive vertices, one-sided Wilcoxon, Holm across the four):** G1: gated − competitor < 0 at 3.0 mm and at 4.0 mm; G2: gated − model < 0 at 3.0 mm and at 4.0 mm. If G1 and G2 hold at a grid, the rule beats both pure strategies there.
- **Descriptive:** D_CWL-P (the dose contrast with CWL-P bins, as D_CWL-M), the share of responsive vertices sent to the model, and the gated estimator's polar-angle error.

##### B. Registration and the pericalcarine shortfall (descriptive)

Per test subject: the boundary-based registration cost, the 95th percentile of the absolute field (Hz) over the brain and the median absolute field at the responsive pericalcarine vertices, against Δ over responsive vertices and Δ in the pericalcarine (model − competitor). Spearman correlations (two-sided, n = 35) and a split at the median registration cost. No test is confirmatory.

##### C. Leakage maps of the 3T 2.0 mm acquisition as a resource (descriptive)

Per included subject, the far-leak maps of standard mapping at 2.0 mm and at the 3.0 / 4.0 mm emulations, carried from the subject's FreeSurfer registration sphere to fsaverage (164k) by nearest-neighbour interpolation on the sphere (three nearest vertices, inverse-distance weights); group mean and share of subjects above 5% per vertex; region table (FreeSurfer aparc, area-weighted mean far leak and share above 10%, mean over subjects), as the HCP table of Section 4.2. Released as GIFTI.

##### Compute

All on hebb inside `/analysis/halilaziz/halil/GEO_CONST`: `nyu_emul.py --gate` reruns every included subject at the chosen settings (reproducing the published scores as a check) and adds CWL-P, the gated scores, the field statistics and the maps; `nyu_groupmaps.py` builds C. Results copied to `results/week17/`.

Hash in `PLAN_NYU_GATE.sha1`, recorded before the first run.

#### S11.12 Contamination from an anatomical atlas (`PLAN_ATLAS.md`, SHA-1 6336f6c5785b0f-b27a66bbdfeba3376b4c476415, recorded 2026-10-13 after the results of S11.10 and S11.11 and before any atlas quantity was computed; SHA-256 d4b221e469ef280851193cdca847513651b182de866d5cc2056a23490d40d947)

User's request (2026-10-13): "run the atlas-based contamination test". In the NYU Retinotopy data, contamination computed with the source half's native-resolution map as the response estimate (CWL-M) predicted the model's gain at both coarsenings (N3, PLAN_NYU.md), while contamination computed from the coarse data alone (CWL-P) did not (PLAN_NYU_GATE.md, descriptive). The question here: does contamination computed from an anatomical retinotopy atlas, which uses no functional data of the subject, predict where the model gains at 3T? If it does, a study could map where correction should pay before acquiring data.

**Measure**

- **Atlas.** The anatomical retinotopy template of Benson et al. (2014), version 4.0 as distributed with neuropythy 0.13.0 (PyPI source archive; files `?h.benson14_{angle,eccen,varea}.v4_0.mgz` on fsaverage, 163,842 vertices per hemisphere; polar angle 0–180° from the upper to the lower vertical meridian in both hemispheres; visual areas 1–12). Only the template files are used; no neuropythy code is run.
- **Transfer to the subject.** For every vertex of the subject's FreeSurfer mesh, the three nearest fsaverage vertices on the registration spheres (`?h.sphere.reg`, unit-normalised), inverse-distance weights: polar angle interpolated on the unit circle, eccentricity linearly, the area label from the nearest vertex.
- **Response estimate.** $x = \exp(i\varphi)$ where the vertex has an area label (1–12) and eccentricity ≤ 12.4° (the NYU stimulus radius, Himmelberg et al., 2021), $x = 0$ elsewhere; $\varphi$ is the polar angle in one visual-field convention for both hemispheres (left hemisphere $\varphi = 90° - \text{angle}$, right hemisphere $\varphi = 90° + \text{angle}$). CWL depends only on differences of x, so the choice of phase origin and direction does not matter.
- **CWL-A.** The contamination-weighted leak of standard mapping at the coarse grid with this x, computed exactly as CWL-M and CWL-P (same far criterion, normalisation by the RMS of x over the responsive vertices, bins 0–2, 2–5, 5–10, 10–20, > 20%, responsive vertices with < 2% far leak at 2.0 mm, ≥ 30 vertices per bin). D_CWL-A = gain in the highest bin with ≥ 30 vertices − gain in the 0–2% bin, gain = residual(model) − residual(competitor).

**Data, settings, subjects**

As PLAN_NYU.md and PLAN_NYU_GATE.md: the 35 included test subjects; model 12 mm / 2 at 3.0 mm and 6 mm / 1 at 4.0 mm; competitor upsampling + 2 mm smoothing; responsive vertices $R^2 \geq 0.2$. `nyu_emul.py` reruns every test subject at these settings with the atlas response added to the contamination pass; the published scores, D_CWL-M and D_CWL-P must be reproduced exactly (check).

**Tests (35 test subjects)**

- **A1:** D_CWL-A < 0 at 3.0 mm. **A2:** D_CWL-A < 0 at 4.0 mm.
- One-sided Wilcoxon signed-rank test, bootstrap 95% interval of the mean (10,000 resamples, seed 0), Holm across the two.
- Interpretation fixed now: if A1 and A2 hold, an anatomy-only map predicts where correction pays at 3T; if both fail, the 3T dose-response needs a response map finer than the coarse data; one of two is reported as such, grid by grid.

**Descriptive**

- D_CWL-A − D_CWL-M and D_CWL-A − D_CWL-P per subject (two-sided Wilcoxon).
- Gain by CWL-A bin (mean over subjects with the bin).
- Per subject, Spearman correlation over the responsive vertices of CWL-A with CWL-M, and of CWL-P with CWL-M (median over subjects): how close each coarse-free or coarse-only measure comes to the fine-data measure.
- Variant: the atlas restricted to V1–V3 (areas 1–3), D_CWL-A3.

**Not part of this plan**

HCP: the per-subject HCP surfaces on TRUBA were cleared after the cohort run; an HCP test would follow under its own plan, and only if A1 and A2 hold. No gated estimator is run.

**Compute**

hebb, inside `/analysis/halilaziz/halil/GEO_CONST`: template files extracted to `data/atlases/benson14_v4_0/surf/`; per-subject atlas maps by `src/week18/atlas_map.py` (fMRIPrep container, FreeSurfer's fsaverage) into `results/nyu/atlas/`; scores by `nyu_emul.py --atlas` into `results/nyu/emul_atlas/`; summary `src/week18/atlas_summary.py` locally; results copied to `results/week18/`.

Hash in `PLAN_ATLAS.sha1`, recorded before the first run.

##### S11.4 Confirmatory cohort (`COHORT_PLAN.md`, SHA-1 21822393451d14655ced1bac656c7e7a7141dbc0, recorded 2026-10-02 before the first fit of any cohort subject; SHA-256 9a8299f121292d036b939531011e9e039ad4a58fa860dfcaf8ddf7387e50deee)

**What is fixed before the data are seen**

- **Subjects.** All HCP 7T retinotopy subjects not in the pilot (164), minus 5 relatives of the 5 tuning subjects and 2 whose S3 packages are incomplete ([HCP ID withheld]: no concatenated retinotopy run; [HCP ID withheld]: no MSMAll spheres): **158 subjects in 86 families**. The 15 pilot test subjects were used to develop the two new analyses below and are reported separately.
- **Settings.** Everything is frozen at the week-3 choices: model prior 2.4 mm (12 mm, 0.125), 3.2 mm (24 mm, 0.25); competitors as chosen on the 5 tuning subjects (evaluate.py re-derives them from the stored tuning estimates); the contamination measures as in the first and second pre-declared analyses. No setting is tuned on any new subject.
- **Pipeline per subject.** Week-1 maps, week-2 operator / leakage / wedge maps, ring maps (week 13), week-3 fits (wedge; PSF stretched 0.85 / 1.15; wedge from the first and from the second half of each source run; ring), contamination maps (CWL-M/P, S2/S4/S8/Q for the wedges; CWL-M/P for the rings).

- **Unit of analysis and family structure.** The subject. Twins and siblings are not independent, so the **primary tests use one subject per family: the lowest subject ID of each family** (86 subjects). Secondary: all 158 subjects, with the same tests and, in addition, a family-level bootstrap (resample families) for the interval of the mean.
- **Inclusion per test.** A subject enters a test when every stratum the test needs has ≥ 30 vertices (as in weeks 3–9). A subject whose pipeline fails a check that the pilot pipeline also applied (operator column mass, coherent-vertex count > 0) is excluded from all tests and listed.
- **Statistics.** As in the pre-declared analyses: one-sided Wilcoxon signed-rank across subjects (direction stated per hypothesis), bootstrap 95% interval of the mean (10,000 resamples, seed 0), subjects in the predicted direction. Secondary tests Holm-corrected within their family of tests. The pilot results are not combined with the new cohort.

**Hypotheses**

Notation: Δ = residual(model) − residual(competitor) (negative: model better). Unless stated, the competitor is the one of the pre-declared analyses, upsampling + 1.6 mm mapping + 8 mm smoothing. Strata by the dose variable at the coarse grid among vertices with < 2% far leak at 1.6 mm; bins 0–2, 2–5, 5–10, 10–20, > 20%; D_x = Δ(highest bin with ≥ 30 vertices) − Δ(0–2% bin) for the dose variable x (far leak, CWL-M, CWL-P, CWL-S4, CWL-Q), computed by `summarize_cwl.py` / `summarize_cwl2.py` exactly as in `STATS_PLAN.md`. "All coherent" gains use the per-subject better of the two smoothing competitors, as in Section 4.8. Sources of the numbers per batch: `eval/hcp_cwl_summary.json`, `eval/hcp_cwl2_summary.json`, `eval/ring_cwl_summary.json` (ring), `eval/dose_*/hcp_dose_response.json` (far-leak bins, competitor upsampling + 8 mm smoothing) and `eval/*/week3_test_scores.json` (all-coherent and PSF rows).

**H1 — confirmatory replication (wedges).** At 2.4 and 3.2 mm, D_CWL-M < 0 (primary, as the first analysis' primary) and D_CWL-P < 0 (the coarse-data-only measure). Four tests; the primary family is {2.4 mm CWL-M, 3.2 mm CWL-M}. Secondary: D_far < 0; the overall gain Δ(all coherent) < 0; the subject counts and sizes of Section 4.3 restated.

**H2 — rings (same subjects, same acquisition).** The expectation before the pilot was that rings, which drive both banks of the calcarine alike, would show no far-leak dose-response. The exploratory look at the 15 pilot subjects (recorded below) did not bear this out: the ring runs have a far-leak dose-response of their own at 2.4 mm, as strong as the wedges', and a weaker one at 3.2 mm, and contamination predicts the ring gain at both grids. Eccentricity differs across much of the far cortex (foveal and peripheral representations are far apart on the sheet), so the far part of the kernel does carry different responses for rings too. The confirmatory hypotheses are therefore:

- H2a: at 3.2 mm, D_far(wedge) − D_far(ring) < 0 (the plain-leak dose-response is steeper for the stimulus with contrast across sulcal banks; pilot −0.016, 15/15). At 2.4 mm the same difference is reported with its interval, without a directional prediction (pilot −0.001, 8/15).

- H2b: D_CWL-M(ring) < 0 at 2.4 and 3.2 mm (contamination predicts the gain for rings as for wedges; pilot −0.028 and −0.027, 13/15 each).
- H2c (secondary): at 3.2 mm, D_CWL-M(ring) − D_far(ring) < 0 (for rings, contamination separates the vertices where the model helps more sharply than the plain leak; pilot −0.027 against −0.007).
- Descriptive: D_far(ring) with its interval at both grids; the model's overall gain for rings (pilot −0.005 and −0.012).

**H3 — source noise inside one dataset.** Fits from half a source run (4 cycles) against the same full-run reference. The expectation before the pilot was that, if part of the model's gain were denoising, the gain where no unmixing is needed would grow with source noise. The exploratory look showed the opposite direction: the gain in the 0–2% far-leak bin shrank with half runs (2.4 mm −0.0086 → −0.0045, 14 of 15 subjects shrank; 3.2 mm −0.0165 → −0.0140, 13 of 15), while the leak-dependent part of the gain did not change. The confirmatory hypotheses are therefore:

- H3a: Δ_clean(half) − Δ_clean(full) > 0 at 2.4 and 3.2 mm (the model's gain where no unmixing is needed decreases with source noise: it is not a denoising gain), where Δ_clean = Δ in the 0–2% far-leak bin of the far-leak dose-response (competitor upsampling + 8 mm smoothing) and "half" is the mean of the first- and second-half fits (one-sided).
- H3b: D_far(half) − D_far(full) (same source): the 95% interval includes 0 (the leak-dependent gain is unchanged). Reported as an interval, not a significance test.
- H3c (secondary): Δ_clean(half) < 0 at both grids (the model still beats the competitor with half the data; pilot 10/15 at 2.4 mm, 15/15 at 3.2 mm).
- Descriptive: the noise ceiling (split-half agreement) of half- and full-run sources.

**H4 — PSF sensitivity (from Section 4.8, now with n ≈ 86).** With the PSF assumed 15% too wide, Δ(all coherent) < 0 at 2.4 mm (the pilot's p = 0.23 case); with 15% too narrow, Δ(all coherent) < 0 at both grids (secondary). Competitor: the per-subject better of the two smoothing competitors, as in the rows B3–B6 of Section 4.8.

##### What the plan does not cover

- The leak-gated estimator, the averaging-kernel and known-truth analyses of week 3 (not recomputed for the cohort).
- Group maps of the gain on fs_LR (descriptive; produced from the same fits, no test).
- Bars (RETBAR1/2): not periodic, would need pRF fitting; not analysed.

##### Changes after hashing

None are allowed; a change found necessary is recorded as a deviation with its reason, as in week 11.

**Record of the exploratory look (15 pilot test subjects, before hashing)**

`results/week13/pilot_explore.md` / `.json`, from `results/week13/cohort/pilot/eval/`. The pilot pipeline reproduced the published pilot numbers (D_CWL-M −0.023 / −0.037 at 2.4 / 3.2 mm, 11/15 and 15/15; overall gain −0.007 / −0.013). New:

| quantity (pilot, exploratory) | 2.4 mm | 3.2 mm |
|---|---|---|
| D_far(ring) | −0.024 (13/15) | −0.007 (10/15) |
| D_far(wedge) − D_far(ring) | −0.001 (8/15) | −0.016 (15/15) |
| D_CWL-M(ring) | −0.028 (13/15) | −0.027 (13/15) |
| ring: overall gain | −0.005 (12/15) | −0.012 (14/15) |
| Δ_clean(full) → Δ_clean(half) | −0.0086 → −0.0045 | −0.0165 → −0.0140 |
| Δ_clean(half) − Δ_clean(full) | +0.0041 (14/15 > 0) | +0.0024 (13/15 > 0) |
| D_far(half) − D_far(full) | +0.001 [−0.003, +0.004] | −0.000 [−0.003, +0.002] |

H2 and H3 above were rewritten after this look; H1 and H4 were not changed. The pilot subjects are not part of the confirmatory cohort.

- Plan hash: `COHORT_PLAN.sha1` (SHA-1 and time), written before the first fit of any of the 158 subjects.

**S11.5 Amendment 1: sensitivity set (`COHORT_PLAN_AMENDMENT_1.md`, SHA-1 580b52b6906bf982c4923a91a19676622af06bce of the unredacted file, recorded 2026-10-03 after an interim look at 8 of 16 batches; verbatim except that the five subject IDs are withheld; SHA-256 of this redacted text 23d02dfec479db0d3367d659007d70ad69327c50e73d36b79c0e4bac3d43209e)**

Decided by the user after the interim look at 8 of 16 batches ("add the 6 as a sensitivity set and continue"). The hashed plan (`COHORT_PLAN.md`, SHA-1 21822393…) is not changed; this file is hashed separately (`COHORT_PLAN_AMENDMENT_1.sha1`).

**The set**

The six HCP 7T subjects excluded from the confirmatory cohort were [five subject IDs withheld in this copy: restricted family information] (relatives of the five tuning subjects; one of them also lacks the MSMAll spheres, which only the fs_LR group maps need) and [HCP ID withheld] (no concatenated retinotopy run in the HCP release; it cannot be processed by this pipeline). **The sensitivity set is therefore the five processable subjects**, run through the identical pipeline (stage 0, A, B, group) as `batch_17`.

**How it is used**

- It is **not** part of the primary or secondary tests of H1–H4, whose subject set stays as hashed (158 subjects, 86 families).
- Two additional, clearly labelled analyses:

1. the five alone, descriptive (means and intervals; no tests with n = 5);
2. a pooled sensitivity analysis of every row of the plan over cohort + sensitivity set (163 subjects), with the same family rule (lowest subject ID per family), to show whether excluding the tuning subjects' relatives changed any conclusion.

- Rationale for the original exclusion stands: these subjects share families with the five subjects on which the settings were chosen, so they are not independent of the tuning data. The sensitivity analysis quantifies how much that matters.

**Implementation**

`work/batches/batch_17.txt` on TRUBA; fetched by a one-off; stage 0 queued by it, fits by the fits pump; scored by the same group job. Locally, `src/week13/cohort_stats.py` excludes `batch_17` and `pilot` from the default batch list; the sensitivity analyses are run by naming the batches explicitly.

##### S11.6 Far-part ablation (`ABLATION_PLAN.md`, SHA-1 f81590bcf70e6309f4f61a7068dc312c4709521f, recorded 2026-10-04 before the first fit; SHA-256 a349f729b16412df8f3ce7b201af0057c07842938f29567ead15e7a51bf4df2b)

Both reviewer passes asked for the direct test of the paper's mechanism claim: invert with the far part of the forward operator removed and see whether the dose-dependent gain disappears while the gain where no far mixing exists stays.

**Fixed before the fits**

- **Subjects, data, settings.** The 15 development subjects (the week-3 test set), wedge runs (full), emulated 2.4 and 3.2 mm; prior 12 mm / 0.125 at 2.4 mm and 24 mm / 0.25 at 3.2 mm; competitor upsampling + 1.6 mm mapping + 8 mm smoothing; the cross-run residual against the other run's 1.6 mm map, as in week 3. Nothing is re-tuned.
- **Operators.** Full K as in week 3. *Near-only* K_near: for every voxel, the entries to unknowns that are far from the voxel's reference unknown — far = the other hemisphere or more than 15 mm along the sheet on the area-scaled sphere, the definition of the leakage maps; reference = the unknown with the largest |K| entry of that voxel — are set to zero. *Renormalised near* K_nearn: K_near with each voxel row rescaled to the signed row sum of K (the "far cortex responds like the near cortex" null). Everything else is identical: noise weights, prior, PCG tolerance 1e-4, rendering through the full 1.6 mm operator.
- **Quantities.** Per subject and grid, for the full, near-only and renormalised-near models and the competitor: the cross-run residual in the far-leak bins of the dose-response (coherent vertices clean at 1.6 mm: 0–2, 2–5, 5–10, 10–20, > 20%), over all coherent vertices, where leak > 10% and in the pericalcarine; Δ = model − competitor per bin; D_far = Δ(highest bin with ≥

30 vertices) − Δ(0–2%); Δ_clean = Δ(0–2%). Also the share of the operator's mass removed per voxel (descriptive).

- **Predictions and tests** (n = 15; one-sided Wilcoxon where a direction is stated; bootstrap 95% of the mean, 10,000 resamples, seed 0):
  - P1: the dose-dependent gain shrinks without the far part: D_far(near) − D_far(full) > 0 at both grids.
  - P2: the gain where no far mixing exists is unchanged: Δ_clean(near) − Δ_clean(full), 95% interval; |mean| < 0.003 expected.
  - P3: the near-only model is worse overall: gain(near) − gain(full) > 0 over all coherent vertices, at both grids.
  - P4: P1–P3 repeated for the renormalised variant.
- **Interpretation, stated in advance.** P1 and P2 both holding means the dose-dependent gain is far unmixing and the leak-independent gain is near-field (deconvolution within 15 mm and exact ribbon sampling). P1 failing (D_far unchanged without the far part) means the dose-response is not produced by the far part of the operator, and the mechanism claim is withdrawn. P2 failing (the clean-bin gain also shrinks) means the far part matters even where the far leak is < 2%.
- **Status.** Exploratory, reviewer-requested, on the development set; reported with its n in the main text and in full in the Supplementary material. Hash recorded in `ABLATION_PLAN.sha1` before the first fit; the job driver refuses to submit fits without it.

##### S11.7 Real coarser sessions (`SF_OPERATOR_PLAN.md`, SHA-1 b733b5af82f367c767be32066c27fc5106d53056, recorded 2026-10-04 before running; SHA-256 26c21615aadab922514ca4aea0c14afc6c95eeb93453ed2f1c80e9adfa96792a)

Section 4.1 validates the geometry of the operator's far part against *emulated* coarsening of the same HCP data. StudyForrest has real sessions of the same subjects at 7T 0.8, 1.4, 2.0 and 3.0 mm (provider-distortion-corrected; registered to the subject's FreeSurfer anatomy in week 4). Here the week-2 test is repeated across real sessions.

###### Fixed before running

- **Subjects and sessions.** The 7 StudyForrest subjects, 7T sessions only; every ordered pair fine → coarse (0.8 → 1.4, 2.0, 3.0; 1.4 → 2.0, 3.0; 2.0 → 3.0). Operators, ribbon weights, registrations and session grids as in week 4 (`sf_model.py`): the native acquisition kernel of each session, surfaces moved into each session's EPI space.
- **Field estimate.** As in week 2: the fine session's standard map divided by its partial-volume dilution (floor 0.3), on the unknowns covered by the fine session; no model fit is involved.
- **Predictors of the coarse session's standard map** (A_c = W_c K_c, split per vertex into near and far columns with the leakage-map definition: other hemisphere or > 15 mm on the area-

scaled FreeSurfer sphere): *naive* (the fine map itself), *near only* (A_near $\hat{f}$), *near + far* (A $\hat{f}$), *near renormalised* (A_near $\hat{f}$ scaled to the full row mass), *near + shuffled far* (the far contributions permuted across the vertices of the stratum, as in week 2).

- **Score and strata.** Residual energy fraction after the best real scaling per hemifield map (`resid_real`), on vertices covered by both sessions: all covered; visual cortex; visual cortex by the far leak of standard mapping at the coarse session (< 2%, > 5%, > 10%, > 20%); pericalcarine. A stratum counts with ≥ 50 vertices.
- **Predictions** (per pair type over subjects, and per subject pooled over its pairs; one-sided Wilcoxon where a direction is stated, n ≤ 7; the shuffled control calibrates the comparison):
    - D1: near + far beats near only where the predicted far leak is high (visual, leak > 10%): resid(full) < resid(near).
    - D2: the true far part beats a shuffled far part: resid(shuffled) > resid(full) in the same stratum.
    - D3: geometry, not mass: resid(full) < resid(renormalised near) in the same stratum.
    - Descriptive: the same over all visual vertices and where leak < 2% (expected: no difference).
- **Interpretation, stated in advance.** D1–D3 holding is the first validation of the far part on real coarser acquisitions (with the usual cross-session confounds raising every residual). D2 failing means the far part's geometry is not supported by real coarse data at these resolutions. Between-session differences in the pairs that involve 0.8 mm (slab coverage) may make those pairs uninformative; the 1.4 → 3.0 mm pair is the primary one.
- **Status.** Exploratory, reviewer-requested; hashed in `SF_OPERATOR_PLAN.sha1` before running.

##### S11.8 Kernel-width scan, re-tuned half-run settings, correlated noise, far criterion (`PLAN_BEFG.md`, SHA-1 69292d8515ca267491cd4acd80a09fc922be07e3, recorded 2026-10-05 before the first fit; SHA-256 9db5178b50ed1f44eed8bb1e7215a04ace71707bd2e8d2a4231ba1831e3e1c35)

###### B. PSF-scale scan (5 tuning subjects)

- **Fits.** Chosen settings (12 mm / 0.125 at 2.4 mm, 24 mm / 0.25 at 3.2 mm), wedge runs, with the inversion's coarse-grid kernels stretched by α = 0.7, 0.85, 1.15, 1.3 on all axes, and by 0.85 and 1.15 on the phase-encoding axis only (y) and on the slice axis only (z); α = 1 from the week-3 tuning fits. Data and rendering unchanged (as in the week-3 ±15% test).
- **Readout.** Cross-run residual over all coherent vertices, where leak > 10% (clean at 1.6 mm) and in the pericalcarine, per α and axis set; mean over the 5 subjects and the subject count favouring each α over α = 1.
- **Expectation stated in advance.** If the modelled native PSF is right, the residual is lowest at α = 1 on every axis set. The week-3 and cohort results (−15% fits slightly better) predict a minimum below 1; the axis sets say which axis drives it. Descriptive (n = 5); no test. The

PSF-component table (`psf_components.md`) gives each component's FWHM contribution and the provenance of each parameter.

**E. H3 with settings re-tuned for half-run sources**

- **E1 (5 tuning subjects).** Fits from each half run (4 cycles) at 16 prior settings per grid (range 6, 12, 24, 48 mm × amplitude scale 0.0625, 0.125, 0.25, 0.5), competitors at smoothing FWHM 2–16 mm; the week-3 selection rule (lowest mean residual over the 5 subjects, all coherent vertices, halves averaged) picks the half-run settings of the model, of standard + smoothing and of upsampling + smoothing.
- **E2 (15 development subjects).** Δ_clean(half) and D_far(half) recomputed with the re-tuned settings (model refitted at the re-tuned prior if it differs from the full-run one; competitors from the saved arrays), against Δ_clean(full) and D_far(full) at the full-run settings. H3a′: Δ_clean(half, re-tuned) − Δ_clean(full) > 0 (one-sided Wilcoxon, n = 15); H3b′ as an interval.
- **Rule.** If H3a′ holds, H3's conclusion ("not denoising") does not depend on the fixed-settings confound. If it reverses (the gain where no far mixing exists grows with noisier sources once both methods are tuned for them), the denoising explanation revives and the manuscript's statement is withdrawn.

**F. Correlated-noise likelihood on the HCP development set (15 subjects)**

- **Fits.** The week-6 likelihood (stationary noise correlation, spectrum from the data's own noise, applied with FFTs; `w6lib`) at the chosen settings, both grids; the spectrum from the complex Fourier components at the eight noise bins neighbouring the stimulus frequency, each scaled to unit variance per voxel, on the coarse grid of each run.
- **Readout.** Residual over all coherent vertices, where leak > 10%, in the far-leak bins; D_far; against the white-noise model and the competitor. Paired differences corr − white (two-sided, n = 15; descriptive).
- **Expectation stated in advance.** From week 6 (StudyForrest): small changes, if any, concentrated at coarse grids. The question is whether the misspecified white likelihood costs accuracy or changes the dose-response; either way the result is reported.

**G. The "far" criterion: true geodesics and the 15 mm threshold**

- **G1 (20 development subjects, operator only).** The leakage maps recomputed with the sphere proxy at thresholds 10, 15 and 20 mm, and with "far = other hemisphere only"; mean far leak, cortex above 5% and 10%, pericalcarine, at 1.6 / 2.4 / 3.2 mm.
- **G2 (5 tuning subjects).** The far criterion recomputed with true geodesic distances on the midthickness mesh (Dijkstra on the triangle graph, limit 25 mm; pairs beyond the limit are far) for the same operator rows; per-vertex agreement of the geodesic and sphere-proxy leak fractions (correlation, mean difference, share of vertices whose > 5% / > 10% label changes),

the share of operator mass whose far/near label flips, and the resulting group-figure differences.

- Descriptive.

Hashes in `PLAN_BEFG.sha1`, recorded before the first job of B, E and F was submitted (G is operator-only and descriptive).

##### S11.9 Addendum: the correlated-noise likelihood with its own prior settings (`PLAN_F2_ADDENDUM.md`, SHA-1 bc984c0bdeee88ec5181caafcffe1341b465a2ab, recorded 2026-10-05 after the first three subjects of F and before any F2 fit; SHA-256 0956c8def95d39bd974e633a988df782ddf8c26ec4522914ed3beae4bb91621b)

**Why, stated openly.** F as planned fits the correlated-noise likelihood at the prior settings chosen for the white-noise likelihood (12 mm / 0.125 at 2.4 mm, 24 mm / 0.25 at 3.2 mm). In the first three development subjects to finish, the correlated fit was worse than the white fit in every stratum (all coherent vertices +0.054 at 2.4 mm and +0.025 at 3.2 mm, 0 of 3 better), and the estimated noise spectrum is dominated by large spatial scales (480–630 times the white level in the lowest radial bin, 0.03–0.04 in the highest). Changing the likelihood changes the balance between data and prior, so a comparison at the white likelihood's settings confounds the likelihood with its tuning — the confound that E removes for H3. F2 removes it for F. It is added after seeing those three subjects; F itself is reported as planned.

**Design**

- **Fits.** The week-16 correlated-noise fit (`src/week16/hcp_corr.py`, unchanged: spectrum from the eight neighbouring noise bins of both runs, floor 0.1 of white, all valid voxels as rows) on a grid of prior settings, range $r \in \{6, 12, 24, 48\}$ mm × amplitude scale $s \in \{0.03125, 0.0625, 0.125, 0.25, 0.5, 1\}$, both grids (2.4, 3.2 mm), full wedge runs, for the 5 tuning subjects and the 15 development subjects. Scored with `src/week16/variant_eval.py` together with the subject's white-noise fits from week 3 (tuning subjects: the week-3 tuning grid; development subjects: the chosen settings and the competitors).
- **Selection (tuning subjects only).** Per grid, the correlated-likelihood setting with the lowest mean cross-run residual over all coherent vertices across the 5 tuning subjects — the week-3 rule. The white likelihood keeps its week-3 choice.
- **Comparison (15 development subjects).** corr(re-tuned) − white(chosen), per subject, over all coherent vertices (primary readout), where leak > 10% (clean at 1.6 mm), in the pericalcarine and in the far-leak bins, and for D_far; mean, subjects in favour, bootstrap 95% interval, two-sided Wilcoxon (descriptive, $n = 15$). Reported alongside: corr at the white settings (F, reproduced inside F2 as a check), and the best correlated setting chosen on the 15 development subjects themselves (an optimistic bound, not a test).
- **Rule.** If corr(re-tuned) does not have a lower mean residual than white(chosen) over all coherent vertices, the manuscript states that the white likelihood did not cost accuracy in HCP

emulation and that the correlated likelihood estimated from neighbouring frequency bins did not help, with or without re-tuning. If it does, the manuscript reports the improvement, its size and where it occurs, and the Limitations entry on the noise model is revised accordingly. Either way D_far under both likelihoods is reported.

Hash in `PLAN_F2_ADDENDUM.sha1`, recorded before the first F2 job is submitted.

**S11.10 External replication, NYU Retinotopy Dataset (`PLAN_NYU.md`, SHA-1 b815c95167c6-caa1bfe625566d0af9ecb0885d7a, recorded 2026-10-06 before the first fit; SHA-256 7dc2ff0f98cc1a560ed71dd2103b27d9ba62056f03c1e0ad5a7473bd817ba993)**

**Question**

Do the model's advantage over the best competitor and its growth with far leak and with contamination, found in HCP 7T emulation (15 development subjects; 158-subject confirmatory cohort), replicate in an independent dataset that differs in site, scanner (Siemens Prisma), field strength (3T), voxel size (2.0 mm), stimulus (sweeping bars instead of rotating wedges) and preprocessing (ours, from the raw data)?

**Data**

OpenNeuro ds003787 v1.0.0 (NYU Retinotopy Dataset; Himmelberg, Kurzawski, Benson, Pelli, Carrasco, Winawer; CC0): 44 subjects, 4–12 bar-mapping runs each (192 volumes, TR 1 s, 2.0 mm isotropic, multiband 6, phase encoding P→A, effective echo spacing 0.58 ms, 104 phase-encoding lines, no partial Fourier), one spin-echo AP/PA field-map pair per session, FreeSurfer 6 reconstructions (provided), and one aperture sequence (100 × 100 × 192, binary; identical in every run). Downloaded to hebb (`GEO_CONST/data/ds003787`, 97.3 GB subset: raw data, FreeSurfer, apertures, ROIs, the authors' pRF maps); nothing is written outside `/analysis/halilaziz/halil/GEO_CONST`.

**Split, fixed now:** `numpy.random.default_rng(17).permutation` over the 44 sorted subject IDs; the first 8 are the tuning subjects (sub-wlsubj057, 062, 084, 087, 094, 118, 121, 126), the other 36 are the test subjects (`nyu_split.json`).

**Exclusions (decided before any result):** a run is excluded if its maximum absolute translation relative to the reference exceeds 4 mm (two voxels); a subject is excluded if fewer than 4 runs remain, if no field-map pair is usable, or if the boundary-based registration fails in every start (cost > 0.8). Excluded runs and subjects are listed before any test-subject result is summarised.

**Preprocessing (per subject; FSL tools in the fMRIPrep container on hebb)**

1. topup on the AP/PA spin-echo pair (acquisition parameters from the sidecars) → field;

2. motion correction of every run to the run-1 single-band reference, itself registered rigidly to the P→A spin-echo image, in one spline resampling into the field-map grid (same 2.0 mm geometry);
3. distortion correction of every run with applytopup (P→A line, Jacobian modulation);
4. registration of the corrected session mean to the FreeSurfer anatomy: FLIRT from three starts, then boundary-based registration with the FreeSurfer white matter, lowest cost kept (as for ds002702);
5. brain mask as for ds002702.

**Polar-angle maps (the analogue of the HCP wedge Fourier coefficient)**

Per run and grid, a GLM with three stimulus regressors — the aperture's coverage of the stimulated field weighted by 1, cos θ and sin θ (θ = polar angle of each aperture pixel about the centre) — each convolved with the SPM double-gamma HRF, plus the temporal derivative of the first, Legendre drifts of order 0–3 and the six motion parameters. In the aperture sequence the three stimulus regressors are uncorrelated ($|r| < 0.01$). The complex map z = β_cos + i β_sin (percent signal change) is the first angular harmonic of each voxel's response: its phase is a linear estimate of the polar angle, as the phase of the wedge Fourier coefficient is in HCP. Noise variance of z per voxel: $\sigma^2$_res · [$(X^{T}X)^{-1}$_cos,cos + $(X^{T}X)^{-1}$_sin,sin], with $\sigma^2$_res the GLM residual variance in percent signal change. Halves: odd and even runs (A, B); z averaged within a half, variance divided by the number of runs.

**Emulation, operators, methods**

- **Emulation:** every corrected time series is coarsened frame by frame to 3.0 mm (×1.5) and 4.0 mm (×2.0) with the k-space crop and Tukey window ($\alpha = 0.5$) used for HCP, and the GLM is rerun on the coarse series.
- **Operators:** the FreeSurfer-anatomy operator of the ds002702 analysis (`kay_emul.py`: unknowns on the white–pial columns, Workbench ribbon weights per grid), with the acquisition PSF from the run sidecars (effective echo spacing, phase-encoding lines and axis; T2* 45 ms at 3T; taper 0.3, slice edge 0.15, interpolation blur 0.3 voxel) and the coarse kernels by the HCP emulation rule. Far leak and contamination exactly as in the paper (other hemisphere, or > 15 mm on the area-scaled registration sphere).
- **Prediction:** source half coarsened, target = the other half's native 2.0 mm standard map; both directions, averaged. Methods: standard mapping of the coarse data; + surface smoothing (FWHM 2, 4, 6, 8, 10, 12, 14, 16 mm); k-space upsampling to 2.0 mm; upsampling + smoothing (same FWHMs); the model (white-noise likelihood, Matérn prior; range $r \in \{6, 12, 24, 48\}$ mm × amplitude $s \in \{0.0625, 0.125, 0.25, 0.5\} \times \sigma_0$, $\sigma_0$ as for ds002702), rendered at 2.0 mm.

- **Score:** the HCP residual — energy of the target left after the best complex scaling of the prediction — per vertex set, averaged over the two directions.

**Vertex sets**

- **responsive:** vertices whose run-averaged native 2.0 mm standard-mapped time series (all runs) is explained by the three stimulus regressors with $R^2 \geq 0.2$ (after drifts and motion); the analogue of HCP's coherent vertices.
- **clean, leak > 10%:** responsive, far leak < 2% at 2.0 mm, far leak > 10% at the coarse grid.
- **pericalcarine:** responsive and in the FreeSurfer pericalcarine label.
- **Dose bins:** responsive vertices with < 2% far leak at 2.0 mm, binned by the dose variable at the coarse grid (0–2, 2–5, 5–10, 10–20, > 20%); a subject enters a bin with ≥ 30 vertices; D_x = Δ(highest bin with ≥ 30 vertices) − Δ(0–2% bin). Dose variables: far leak; CWL-M (contamination with the source half's native 2.0 mm map as the response estimate, normalised as in the paper).

**Selection (tuning subjects only)**

The week-3 rule: per grid, the model setting, the smoothing FWHM and the upsampling + smoothing FWHM with the lowest mean residual over responsive vertices of the 8 tuning subjects. The **competitor** of the tests is the better of the two chosen competitors on the tuning subjects. The choices are written to `SETTINGS_NYU.txt` before any test subject is fitted. The tuning subjects' runs may be used to find and fix implementation errors; the test subjects are not looked at until then.

**Hypotheses (36 test subjects; Δ = residual(model) − residual(competitor), negative favours the model)**

Primary, at both coarse grids (3.0 mm, ×1.5; 4.0 mm, ×2.0), one-sided Wilcoxon signed-rank, Holm across the six tests:

- **N1** — Δ over responsive vertices < 0 (the overall advantage replicates);
- **N2** — D_far < 0 (the gain grows with far leak);
- **N3** — D_CWL-M < 0 (the gain grows with contamination).

Secondary (Holm within the family): Δ(clean, leak > 10%) < 0; Δ(pericalcarine) < 0. Descriptive: the polar-angle error in degrees (mean absolute angular difference between the phases of prediction and target over responsive vertices); the far-leak figures of a 3T 2.0 mm acquisition and its 3.0 / 4.0 mm emulations (mean far leak, share of cortex above 5% and 10%, all included subjects). Statistics as in the confirmatory cohort: subjects in the predicted direction, mean, bootstrap 95% interval (10,000 resamples, seed 0), one-sided p.

**Decision rules**

- N1 holds at both grids (Holm): the paper states that the HCP emulation result replicates at another site, scanner, field strength, voxel size and stimulus. N1 fails at a grid: the paper states that it did not replicate there, with the numbers, and the abstract and Discussion are revised accordingly.
- N2 / N3: replicated or not, stated per grid as above.
- Whatever the outcome, the analysis is reported in the manuscript (a new results subsection and a supplement section) and the release.

**Deviations**

Any change to this plan (a failed preprocessing step, a changed threshold, an implementation fix that alters a definition) is written into `DEVIATIONS_NYU.md` with its reason and time before the test subjects' results are summarised.

Hash in `PLAN_NYU.sha1`, recorded before the first fit.

**Deviations (`DEVIATIONS_NYU.md`, recorded before any test-subject result)**

1. **Prior grid extended (2026-10-06, about 14:50 local; no test subject analysed).** In the first tuning subject (sub-wlsubj057, 3.0 mm, direction A→B) the model's residual fell steadily towards the planned grid's weakest prior and shortest range (best r = 6 mm, s = 0.5; e.g. s 0.0625 → 0.5 at r = 6: 0.633 → 0.528), so the optimum lay outside the planned grid r ∈ {6, 12, 24, 48} × s ∈ {0.0625, 0.125, 0.25, 0.5}. The grid was extended to r ∈ {3, 6, 12, 24, 48} × s ∈ {0.0625, 0.125, 0.25, 0.5, 1, 2, 4} for all tuning subjects. The same extension was needed for ds002702 (week 7), whose FreeSurfer-based pipeline uses the same $\sigma 0$ definition. The selection rule is unchanged.
2. **Implementation fixes in preprocessing (no change of definition).** (a) Three subjects (sub-wlsubj086, 087, 088) have no single-band reference; the motion-correction reference is then the mean of run 1's first 10 volumes. (b) Ten runs of one subject and one subject's field maps are named without the `acq-PA` / `run-01` entities; files are matched with or without them. (c) FSL tools that exit with status 0 after failing to read an image are caught by checking their output files.

**S12. Software and compute**

Python 3.12 (numpy, scipy, nibabel) inside the XCP-D container (pennlinc/xcp_d; Connectome Workbench `wb_command` 2.0.1 for ribbon weights and resampling). FSL 6 inside the fMRIPrep container (nipreps/fmriprep 24.1.1: mcflirt, FLIRT with BBR, fugue, fsl_prepare_fieldmap). Apptainer.

Compute ran on TRUBA (SLURM debug partition, 8–28 cores per job) and a lab server (Apptainer, NFS storage). Typical costs:

- operator and ribbon weights: 30–50 min per HCP subject (about 50 min on the cohort's 28-core jobs);
- one model fit: 1–25 min, depending on grid and prior;
- contamination maps: 4–20 min per cell.

Workbench's volume-to-surface mapping must not run concurrently on one node (it crashes); calls are serialised with a node-local lock.

The confirmatory cohort (S13) ran on TRUBA's debug partition in 17 batches of up to 10 subjects, three stages per subject (maps and operator; the wedge fit and its contamination maps; the PSF, half-run and ring fits) and one scoring job per batch; per subject about 50 min for the operator, 13–25 min per fit and about 10 min per contamination map on 28 cores. Jobs are named by stage and subject and resubmitted when the scheduler's per-user cap refuses them; a hash of the analysis plan must exist before any fit is submitted. Software inside the XCP-D image (built 2026-05-18): Python 3.12.12, numpy 2.2.6, scipy 1.15.2, nibabel 5.3.2; Apptainer 1.3.3. A re-run of the cohort downloads about 2.8 GB per subject (≈ 440 GB for 158 subjects). The reviewer-requested tests (S14) cost about 1 h per subject for the ablation (two operator variants, two grids), 45 min per subject for the three-kernel leakage maps and 1–2 h per StudyForrest subject (four session operators and weights). The second set (S15) cost, per tuning subject, eight stretched-kernel fits per grid (≈ 15 min each) and sixteen half-run prior settings per grid and half; per development subject, four half-run fits and two correlated-noise fits with their evaluations (≈ 2.5 h); the threshold maps 20 min and the geodesic check (Dijkstra from every vertex of the region, limit 25 mm, 28 cores) about 1 h per subject. The NYU replication (S16) ran on the lab server: preprocessing (FSL topup, mcflirt, applytopup, BBR) about 19 min per subject; per tuning subject 4.5–6 h for the operators, GLMs and the 35 prior settings at both grids; per test subject 28–77 min at the chosen settings (12 subjects in parallel, 6 threads each). The rerun of S16.3–S16.5 repeated the chosen settings for all 43 included subjects and the group maps took under a minute. The atlas analysis (S16.6) reran the 35 test subjects at the chosen settings; the server was shared and its cores clocked down, so it took a median of 4.0 h per subject (two queues, 23 subjects at a time).

##### S13. Confirmatory cohort (158 further HCP subjects)

Plan: S11.4–S11.5; methods: S17, Section 3.9; condensed table: S17, Section 4.9 and main text Section 4.4. The tables below are copied from the result files of `src/13_hcp_cohort/cohort_stats.py` (one row per pre-declared quantity; the primary set is one subject per family, the lowest subject ID; "all subjects" adds a family-level bootstrap of the mean). Negative means favour the model.

#### S13.1 Development set: the 15 held-out subjects of the main analyses, run through the batch pipeline (exploratory look before hashing)

15 subjects; no family rule

| hyp. | role | test | n | mean | median | in direction | bootstrap 95% | p | p (Holm) | all subjects: n, mean, family-bootstrap 95% |
|---|---|---|---|---|---|---|---|---|---|---|
| H1 | primary | D_CWL-M < 0, 2.4 mm | 15 | −0.0234 | −0.0280 | 11/15 | [−0.0384, −0.0077] | 0.008 | – | 15, −0.0234, [−0.0384, −0.0077] |
| H1 | primary | D_CWL-M < 0, 3.2 mm | 15 | −0.0366 | −0.0347 | 15/15 | [−0.0453, −0.0286] | < 0.001 | – | 15, −0.0366, [−0.0453, −0.0286] |
| H1 | secondary | D_CWL-P < 0, 2.4 mm | 15 | −0.0414 | −0.0377 | 14/15 | [−0.0564, −0.0273] | < 0.001 | < 0.001 | 15, −0.0414, [−0.0564, −0.0273] |
| H1 | secondary | D_far < 0, 2.4 mm | 15 | −0.0244 | −0.0255 | 12/15 | [−0.0370, −0.0115] | 0.003 | 0.007 | 15, −0.0244, [−0.0370, −0.0115] |
| H1 | secondary | gain, all coherent < 0, 2.4 mm | 15 | −0.0070 | −0.0071 | 12/15 | [−0.0113, −0.0025] | 0.006 | 0.007 | 15, −0.0070, [−0.0113, −0.0025] |
| H1 | descriptive | D_CWL-S4, 2.4 mm | 15 | −0.0276 | −0.0301 | 12/15 | [−0.0401, −0.0155] | 0.001 | – | 15, −0.0276, [−0.0401, −0.0155] |
| H1 | descriptive | D_CWL-Q, 2.4 mm | 15 | −0.0311 | −0.0237 | 14/15 | [−0.0418, −0.0210] | < 0.001 | – | 15, −0.0311, [−0.0418, −0.0210] |
| H1 | secondary | D_CWL-P < 0, 3.2 mm | 15 | −0.0365 | −0.0420 | 15/15 | [−0.0463, −0.0263] | < 0.001 | < 0.001 | 15, −0.0365, [−0.0463, −0.0263] |
| H1 | secondary | D_far < 0, 3.2 mm | 15 | −0.0229 | −0.0244 | 13/15 | [−0.0310, −0.0145] | < 0.001 | < 0.001 | 15, −0.0229, [−0.0310, −0.0145] |
| H1 | secondary | gain, all coherent < 0, 3.2 mm | 15 | −0.0127 | −0.0133 | 14/15 | [−0.0174, −0.0070] | 0.002 | 0.006 | 15, −0.0127, [−0.0174, −0.0070] |
| H1 | descriptive | D_CWL-S4, 3.2 mm | 15 | −0.0330 | −0.0289 | 15/15 | [−0.0410, −0.0251] | < 0.001 | – | 15, −0.0330, [−0.0410, −0.0251] |
| H1 | descriptive | D_CWL-Q, 3.2 mm | 15 | −0.0278 | −0.0301 | 15/15 | [−0.0366, −0.0193] | < 0.001 | – | 15, −0.0278, [−0.0366, −0.0193] |
| H2 | interval | D_far(wedge) − D_far(ring), 2.4 mm | 15 | −0.0007 | −0.0021 | 8/15 | [−0.0149, +0.0154] | 0.319 | – | 15, −0.0007, [−0.0149, +0.0154] |
| H2 | descriptive | D_far(ring), 2.4 mm | 15 | −0.0237 | −0.0318 | 13/15 | [−0.0341, −0.0134] | < 0.001 | – | 15, −0.0237, [−0.0341, −0.0134] |
| H2 | H2b | D_CWL-M(ring) < 0, 2.4 mm | 15 | −0.0277 | −0.0287 | 13/15 | [−0.0410, −0.0148] | < 0.001 | – | 15, −0.0277, [−0.0410, −0.0148] |
| H2 | H2a | D_far(wedge) − D_far(ring) < 0, 3.2 mm | 15 | −0.0160 | −0.0167 | 15/15 | [−0.0218, −0.0103] | < 0.001 | – | 15, −0.0160, [−0.0218, −0.0103] |
| H2 | descriptive | D_far(ring), 3.2 mm | 15 | −0.0068 | −0.0060 | 10/15 | [−0.0146, −0.0002] | 0.068 | – | 15, −0.0068, [−0.0146, −0.0002] |
| H2 | H2b | D_CWL-M(ring) < 0, 3.2 mm | 15 | −0.0270 | −0.0299 | 13/15 | [−0.0372, −0.0160] | < 0.001 | – | 15, −0.0270, [−0.0372, −0.0160] |
| H2 | secondary | D_CWL-M(ring) − D_far(ring) < 0, 3.2 mm | 15 | −0.0202 | −0.0184 | 14/15 | [−0.0280, −0.0123] | < 0.001 | < 0.001 | 15, −0.0202, [−0.0280, −0.0123] |
| H2 | descriptive | ring: gain, all coherent, 2.4 mm | 15 | −0.0053 | −0.0038 | 12/15 | [−0.0092, −0.0018] | 0.009 | – | 15, −0.0053, [−0.0092, −0.0018] |
| H2 | descriptive | ring: gain, all coherent, 3.2 mm | 15 | −0.0120 | −0.0103 | 14/15 | [−0.0174, −0.0065] | 0.001 | – | 15, −0.0120, [−0.0174, −0.0065] |
| H3 | H3a | Δ_clean(half) − Δ_clean(full) > 0, 2.4 mm | 15 | +0.0041 | +0.0059 | 14/15 | [+0.0009, +0.0064] | 0.004 | – | 15, +0.0041, [+0.0009, +0.0064] |
| H3 | interval | D_far(half) − D_far(full), 2.4 mm (interval should include 0 or be positive) | 15 | +0.0008 | +0.0005 | 6/15 | [−0.0027, +0.0045] | 0.640 | – | 15, +0.0008, [−0.0027, +0.0045] |
| H3 | descriptive | Δ_clean(full), 2.4 mm | 15 | −0.0086 | −0.0058 | 14/15 | [−0.0127, −0.0050] | < 0.001 | – | 15, −0.0086, [−0.0127, −0.0050] |
| H3 | secondary | Δ_clean(half) < 0, 2.4 mm | 15 | −0.0045 | −0.0047 | 10/15 | [−0.0088, −0.0003] | 0.047 | 0.047 | 15, −0.0045, [−0.0088, −0.0003] |
| H3 | H3a | Δ_clean(half) − Δ_clean(full) > 0, 3.2 mm | 15 | +0.0024 | +0.0027 | 13/15 | [−0.0001, +0.0045] | 0.011 | – | 15, +0.0024, [−0.0001, +0.0045] |

| hyp. | role | test | n | mean | median | in direction | bootstrap 95% | p | p (Holm) | all subjects: n, mean, family-bootstrap 95% |
|---|---|---|---|---|---|---|---|---|---|---|
| H3 | interval | D_far(half) − D_far(full), 3.2 mm (interval should include 0 or be positive) | 15 | −0.0003 | +0.0003 | 7/15 | [−0.0026, +0.0019] | 0.555 | – | 15, −0.0003, [−0.0026, +0.0019] |
| H3 | descriptive | Δ_clean(full), 3.2 mm | 15 | −0.0165 | −0.0152 | 15/15 | [−0.0212, −0.0120] | < 0.001 | – | 15, −0.0165, [−0.0212, −0.0120] |
| H3 | secondary | Δ_clean(half) < 0, 3.2 mm | 15 | −0.0140 | −0.0123 | 15/15 | [−0.0177, −0.0107] | < 0.001 | < 0.001 | 15, −0.0140, [−0.0177, −0.0107] |
| H4 | primary | PSF 15% too wide: gain, all coherent < 0, 2.4 mm | 0 | – | – | – | – | – | – | – |
| H4 | secondary | PSF 15% too wide: gain, all coherent < 0, 3.2 mm | 0 | – | – | – | – | – | – | – |
| H4 | secondary | PSF 15% too narrow: gain, all coherent < 0, 2.4 mm | 0 | – | – | – | – | – | – | – |
| H4 | secondary | PSF 15% too narrow: gain, all coherent < 0, 3.2 mm | 0 | – | – | – | – | – | – | – |

### S13.2 Confirmatory cohort in full

158 subjects with results (16 batches); primary set: 86 (lowest: one subject per family)

| hyp. | role | test | n | mean | median | in direction | bootstrap 95% | p | p (Holm) | all subjects: n, mean, family-bootstrap 95% |
|---|---|---|---|---|---|---|---|---|---|---|
| H1 | primary | D_CWL-M < 0, 2.4 mm | 86 | −0.0264 | −0.0260 | 64/86 | [−0.0332, −0.0197] | < 0.001 | – | 158, −0.0250, [−0.0310, −0.0191] |
| H1 | primary | D_CWL-M < 0, 3.2 mm | 86 | −0.0219 | −0.0183 | 77/86 | [−0.0262, −0.0177] | < 0.001 | – | 158, −0.0188, [−0.0224, −0.0152] |
| H1 | secondary | D_CWL-P < 0, 2.4 mm | 86 | −0.0339 | −0.0283 | 70/86 | [−0.0418, −0.0264] | < 0.001 | < 0.001 | 158, −0.0302, [−0.0365, −0.0242] |
| H1 | secondary | D_far < 0, 2.4 mm | 86 | −0.0201 | −0.0192 | 57/86 | [−0.0271, −0.0133] | < 0.001 | < 0.001 | 158, −0.0189, [−0.0246, −0.0135] |
| H1 | secondary | gain, all coherent < 0, 2.4 mm | 86 | −0.0087 | −0.0080 | 82/86 | [−0.0099, −0.0075] | < 0.001 | < 0.001 | 158, −0.0089, [−0.0099, −0.0079] |
| H1 | descriptive | D_CWL-S4, 2.4 mm | 86 | −0.0185 | −0.0195 | 61/86 | [−0.0254, −0.0117] | < 0.001 | – | 158, −0.0186, [−0.0248, −0.0128] |
| H1 | descriptive | D_CWL-Q, 2.4 mm | 86 | −0.0207 | −0.0181 | 62/86 | [−0.0278, −0.0137] | < 0.001 | – | 158, −0.0198, [−0.0249, −0.0149] |
| H1 | secondary | D_CWL-P < 0, 3.2 mm | 86 | −0.0211 | −0.0194 | 71/86 | [−0.0259, −0.0163] | < 0.001 | < 0.001 | 158, −0.0182, [−0.0221, −0.0143] |
| H1 | secondary | D_far < 0, 3.2 mm | 86 | −0.0049 | −0.0054 | 48/86 | [−0.0092, −0.0008] | 0.015 | 0.015 | 158, −0.0038, [−0.0072, −0.0004] |
| H1 | secondary | gain, all coherent < 0, 3.2 mm | 86 | −0.0128 | −0.0131 | 85/86 | [−0.0141, −0.0114] | < 0.001 | < 0.001 | 158, −0.0126, [−0.0136, −0.0115] |
| H1 | descriptive | D_CWL-S4, 3.2 mm | 86 | −0.0149 | −0.0153 | 67/86 | [−0.0190, −0.0109] | < 0.001 | – | 158, −0.0121, [−0.0155, −0.0086] |
| H1 | descriptive | D_CWL-Q, 3.2 mm | 86 | −0.0150 | −0.0141 | 66/86 | [−0.0194, −0.0107] | < 0.001 | – | 158, −0.0124, [−0.0158, −0.0091] |
| H2 | interval | D_far(wedge) − D_far(ring), 2.4 mm | 86 | −0.0076 | −0.0016 | 48/86 | [−0.0142, −0.0010] | 0.027 | – | 158, −0.0077, [−0.0126, −0.0028] |
| H2 | descriptive | D_far(ring), 2.4 mm | 86 | −0.0125 | −0.0080 | 53/86 | [−0.0195, −0.0060] | 0.002 | – | 158, −0.0112, [−0.0166, −0.0058] |
| H2 | H2b | D_CWL-M(ring) < 0, 2.4 mm | 86 | −0.0263 | −0.0260 | 71/86 | [−0.0327, −0.0199] | < 0.001 | – | 158, −0.0268, [−0.0326, −0.0209] |
| H2 | H2a | D_far(wedge) − D_far(ring) < 0, 3.2 mm | 86 | −0.0047 | −0.0047 | 46/86 | [−0.0087, −0.0006] | 0.010 | – | 158, −0.0041, [−0.0065, −0.0016] |
| H2 | descriptive | D_far(ring), 3.2 mm | 86 | −0.0002 | +0.0006 | 40/86 | [−0.0036, +0.0031] | 0.554 | – | 158, +0.0003, [−0.0027, +0.0032] |

| hyp. | role | test | n | mean | median | in direction | bootstrap 95% | p | p (Holm) | all subjects: n, mean, family-bootstrap 95% |
|---|---|---|---|---|---|---|---|---|---|---|
| H2 | H2b | D_CWL-M(ring) < 0, 3.2 mm | 86 | −0.0209 | −0.0206 | 79/86 | [−0.0243, −0.0176] | < 0.001 | – | 158, −0.0210, [−0.0239, −0.0179] |
| H2 | secondary | D_CWL-M(ring) − D_far(ring) < 0, 3.2 mm | 86 | −0.0207 | −0.0199 | 78/86 | [−0.0243, −0.0170] | < 0.001 | < 0.001 | 158, −0.0213, [−0.0243, −0.0182] |
| H2 | descriptive | ring: gain, all coherent, 2.4 mm | 86 | −0.0070 | −0.0076 | 76/86 | [−0.0085, −0.0056] | < 0.001 | – | 158, −0.0075, [−0.0087, −0.0064] |
| H2 | descriptive | ring: gain, all coherent, 3.2 mm | 86 | −0.0130 | −0.0131 | 84/86 | [−0.0144, −0.0117] | < 0.001 | – | 158, −0.0130, [−0.0141, −0.0119] |
| H3 | H3a | Δ_clean(half) − Δ_clean(full) > 0, 2.4 mm | 86 | +0.0032 | +0.0032 | 71/86 | [+0.0023, +0.0041] | < 0.001 | – | 158, +0.0032, [+0.0026, +0.0039] |
| H3 | interval | D_far(half) − D_far(full), 2.4 mm (interval should include 0 or be positive) | 86 | −0.0002 | +0.0003 | 42/86 | [−0.0020, +0.0016] | 0.578 | – | 158, +0.0004, [−0.0009, +0.0017] |
| H3 | descriptive | Δ_clean(full), 2.4 mm | 86 | −0.0096 | −0.0086 | 80/86 | [−0.0110, −0.0081] | < 0.001 | – | 158, −0.0103, [−0.0115, −0.0091] |
| H3 | secondary | Δ_clean(half) < 0, 2.4 mm | 86 | −0.0063 | −0.0063 | 71/86 | [−0.0082, −0.0046] | < 0.001 | < 0.001 | 158, −0.0070, [−0.0084, −0.0056] |
| H3 | H3a | Δ_clean(half) − Δ_clean(full) > 0, 3.2 mm | 86 | +0.0010 | +0.0007 | 53/86 | [+0.0002, +0.0018] | 0.007 | – | 158, +0.0010, [+0.0005, +0.0016] |
| H3 | interval | D_far(half) − D_far(full), 3.2 mm (interval should include 0 or be positive) | 86 | +0.0004 | +0.0004 | 39/86 | [−0.0007, +0.0015] | 0.671 | – | 158, −0.0002, [−0.0010, +0.0008] |
| H3 | descriptive | Δ_clean(full), 3.2 mm | 86 | −0.0161 | −0.0153 | 86/86 | [−0.0178, −0.0146] | < 0.001 | – | 158, −0.0160, [−0.0172, −0.0148] |
| H3 | secondary | Δ_clean(half) < 0, 3.2 mm | 86 | −0.0152 | −0.0149 | 85/86 | [−0.0168, −0.0136] | < 0.001 | < 0.001 | 158, −0.0150, [−0.0162, −0.0138] |
| H4 | primary | PSF 15% too wide: gain, all coherent < 0, 2.4 mm | 86 | −0.0038 | −0.0032 | 63/86 | [−0.0051, −0.0025] | < 0.001 | – | 158, −0.0041, [−0.0052, −0.0030] |
| H4 | secondary | PSF 15% too wide: gain, all coherent < 0, 3.2 mm | 86 | −0.0050 | −0.0056 | 64/86 | [−0.0065, −0.0036] | < 0.001 | < 0.001 | 158, −0.0048, [−0.0059, −0.0036] |
| H4 | secondary | PSF 15% too narrow: gain, all coherent < 0, 2.4 mm | 86 | −0.0109 | −0.0106 | 85/86 | [−0.0121, −0.0097] | < 0.001 | < 0.001 | 158, −0.0111, [−0.0121, −0.0101] |
| H4 | secondary | PSF 15% too narrow: gain, all coherent < 0, 3.2 mm | 86 | −0.0158 | −0.0164 | 85/86 | [−0.0171, −0.0145] | < 0.001 | < 0.001 | 158, −0.0156, [−0.0167, −0.0146] |

#### S13.3 Sensitivity analysis: cohort plus the five relatives of the tuning subjects

163 subjects with results (17 batches); primary set: 91 (lowest: one subject per family)

| hyp. | role | test | n | mean | median | in direction | bootstrap 95% | p | p (Holm) | all subjects: n, mean, family-bootstrap 95% |
|---|---|---|---|---|---|---|---|---|---|---|
| H1 | primary | D_CWL-M < 0, 2.4 mm | 91 | −0.0273 | −0.0257 | 69/91 | [−0.0340, −0.0207] | < 0.001 | – | 163, −0.0255, [−0.0313, −0.0198] |
| H1 | primary | D_CWL-M < 0, 3.2 mm | 91 | −0.0217 | −0.0178 | 82/91 | [−0.0256, −0.0178] | < 0.001 | – | 163, −0.0187, [−0.0222, −0.0153] |
| H1 | secondary | D_CWL-P < 0, 2.4 mm | 91 | −0.0336 | −0.0281 | 74/91 | [−0.0410, −0.0265] | < 0.001 | < 0.001 | 163, −0.0302, [−0.0361, −0.0243] |
| H1 | secondary | D_far < 0, 2.4 mm | 91 | −0.0192 | −0.0191 | 59/91 | [−0.0260, −0.0127] | < 0.001 | < 0.001 | 163, −0.0184, [−0.0240, −0.0131] |
| H1 | secondary | gain, all coherent < 0, 2.4 mm | 91 | −0.0087 | −0.0080 | 87/91 | [−0.0098, −0.0075] | < 0.001 | < 0.001 | 163, −0.0089, [−0.0099, −0.0078] |
| H1 | descriptive | D_CWL-S4, 2.4 mm | 91 | −0.0183 | −0.0194 | 64/91 | [−0.0250, −0.0118] | < 0.001 | – | 163, −0.0184, [−0.0242, −0.0128] |
| H1 | descriptive | D_CWL-Q, 2.4 mm | 91 | −0.0204 | −0.0178 | 66/91 | [−0.0271, −0.0136] | < 0.001 | – | 163, −0.0197, [−0.0246, −0.0149] |
| H1 | secondary | D_CWL-P < 0, 3.2 mm | 91 | −0.0210 | −0.0192 | 76/91 | [−0.0256, −0.0165] | < 0.001 | < 0.001 | 163, −0.0183, [−0.0219, −0.0145] |

| hyp. | role | test | n | mean | median | in direction | bootstrap 95% | p | p (Holm) | all subjects: n, mean, family-bootstrap 95% |
|---|---|---|---|---|---|---|---|---|---|---|
| H1 | secondary | D_far < 0, 3.2 mm | 91 | −0.0055 | −0.0061 | 53/91 | [−0.0096, −0.0014] | 0.006 | 0.006 | 163, −0.0041, [−0.0075, −0.0009] |
| H1 | secondary | gain, all coherent < 0, 3.2 mm | 91 | −0.0128 | −0.0132 | 90/91 | [−0.0141, −0.0115] | < 0.001 | < 0.001 | 163, −0.0126, [−0.0136, −0.0115] |
| H1 | descriptive | D_CWL-S4, 3.2 mm | 91 | −0.0153 | −0.0152 | 72/91 | [−0.0192, −0.0114] | < 0.001 | – | 163, −0.0124, [−0.0158, −0.0089] |
| H1 | descriptive | D_CWL-Q, 3.2 mm | 91 | −0.0150 | −0.0141 | 71/91 | [−0.0191, −0.0110] | < 0.001 | – | 163, −0.0125, [−0.0158, −0.0092] |
| H2 | interval | D_far(wedge) − D_far(ring), 2.4 mm | 91 | −0.0065 | −0.0012 | 50/91 | [−0.0131, +0.0001] | 0.048 | – | 163, −0.0071, [−0.0120, −0.0023] |
| H2 | descriptive | D_far(ring), 2.4 mm | 91 | −0.0127 | −0.0091 | 57/91 | [−0.0193, −0.0063] | < 0.001 | – | 163, −0.0113, [−0.0164, −0.0060] |
| H2 | H2b | D_CWL-M(ring) < 0, 2.4 mm | 91 | −0.0263 | −0.0259 | 75/91 | [−0.0327, −0.0199] | < 0.001 | – | 163, −0.0268, [−0.0325, −0.0208] |
| H2 | H2a | D_far(wedge) − D_far(ring) < 0, 3.2 mm | 91 | −0.0051 | −0.0053 | 49/91 | [−0.0089, −0.0013] | 0.004 | – | 163, −0.0043, [−0.0068, −0.0019] |
| H2 | descriptive | D_far(ring), 3.2 mm | 91 | −0.0003 | +0.0006 | 43/91 | [−0.0036, +0.0029] | 0.509 | – | 163, +0.0002, [−0.0026, +0.0031] |
| H2 | H2b | D_CWL-M(ring) < 0, 3.2 mm | 91 | −0.0206 | −0.0199 | 84/91 | [−0.0239, −0.0172] | < 0.001 | – | 163, −0.0207, [−0.0236, −0.0178] |
| H2 | secondary | D_CWL-M(ring) − D_far(ring) < 0, 3.2 mm | 91 | −0.0202 | −0.0205 | 81/91 | [−0.0238, −0.0167] | < 0.001 | < 0.001 | 163, −0.0210, [−0.0240, −0.0180] |
| H2 | descriptive | ring: gain, all coherent, 2.4 mm | 91 | −0.0070 | −0.0076 | 80/91 | [−0.0084, −0.0056] | < 0.001 | – | 163, −0.0075, [−0.0086, −0.0064] |
| H2 | descriptive | ring: gain, all coherent, 3.2 mm | 91 | −0.0130 | −0.0132 | 89/91 | [−0.0144, −0.0117] | < 0.001 | – | 163, −0.0130, [−0.0140, −0.0119] |
| H3 | H3a | Δ_clean(half) − Δ_clean(full) > 0, 2.4 mm | 91 | +0.0032 | +0.0033 | 75/91 | [+0.0024, +0.0041] | < 0.001 | – | 163, +0.0032, [+0.0026, +0.0039] |
| H3 | interval | D_far(half) − D_far(full), 2.4 mm (interval should include 0 or be positive) | 91 | +0.0000 | +0.0005 | 44/91 | [−0.0017, +0.0017] | 0.674 | – | 163, +0.0005, [−0.0008, +0.0017] |
| H3 | descriptive | Δ_clean(full), 2.4 mm | 91 | −0.0095 | −0.0087 | 84/91 | [−0.0110, −0.0082] | < 0.001 | – | 163, −0.0102, [−0.0114, −0.0090] |
| H3 | secondary | Δ_clean(half) < 0, 2.4 mm | 91 | −0.0063 | −0.0063 | 75/91 | [−0.0081, −0.0046] | < 0.001 | < 0.001 | 163, −0.0070, [−0.0083, −0.0057] |
| H3 | H3a | Δ_clean(half) − Δ_clean(full) > 0, 3.2 mm | 91 | +0.0010 | +0.0008 | 57/91 | [+0.0002, +0.0017] | 0.005 | – | 163, +0.0010, [+0.0005, +0.0016] |
| H3 | interval | D_far(half) − D_far(full), 3.2 mm (interval should include 0 or be positive) | 91 | +0.0005 | +0.0004 | 39/91 | [−0.0005, +0.0016] | 0.812 | – | 163, −0.0000, [−0.0009, +0.0009] |
| H3 | descriptive | Δ_clean(full), 3.2 mm | 91 | −0.0161 | −0.0153 | 91/91 | [−0.0177, −0.0146] | < 0.001 | – | 163, −0.0160, [−0.0172, −0.0148] |
| H3 | secondary | Δ_clean(half) < 0, 3.2 mm | 91 | −0.0152 | −0.0150 | 90/91 | [−0.0167, −0.0137] | < 0.001 | < 0.001 | 163, −0.0150, [−0.0161, −0.0138] |
| H4 | primary | PSF 15% too wide: gain, all coherent < 0, 2.4 mm | 91 | −0.0038 | −0.0035 | 67/91 | [−0.0050, −0.0025] | < 0.001 | – | 163, −0.0040, [−0.0051, −0.0030] |
| H4 | secondary | PSF 15% too wide: gain, all coherent < 0, 3.2 mm | 91 | −0.0051 | −0.0056 | 68/91 | [−0.0065, −0.0037] | < 0.001 | < 0.001 | 163, −0.0048, [−0.0060, −0.0037] |
| H4 | secondary | PSF 15% too narrow: gain, all coherent < 0, 2.4 mm | 91 | −0.0109 | −0.0106 | 90/91 | [−0.0120, −0.0097] | < 0.001 | < 0.001 | 163, −0.0111, [−0.0121, −0.0100] |
| H4 | secondary | PSF 15% too narrow: gain, all coherent < 0, 3.2 mm | 91 | −0.0158 | −0.0164 | 90/91 | [−0.0171, −0.0146] | < 0.001 | < 0.001 | 163, −0.0157, [−0.0166, −0.0146] |

#### S13.4 The five relatives alone (descriptive)

5 subjects; no family rule

| hyp. | role | test | n | mean | median | in direction | bootstrap 95% | p | p (Holm) | all subjects: n, mean, family-bootstrap 95% |
|---|---|---|---|---|---|---|---|---|---|---|
| H1 | primary | D_CWL-M < 0, 2.4 mm | 5 | −0.0429 | −0.0161 | 5/5 | [−0.0844, −0.0128] | 0.031 | – | 5, −0.0429, [−0.0844, −0.0128] |
| H1 | primary | D_CWL-M < 0, 3.2 mm | 5 | −0.0184 | −0.0152 | 5/5 | [−0.0270, −0.0122] | 0.031 | – | 5, −0.0184, [−0.0270, −0.0122] |
| H1 | secondary | D_CWL-P < 0, 2.4 mm | 5 | −0.0286 | −0.0202 | 4/5 | [−0.0542, −0.0080] | 0.062 | 0.188 | 5, −0.0286, [−0.0542, −0.0080] |
| H1 | secondary | D_far < 0, 2.4 mm | 5 | −0.0031 | +0.0087 | 2/5 | [−0.0278, +0.0216] | 0.500 | 0.500 | 5, −0.0031, [−0.0278, +0.0216] |
| H1 | secondary | gain, all coherent < 0, 2.4 mm | 5 | −0.0082 | −0.0110 | 5/5 | [−0.0117, −0.0041] | 0.031 | 0.188 | 5, −0.0082, [−0.0117, −0.0041] |
| H1 | descriptive | D_CWL-S4, 2.4 mm | 5 | −0.0135 | −0.0147 | 3/5 | [−0.0420, +0.0109] | 0.312 | – | 5, −0.0135, [−0.0420, +0.0109] |
| H1 | descriptive | D_CWL-Q, 2.4 mm | 5 | −0.0145 | −0.0100 | 4/5 | [−0.0285, −0.0022] | 0.062 | – | 5, −0.0145, [−0.0285, −0.0022] |
| H1 | secondary | D_CWL-P < 0, 3.2 mm | 5 | −0.0198 | −0.0143 | 5/5 | [−0.0282, −0.0128] | 0.031 | 0.188 | 5, −0.0198, [−0.0282, −0.0128] |
| H1 | secondary | D_far < 0, 3.2 mm | 5 | −0.0143 | −0.0088 | 5/5 | [−0.0225, −0.0069] | 0.031 | 0.188 | 5, −0.0143, [−0.0225, −0.0069] |
| H1 | secondary | gain, all coherent < 0, 3.2 mm | 5 | −0.0138 | −0.0169 | 5/5 | [−0.0178, −0.0091] | 0.031 | 0.188 | 5, −0.0138, [−0.0178, −0.0091] |
| H1 | descriptive | D_CWL-S4, 3.2 mm | 5 | −0.0212 | −0.0141 | 5/5 | [−0.0423, −0.0068] | 0.031 | – | 5, −0.0212, [−0.0423, −0.0068] |
| H1 | descriptive | D_CWL-Q, 3.2 mm | 5 | −0.0145 | −0.0079 | 5/5 | [−0.0262, −0.0057] | 0.031 | – | 5, −0.0145, [−0.0262, −0.0057] |
| H2 | interval | D_far(wedge) − D_far(ring), 2.4 mm | 5 | +0.0118 | +0.0213 | 2/5 | [−0.0174, +0.0400] | 0.844 | – | 5, +0.0118, [−0.0174, +0.0400] |
| H2 | descriptive | D_far(ring), 2.4 mm | 5 | −0.0149 | −0.0126 | 4/5 | [−0.0264, −0.0034] | 0.062 | – | 5, −0.0149, [−0.0264, −0.0034] |
| H2 | H2b | D_CWL-M(ring) < 0, 2.4 mm | 5 | −0.0259 | −0.0083 | 4/5 | [−0.0681, +0.0070] | 0.156 | – | 5, −0.0259, [−0.0681, +0.0070] |
| H2 | H2a | D_far(wedge) − D_far(ring) < 0, 3.2 mm | 5 | −0.0119 | −0.0104 | 3/5 | [−0.0231, −0.0008] | 0.156 | – | 5, −0.0119, [−0.0231, −0.0008] |
| H2 | descriptive | D_far(ring), 3.2 mm | 5 | −0.0024 | −0.0081 | 3/5 | [−0.0148, +0.0127] | 0.406 | – | 5, −0.0024, [−0.0148, +0.0127] |
| H2 | H2b | D_CWL-M(ring) < 0, 3.2 mm | 5 | −0.0142 | −0.0052 | 5/5 | [−0.0307, −0.0018] | 0.031 | – | 5, −0.0142, [−0.0307, −0.0018] |
| H2 | secondary | D_CWL-M(ring) − D_far(ring) < 0, 3.2 mm | 5 | −0.0117 | −0.0241 | 3/5 | [−0.0294, +0.0059] | 0.156 | 0.156 | 5, −0.0117, [−0.0294, +0.0059] |
| H2 | descriptive | ring: gain, all coherent, 2.4 mm | 5 | −0.0066 | −0.0107 | 4/5 | [−0.0121, −0.0008] | 0.094 | – | 5, −0.0066, [−0.0121, −0.0008] |
| H2 | descriptive | ring: gain, all coherent, 3.2 mm | 5 | −0.0130 | −0.0137 | 5/5 | [−0.0201, −0.0060] | 0.031 | – | 5, −0.0130, [−0.0201, −0.0060] |
| H3 | H3a | Δ_clean(half) − Δ_clean(full) > 0, 2.4 mm | 5 | +0.0035 | +0.0053 | 4/5 | [−0.0007, +0.0061] | 0.094 | – | 5, +0.0035, [−0.0007, +0.0061] |
| H3 | interval | D_far(half) − D_far(full), 2.4 mm (interval should include 0 or be positive) | 5 | +0.0038 | +0.0034 | 2/5 | [−0.0013, +0.0090] | 0.906 | – | 5, +0.0038, [−0.0013, +0.0090] |
| H3 | descriptive | Δ_clean(full), 2.4 mm | 5 | −0.0088 | −0.0113 | 4/5 | [−0.0150, −0.0027] | 0.062 | – | 5, −0.0088, [−0.0150, −0.0027] |
| H3 | secondary | Δ_clean(half) < 0, 2.4 mm | 5 | −0.0053 | −0.0069 | 4/5 | [−0.0096, −0.0001] | 0.094 | 0.094 | 5, −0.0053, [−0.0096, −0.0001] |
| H3 | H3a | Δ_clean(half) − Δ_clean(full) > 0, 3.2 mm | 5 | +0.0006 | +0.0015 | 4/5 | [−0.0019, +0.0022] | 0.312 | – | 5, +0.0006, [−0.0019, +0.0022] |
| H3 | interval | D_far(half) − D_far(full), 3.2 mm (interval should include 0 or be positive) | 5 | +0.0038 | +0.0037 | 0/5 | [+0.0016, +0.0063] | 1.000 | – | 5, +0.0038, [+0.0016, +0.0063] |
| H3 | descriptive | Δ_clean(full), 3.2 mm | 5 | −0.0159 | −0.0158 | 5/5 | [−0.0217, −0.0095] | 0.031 | – | 5, −0.0159, [−0.0217, −0.0095] |
| H3 | secondary | Δ_clean(half) < 0, 3.2 mm | 5 | −0.0152 | −0.0184 | 5/5 | [−0.0204, −0.0084] | 0.031 | 0.062 | 5, −0.0152, [−0.0204, −0.0084] |

| hyp. | role | test | n | mean | median | in direction | bootstrap 95% | p | p (Holm) | all subjects: n, mean, family-bootstrap 95% |
|---|---|---|---|---|---|---|---|---|---|---|
| H4 | primary | PSF 15% too wide: gain, all coherent < 0, 2.4 mm | 5 | −0.0031 | −0.0071 | 4/5 | [−0.0075, +0.0025] | 0.094 | – | 5, −0.0031, [−0.0075, +0.0025] |
| H4 | secondary | PSF 15% too wide: gain, all coherent < 0, 3.2 mm | 5 | −0.0068 | −0.0110 | 4/5 | [−0.0121, −0.0012] | 0.094 | 0.094 | 5, −0.0068, [−0.0121, −0.0012] |
| H4 | secondary | PSF 15% too narrow: gain, all coherent < 0, 2.4 mm | 5 | −0.0100 | −0.0118 | 5/5 | [−0.0133, −0.0068] | 0.031 | 0.094 | 5, −0.0100, [−0.0133, −0.0068] |
| H4 | secondary | PSF 15% too narrow: gain, all coherent < 0, 3.2 mm | 5 | −0.0162 | −0.0180 | 5/5 | [−0.0199, −0.0122] | 0.031 | 0.094 | 5, −0.0162, [−0.0199, −0.0122] |

##### S13.5 Gain by bin (main-text Fig. 7)

Model − competitor (upsampling + 1.6 mm mapping + 8 mm smoothing), mean ± SE over subjects, by bin of the dose variable at the coarse grid among vertices with < 2% far leak at 1.6 mm; a subject enters a bin with ≥ 30 vertices (`cohort_curves.md`, written by `src/13_hcp_cohort/fig_cohort.py` with the figure).

| grid | gain by | 0–2% | 2–5% | 5–10% | 10–20% | > 20% |
|---|---|---|---|---|---|---|
| 2.4 mm | far leak, wedges (full source run) | −0.0103 ± 0.0006 (n = 158) | −0.0118 ± 0.0007 (n = 158) | −0.0139 ± 0.0010 (n = 158) | −0.0144 ± 0.0015 (n = 158) | −0.0298 ± 0.0028 (n = 154) |
| 2.4 mm | far leak, wedges (half source run) | −0.0070 ± 0.0005 (n = 158) | −0.0079 ± 0.0006 (n = 158) | −0.0091 ± 0.0007 (n = 158) | −0.0111 ± 0.0012 (n = 158) | −0.0263 ± 0.0021 (n = 154) |
| 2.4 mm | far leak, rings | −0.0092 ± 0.0006 (n = 158) | −0.0101 ± 0.0006 (n = 158) | −0.0109 ± 0.0010 (n = 158) | −0.0079 ± 0.0013 (n = 157) | −0.0206 ± 0.0027 (n = 157) |
| 2.4 mm | contamination (CWL-M), wedges | −0.0077 ± 0.0006 (n = 158) | −0.0149 ± 0.0007 (n = 158) | −0.0202 ± 0.0010 (n = 158) | −0.0216 ± 0.0015 (n = 158) | −0.0334 ± 0.0029 (n = 153) |
| 2.4 mm | contamination (CWL-M), rings | −0.0077 ± 0.0006 (n = 158) | −0.0096 ± 0.0006 (n = 158) | −0.0137 ± 0.0009 (n = 158) | −0.0177 ± 0.0014 (n = 158) | −0.0353 ± 0.0028 (n = 154) |
| 3.2 mm | far leak, wedges (full source run) | −0.0160 ± 0.0006 (n = 158) | −0.0130 ± 0.0006 (n = 158) | −0.0176 ± 0.0008 (n = 158) | −0.0161 ± 0.0011 (n = 158) | −0.0198 ± 0.0017 (n = 158) |
| 3.2 mm | far leak, wedges (half source run) | −0.0150 ± 0.0005 (n = 158) | −0.0113 ± 0.0005 (n = 158) | −0.0157 ± 0.0006 (n = 158) | −0.0155 ± 0.0008 (n = 158) | −0.0189 ± 0.0013 (n = 158) |
| 3.2 mm | far leak, rings | −0.0157 ± 0.0006 (n = 158) | −0.0132 ± 0.0006 (n = 158) | −0.0162 ± 0.0008 (n = 158) | −0.0137 ± 0.0010 (n = 158) | −0.0152 ± 0.0014 (n = 157) |
| 3.2 mm | contamination (CWL-M), wedges | −0.0109 ± 0.0006 (n = 158) | −0.0130 ± 0.0005 (n = 158) | −0.0189 ± 0.0006 (n = 158) | −0.0242 ± 0.0010 (n = 158) | −0.0297 ± 0.0018 (n = 158) |
| 3.2 mm | contamination (CWL-M), rings | −0.0129 ± 0.0007 (n = 158) | −0.0107 ± 0.0005 (n = 158) | −0.0147 ± 0.0006 (n = 158) | −0.0198 ± 0.0008 (n = 158) | −0.0340 ± 0.0014 (n = 157) |

##### S13.6 The dose-response contrast under other bin conventions

158 subjects, primary set 86. Per-subject gains Δ by bin (model − upsampling + 8 mm smoothing; vertices clean at 1.6 mm) from the contamination summaries of the cohort batches; a bin counts when it has ≥ 30 vertices. "D (as planned)": highest available bin − 0–2% bin. One-sided Wilcoxon (negative predicted), bootstrap 95% of the mean. Source: `src/14_robustness_checks/dose_sensitivity.py`.

| stimulus | dose variable | grid | definition | n | mean | in direction | bootstrap 95% | p |
|---|---|---|---|---|---|---|---|---|
| wedges | contamination (CWL-M) | 2.4 mm | D (as planned) | 86 | −0.0264 | 64/86 | [−0.0332, −0.0197] | < 0.001 |
| wedges | contamination (CWL-M) | 2.4 mm | reference bin 2–5% | 86 | −0.0184 | 59/86 | [−0.0253, −0.0116] | < 0.001 |
| wedges | contamination (CWL-M) | 2.4 mm | top bin fixed at 10–20% | 86 | −0.0191 | 69/86 | [−0.0232, −0.0151] | < 0.001 |

| stimulus | dose variable | grid | definition | n | mean | in direction | bootstrap 95% | p |
|---|---|---|---|---|---|---|---|---|
| wedges | contamination (CWL-M) | 2.4 mm | slope per bin step | 86 | −0.0064 | 67/86 | [−0.0080, −0.0049] | < 0.001 |
| wedges | contamination (CWL-M) | 2.4 mm | Spearman ρ (Δ vs bin) | 86 | −0.4907 | 63/86 | [−0.6047, −0.3733] | < 0.001 |
| wedges | contamination (CWL-M) | 2.4 mm | highest available bin | 86 | > 20% in 83, 10–20% in 3, lower in 0 | | | |
| wedges | contamination (CWL-M) | 3.2 mm | D (as planned) | 86 | −0.0219 | 77/86 | [−0.0262, −0.0177] | < 0.001 |
| wedges | contamination (CWL-M) | 3.2 mm | reference bin 2–5% | 86 | −0.0196 | 73/86 | [−0.0241, −0.0154] | < 0.001 |
| wedges | contamination (CWL-M) | 3.2 mm | top bin fixed at 10–20% | 86 | −0.0152 | 79/86 | [−0.0174, −0.0130] | < 0.001 |
| wedges | contamination (CWL-M) | 3.2 mm | slope per bin step | 86 | −0.0057 | 81/86 | [−0.0066, −0.0048] | < 0.001 |
| wedges | contamination (CWL-M) | 3.2 mm | Spearman ρ (Δ vs bin) | 86 | −0.6826 | 77/86 | [−0.7605, −0.5988] | < 0.001 |
| wedges | contamination (CWL-M) | 3.2 mm | highest available bin | 86 | > 20% in 86, 10–20% in 0, lower in 0 | | | |
| wedges | contamination, coarse data only (CWL-P) | 2.4 mm | D (as planned) | 86 | −0.0339 | 70/86 | [−0.0418, −0.0264] | < 0.001 |
| wedges | contamination, coarse data only (CWL-P) | 2.4 mm | reference bin 2–5% | 86 | −0.0267 | 63/86 | [−0.0344, −0.0194] | < 0.001 |
| wedges | contamination, coarse data only (CWL-P) | 2.4 mm | top bin fixed at 10–20% | 86 | −0.0179 | 67/86 | [−0.0225, −0.0135] | < 0.001 |
| wedges | contamination, coarse data only (CWL-P) | 2.4 mm | slope per bin step | 86 | −0.0079 | 69/86 | [−0.0097, −0.0062] | < 0.001 |
| wedges | contamination, coarse data only (CWL-P) | 2.4 mm | Spearman ρ (Δ vs bin) | 86 | −0.5523 | 69/86 | [−0.6581, −0.4430] | < 0.001 |
| wedges | contamination, coarse data only (CWL-P) | 2.4 mm | highest available bin | 86 | > 20% in 83, 10–20% in 3, lower in 0 | | | |
| wedges | contamination, coarse data only (CWL-P) | 3.2 mm | D (as planned) | 86 | −0.0211 | 71/86 | [−0.0259, −0.0163] | < 0.001 |
| wedges | contamination, coarse data only (CWL-P) | 3.2 mm | reference bin 2–5% | 86 | −0.0200 | 71/86 | [−0.0249, −0.0153] | < 0.001 |
| wedges | contamination, coarse data only (CWL-P) | 3.2 mm | top bin fixed at 10–20% | 86 | −0.0125 | 73/86 | [−0.0149, −0.0101] | < 0.001 |
| wedges | contamination, coarse data only (CWL-P) | 3.2 mm | slope per bin step | 86 | −0.0054 | 77/86 | [−0.0064, −0.0043] | < 0.001 |
| wedges | contamination, coarse data only (CWL-P) | 3.2 mm | Spearman ρ (Δ vs bin) | 86 | −0.6035 | 73/86 | [−0.6895, −0.5116] | < 0.001 |
| wedges | contamination, coarse data only (CWL-P) | 3.2 mm | highest available bin | 86 | > 20% in 86, 10–20% in 0, lower in 0 | | | |
| wedges | plain far leak | 2.4 mm | D (as planned) | 86 | −0.0201 | 57/86 | [−0.0271, −0.0133] | < 0.001 |
| wedges | plain far leak | 2.4 mm | reference bin 2–5% | 86 | −0.0179 | 58/86 | [−0.0249, −0.0112] | < 0.001 |
| wedges | plain far leak | 2.4 mm | top bin fixed at 10–20% | 86 | −0.0057 | 55/86 | [−0.0096, −0.0017] | 0.005 |
| wedges | plain far leak | 2.4 mm | slope per bin step | 86 | −0.0044 | 56/86 | [−0.0060, −0.0029] | < 0.001 |
| wedges | plain far leak | 2.4 mm | Spearman ρ (Δ vs bin) | 86 | −0.2698 | 55/86 | [−0.3896, −0.1500] | < 0.001 |
| wedges | plain far leak | 2.4 mm | highest available bin | 86 | > 20% in 84, 10–20% in 2, lower in 0 | | | |
| wedges | plain far leak | 3.2 mm | D (as planned) | 86 | −0.0049 | 48/86 | [−0.0092, −0.0008] | 0.015 |
| wedges | plain far leak | 3.2 mm | reference bin 2–5% | 86 | −0.0087 | 59/86 | [−0.0131, −0.0044] | < 0.001 |

| stimulus | dose variable | grid | definition | n | mean | in direction | bootstrap 95% | p |
|---|---|---|---|---|---|---|---|---|
| wedges | plain far leak | 3.2 mm | top bin fixed at 10–20% | 86 | +0.0002 | 41/86 | [−0.0023, +0.0026] | 0.642 |
| wedges | plain far leak | 3.2 mm | slope per bin step | 86 | −0.0013 | 51/86 | [−0.0023, −0.0004] | 0.006 |
| wedges | plain far leak | 3.2 mm | Spearman ρ (Δ vs bin) | 86 | −0.1233 | 50/86 | [−0.2430, −0.0000] | 0.027 |
| wedges | plain far leak | 3.2 mm | highest available bin | 86 | > 20% in 86, 10–20% in 0, lower in 0 | | | |
| rings | contamination (CWL-M) | 2.4 mm | D (as planned) | 86 | −0.0263 | 71/86 | [−0.0327, −0.0199] | < 0.001 |
| rings | contamination (CWL-M) | 2.4 mm | reference bin 2–5% | 86 | −0.0248 | 71/86 | [−0.0308, −0.0188] | < 0.001 |
| rings | contamination (CWL-M) | 2.4 mm | top bin fixed at 10–20% | 86 | −0.0111 | 68/86 | [−0.0148, −0.0074] | < 0.001 |
| rings | contamination (CWL-M) | 2.4 mm | slope per bin step | 86 | −0.0062 | 73/86 | [−0.0076, −0.0047] | < 0.001 |
| rings | contamination (CWL-M) | 2.4 mm | Spearman ρ (Δ vs bin) | 86 | −0.5372 | 71/86 | [−0.6488, −0.4163] | < 0.001 |
| rings | contamination (CWL-M) | 2.4 mm | highest available bin | 86 | > 20% in 83, 10–20% in 3, lower in 0 | | | |
| rings | contamination (CWL-M) | 3.2 mm | D (as planned) | 86 | −0.0209 | 79/86 | [−0.0243, −0.0176] | < 0.001 |
| rings | contamination (CWL-M) | 3.2 mm | reference bin 2–5% | 86 | −0.0238 | 82/86 | [−0.0273, −0.0205] | < 0.001 |
| rings | contamination (CWL-M) | 3.2 mm | top bin fixed at 10–20% | 86 | −0.0071 | 67/86 | [−0.0093, −0.0049] | < 0.001 |
| rings | contamination (CWL-M) | 3.2 mm | slope per bin step | 86 | −0.0052 | 80/86 | [−0.0060, −0.0043] | < 0.001 |
| rings | contamination (CWL-M) | 3.2 mm | Spearman ρ (Δ vs bin) | 86 | −0.6581 | 79/86 | [−0.7395, −0.5686] | < 0.001 |
| rings | contamination (CWL-M) | 3.2 mm | highest available bin | 86 | > 20% in 85, 10–20% in 1, lower in 0 | | | |
| rings | plain far leak | 2.4 mm | D (as planned) | 86 | −0.0125 | 53/86 | [−0.0195, −0.0060] | 0.002 |
| rings | plain far leak | 2.4 mm | reference bin 2–5% | 86 | −0.0119 | 50/86 | [−0.0186, −0.0056] | 0.003 |
| rings | plain far leak | 2.4 mm | top bin fixed at 10–20% | 85 | −0.0001 | 44/85 | [−0.0032, +0.0028] | 0.369 |
| rings | plain far leak | 2.4 mm | slope per bin step | 86 | −0.0022 | 50/86 | [−0.0038, −0.0005] | 0.004 |
| rings | plain far leak | 2.4 mm | Spearman ρ (Δ vs bin) | 86 | −0.1233 | 50/86 | [−0.2593, +0.0093] | 0.046 |
| rings | plain far leak | 2.4 mm | highest available bin | 86 | > 20% in 85, 10–20% in 0, lower in 1 | | | |
| rings | plain far leak | 3.2 mm | D (as planned) | 86 | −0.0002 | 40/86 | [−0.0036, +0.0031] | 0.554 |
| rings | plain far leak | 3.2 mm | reference bin 2–5% | 86 | −0.0032 | 50/86 | [−0.0064, +0.0000] | 0.026 |
| rings | plain far leak | 3.2 mm | top bin fixed at 10–20% | 86 | +0.0029 | 31/86 | [+0.0005, +0.0052] | 0.994 |
| rings | plain far leak | 3.2 mm | slope per bin step | 86 | −0.0001 | 43/86 | [−0.0009, +0.0007] | 0.465 |
| rings | plain far leak | 3.2 mm | Spearman ρ (Δ vs bin) | 86 | +0.0360 | 42/86 | [−0.0860, +0.1570] | 0.698 |
| rings | plain far leak | 3.2 mm | highest available bin | 86 | > 20% in 85, 10–20% in 1, lower in 0 | | | |

#### H2a: D_far(wedges) − D_far(rings) at 3.2 mm under each definition

| definition | n | mean | in direction | bootstrap 95% | p |
|---|---|---|---|---|---|
| D (as planned) | 86 | −0.0047 | 46/86 | [−0.0087, −0.0006] | 0.010 |
| reference bin 2–5% | 86 | −0.0054 | 47/86 | [−0.0093, −0.0015] | 0.009 |
| top bin fixed at 10–20% | 86 | −0.0027 | 51/86 | [−0.0048, −0.0005] | 0.013 |
| slope per bin step | 86 | −0.0012 | 49/86 | [−0.0021, −0.0003] | 0.004 |
| Spearman ρ (Δ vs bin) | 86 | −0.1593 | 44/86 | [−0.2837, −0.0337] | 0.008 |

##### S13.7 The eight primary tests adjusted across their family (post hoc)

The cohort plan (S11.4) fixed eight primary one-sided tests and no correction across them (Holm only within each family of secondary tests). This table, added after the reviewer round of 2026-10-27, applies Holm's step-down and Bonferroni corrections across the eight, from the exact p-values of `cohort_final.json` (primary set: one subject per family, n = 86; one-sided Wilcoxon signed-rank; bootstrap 95% interval of the mean). Source: `src/13_hcp_cohort/primary_holm.py`.

| no. | hyp. | test | n | mean | in direction | bootstrap 95% | p (one-sided) | p (Holm, 8 tests) | p (Bonferroni, 8 tests) |
|---|---|---|---|---|---|---|---|---|---|
| P1 | H1 | D_CWL-M < 0, 2.4 mm | 86 | −0.0264 | 64/86 | [−0.0332, −0.0197] | < 0.001 | < 0.001 | < 0.001 |
| P2 | H1 | D_CWL-M < 0, 3.2 mm | 86 | −0.0219 | 77/86 | [−0.0262, −0.0177] | < 0.001 | < 0.001 | < 0.001 |
| P3 | H2 | D_far(wedge) − D_far(ring) < 0, 3.2 mm | 86 | −0.0047 | 46/86 | [−0.0087, −0.0006] | 0.010 | 0.013 | 0.077 |
| P4 | H2 | D_CWL-M(ring) < 0, 2.4 mm | 86 | −0.0263 | 71/86 | [−0.0327, −0.0199] | < 0.001 | < 0.001 | < 0.001 |
| P5 | H2 | D_CWL-M(ring) < 0, 3.2 mm | 86 | −0.0209 | 79/86 | [−0.0243, −0.0176] | < 0.001 | < 0.001 | < 0.001 |
| P6 | H3 | Δ_clean(half) − Δ_clean(full) > 0, 2.4 mm | 86 | +0.0032 | 71/86 | [+0.0023, +0.0041] | < 0.001 | < 0.001 | < 0.001 |
| P7 | H3 | Δ_clean(half) − Δ_clean(full) > 0, 3.2 mm | 86 | +0.0010 | 53/86 | [+0.0002, +0.0018] | 0.007 | 0.013 | 0.053 |
| P8 | H4 | PSF 15% too wide: gain, all coherent < 0, 2.4 mm | 86 | −0.0038 | 63/86 | [−0.0051, −0.0025] | < 0.001 | < 0.001 | < 0.001 |

Largest Holm-adjusted p: 0.013 (P3 and P7; the two smallest effects). All eight remain below 0.05 under Holm; under Bonferroni P3 and P7 do not (0.053 and 0.077).

#### S14. Reviewer-requested tests: far-part ablation, realistic kernels, real coarser sessions

Three analyses asked for by the reviewer round, each run under a plan hashed first (S11.6, S11.7; the leakage maps of S14.2 are descriptive). Tables are copied from the result files.

##### S14.1 Far-part ablation of the inversion (15 development subjects)

Cross-run residual against the other run's 1.6 mm map, mean over subjects; Δ = model − competitor (upsampling + 8 mm smoothing); D_far = Δ(highest bin ≥ 30 vertices) − Δ(0–2%). "near": far entries of the operator removed per voxel; "nearn": the same, rows renormalised. Source: `src/15_far_part_tests/ablation_eval.py` from `ablation_eval/`.

**Emulated 2.4 mm**

####### Residuals (mean over subjects)

| stratum | model, full operator | model, near-only operator | model, near-only renormalised | upsampling + 1.6 mm mapping + smoothing 8 mm | standard + smoothing 10 mm | upsampling + 1.6 mm mapping | standard mapping |
|---|---|---|---|---|---|---|---|
| all coherent | 0.3131 | 0.3126 | 0.3133 | 0.3212 | 0.3412 | 0.3556 | 0.4218 |
| pericalcarine | 0.2398 | 0.2397 | 0.2399 | 0.2482 | 0.2685 | 0.2774 | 0.3677 |
| clean, leak > 10% | 0.3428 | 0.3434 | 0.3449 | 0.3671 | 0.3986 | 0.3935 | 0.5295 |
| 0–2% | 0.3094 | 0.3090 | 0.3094 | 0.3180 | 0.3368 | 0.3497 | 0.4128 |
| 2–5% | 0.2855 | 0.2851 | 0.2854 | 0.2958 | 0.3171 | 0.3203 | 0.3863 |
| 5–10% | 0.3005 | 0.2993 | 0.3004 | 0.3148 | 0.3401 | 0.3399 | 0.4325 |
| 10–20% | 0.3389 | 0.3390 | 0.3408 | 0.3582 | 0.3873 | 0.3919 | 0.5155 |
| > 20% | 0.3260 | 0.3283 | 0.3297 | 0.3611 | 0.3996 | 0.3631 | 0.5295 |

####### Gain over the competitor by far-leak bin (mean over subjects)

| model | 0–2% | 2–5% | 5–10% | 10–20% | > 20% | all coherent | D_far | Δ_clean |
|---|---|---|---|---|---|---|---|---|
| model, full operator | −0.0086 | −0.0103 | −0.0143 | −0.0193 | −0.0351 | −0.0082 | −0.0244 | −0.0086 |
| model, near-only operator | −0.0090 | −0.0108 | −0.0155 | −0.0192 | −0.0328 | −0.0086 | −0.0219 | −0.0090 |
| model, near-only renormalised | −0.0086 | −0.0104 | −0.0144 | −0.0175 | −0.0314 | −0.0079 | −0.0214 | −0.0086 |

####### Predictions

| prediction | n | mean | median | in direction | bootstrap 95% | p (one-sided) | p (two-sided) |
|---|---|---|---|---|---|---|---|
| P1 D_far(near) − D_far(full) > 0 | 15 | +0.0025 | +0.0016 | 10/15 | [+0.0001, +0.0050] | 0.060 | 0.121 |
| P2 Δ_clean(near) − Δ_clean(full) (interval) | 15 | −0.0004 | −0.0003 | 2/15 | [−0.0006, −0.0002] | – | 0.001 |
| P3 gain(near) − gain(full) > 0, all coherent | 15 | −0.0005 | −0.0003 | 2/15 | [−0.0007, −0.0002] | 0.999 | 0.002 |
| top-bin gain(near) − top-bin gain(full) > 0 | 14 | +0.0023 | +0.0015 | 9/14 | [−0.0003, +0.0051] | 0.086 | 0.173 |
| P1 D_far(nearn) − D_far(full) > 0 | 15 | +0.0030 | +0.0026 | 9/15 | [−0.0010, +0.0071] | 0.151 | 0.303 |
| P2 Δ_clean(nearn) − Δ_clean(full) (interval) | 15 | +0.0000 | −0.0001 | 5/15 | [−0.0004, +0.0004] | – | 0.890 |
| P3 gain(nearn) − gain(full) > 0, all coherent | 15 | +0.0002 | +0.0003 | 10/15 | [−0.0003, +0.0007] | 0.126 | 0.252 |
| top-bin gain(nearn) − top-bin gain(full) > 0 | 14 | +0.0037 | +0.0030 | 9/14 | [−0.0004, +0.0080] | 0.097 | 0.194 |

**Emulated 3.2 mm**

####### Residuals (mean over subjects)

| stratum | model, full operator | model, near-only operator | model, near-only renormalised | upsampling + 1.6 mm mapping + smoothing 8 mm | standard + smoothing 10 mm | upsampling + 1.6 mm mapping | standard mapping |
|---|---|---|---|---|---|---|---|
| all coherent | 0.3248 | 0.3250 | 0.3249 | 0.3385 | 0.3520 | 0.3798 | 0.4579 |
| pericalcarine | 0.2505 | 0.2512 | 0.2503 | 0.2628 | 0.2746 | 0.3052 | 0.4089 |
| clean, leak > 10% | 0.3439 | 0.3440 | 0.3456 | 0.3712 | 0.3921 | 0.4111 | 0.5336 |
| 0–2% | 0.3256 | 0.3260 | 0.3258 | 0.3421 | 0.3554 | 0.3738 | 0.4990 |
| 2–5% | 0.3022 | 0.3026 | 0.3023 | 0.3160 | 0.3302 | 0.3501 | 0.3937 |
| 5–10% | 0.3113 | 0.3117 | 0.3110 | 0.3299 | 0.3470 | 0.3656 | 0.4355 |
| 10–20% | 0.3355 | 0.3358 | 0.3365 | 0.3539 | 0.3736 | 0.3944 | 0.4943 |
| > 20% | 0.3514 | 0.3518 | 0.3545 | 0.3908 | 0.4126 | 0.4270 | 0.5707 |

####### Gain over the competitor by far-leak bin (mean over subjects)

| model | 0–2% | 2–5% | 5–10% | 10–20% | > 20% | all coherent | D_far | Δ_clean |
|---|---|---|---|---|---|---|---|---|
| model, full operator | −0.0165 | −0.0137 | −0.0186 | −0.0184 | −0.0394 | −0.0137 | −0.0229 | −0.0165 |
| model, near-only operator | −0.0160 | −0.0133 | −0.0181 | −0.0181 | −0.0390 | −0.0135 | −0.0229 | −0.0160 |
| model, near-only renormalised | −0.0162 | −0.0136 | −0.0188 | −0.0174 | −0.0363 | −0.0137 | −0.0200 | −0.0162 |

####### Predictions

| prediction | n | mean | median | in direction | bootstrap 95% | p (one-sided) | p (two-sided) |
|---|---|---|---|---|---|---|---|
| P1 D_far(near) − D_far(full) > 0 | 15 | −0.0000 | −0.0021 | 7/15 | [−0.0022, +0.0022] | 0.467 | 0.934 |
| P2 Δ_clean(near) − Δ_clean(full) (interval) | 15 | +0.0004 | +0.0003 | 11/15 | [+0.0001, +0.0007] | – | 0.041 |
| P3 gain(near) − gain(full) > 0, all coherent | 15 | +0.0002 | +0.0003 | 9/15 | [−0.0002, +0.0006] | 0.262 | 0.524 |
| top-bin gain(near) − top-bin gain(full) > 0 | 15 | +0.0004 | −0.0007 | 7/15 | [−0.0018, +0.0026] | 0.339 | 0.679 |
| P1 D_far(nearn) − D_far(full) > 0 | 15 | +0.0028 | +0.0044 | 10/15 | [−0.0019, +0.0071] | 0.053 | 0.107 |
| P2 Δ_clean(nearn) − Δ_clean(full) (interval) | 15 | +0.0002 | +0.0003 | 9/15 | [−0.0002, +0.0007] | – | 0.359 |
| P3 gain(nearn) − gain(full) > 0, all coherent | 15 | +0.0000 | +0.0004 | 10/15 | [−0.0007, +0.0007] | 0.402 | 0.804 |
| top-bin gain(nearn) − top-bin gain(full) > 0 | 15 | +0.0031 | +0.0047 | 10/15 | [−0.0019, +0.0075] | 0.053 | 0.107 |

##### S14.2 Leakage of standard mapping for realistic native-acquisition kernels (20 development subjects)

Area-weighted means per hemisphere, averaged over hemispheres and subjects (± SD over subjects). "Emulated": the kernel of the k-space emulation used throughout the paper (native 1.6 mm kernel times a Tukey window at the coarse Nyquist). "Native": the week-2 native PSF model evaluated for an acquisition at that voxel size (Tukey-tapered sinc in plane, T2* decay over the echo train along PE with the same echo spacing and FOV, erf slice profile, interpolation blur). "PF 7/8": the same with the PE band reduced to 7/8 (the magnitude PSF of a zero-filled partial-Fourier reconstruction). Far = other hemisphere or > 15 mm on the area-scaled sphere. Source: `src/15_far_part_tests/leak_realistic.py`; per-subject files in `leak_realistic/`.

##### Emulated grid 2.4 mm

| quantity | emulated kernel (Tukey window; as published) | native-acquisition kernel at that voxel size | native kernel with partial Fourier 7/8 (magnitude approximation) |
|---|---|---|---|
| effective PSF FWHM x / y / z (mm) | 3.83 / 4.08 / 3.96 | 3.77 / 3.83 / 3.74 | 3.77 / 4.32 / 3.74 |
| mean far leak (%) | 2.56 ± 0.54 | 1.27 ± 0.56 | 2.05 ± 0.71 |
| cortex with far leak > 5% (%) | 17.95 ± 3.01 | 14.93 ± 2.82 | 18.03 ± 3.36 |
| > 10% (%) | 9.76 ± 2.28 | 7.98 ± 2.00 | 9.47 ± 2.27 |
| mean gross far weight (%) | 11.78 ± 0.87 | 13.27 ± 0.85 | 14.15 ± 0.89 |
| cortex with gross far weight > 20% (%) | 17.88 ± 2.74 | 21.68 ± 2.75 | 24.50 ± 2.83 |
| pericalcarine far leak (%) | 5.03 ± 1.42 | 3.56 ± 1.40 | 4.76 ± 1.67 |
| dilution of a unit field | 0.733 ± 0.015 | 0.751 ± 0.014 | 0.740 ± 0.014 |

##### Emulated grid 3.2 mm

| quantity | emulated kernel (Tukey window; as published) | native-acquisition kernel at that voxel size | native kernel with partial Fourier 7/8 (magnitude approximation) |
|---|---|---|---|
| effective PSF FWHM x / y / z (mm) | 5.07 / 5.36 / 5.11 | 4.99 / 4.98 / 4.94 | 4.99 / 5.65 / 4.94 |
| mean far leak (%) | 3.62 ± 0.69 | 2.42 ± 0.83 | 3.78 ± 1.06 |
| cortex with far leak > 5% (%) | 25.85 ± 2.70 | 24.26 ± 3.03 | 29.83 ± 4.19 |
| > 10% (%) | 16.84 ± 2.52 | 15.18 ± 2.46 | 18.07 ± 2.87 |
| mean gross far weight (%) | 21.10 ± 1.23 | 24.87 ± 1.07 | 25.99 ± 1.09 |
| cortex with gross far weight > 20% (%) | 45.61 ± 3.67 | 58.60 ± 3.04 | 61.85 ± 2.97 |
| pericalcarine far leak (%) | 6.45 ± 1.82 | 5.21 ± 1.95 | 7.01 ± 2.23 |
| dilution of a unit field | 0.685 ± 0.017 | 0.697 ± 0.016 | 0.686 ± 0.015 |

#### S14.3 The operator's far part on real coarser StudyForrest sessions (7 subjects)

Residual energy fraction of the coarse session's standard map predicted from the fine session's field estimate through the coarse session's operator (mean over subjects); tests: D1 full < near, D2 shuffled > full, D3 renorm > full (one-sided Wilcoxon).

### r08->r14

| stratum | n subj. | n vertices (mean) | fine map itself | near part only | near + far | near, renormalised | near + shuffled far | D1 full – near | D2 shuffled – full | D3 renorm – full |
|---|---|---|---|---|---|---|---|---|---|---|
| all covered | 7 | 73092 | 0.642 | 0.519 | 0.517 | 0.520 | 0.521 | −0.0020 (6/7, p 0.023) | +0.0035 (7/7, p 0.008) | +0.0023 (7/7, p 0.008) |
| visual cortex | 7 | 26564 | 0.495 | 0.390 | 0.388 | 0.390 | 0.391 | −0.0020 (6/7, p 0.039) | +0.0028 (7/7, p 0.008) | +0.0018 (7/7, p 0.008) |
| visual, far leak < 2% | 7 | 20689 | 0.482 | 0.381 | 0.381 | 0.381 | 0.381 | −0.0001 (5/7, p 0.078) | +0.0001 (4/7, p 0.188) | +0.0001 (6/7, p 0.109) |
| visual, > 5% | 7 | 3488 | 0.614 | 0.483 | 0.466 | 0.483 | 0.491 | −0.0171 (6/7, p 0.016) | +0.0251 (7/7, p 0.008) | +0.0172 (7/7, p 0.008) |
| visual, > 10% | 7 | 1883 | 0.635 | 0.500 | 0.477 | 0.500 | 0.515 | −0.0232 (6/7, p 0.016) | +0.0385 (7/7, p 0.008) | +0.0232 (7/7, p 0.008) |
| visual, > 20% | 7 | 462 | 0.605 | 0.470 | 0.442 | 0.470 | 0.508 | −0.0288 (6/7, p 0.023) | +0.0663 (7/7, p 0.008) | +0.0283 (6/7, p 0.039) |
| all covered, > 10% | 7 | 3488 | 0.725 | 0.600 | 0.574 | 0.597 | 0.619 | −0.0264 (7/7, p 0.008) | +0.0451 (7/7, p 0.008) | +0.0235 (7/7, p 0.008) |
| pericalcarine | 7 | 4416 | 0.415 | 0.315 | 0.312 | 0.316 | 0.317 | −0.0025 (5/7, p 0.148) | +0.0047 (6/7, p 0.023) | +0.0036 (7/7, p 0.008) |

### r08->r20

| stratum | n subj. | n vertices (mean) | fine map itself | near part only | near + far | near, renormalised | near + shuffled far | D1 full – near | D2 shuffled – full | D3 renorm – full |
|---|---|---|---|---|---|---|---|---|---|---|
| all covered | 7 | 70074 | 0.632 | 0.480 | 0.476 | 0.480 | 0.484 | −0.0044 (7/7, p 0.008) | +0.0079 (7/7, p 0.008) | +0.0041 (7/7, p 0.008) |
| visual cortex | 7 | 25015 | 0.505 | 0.384 | 0.380 | 0.383 | 0.387 | −0.0043 (7/7, p 0.008) | +0.0071 (7/7, p 0.008) | +0.0033 (7/7, p 0.008) |
| visual, far leak < 2% | 7 | 16463 | 0.496 | 0.379 | 0.379 | 0.380 | 0.380 | −0.0001 (4/7, p 0.344) | +0.0002 (4/7, p 0.109) | +0.0005 (7/7, p 0.008) |
| visual, > 5% | 7 | 4766 | 0.556 | 0.412 | 0.391 | 0.413 | 0.432 | −0.0202 (7/7, p 0.008) | +0.0402 (7/7, p 0.008) | +0.0214 (7/7, p 0.008) |

| stratum | n subj. | n vertices (mean) | fine map itself | near part only | near + far | near, renormalised | near + shuffled far | D1 full − near | D2 shuffled − full | D3 renorm − full |
|---|---|---|---|---|---|---|---|---|---|---|
| visual, > 10% | 7 | 3126 | 0.584 | 0.438 | 0.408 | 0.439 | 0.469 | −0.0299 (7/7, p 0.008) | +0.0607 (7/7, p 0.008) | +0.0311 (7/7, p 0.008) |
| visual, > 20% | 7 | 1362 | 0.607 | 0.475 | 0.426 | 0.473 | 0.527 | −0.0483 (7/7, p 0.008) | +0.1010 (7/7, p 0.008) | +0.0467 (7/7, p 0.008) |
| all covered, > 10% | 7 | 7165 | 0.691 | 0.543 | 0.508 | 0.539 | 0.575 | −0.0351 (7/7, p 0.008) | +0.0663 (7/7, p 0.008) | +0.0304 (7/7, p 0.008) |
| pericalcarine | 7 | 4116 | 0.445 | 0.325 | 0.320 | 0.326 | 0.332 | −0.0049 (3/7, p 0.289) | +0.0114 (7/7, p 0.008) | +0.0063 (6/7, p 0.016) |

# r08->r30

| stratum | n subj. | n vertices (mean) | fine map itself | near part only | near + far | near, renormalised | near + shuffled far | D1 full − near | D2 shuffled − full | D3 renorm − full |
|---|---|---|---|---|---|---|---|---|---|---|
| all covered | 7 | 57221 | 0.673 | 0.485 | 0.471 | 0.477 | 0.497 | −0.0141 (7/7, p 0.008) | +0.0262 (7/7, p 0.008) | +0.0069 (7/7, p 0.008) |
| visual cortex | 7 | 19186 | 0.546 | 0.390 | 0.375 | 0.381 | 0.401 | −0.0144 (7/7, p 0.008) | +0.0258 (7/7, p 0.008) | +0.0058 (7/7, p 0.008) |
| visual, far leak < 2% | 7 | 7240 | 0.559 | 0.394 | 0.395 | 0.395 | 0.395 | +0.0010 (2/7, p 0.961) | −0.0005 (3/7, p 0.852) | −0.0005 (4/7, p 0.711) |
| visual, > 5% | 7 | 7143 | 0.551 | 0.397 | 0.365 | 0.385 | 0.428 | −0.0320 (7/7, p 0.008) | +0.0637 (7/7, p 0.008) | +0.0199 (7/7, p 0.008) |
| visual, > 10% | 7 | 4804 | 0.584 | 0.418 | 0.377 | 0.409 | 0.468 | −0.0407 (7/7, p 0.008) | +0.0903 (7/7, p 0.008) | +0.0316 (7/7, p 0.008) |
| visual, > 20% | 7 | 2941 | 0.588 | 0.417 | 0.369 | 0.412 | 0.494 | −0.0482 (7/7, p 0.008) | +0.1251 (7/7, p 0.008) | +0.0434 (7/7, p 0.008) |
| all covered, > 10% | 7 | 14225 | 0.716 | 0.538 | 0.493 | 0.524 | 0.594 | −0.0450 (7/7, p 0.008) | +0.1012 (7/7, p 0.008) | +0.0305 (7/7, p 0.008) |
| pericalcarine | 7 | 3150 | 0.463 | 0.288 | 0.271 | 0.281 | 0.307 | −0.0169 (5/7, p 0.078) | +0.0364 (7/7, p 0.008) | +0.0106 (7/7, p 0.008) |

# r14->r20

| stratum | n subj. | n vertices (mean) | fine map itself | near part only | near + far | near, renormalised | near + shuffled far | D1 full − near | D2 shuffled − full | D3 renorm − full |
|---|---|---|---|---|---|---|---|---|---|---|
| all covered | 7 | 145365 | 0.424 | 0.378 | 0.375 | 0.377 | 0.382 | −0.0024 (5/7, p 0.055) | +0.0073 (7/7, p 0.008) | +0.0018 (6/7, p 0.039) |
| visual cortex | 7 | 30855 | 0.261 | 0.213 | 0.210 | 0.212 | 0.218 | −0.0026 (5/7, p 0.055) | +0.0075 (7/7, p 0.008) | +0.0022 (6/7, p 0.023) |
| visual, far leak < 2% | 7 | 20323 | 0.260 | 0.207 | 0.207 | 0.207 | 0.207 | −0.0000 (3/7, p 0.531) | +0.0002 (6/7, p 0.023) | +0.0003 (6/7, p 0.016) |
| visual, > 5% | 7 | 5643 | 0.293 | 0.255 | 0.242 | 0.255 | 0.285 | −0.0123 (5/7, p 0.078) | +0.0422 (7/7, p 0.008) | +0.0122 (6/7, p 0.055) |
| visual, > 10% | 7 | 3598 | 0.312 | 0.274 | 0.256 | 0.275 | 0.322 | −0.0175 (5/7, p 0.055) | +0.0658 (7/7, p 0.008) | +0.0185 (6/7, p 0.055) |
| visual, > 20% | 7 | 1509 | 0.362 | 0.323 | 0.293 | 0.326 | 0.404 | −0.0305 (5/7, p 0.078) | +0.1109 (7/7, p 0.008) | +0.0330 (6/7, p 0.055) |
| all covered, > 10% | 7 | 10979 | 0.420 | 0.387 | 0.367 | 0.382 | 0.437 | −0.0195 (6/7, p 0.023) | +0.0698 (7/7, p 0.008) | +0.0143 (6/7, p 0.078) |
| pericalcarine | 7 | 4169 | 0.206 | 0.163 | 0.160 | 0.162 | 0.171 | −0.0029 (4/7, p 0.234) | +0.0108 (7/7, p 0.008) | +0.0023 (6/7, p 0.016) |

# r14->r30

| stratum | n subj. | n vertices (mean) | fine map itself | near part only | near + far | near, renormalised | near + shuffled far | D1 full − near | D2 shuffled − full | D3 renorm − full |
|---|---|---|---|---|---|---|---|---|---|---|
| all covered | 7 | 121817 | 0.559 | 0.491 | 0.481 | 0.486 | 0.506 | −0.0103 (7/7, p 0.008) | +0.0247 (7/7, p 0.008) | +0.0047 (6/7, p 0.016) |
| visual cortex | 7 | 23770 | 0.383 | 0.303 | 0.289 | 0.296 | 0.320 | −0.0141 (7/7, p 0.008) | +0.0304 (7/7, p 0.008) | +0.0064 (7/7, p 0.008) |
| visual, far leak < 2% | 7 | 8963 | 0.396 | 0.303 | 0.304 | 0.304 | 0.304 | +0.0014 (1/7, p 0.984) | −0.0008 (2/7, p 0.891) | −0.0006 (2/7, p 0.922) |
| visual, > 5% | 7 | 8689 | 0.379 | 0.319 | 0.286 | 0.306 | 0.365 | −0.0324 (7/7, p 0.008) | +0.0788 (7/7, p 0.008) | +0.0197 (7/7, p 0.008) |
| visual, > 10% | 7 | 5689 | 0.406 | 0.344 | 0.303 | 0.337 | 0.417 | −0.0411 (7/7, p 0.008) | +0.1135 (7/7, p 0.008) | +0.0333 (7/7, p 0.008) |
| visual, > 20% | 7 | 3381 | 0.423 | 0.363 | 0.310 | 0.358 | 0.466 | −0.0522 (7/7, p 0.008) | +0.1562 (7/7, p 0.008) | +0.0478 (7/7, p 0.008) |
| all covered, > 10% | 7 | 26332 | 0.565 | 0.516 | 0.480 | 0.504 | 0.590 | −0.0355 (7/7, p 0.008) | +0.1096 (7/7, p 0.008) | +0.0241 (6/7, p 0.016) |
| pericalcarine | 7 | 3176 | 0.299 | 0.219 | 0.201 | 0.207 | 0.243 | −0.0185 (6/7, p 0.023) | +0.0420 (7/7, p 0.008) | +0.0063 (7/7, p 0.008) |

# r20->r30

| stratum | n subj. | n vertices (mean) | fine map itself | near part only | near + far | near, renormalised | near + shuffled far | D1 full − near | D2 shuffled − full | D3 renorm − full |
|---|---|---|---|---|---|---|---|---|---|---|
| all covered | 7 | 151880 | 0.516 | 0.449 | 0.438 | 0.443 | 0.461 | −0.0114 (7/7, p 0.008) | +0.0225 (7/7, p 0.008) | +0.0047 (7/7, p 0.008) |
| visual cortex | 7 | 24101 | 0.330 | 0.258 | 0.243 | 0.249 | 0.274 | −0.0149 (7/7, p 0.008) | +0.0303 (7/7, p 0.008) | +0.0056 (6/7, p 0.016) |
| visual, far leak < 2% | 7 | 9035 | 0.348 | 0.255 | 0.256 | 0.256 | 0.256 | +0.0009 (2/7, p 0.922) | −0.0003 (3/7, p 0.656) | −0.0003 (3/7, p 0.656) |
| visual, > 5% | 7 | 8867 | 0.324 | 0.278 | 0.244 | 0.262 | 0.322 | −0.0335 (7/7, p 0.008) | +0.0780 (7/7, p 0.008) | +0.0181 (7/7, p 0.008) |
| visual, > 10% | 7 | 5802 | 0.335 | 0.292 | 0.250 | 0.280 | 0.363 | −0.0424 (7/7, p 0.008) | +0.1133 (7/7, p 0.008) | +0.0302 (7/7, p 0.008) |
| visual, > 20% | 7 | 3414 | 0.341 | 0.302 | 0.249 | 0.293 | 0.408 | −0.0530 (7/7, p 0.008) | +0.1591 (7/7, p 0.008) | +0.0440 (7/7, p 0.008) |
| all covered, > 10% | 7 | 31295 | 0.505 | 0.464 | 0.425 | 0.451 | 0.531 | −0.0388 (7/7, p 0.008) | +0.1055 (7/7, p 0.008) | +0.0258 (7/7, p 0.008) |
| pericalcarine | 7 | 3060 | 0.263 | 0.189 | 0.168 | 0.175 | 0.209 | −0.0205 (7/7, p 0.008) | +0.0409 (7/7, p 0.008) | +0.0066 (6/7, p 0.016) |

### pooled over pairs

| stratum | n subj. | n vertices (mean) | fine map itself | near part only | near + far | near, renormalised | near + shuffled far | D1 full − near | D2 shuffled − full | D3 renorm − full |
|---|---|---|---|---|---|---|---|---|---|---|
| all covered | 7 | 103241 | 0.574 | 0.467 | 0.460 | 0.464 | 0.475 | −0.0074 (7/7, p 0.008) | +0.0154 (7/7, p 0.008) | +0.0041 (7/7, p 0.008) |
| visual cortex | 7 | 24915 | 0.420 | 0.323 | 0.314 | 0.319 | 0.332 | −0.0087 (7/7, p 0.008) | +0.0173 (7/7, p 0.008) | +0.0042 (7/7, p 0.008) |
| visual, far leak < 2% | 7 | 13785 | 0.424 | 0.320 | 0.320 | 0.320 | 0.320 | +0.0005 (2/7, p 0.961) | −0.0002 (3/7, p 0.766) | −0.0001 (3/7, p 0.711) |
| visual, > 5% | 7 | 6433 | 0.453 | 0.357 | 0.333 | 0.351 | 0.387 | −0.0246 (7/7, p 0.008) | +0.0547 (7/7, p 0.008) | +0.0181 (7/7, p 0.008) |
| visual, > 10% | 7 | 4150 | 0.476 | 0.378 | 0.345 | 0.373 | 0.426 | −0.0325 (7/7, p 0.008) | +0.0804 (7/7, p 0.008) | +0.0280 (7/7, p 0.008) |

| stratum | n subj. | n vertices (mean) | fine map itself | near part only | near + far | near, renormalised | near + shuffled far | D1 full − near | D2 shuffled − full | D3 renorm − full |
|---|---|---|---|---|---|---|---|---|---|---|
| visual, > 20% | 7 | 2178 | 0.488 | 0.392 | 0.348 | 0.389 | 0.468 | −0.0435 (7/7, p 0.008) | +0.1198 (7/7, p 0.008) | +0.0406 (7/7, p 0.008) |
| all covered, > 10% | 7 | 15581 | 0.604 | 0.508 | 0.475 | 0.499 | 0.558 | −0.0334 (7/7, p 0.008) | +0.0829 (7/7, p 0.008) | +0.0248 (7/7, p 0.008) |
| pericalcarine | 7 | 3681 | 0.349 | 0.250 | 0.239 | 0.245 | 0.263 | −0.0110 (5/7, p 0.039) | +0.0244 (7/7, p 0.008) | +0.0059 (7/7, p 0.008) |

#### S15. Reviewer-requested tests, second set: kernel-width scan, re-tuned half-run settings, correlated noise, the far criterion

Four analyses asked for by the reviewer round, run under one plan hashed first (S11.8) and one addendum (S11.9, S15.7). Two descriptive additions were made after the plan: the calibration of the graph geodesic against Workbench's exact geodesic, with gross far fractions and pair statistics (S15.5), and the leakage maps with the kernels narrowed to the scan's optimum (S15.2). Tables are copied from the result files.

##### S15.1 Components of the native PSF model, and the kernel-width scan (5 tuning subjects)

Parameters: voxel 1.6 mm; effective echo spacing 0.32 ms and FOV 208.0 mm along PE (HCP 7T protocol; iPAT 2 folded into the effective spacing); T2* 28.0 ms (grey matter at 7T, literature value); in-plane Tukey taper 0.3, slice edge 0.15 voxel and interpolation blur σ 0.4 mm are modelling assumptions. Partial Fourier (7/8 in the HCP 7T protocol) is not in the native model.

| axis | components included | FWHM (mm) | voxels |
|---|---|---|---|
| x (frequency encoding) | sinc of the voxel (box in k-space) | 1.94 | 1.21 |
| x (frequency encoding) | + Tukey taper 0.3 (modelling assumption; no raw filter in the protocol) | 2.26 | 1.41 |
| x (frequency encoding) | + interpolation blur, σ = 0.4 mm (one-step resampling of the HCP pipeline; assumption) | 2.34 | 1.46 |
| y (phase encoding) | sinc of the voxel (box in k-space) | 1.94 | 1.21 |
| y (phase encoding) | + Tukey taper 0.3 (modelling assumption; no raw filter in the protocol) | 2.26 | 1.41 |
| y (phase encoding) | + T2* decay over the echo train (effective echo spacing 0.32 ms, FOV 208.0 mm, T2* 28.0 ms) | 2.50 | 1.56 |
| y (phase encoding) | + interpolation blur, σ = 0.4 mm (one-step resampling of the HCP pipeline; assumption) | 2.60 | 1.62 |
| z (slice) | slice profile: box of the slice thickness with erf edge 0.15 voxel (assumption), band-limited to 1.2× Nyquist | 2.12 | 1.32 |
| z (slice) | + interpolation blur, σ = 0.4 mm | 2.24 | 1.40 |

##### Emulated coarse kernels (native model times the Tukey-0.5 window at the coarse Nyquist)

| grid | axis | FWHM (mm) | voxels |
|---|---|---|---|
| 2.4 mm | x | 3.84 | 1.60 |
| 2.4 mm | y | 4.08 | 1.70 |
| 2.4 mm | z | 3.94 | 1.64 |
| 3.2 mm | x | 5.12 | 1.60 |
| 3.2 mm | y | 5.34 | 1.67 |
| 3.2 mm | z | 5.18 | 1.62 |

Reading: a bare sinc has a main lobe of 1.21 voxels; the taper, the T2* decay along PE and the interpolation blur bring the native kernel to 1.46 (x), 1.62 (y) and 1.40 (z) voxels. The emulated coarse kernels inherit this width (1.6–1.7 voxels).

α stretches the inversion's coarse-grid kernels (data and rendering unchanged); "all" = all three axes, "y" = phase-encoding axis only, "z" = slice axis only. "better than α = 1": subjects in which the stretched kernel gives the lower residual.

### 2.4 mm

| α | axes | all coherent | better than α = 1 | leak > 10% | better | pericalcarine | better |
|---|---|---|---|---|---|---|---|
| 1 | – | 0.3126 | – | 0.3725 | – | 0.2156 | – |
| 0.7 | all | 0.3100 | 4/5 | 0.3715 | 4/5 | 0.2135 | 3/5 |
| 1.3 | all | 0.3227 | 0/5 | 0.3810 | 1/5 | 0.2246 | 0/5 |
| 0.85 | all | 0.3103 | 5/5 | 0.3704 | 4/5 | 0.2134 | 4/5 |
| 1.15 | all | 0.3167 | 0/5 | 0.3761 | 1/5 | 0.2194 | 1/5 |
| 0.85 | y | 0.3112 | 5/5 | 0.3710 | 4/5 | 0.2147 | 5/5 |
| 0.85 | z | 0.3123 | 4/5 | 0.3720 | 4/5 | 0.2154 | 4/5 |
| 1.15 | y | 0.3143 | 0/5 | 0.3738 | 0/5 | 0.2167 | 1/5 |
| 1.15 | z | 0.3135 | 0/5 | 0.3738 | 1/5 | 0.2166 | 1/5 |

### 3.2 mm

| α | axes | all coherent | better than α = 1 | leak > 10% | better | pericalcarine | better |
|---|---|---|---|---|---|---|---|
| 1 | – | 0.3214 | – | 0.3279 | – | 0.2237 | – |
| 0.7 | all | 0.3178 | 4/5 | 0.3229 | 3/5 | 0.2199 | 4/5 |
| 1.3 | all | 0.3379 | 0/5 | 0.3441 | 0/5 | 0.2397 | 0/5 |
| 0.85 | all | 0.3182 | 5/5 | 0.3239 | 5/5 | 0.2206 | 5/5 |
| 1.15 | all | 0.3283 | 0/5 | 0.3351 | 0/5 | 0.2304 | 0/5 |
| 0.85 | y | 0.3196 | 5/5 | 0.3269 | 3/5 | 0.2225 | 4/5 |
| 0.85 | z | 0.3209 | 5/5 | 0.3267 | 5/5 | 0.2233 | 4/5 |
| 1.15 | y | 0.3241 | 0/5 | 0.3293 | 0/5 | 0.2257 | 1/5 |
| 1.15 | z | 0.3230 | 0/5 | 0.3305 | 0/5 | 0.2254 | 2/5 |

#### S15.2 Leakage maps with the kernels narrowed to the scan optimum (20 development subjects)

α scales the inversion's kernels on all three axes (α = 1 is the paper's PSF model; the scan preferred 0.85 and 0.7); emulated coarse kernels, sphere criterion at 15 mm. FWHM = full width at half maximum of the kernel along x / y / z (mm, mean over subjects). Area-weighted means per hemisphere, averaged over hemispheres and subjects (± SD).

### 1.6 mm

| α | kernel FWHM x / y / z (mm) | net far leak (%) | gross far (%) | cortex > 5% (%) | cortex > 10% (%) | pericalcarine (%) |
|---|---|---|---|---|---|---|
| 1 | 2.56 / 2.70 / 2.55 | 2.18 ± 0.35 | 3.56 ± 0.45 | 13.66 ± 2.96 | 4.75 ± 1.63 | 3.88 ± 0.94 |

| α | kernel FWHM x / y / z (mm) | net far leak (%) | gross far (%) | cortex > 5% (%) | cortex > 10% (%) | pericalcarine (%) |
|---|---|---|---|---|---|---|
| 0.85 | 2.17 / 2.29 / 2.16 | 1.71 ± 0.30 | 2.61 ± 0.37 | 10.69 ± 2.73 | 3.28 ± 1.22 | 3.07 ± 0.80 |
| 0.7 | 1.79 / 1.89 / 1.78 | 1.30 ± 0.26 | 1.99 ± 0.32 | 8.05 ± 2.39 | 2.19 ± 0.86 | 2.38 ± 0.68 |

### 2.4 mm

| α | kernel FWHM x / y / z (mm) | net far leak (%) | gross far (%) | cortex > 5% (%) | cortex > 10% (%) | pericalcarine (%) |
|---|---|---|---|---|---|---|
| 1 | 3.83 / 4.08 / 3.96 | 2.56 ± 0.54 | 11.78 ± 0.87 | 17.95 ± 3.01 | 9.76 ± 2.28 | 5.03 ± 1.42 |
| 0.85 | 3.26 / 3.46 / 3.37 | 1.97 ± 0.44 | 8.47 ± 0.74 | 14.09 ± 2.78 | 7.12 ± 1.87 | 4.02 ± 1.21 |
| 0.7 | 2.68 / 2.85 / 2.77 | 1.55 ± 0.36 | 6.18 ± 0.63 | 11.37 ± 2.48 | 5.37 ± 1.51 | 3.18 ± 0.99 |

### 3.2 mm

| α | kernel FWHM x / y / z (mm) | net far leak (%) | gross far (%) | cortex > 5% (%) | cortex > 10% (%) | pericalcarine (%) |
|---|---|---|---|---|---|---|
| 1 | 5.07 / 5.36 / 5.11 | 3.62 ± 0.69 | 21.10 ± 1.23 | 25.85 ± 2.70 | 16.84 ± 2.52 | 6.45 ± 1.82 |
| 0.85 | 4.31 / 4.55 / 4.35 | 2.78 ± 0.60 | 16.30 ± 1.14 | 20.46 ± 2.62 | 12.79 ± 2.20 | 5.42 ± 1.69 |
| 0.7 | 3.55 / 3.75 / 3.58 | 2.14 ± 0.51 | 12.43 ± 1.00 | 16.09 ± 2.37 | 9.81 ± 1.87 | 4.39 ± 1.54 |

#### S15.3 Settings re-tuned on half-run sources (5 tuning subjects), and H3 with the re-tuned settings (15 development subjects)

Cross-run residual against the other run's full-run 1.6 mm map, averaged over the two halves and the subjects; the week-3 rule picks the lowest mean over all coherent vertices (first table) and, secondarily, where leak > 10% at the coarse grid. Full-run choices from week 3 for comparison.

### 2.4 mm

Half-run choice (all coherent): model **model_r24_s0.25**, smoothing **smooth10**, upsampling + smoothing **upsmooth8**; full-run choice (week 3): model_r12_s0.125, smooth10, upsmooth8. Where leak > 10%: model_r48_s0.5, smooth10, upsmooth8.

| method | all coherent | leak > 10% |
|---|---|---|
| model_r24_s0.25 | 0.3258 | 0.3797 |
| model_r48_s0.5 | 0.3261 | 0.3793 |
| model_r12_s0.125 | 0.3286 | 0.3868 |
| model_r12_s0.25 | 0.3295 | 0.3836 |
| model_r48_s0.25 | 0.3309 | 0.3880 |
| upsmooth8 | 0.3310 | 0.3908 |
| model_r24_s0.5 | 0.3317 | 0.3839 |
| upsmooth10 | 0.3323 | 0.3924 |
| model_r24_s0.125 | 0.3327 | 0.3913 |
| upsmooth6 | 0.3328 | 0.3924 |
| upsmooth12 | 0.3356 | 0.3958 |
| model_r6_s0.25 | 0.3394 | 0.3989 |
| upsmooth4 | 0.3396 | 0.3987 |
| upsmooth14 | 0.3401 | 0.4005 |
| model_r6_s0.125 | 0.3416 | 0.4053 |

| method | all coherent | leak > 10% |
|---|---|---|
| smooth10 | 0.3434 | 0.4089 |
| model_r12_s0.5 | 0.3439 | 0.3983 |
| smooth8 | 0.3440 | 0.4122 |
| smooth12 | 0.3452 | 0.4092 |
| upsmooth16 | 0.3453 | 0.4060 |
| model_r12_s0.0625 | 0.3483 | 0.4123 |
| smooth6 | 0.3485 | 0.4214 |
| smooth14 | 0.3485 | 0.4116 |
| model_r48_s0.125 | 0.3507 | 0.4114 |
| upsmooth2 | 0.3528 | 0.4098 |
| smooth16 | 0.3528 | 0.4154 |
| model_r24_s0.0625 | 0.3574 | 0.4205 |
| model_r6_s0.5 | 0.3584 | 0.4179 |
| smooth4 | 0.3595 | 0.4405 |
| upsample | 0.3641 | 0.4197 |
| model_r6_s0.0625 | 0.3726 | 0.4382 |
| smooth2 | 0.3809 | 0.4754 |
| model_r48_s0.0625 | 0.3872 | 0.4496 |
| standard | 0.4165 | 0.5329 |

### 3.2 mm

Half-run choice (all coherent): model **model_r48_s0.5**, smoothing **smooth10**, upsampling + smoothing **upsmooth10**; full-run choice (week 3): model_r24_s0.25, smooth10, upsmooth8. Where leak > 10%: model_r12_s0.25, smooth10, upsmooth8.

| method | all coherent | leak > 10% |
|---|---|---|
| model_r48_s0.5 | 0.3347 | 0.3409 |
| model_r24_s0.25 | 0.3348 | 0.3411 |
| model_r12_s0.25 | 0.3373 | 0.3406 |
| model_r24_s0.5 | 0.3388 | 0.3424 |
| upsmooth10 | 0.3410 | 0.3457 |
| upsmooth8 | 0.3410 | 0.3456 |
| model_r12_s0.125 | 0.3423 | 0.3477 |
| upsmooth12 | 0.3431 | 0.3482 |
| model_r48_s0.25 | 0.3439 | 0.3540 |
| upsmooth6 | 0.3446 | 0.3491 |
| upsmooth14 | 0.3468 | 0.3522 |
| model_r24_s0.125 | 0.3473 | 0.3573 |
| model_r6_s0.25 | 0.3494 | 0.3508 |
| smooth10 | 0.3507 | 0.3607 |
| model_r12_s0.5 | 0.3512 | 0.3538 |
| upsmooth16 | 0.3513 | 0.3571 |
| smooth12 | 0.3520 | 0.3613 |
| smooth8 | 0.3521 | 0.3632 |
| upsmooth4 | 0.3531 | 0.3581 |
| smooth14 | 0.3548 | 0.3638 |
| smooth6 | 0.3574 | 0.3711 |

| method | all coherent | leak > 10% |
|---|---|---|
| smooth16 | 0.3587 | 0.3676 |
| model_r6_s0.125 | 0.3632 | 0.3656 |
| model_r6_s0.5 | 0.3654 | 0.3675 |
| upsmooth2 | 0.3664 | 0.3724 |
| smooth4 | 0.3694 | 0.3880 |
| model_r48_s0.125 | 0.3698 | 0.3831 |
| model_r12_s0.0625 | 0.3726 | 0.3795 |
| upsample | 0.3750 | 0.3820 |
| model_r24_s0.0625 | 0.3795 | 0.3915 |
| smooth2 | 0.3926 | 0.4218 |
| model_r6_s0.0625 | 0.4047 | 0.4089 |
| model_r48_s0.0625 | 0.4110 | 0.4251 |
| standard | 0.4465 | 0.4787 |

Full runs: model `model_r12_s0.125|model_r24_s0.25`, competitor `upsmooth8|upsmooth8` (week-3 choices). Half runs: model `model_r24_s0.25|model_r48_s0.5`, competitor `upsmooth8|upsmooth10` (chosen on the tuning subjects' half runs, E1). Δ_clean = gain in the 0–2% far-leak bin; D_far = Δ(highest bin) − Δ(0–2%); half = mean of the two halves.

| grid | quantity | mean | in direction | bootstrap 95% | p (one-sided) | p (two-sided) |
|---|---|---|---|---|---|---|
| 2.4 mm | H3a′: Δ_clean(half, re-tuned) − Δ_clean(full) > 0 | +0.0026 | 14/15 | [+0.0011, +0.0037] | 0.003 | 0.007 |
| 2.4 mm | H3a (fixed settings, for reference): Δ_clean(half, full-run settings) − Δ_clean(full) > 0 | +0.0041 | 14/15 | [+0.0009, +0.0064] | 0.004 | 0.008 |
| 2.4 mm | H3b′: D_far(half, re-tuned) − D_far(full) (interval) | −0.0000 | 8/15 | [−0.0028, +0.0025] | – | 0.978 |
| 2.4 mm | H3c′: Δ_clean(half, re-tuned) < 0 | −0.0060 | 11/15 | [−0.0101, −0.0022] | 0.006 | 0.012 |
| 2.4 mm | Δ_clean(full) (reference) | −0.0086 | 14/15 | [−0.0127, −0.0050] | < 0.001 | < 0.001 |
| 2.4 mm | half-run model gain, all coherent (re-tuned) | −0.0052 | 12/15 | [−0.0088, −0.0018] | 0.008 | 0.015 |
| 3.2 mm | H3a′: Δ_clean(half, re-tuned) − Δ_clean(full) > 0 | +0.0022 | 13/15 | [+0.0007, +0.0036] | 0.006 | 0.012 |
| 3.2 mm | H3a (fixed settings, for reference): Δ_clean(half, full-run settings) − Δ_clean(full) > 0 | +0.0024 | 13/15 | [−0.0001, +0.0045] | 0.011 | 0.022 |
| 3.2 mm | H3b′: D_far(half, re-tuned) − D_far(full) (interval) | +0.0028 | 6/15 | [+0.0007, +0.0049] | – | 0.035 |
| 3.2 mm | H3c′: Δ_clean(half, re-tuned) < 0 | −0.0143 | 14/15 | [−0.0185, −0.0101] | < 0.001 | < 0.001 |
| 3.2 mm | Δ_clean(full) (reference) | −0.0165 | 15/15 | [−0.0212, −0.0120] | < 0.001 | < 0.001 |
| 3.2 mm | half-run model gain, all coherent (re-tuned) | −0.0106 | 14/15 | [−0.0141, −0.0070] | < 0.001 | < 0.001 |

##### S15.4 The far criterion at other thresholds (20 development subjects)

Sphere proxy at 10 / 15 / 20 mm along the sheet (15 mm is the paper's convention) and "other hemisphere only"; emulated kernels; area-weighted means per hemisphere, averaged over hemispheres and subjects (± SD).

## 1.6 mm

| far criterion | mean far leak (%) | cortex > 5% (%) | cortex > 10% (%) | pericalcarine (%) |
|---|---|---|---|---|
| > 10 mm | 3.53 ± 0.48 | 24.39 ± 4.04 | 8.77 ± 2.52 | 5.84 ± 1.16 |
| > 15 mm (used) | 2.18 ± 0.35 | 13.66 ± 2.96 | 4.75 ± 1.63 | 3.88 ± 0.94 |
| > 20 mm | 1.42 ± 0.25 | 8.27 ± 2.08 | 2.78 ± 1.04 | 2.55 ± 0.80 |
| other hemisphere only | 0.21 ± 0.04 | 0.84 ± 0.27 | 0.22 ± 0.08 | 0.30 ± 0.15 |

## 2.4 mm

| far criterion | mean far leak (%) | cortex > 5% (%) | cortex > 10% (%) | pericalcarine (%) |
|---|---|---|---|---|
| > 10 mm | 4.38 ± 0.73 | 30.03 ± 3.84 | 16.90 ± 3.21 | 7.56 ± 1.65 |
| > 15 mm (used) | 2.56 ± 0.54 | 17.95 ± 3.01 | 9.76 ± 2.28 | 5.03 ± 1.42 |
| > 20 mm | 1.60 ± 0.39 | 11.42 ± 2.25 | 6.04 ± 1.59 | 3.34 ± 1.18 |
| other hemisphere only | 0.23 ± 0.06 | 1.71 ± 0.39 | 0.85 ± 0.24 | 0.31 ± 0.24 |

## 3.2 mm

| far criterion | mean far leak (%) | cortex > 5% (%) | cortex > 10% (%) | pericalcarine (%) |
|---|---|---|---|---|
| > 10 mm | 5.09 ± 0.94 | 36.40 ± 3.25 | 25.38 ± 3.16 | 9.52 ± 1.95 |
| > 15 mm (used) | 3.62 ± 0.69 | 25.85 ± 2.70 | 16.84 ± 2.52 | 6.45 ± 1.82 |
| > 20 mm | 2.26 ± 0.51 | 17.24 ± 2.13 | 10.85 ± 1.87 | 4.14 ± 1.43 |
| other hemisphere only | 0.35 ± 0.09 | 2.78 ± 0.43 | 1.79 ± 0.35 | 0.41 ± 0.52 |

##### S15.5 True geodesic against the sphere proxy (5 tuning subjects)

Geodesic = Dijkstra on the edge graph of the native midthickness mesh (every undirected edge once), calibrated against Workbench's exact geodesic (`wb_command -surface-geodesic-distance`, 60 random ROI vertices per hemisphere, targets at 5–25 mm): graph / exact median 1.0350 (subjects 1.0347–1.0354; IQR within subjects about 1.02–1.05), so the far label is also evaluated at the calibrated graph threshold 15.53 mm ("calibrated"). Per hemisphere, averaged over hemispheres and subjects. Net = signed far weight over the row sum; gross = |A| far share. "label change": share of cortex (area-weighted) whose > 5% / > 10% far-leak label differs between the geodesic (uncalibrated) and sphere criteria; "mass flip": share of the operator's mass (|A|) whose far/near label differs.

| grid | net far leak: geodesic / calibrated / sphere (%) | gross far: geodesic / sphere (%) | > 5%: geodesic / calibrated / sphere (%) | > 10%: geodesic / calibrated / sphere (%) | r | median: geodesic / sphere (%) | label change > 5% / > 10% (%) | mass flip (%) |
|---|---|---|---|---|---|---|---|---|
| 1.6 mm | 2.27 / 2.17 / 2.30 | 3.32 / 3.37 | 14.1 / 13.4 / 15.0 | 5.1 / 4.8 / 5.5 | 0.891 | 0.70 / 0.64 | 4.3 / 1.6 | 0.9 |
| 2.4 mm | 2.58 / 2.48 / 2.83 | 9.78 / 9.70 | 18.3 / 17.5 / 19.7 | 10.1 / 9.7 / 11.2 | 0.858 | 0.60 / 0.73 | 6.1 / 3.4 | 2.7 |
| 3.2 mm | 3.58 / 3.38 / 4.03 | 17.33 / 16.84 | 26.1 / 25.0 / 27.7 | 17.2 / 16.4 / 18.6 | 0.898 | 0.82 / 1.04 | 9.2 / 5.7 | 4.3 |

Pairs (vertex, unknown) of A in the same hemisphere, pooled over hemispheres and subjects, by sphere-proxy distance: mean geodesic / sphere ratio and the share of pairs whose far label differs (geodesic > 15 mm but sphere ≤ 15 mm, and the reverse). Pairs beyond the geodesic limit are not in the ratio.

| grid | sphere distance (mm) | pairs (millions) | geodesic / sphere | geodesic far, sphere near (%) | geodesic near, sphere far (%) |
|---|---|---|---|---|---|
| 1.6 mm | 5-10 | 329.4 | 1.164 | 1.7 | 0.0 |
| 1.6 mm | 10-15 | 251.6 | 1.046 | 25.1 | 0.0 |
| 1.6 mm | 15-20 | 139.2 | 0.983 | 0.0 | 35.3 |
| 1.6 mm | 20-25 | 63.0 | 0.885 | 0.0 | 12.8 |
| 2.4 mm | 5-10 | 415.7 | 1.231 | 4.7 | 0.0 |
| 2.4 mm | 10-15 | 644.8 | 1.173 | 46.3 | 0.0 |
| 2.4 mm | 15-20 | 643.8 | 1.069 | 0.0 | 18.2 |
| 2.4 mm | 20-25 | 382.1 | 0.919 | 0.0 | 6.2 |
| 3.2 mm | 5-10 | 408.5 | 1.234 | 5.0 | 0.0 |
| 3.2 mm | 10-15 | 664.7 | 1.189 | 49.0 | 0.0 |
| 3.2 mm | 15-20 | 744.4 | 1.093 | 0.0 | 15.4 |
| 3.2 mm | 20-25 | 480.7 | 0.937 | 0.0 | 4.8 |

##### S15.6 Correlated-noise likelihood on HCP (15 development subjects)

Cross-run residual (mean over subjects); Δ = model − competitor (upsampling + 8 mm smoothing); D_far = Δ(highest bin) − Δ(0–2%).

#### 2.4 mm

| stratum | white | correlated | competitor | corr – white (mean, subjects corr better, two-sided p) |
|---|---|---|---|---|
| all coherent | 0.3131 | 0.3670 | 0.3212 | +0.0539, 0/15, p < 0.001 |
| pericalcarine | 0.2398 | 0.3047 | 0.2482 | +0.0649, 0/15, p < 0.001 |
| clean, leak > 10% | 0.3428 | 0.4106 | 0.3671 | +0.0678, 0/15, p < 0.001 |
| 0–2% | 0.3094 | 0.3574 | 0.3180 | +0.0480, 0/15, p < 0.001 |
| 2–5% | 0.2855 | 0.3313 | 0.2958 | +0.0458, 0/15, p < 0.001 |
| 5–10% | 0.3005 | 0.3474 | 0.3148 | +0.0469, 0/15, p < 0.001 |
| 10–20% | 0.3389 | 0.4040 | 0.3582 | +0.0651, 0/15, p < 0.001 |
| > 20% | 0.3260 | 0.4024 | 0.3611 | +0.0764, 0/14, p < 0.001 |

D_far: white −0.0244, correlated +0.0038; corr − white +0.0283 [+0.0112, +0.0492], p 0.004

#### 3.2 mm

| stratum | white | correlated | competitor | corr – white (mean, subjects corr better, two-sided p) |
|---|---|---|---|---|
| all coherent | 0.3248 | 0.3503 | 0.3385 | +0.0255, 0/15, p < 0.001 |
| pericalcarine | 0.2505 | 0.2745 | 0.2628 | +0.0240, 5/15, p 0.018 |
| clean, leak > 10% | 0.3439 | 0.3652 | 0.3712 | +0.0212, 2/15, p < 0.001 |
| 0–2% | 0.3256 | 0.3515 | 0.3421 | +0.0260, 0/15, p < 0.001 |
| 2–5% | 0.3022 | 0.3273 | 0.3160 | +0.0251, 0/15, p < 0.001 |
| 5–10% | 0.3113 | 0.3332 | 0.3299 | +0.0219, 0/15, p < 0.001 |

| stratum | white | correlated | competitor | corr − white (mean, subjects corr better, two-sided p) |
|---|---|---|---|---|
| 10–20% | 0.3355 | 0.3572 | 0.3539 | +0.0217, 3/15, p < 0.001 |
| > 20% | 0.3514 | 0.3730 | 0.3908 | +0.0215, 1/15, p < 0.001 |

D_far: white −0.0229, correlated −0.0273; corr − white −0.0044 [−0.0114, +0.0027], p 0.252

Per subject and run (2 runs), the rendered 1.6 mm prediction of each fit over the analysis vertices: amplitude ratio |corr| / |white| (root mean square), magnitude of the complex correlation, and both after removing each hemisphere's mean. `src/16_sensitivity_analyses/corr_diag.py`; mean [min, max] over subjects × runs. The noise spectrum (radial power relative to white, from the fit logs of the 15 subjects) is given for context.

| grid | amplitude ratio | correlation | amplitude ratio, hemisphere means removed | correlation, means removed | noise power at wavelengths > 20 mm (× white) | lowest radial bin (× white) |
|---|---|---|---|---|---|---|
| 2.4 mm | 0.73 [0.63, 0.81] | 0.95 [0.92, 0.98] | 0.73 [0.64, 0.81] | 0.95 [0.91, 0.97] | 44–120 | 153–546 |
| 3.2 mm | 0.84 [0.75, 0.91] | 0.97 [0.94, 0.98] | 0.84 [0.75, 0.91] | 0.97 [0.94, 0.98] | 41–133 | 171–748 |

Reading: the correlated fit reproduces the white fit's pattern (r 0.92–0.98) at 63–81% (2.4 mm) and 75–91% (3.2 mm) of its amplitude, and the shrinkage is the same with the hemisphere means removed, so it is not confined to the coarsest scale. The estimated spectrum puts tens to hundreds of times the white power at the large spatial scales over which the retinotopic map varies, so the likelihood discounts them.

##### S15.7 The correlated-noise likelihood with its own prior settings (addendum, S11.9)

Plan: `PLAN_F2_ADDENDUM.md`. Correlated fits on r ∈ {6, 12, 24, 48} mm × s ∈ {1/32 … 1}; the week-3 rule (lowest mean cross-run residual over all coherent vertices of the tuning subjects) picks the correlated setting per grid; the white likelihood keeps its week-3 choice. Negative differences favour the correlated likelihood.

#### 2.4 mm

Tuning subjects, mean residual over all coherent vertices (white likelihood at its choice `model_r12_s0.125`: 0.3126); choice **model_r24_s0.25corr** (0.3406).

| r \ s | 0.03125 | 0.0625 | 0.125 | 0.25 | 0.5 | 1 |
|---|---|---|---|---|---|---|
| 6 mm | 0.4993 | 0.4470 | 0.3964 | 0.3769 | 0.3834 | 0.4035 |
| 12 mm | 0.4219 | 0.3724 | 0.3490 | 0.3505 | 0.3643 | 0.3838 |
| 24 mm | 0.4102 | 0.3662 | 0.3431 | **0.3406** | 0.3503 | 0.3660 |
| 48 mm | 0.4416 | 0.3959 | 0.3590 | 0.3414 | 0.3412 | 0.3513 |

Development subjects (n = 15): residual by stratum, mean over subjects; differences paired per subject.

| stratum | white `model_r12_s0.125` | correlated, re-tuned `model_r24_s0.25corr` | correlated at the white settings (F) | competitor `upsmooth8` | re-tuned − white (mean [95%], subjects corr better, p) |
|---|---|---|---|---|---|
| all coherent | 0.3131 | 0.3596 | 0.3670 | 0.3212 | +0.0465 [+0.0361, +0.0570], 0/15, p < 0.001 |
| pericalcarine | 0.2398 | 0.2986 | 0.3047 | 0.2482 | +0.0588 [+0.0347, +0.0851], 1/15, p < 0.001 |
| clean, leak > 10% | 0.3428 | 0.4011 | 0.4106 | 0.3671 | +0.0583 [+0.0427, +0.0743], 0/15, p < 0.001 |
| 0–2% | 0.3094 | 0.3510 | 0.3574 | 0.3180 | +0.0416 [+0.0324, +0.0514], 0/15, p < 0.001 |
| 2–5% | 0.2855 | 0.3241 | 0.3313 | 0.2958 | +0.0386 [+0.0276, +0.0499], 0/15, p < 0.001 |
| 5–10% | 0.3005 | 0.3397 | 0.3474 | 0.3148 | +0.0392 [+0.0265, +0.0521], 1/15, p < 0.001 |
| 10–20% | 0.3389 | 0.3945 | 0.4040 | 0.3582 | +0.0556 [+0.0408, +0.0710], 0/15, p < 0.001 |
| > 20% | 0.3260 | 0.3932 | 0.4024 | 0.3611 | +0.0672 [+0.0436, +0.0928], 0/14, p < 0.001 |

D_far (model − competitor, highest bin − 0–2%): white −0.0244, correlated re-tuned +0.0015; difference +0.0259 [+0.0080, +0.0476], p 0.015. Oracle bound (best correlated setting chosen on these development subjects themselves): `model_r24_s0.25corr`, 0.3596 against white 0.3131 (+0.0465). F reproduced inside F2 (`model_r12_s0.125corr`, same code, warm-started differently): largest per-subject difference from the F run 1.4e-06 over 15 subjects.

### 3.2 mm

Tuning subjects, mean residual over all coherent vertices (white likelihood at its choice `model_r24_s0.25`: 0.3214); choice **model_r48_s0.5corr** (0.3388).

| r \ s | 0.03125 | 0.0625 | 0.125 | 0.25 | 0.5 | 1 |
|---|---|---|---|---|---|---|
| 6 mm | 0.5475 | 0.4951 | 0.4292 | 0.3880 | 0.3871 | 0.4171 |
| 12 mm | 0.4583 | 0.4017 | 0.3609 | 0.3487 | 0.3589 | 0.3864 |
| 24 mm | 0.4345 | 0.3871 | 0.3535 | 0.3400 | 0.3433 | 0.3601 |
| 48 mm | 0.4605 | 0.4171 | 0.3761 | 0.3483 | **0.3388** | 0.3438 |

Development subjects (n = 15): residual by stratum, mean over subjects; differences paired per subject.

| stratum | white `model_r24_s0.25` | correlated, re-tuned `model_r48_s0.5corr` | correlated at the white settings (F) | competitor `upsmooth8` | re-tuned − white (mean [95%], subjects corr better, p) |
|---|---|---|---|---|---|
| all coherent | 0.3248 | 0.3500 | 0.3503 | 0.3385 | +0.0252 [+0.0188, +0.0330], 0/15, p < 0.001 |

| stratum | white `model_r24_s0.25` | correlated, re-tuned `model_r48_s0.5corr` | correlated at the white settings (F) | competitor `upsmooth8` | re-tuned − white (mean [95%], subjects corr better, p) |
|---|---|---|---|---|---|
| pericalcarine | 0.2505 | 0.2747 | 0.2745 | 0.2628 | +0.0242 [+0.0076, +0.0456], 5/15, p 0.018 |
| clean, leak > 10% | 0.3439 | 0.3650 | 0.3652 | 0.3712 | +0.0210 [+0.0121, +0.0315], 2/15, p < 0.001 |
| 0–2% | 0.3256 | 0.3511 | 0.3515 | 0.3421 | +0.0255 [+0.0198, +0.0327], 0/15, p < 0.001 |
| 2–5% | 0.3022 | 0.3268 | 0.3273 | 0.3160 | +0.0246 [+0.0191, +0.0316], 0/15, p < 0.001 |
| 5–10% | 0.3113 | 0.3328 | 0.3332 | 0.3299 | +0.0215 [+0.0146, +0.0300], 0/15, p < 0.001 |
| 10–20% | 0.3355 | 0.3568 | 0.3572 | 0.3539 | +0.0213 [+0.0114, +0.0329], 3/15, p < 0.001 |
| > 20% | 0.3514 | 0.3730 | 0.3730 | 0.3908 | +0.0215 [+0.0121, +0.0319], 1/15, p < 0.001 |

D_far (model − competitor, highest bin − 0–2%): white −0.0229, correlated re-tuned −0.0269; difference −0.0040 [−0.0110, +0.0032], p 0.277. Oracle bound (best correlated setting chosen on these development subjects themselves): `model_r48_s0.5corr`, 0.3500 against white 0.3248 (+0.0252). F reproduced inside F2 (`model_r24_s0.25corr`, same code, warm-started differently): largest per-subject difference from the F run 2.3e-06 over 15 subjects.

#### S16. External replication in the NYU Retinotopy Dataset (3T, 2.0 mm, sweeping bars)

Plans: S11.10 (with its deviations), S11.11 and S11.12; methods: S17, Section 3.10; summary: main text Sections 4.2, 4.4 and 4.5 and Fig. 8. Tables copied from the result files of `src/17_nyu_3t_replication/nyu_summary.py`.

##### S16.1 Settings chosen on the 8 tuning subjects

Mean cross-half residual over responsive vertices (lower is better); the competitor of the tests is the better of the two chosen smoothing competitors.

#### 3.0 mm

Choice: model **model_r12_s2** (0.3064), smoothing **smooth2** (0.3214), upsampling + smoothing **upsmooth2** (0.3030); competitor of the tests **upsmooth2**. Standard mapping 0.3487, upsampling 0.2995, native source half (no coarsening) 0.1586.

| model setting | residual |
|---|---|
| model_r12_s2 | 0.3064 |
| model_r24_s4 | 0.3069 |
| model_r6_s1 | 0.3083 |
| model_r48_s4 | 0.3124 |

| model setting | residual |
|---|---|
| model_r24_s2 | 0.3129 |
| model_r6_s2 | 0.3140 |
| model_r3_s1 | 0.3143 |
| model_r12_s1 | 0.3153 |

### 4.0 mm

Choice: model **model_r6_s1** (0.3615), smoothing **smooth4** (0.3905), upsampling + smoothing **upsmooth2** (0.3687); competitor of the tests **upsmooth2**. Standard mapping 0.4218, upsampling 0.3704, native source half (no coarsening) 0.1626.

| model setting | residual |
|---|---|
| model_r6_s1 | 0.3615 |
| model_r24_s2 | 0.3627 |
| model_r48_s4 | 0.3628 |
| model_r12_s1 | 0.3635 |
| model_r12_s2 | 0.3635 |
| model_r24_s4 | 0.3667 |
| model_r6_s2 | 0.3701 |
| model_r3_s2 | 0.3707 |

#### S16.2 Tests on the 35 included test subjects, descriptive comparisons, and the leak of the 3T acquisition

Δ = residual(model) − residual(competitor), negative favours the model; one-sided Wilcoxon (predicted direction: less), bootstrap 95% interval of the mean (10,000 resamples, seed 0); Holm over the six primary tests (N1–N3 × two grids).

Not included (preprocessing exclusions or missing): sub-wlsubj055.

| grid | test | competitor | model | mean | subjects in direction | bootstrap 95% | p (one-sided) | p (Holm) |
|---|---|---|---|---|---|---|---|---|
| 3.0 mm | N1: Δ responsive | upsmooth2 | model_r12_s2 | +0.0047 | 13/35 | [+0.0012, +0.0084] | 0.990 | 1.000 |
| 3.0 mm | N2: D_far | upsmooth2 | model_r12_s2 | −0.0024 | 19/35 | [−0.0109, +0.0060] | 0.311 | 0.932 |
| 3.0 mm | N3: D_CWL-M | upsmooth2 | model_r12_s2 | −0.0151 | 25/35 | [−0.0258, −0.0045] | 0.004 | 0.018 |
| 3.0 mm | descriptive: Δ responsive vs plain upsampling | upsmooth2 | model_r12_s2 | +0.0085 | 8/35 | [+0.0046, +0.0127] | 1.000 | – |
| 3.0 mm | secondary: Δ clean_leak10 | upsmooth2 | model_r12_s2 | +0.0008 | 15/35 | [−0.0049, +0.0063] | 0.739 | 1.000 |
| 3.0 mm | secondary: Δ calc | upsmooth2 | model_r12_s2 | +0.0125 | 7/35 | [+0.0065, +0.0183] | 1.000 | 1.000 |
| 4.0 mm | N1: Δ responsive | upsmooth2 | model_r6_s1 | −0.0073 | 27/35 | [−0.0109, −0.0039] | < 0.001 | < 0.001 |
| 4.0 mm | N2: D_far | upsmooth2 | model_r6_s1 | +0.0030 | 18/35 | [−0.0043, +0.0107] | 0.701 | 1.000 |
| 4.0 mm | N3: D_CWL-M | upsmooth2 | model_r6_s1 | −0.0113 | 25/35 | [−0.0191, −0.0034] | 0.007 | 0.029 |
| 4.0 mm | descriptive: Δ responsive vs plain upsampling | upsmooth2 | model_r6_s1 | −0.0080 | 27/35 | [−0.0118, −0.0043] | < 0.001 | – |

| grid | test | competitor | model | mean | subjects in direction | bootstrap 95% | p (one-sided) | p (Holm) |
|---|---|---|---|---|---|---|---|---|
| 4.0 mm | secondary: Δ clean_leak10 | upsmooth2 | model_r6_s1 | −0.0135 | 28/35 | [−0.0196, −0.0073] | < 0.001 | < 0.001 |
| 4.0 mm | secondary: Δ calc | upsmooth2 | model_r6_s1 | +0.0092 | 13/35 | [+0.0012, +0.0174] | 0.981 | 1.000 |

##### Descriptive

| grid | polar-angle error, model / competitor (degrees) | difference (mean, subjects model better) |
|---|---|---|
| 3.0 mm | 38.38 / 36.66 | +1.72, 1/35 |
| 4.0 mm | 39.31 / 38.60 | +0.70, 3/35 |

Gain (model − competitor) by bin of the dose variable at the coarse grid (responsive vertices with < 2% far leak at 2.0 mm), mean (subjects with the bin):

| grid | dose | 0–2% | 2–5% | 5–10% | 10–20% | > 20% |
|---|---|---|---|---|---|---|
| 3.0 mm | far leak | +0.0051 (35) | +0.0053 (35) | −0.0007 (35) | −0.0009 (35) | +0.0026 (35) |
| 3.0 mm | CWL-M | +0.0101 (35) | +0.0033 (35) | +0.0006 (35) | −0.0024 (35) | −0.0050 (35) |
| 4.0 mm | far leak | −0.0145 (35) | −0.0125 (35) | −0.0114 (35) | −0.0159 (35) | −0.0115 (35) |
| 4.0 mm | CWL-M | −0.0065 (35) | −0.0153 (35) | −0.0136 (35) | −0.0150 (35) | −0.0179 (35) |

##### Far leak of standard mapping for the 3T 2.0 mm acquisition and its emulations (43 subjects; area-weighted, mean ± SD)

| grid | mean far leak (%) | cortex > 5% (%) | cortex > 10% (%) | other hemisphere (%) |
|---|---|---|---|---|
| 2.0 mm | 3.21 ± 0.45 | 18.80 ± 2.91 | 9.67 ± 2.33 | 0.29 ± 0.09 |
| 3.0 mm | 4.24 ± 0.67 | 27.35 ± 2.56 | 18.12 ± 2.42 | 0.38 ± 0.15 |
| 4.0 mm | 5.00 ± 0.67 | 32.56 ± 2.02 | 24.44 ± 2.07 | 0.58 ± 0.21 |

#### S16.3 A coarse-data-only rule: the model where CWL-P is high, the competitor elsewhere (S11.11 A)

Δ = residual(first) − residual(second) over responsive vertices, negative favours the first; one-sided Wilcoxon (less), bootstrap 95% interval of the mean (10,000 resamples, seed 0); Holm over the four pre-specified tests (G1, G2 × two grids).

Reproducibility of the published week-17 scores (model, competitor, standard mapping, upsampling; test subjects): 3.0 mm max |difference| 0.00e+00 over 140 values; 4.0 mm max |difference| 0.00e+00 over 140 values.

##### Choice of τ (tuning subjects, mean residual over responsive vertices)

| grid | τ = 0 | τ = 0.02 | τ = 0.05 | τ = 0.1 | τ = 0.2 | τ = ∞ (competitor) | model alone |
|---|---|---|---|---|---|---|---|
| 3.0 mm | 0.3064 (100%) | 0.3054 (46%) | 0.3038 (21%) | 0.3034 (9%) | **0.3029** (3%) | 0.3030 (0%) | 0.3064 |
| 4.0 mm | **0.3615** (100%) | 0.3618 (68%) | 0.3641 (37%) | 0.3657 (17%) | 0.3673 (6%) | 0.3687 (0%) | 0.3615 |

(in brackets: share of responsive vertices sent to the model; bold: the chosen τ)

##### Tests (test subjects)

| grid | test | τ | mean | subjects in direction | bootstrap 95% | p (one-sided) | p (Holm, 4) |
|---|---|---|---|---|---|---|---|
| 3.0 mm | G1: gated − competitor | 0.2 | +0.0008 | 9/35 | [+0.0003, +0.0014] | 0.999 | 1.000 |
| 3.0 mm | G2: gated − model | 0.2 | −0.0039 | 22/35 | [−0.0074, −0.0005] | 0.030 | 0.090 |
| 4.0 mm | G1: gated − competitor | 0 | −0.0073 | 27/35 | [−0.0109, −0.0039] | < 0.001 | < 0.001 |
| 4.0 mm | G2: gated − model | 0 | +0.0000 | 0/35 | [+0.0000, +0.0000] | 1.000 | 1.000 |

##### Descriptive (test subjects)

| grid | share sent to the model | D_CWL-P (model − competitor) | D_CWL-M (for comparison) | polar-angle error: gated / model / competitor (°) | gated − model at every τ (mean) |
|---|---|---|---|---|---|
| 3.0 mm | 3% | +0.0001 (21/35, p 0.376) | −0.0151 (25/35) | 36.66 / 38.38 / 36.66 | 0: +0.0000, 0.02: −0.0022, 0.05: −0.0029, 0.1: −0.0034, 0.2: −0.0039, inf: −0.0047 |
| 4.0 mm | 100% | +0.0026 (19/35, p 0.593) | −0.0113 (25/35) | 39.31 / 39.31 / 38.60 | 0: +0.0000, 0.02: +0.0004, 0.05: +0.0028, 0.1: +0.0059, 0.2: +0.0073, inf: +0.0073 |

All τ on the test subjects (descriptive; mean Δ against the competitor, subjects better, share sent to the model):

| grid | τ = 0 | τ = 0.02 | τ = 0.05 | τ = 0.1 | τ = 0.2 | τ = inf |
|---|---|---|---|---|---|---|
| 3.0 mm | +0.0047 (13/35; 100%) | +0.0025 (13/35; 48%) | +0.0018 (12/35; 22%) | +0.0013 (12/35; 9%) | +0.0008 (9/35; 3%) | +0.0000 (0/35; 0%) |
| 4.0 mm | −0.0073 (27/35; 100%) | −0.0069 (28/35; 69%) | −0.0045 (23/35; 38%) | −0.0014 (21/35; 18%) | −0.0000 (16/35; 6%) | +0.0000 (0/35; 0%) |

Gain (model − competitor) by CWL-P bin (responsive vertices with < 2% far leak at 2.0 mm), mean (subjects with the bin):

| grid | 0–2% | 2–5% | 5–10% | 10–20% | > 20% |
|---|---|---|---|---|---|
| 3.0 mm | +0.0050 (35) | +0.0015 (35) | +0.0025 (35) | +0.0015 (35) | +0.0052 (35) |
| 4.0 mm | −0.0108 (35) | −0.0140 (35) | −0.0159 (35) | −0.0121 (35) | −0.0083 (35) |

##### S16.4 Registration, field inhomogeneity and the pericalcarine shortfall (S11.11 B, descriptive)

Predictors: bbr_cost median 0.386 (range 0.296–0.527); p95_brain_hz median 44.5 (range 29.9–56); median_responsive_hz median 4.58 (range 2.56–12.9); median_calc_responsive_hz median 4.12 (range 1.48–18.8).

| grid | outcome (model − competitor) | mean | BBR cost: Spearman ρ (p) | field p95 brain: ρ (p) | field at responsive pericalcarine: ρ (p) | mean, BBR cost ≤ median / > median (Mann–Whitney p) |
|---|---|---|---|---|---|---|
| 3.0 mm | Δ responsive | +0.0047 | +0.05 (0.75) | −0.33 (0.05) | +0.02 (0.91) | +0.0043 (n 18) / +0.0052 (n 17) (0.70) |
| 3.0 mm | Δ pericalcarine | +0.0125 | +0.10 (0.55) | −0.16 (0.37) | +0.09 (0.62) | +0.0103 (n 18) / +0.0149 (n 17) (0.50) |
| 4.0 mm | Δ responsive | −0.0073 | −0.04 (0.83) | −0.37 (0.03) | −0.16 (0.36) | −0.0073 (n 18) / −0.0073 (n 17) (0.99) |
| 4.0 mm | Δ pericalcarine | +0.0092 | +0.12 (0.49) | −0.15 (0.38) | −0.19 (0.27) | +0.0060 (n 18) / +0.0126 (n 17) (0.50) |

Pericalcarine set (test subjects, mean over subjects): responsive vertices and their share of the responsive set.

- 3.0 mm: 2173 pericalcarine responsive vertices per subject (5.0% of the responsive set).
- 4.0 mm: 1500 pericalcarine responsive vertices per subject (4.7% of the responsive set).

##### S16.5 Group leakage maps of the 3T acquisition (S11.11 C, descriptive; Figure S2)

| grid | group-mean far leak (%) | cortex > 5% in the group map (%) | cortex > 10% in the group map (%) |
|---|---|---|---|
| 2.0 mm | 3.00 | 17.3 | 3.4 |
| 3.0 mm | 3.77 | 28.0 | 9.2 |
| 4.0 mm | 4.02 | 32.7 | 15.1 |

Regions with the highest far leak at 2.0 mm (area-weighted mean over the region, mean over subjects; share of the region above 10%):

| region | mean far leak (%) | share above 10% (%) |
|---|---|---|
| cuneus | 7.7 | 28 |
| pericalcarine | 7.5 | 24 |
| transversetemporal | 6.4 | 22 |
| lingual | 6.1 | 21 |
| bankssts | 5.6 | 22 |
| isthmuscingulate | 5.6 | 19 |
| posteriorcingulate | 5.5 | 20 |
| postcentral | 5.0 | 14 |

Regions with the highest far leak at 4.0 mm (area-weighted mean over the region, mean over subjects; share of the region above 10%):

| region | mean far leak (%) | share above 10% (%) |
|---|---|---|
| cuneus | 12.0 | 41 |
| posteriorcingulate | 11.0 | 46 |
| bankssts | 10.2 | 41 |
| transversetemporal | 9.7 | 42 |
| lingual | 8.2 | 32 |
| postcentral | 8.1 | 32 |
| parsopercularis | 7.8 | 35 |
| pericalcarine | 7.6 | 30 |

###### Why the group map's mean is lower than the individual maps' (check added after the maps)

Per subject, the area-weighted mean far leak of standard mapping in subject space (midthickness areas, analysis vertices; as in `nyu_test.md`) and the same map resampled to fsaverage (three nearest vertices on `?h.sphere.reg`, inverse-distance weights) and weighted by fsaverage's midthickness areas, as the group map is. Script: `src/17_nyu_3t_replication/gm_meancheck.py` (descriptive, added after the group maps were computed).

| grid | subject space, mean over subjects (%) | resampled to fsaverage, mean over subjects (%) | subjects lower after resampling |
|---|---|---|---|
| 2.0 mm | 3.21 | 3.00 | 43/43 |
| 3.0 mm | 4.24 | 3.78 | 43/43 |
| 4.0 mm | 5.00 | 4.02 | 43/43 |

The group map's mean (3.00, 3.77, 4.02%) equals the mean of the resampled individual maps: averaging across subjects does not lower it; resampling to the average surface and its area weights does.

| subject | 2.0 subject | 2.0 fsaverage | 3.0 subject | 3.0 fsaverage | 4.0 subject | 4.0 fsaverage |
|---|---|---|---|---|---|---|
| sub-wlsubj001 | 2.85 | 2.68 | 3.81 | 3.40 | 4.74 | 3.90 |
| sub-wlsubj004 | 3.52 | 3.24 | 4.76 | 4.23 | 5.35 | 4.09 |
| sub-wlsubj006 | 2.59 | 2.51 | 3.07 | 2.83 | 4.02 | 3.39 |
| sub-wlsubj007 | 2.65 | 2.49 | 3.16 | 2.79 | 4.00 | 3.29 |
| sub-wlsubj014 | 2.84 | 2.72 | 3.70 | 3.47 | 4.41 | 3.84 |
| sub-wlsubj019 | 3.05 | 2.84 | 3.85 | 3.37 | 4.41 | 3.43 |
| sub-wlsubj023 | 3.23 | 3.17 | 4.67 | 4.44 | 5.64 | 5.12 |
| sub-wlsubj042 | 2.85 | 2.62 | 3.73 | 3.23 | 4.47 | 3.46 |
| sub-wlsubj043 | 3.01 | 2.84 | 4.01 | 3.65 | 5.06 | 4.28 |
| sub-wlsubj045 | 2.52 | 2.34 | 3.32 | 2.85 | 4.32 | 3.42 |
| sub-wlsubj046 | 3.77 | 3.39 | 5.04 | 4.37 | 5.76 | 4.56 |
| sub-wlsubj056 | 3.28 | 3.12 | 4.48 | 4.05 | 5.49 | 4.50 |
| sub-wlsubj057 | 3.56 | 3.26 | 4.64 | 4.07 | 5.27 | 4.26 |
| sub-wlsubj062 | 3.13 | 2.88 | 4.21 | 3.68 | 4.87 | 3.80 |
| sub-wlsubj064 | 3.62 | 3.43 | 4.94 | 4.49 | 5.74 | 4.66 |
| sub-wlsubj067 | 3.19 | 2.97 | 4.15 | 3.68 | 5.00 | 4.08 |
| sub-wlsubj071 | 3.52 | 3.26 | 4.43 | 3.85 | 5.28 | 4.18 |
| sub-wlsubj076 | 3.33 | 3.00 | 4.32 | 3.67 | 5.12 | 3.91 |
| sub-wlsubj079 | 3.83 | 3.50 | 4.93 | 4.32 | 5.61 | 4.43 |
| sub-wlsubj081 | 2.38 | 2.26 | 3.08 | 2.75 | 3.81 | 3.16 |
| sub-wlsubj083 | 3.47 | 3.27 | 4.50 | 4.04 | 5.45 | 4.37 |
| sub-wlsubj084 | 2.76 | 2.63 | 3.44 | 3.16 | 3.99 | 3.24 |
| sub-wlsubj085 | 3.79 | 3.57 | 5.11 | 4.53 | 5.85 | 4.62 |
| sub-wlsubj086 | 3.22 | 2.90 | 4.33 | 3.80 | 5.29 | 4.19 |
| sub-wlsubj087 | 3.01 | 2.84 | 3.94 | 3.52 | 4.73 | 3.82 |
| sub-wlsubj088 | 3.58 | 3.30 | 4.97 | 4.38 | 5.73 | 4.50 |
| sub-wlsubj090 | 3.75 | 3.51 | 5.06 | 4.47 | 5.84 | 4.60 |
| sub-wlsubj091 | 3.67 | 3.58 | 4.95 | 4.62 | 5.02 | 3.89 |
| sub-wlsubj092 | 3.27 | 3.05 | 4.47 | 4.02 | 5.48 | 4.47 |
| sub-wlsubj094 | 2.53 | 2.39 | 3.11 | 2.73 | 3.68 | 2.75 |
| sub-wlsubj095 | 2.46 | 2.36 | 3.34 | 3.06 | 4.20 | 3.42 |
| sub-wlsubj104 | 2.47 | 2.33 | 3.05 | 2.71 | 3.61 | 2.92 |
| sub-wlsubj105 | 2.70 | 2.50 | 3.60 | 3.15 | 4.29 | 3.30 |
| sub-wlsubj109 | 3.68 | 3.36 | 4.94 | 4.34 | 5.94 | 4.75 |
| sub-wlsubj114 | 3.27 | 3.05 | 4.43 | 3.91 | 5.37 | 4.29 |
| sub-wlsubj115 | 3.12 | 2.91 | 4.18 | 3.66 | 4.68 | 3.64 |
| sub-wlsubj116 | 3.26 | 2.98 | 4.39 | 3.89 | 5.17 | 4.13 |
| sub-wlsubj117 | 3.21 | 3.04 | 4.10 | 3.70 | 4.78 | 4.01 |
| sub-wlsubj118 | 4.22 | 4.06 | 5.73 | 5.32 | 6.12 | 5.00 |

| subject | 2.0 subject | 2.0 fsaverage | 3.0 subject | 3.0 fsaverage | 4.0 subject | 4.0 fsaverage |
|---|---|---|---|---|---|---|
| sub-wlsubj120 | 3.64 | 3.38 | 4.46 | 3.85 | 5.14 | 4.06 |
| sub-wlsubj121 | 3.31 | 3.08 | 4.47 | 4.01 | 5.24 | 4.20 |
| sub-wlsubj122 | 3.32 | 3.06 | 4.38 | 3.90 | 5.13 | 4.06 |
| sub-wlsubj126 | 3.73 | 3.52 | 4.91 | 4.45 | 5.86 | 4.81 |

#### S16.6 Contamination from an anatomical atlas (S11.12)

D = gain(highest bin with ≥ 30 vertices) − gain(0–2% bin), gain = residual(model) − residual(competitor); negative: the gain grows with the dose. One-sided Wilcoxon (less), bootstrap 95% interval of the mean (10,000 resamples, seed 0), Holm across A1 and A2.

Reproducibility of the week-17b rerun (model and competitor residuals, D_CWL-M, D_CWL-P): 3.0 mm: largest difference 5.3e-13 (residuals), 6.6e-12 (D) over 35 subjects; 4.0 mm: largest difference 1.1e-16 (residuals), 1.4e-16 (D) over 35 subjects.

##### Tests

| grid | test | mean | subjects with D < 0 | bootstrap 95% | p (one-sided) | p (Holm, 2) |
|---|---|---|---|---|---|---|
| 3.0 mm | A1: D_CWL-A < 0 | −0.0056 | 20/35 | [−0.0131, +0.0017] | 0.122 | 0.122 |
| 4.0 mm | A2: D_CWL-A < 0 | −0.0088 | 24/35 | [−0.0170, −0.0007] | 0.014 | 0.029 |

##### Descriptive

Dose contrasts with each response estimate (mean, subjects with D < 0, one-sided p):

| grid | CWL-A (atlas, all areas) | CWL-A3 (atlas, V1–V3) | CWL-M (native map) | CWL-P (coarse data) | D_CWL-A − D_CWL-M | D_CWL-A − D_CWL-P |
|---|---|---|---|---|---|---|
| 3.0 mm | −0.0056 (20/35, p 0.122) | −0.0081 (20/35, p 0.013) | −0.0151 (25/35, p 0.004) | +0.0001 (21/35, p 0.376) | +0.0095 (two-sided p 0.048) | −0.0057 (two-sided p 0.413) |
| 4.0 mm | −0.0088 (24/35, p 0.014) | +0.0031 (18/35, p 0.654) | −0.0113 (25/35, p 0.007) | +0.0026 (19/35, p 0.593) | +0.0025 (two-sided p 0.481) | −0.0114 (two-sided p 0.030) |

Gain (model − competitor) by CWL-A bin (responsive vertices with < 2% far leak at 2.0 mm), mean (subjects with the bin):

| grid | 0–2% | 2–5% | 5–10% | 10–20% | > 20% |
|---|---|---|---|---|---|
| 3.0 mm | +0.0059 (35) | +0.0016 (35) | +0.0048 (35) | +0.0002 (35) | +0.0003 (35) |
| 4.0 mm | −0.0055 (35) | −0.0124 (35) | −0.0160 (35) | −0.0120 (35) | −0.0143 (35) |

How close each measure comes to the fine-data measure: Spearman correlation with CWL-M over the responsive vertices, median over subjects and directions (and the share of responsive vertices that receive an atlas response):

| grid | CWL-A vs CWL-M | CWL-A3 vs CWL-M | CWL-P vs CWL-M | responsive vertices with an atlas response |
|---|---|---|---|---|
| 3.0 mm | 0.53 | 0.43 | 0.88 | 57% |
| 4.0 mm | 0.53 | 0.46 | 0.81 | 56% |

**S17. The full-length text (draft v1.5, 2026-10-07), Sections 2–4**

The theory, methods and results of the full-length draft that preceded the condensed main text, verbatim except that figures are cited in the main text's numbering (its Figure 10 is Fig. S1). Section numbers inside it refer to its own sections.

#### 2. Theory

##### 2.1 Forward model

Let f be the BOLD response field on the cortical sheet, represented at column quadrature points between the white and pial surfaces (a column-mean component b0, optionally a depth gradient b1). The acquired volume is $y = K f + \varepsilon$. K is the effective-PSF operator:

- readout blur from T2* decay over the echo train, with the effective echo spacing and number of phase-encoding lines;
- an in-plane k-space taper;
- the slice profile;
- interpolation blur;
- each evaluated in the scanner's voxel frame through the registration.

Standard ribbon mapping is a vertex × voxel matrix W (Connectome Workbench weights; Marcus et al., 2011). The standard surface map is therefore $z = W y = A f + W \varepsilon$, with $\mathbf{A = W K}$. Row $a_v$ of A is the averaging kernel of standard mapping at vertex v.

##### 2.2 Far leak

Call cortex *far* from v if it lies in the other hemisphere, or more than 15 mm from v along the surface (measured on the registration sphere scaled to the hemisphere's cortical area, radius $\sqrt{A/4\pi}$): a proxy for the surface geodesic that is least accurate where the sphere is most distorted; the 15 mm is a convention). The **far leak** is the share of row $a_v$ that falls on far cortex: $\lambda_v = \Sigma_{u \in F(v)} a_{vu} / \Sigma_u a_{vu}$, with both sums over the cortical unknowns, so that $\lambda_v$ is a share of the cortical mass of the readout (the non-cortical part of a voxel is not in A). It depends only on the acquisition and the anatomy (Fig. 1A). It is the quantity in the leakage maps. Section 4.2 gives the figures at 10 and 20 mm, for the other hemisphere alone and with true geodesic distances.

##### 2.3 Contamination-weighted leak

Mixing harms a map only if the mixed cortex responds differently. For a response estimate x (maps j), the change of the standard map at v caused by far cortex is

$$c_{v,j} = \Sigma_{u \in F(v)} a_{vu} (x_{u,j} - x_{v,j}) / \Sigma_u a_{vu},$$

and the contamination-weighted leak (CWL) is $\|c\_v\|$ divided by the root-mean-square response over the scored vertex set, so that the bin edges used below (2, 5, 10, 20%) are fractions of the typical response. Four versions differ in where x comes from: CWL-M, the source run's own native-resolution map; CWL-P, the coarse data alone (upsampled and mapped); CWL-S4, the native map smoothed 4 mm on the surface; CWL-Q, the model's own estimate from the coarse data. With x from the coarse data alone, CWL is a data-driven map of where correction is expected to pay. When far cortex responds like the vertex itself ($x\_u \approx x\_v$), CWL is small however large $\lambda\_v$ is.

#### 2.4 Estimator

We estimate f as the posterior mean under a Gaussian likelihood and a Matérn ($\nu = 1$) prior on each hemisphere's surface. The prior is implemented through the SPDE–GMRF link with finite elements on the subject's mesh (Lindgren, Rue & Lindström, 2011). The estimate is rendered into the target's observation domain as $W\_t\ K\_t\ \hat{f}$. Noise is either independent per voxel, with a variance estimated per voxel and run (HCP: the mean power at the eight frequency bins neighbouring the stimulus frequency; StudyForrest and ds002702: the run-to-run variance of the betas), propagated through any emulation and calibrated once on the coarse runs by one factor (observed / propagated variance), or stationary and spatially correlated, with its spectrum estimated from the source's run residuals (StudyForrest only). The prior range and amplitude are hyperparameters (HCP: 12 mm and 0.125 at 2.4 mm, 24 mm and 0.25 at 3.2 mm, chosen on the five tuning subjects; Supplementary S3).

#### 2.5 Resolvability (Supplementary S10)

A sampling-limit theory of what a lattice of voxels can resolve on the folded sheet (a resolvable tangential wavelength of about 3.1–3.5 mm at 3T and 2.5–2.8 mm at 7T map-level CNR) is given in Supplementary S10. It did not predict the achieved resolution of the regularised estimator, which the PSF and the prior set, and it is kept as background only.

### 3. Methods

#### 3.1 Data

| dataset | subjects | acquisitions used | role |
|---|---|---|---|
| HCP 7T retinotopy (Benson et al., 2018) | 20 for development (5 tuning, 15 development; family-wise split fixed in advance); 158 further subjects in 86 families as a confirmatory cohort, plus 5 as a sensitivity set | 7T 1.6 mm, polar-angle wedges (CCW and CW runs) and, in the cohort, eccentricity rings (expanding and contracting runs); 2.4 / 3.2 mm emulated in k-space | operator validation, emulation, leakage maps, dose-response; hashed confirmatory tests (Section 3.9) |
| StudyForrest multi-resolution (Sengupta et al., 2017a, 2017b, 2018) | 7 (3T sessions in 5) | 7T 0.8 / 1.4 / 2.0 / 3.0 mm, 3T 1.4 / 2.0 / 3.0 mm; left / right hemifield orientation stimulus, 10 runs per session | real cross-session test; within-session emulation |
| OpenNeuro ds002702 (Kay et al., 2019, 2020a, 2020b; Markiewicz et al., 2021) | 5 | 7T GE 0.8 mm (5 subjects, 9 runs); 3T GE 2.4 mm (2 subjects); 7T SE 1.05 mm (2 subjects); six eccentricity rings | third within-session dataset; real second acquisitions |

| dataset | subjects | acquisitions used | role |
|---|---|---|---|
| NYU Retinotopy Dataset, OpenNeuro ds003787 (Himmelberg et al., 2021) | 44 (8 tuning, 36 test; 43 included) | 3T 2.0 mm, multiband 6, sweeping bars, 4–12 runs; 3.0 / 4.0 mm emulated in k-space | independent pre-specified replication of the emulation result (Section 3.10) |

HCP restricted family information was used only to form family-wise splits and to pick one subject per family for the cohort's primary tests; no restricted information is reported. ds002702 and ds003787 were processed with the authors' FreeSurfer reconstructions (Fischl, 2012).

##### 3.2 Preprocessing

- **HCP:** the released MNINonLinear time series, mapped with the HCP pipelines' own ribbon weights and goodvoxels masks. Responses are the stimulus-frequency Fourier components of each wedge run.
- **StudyForrest and ds002702:** motion correction to the first run (MCFLIRT; Jenkinson et al., 2002), then registration of the session mean to the FreeSurfer anatomy.
    - StudyForrest: bbregister, anchored within subjects.
    - ds002702: FLIRT from three starts, then boundary-based registration (Greve & Fischl, 2009), on canonically reoriented copies. Registration was checked by the EPI grey–white contrast across the white surface against copies shifted by 2 mm.
    - Then a per-run GLM: SPM-style HRF with temporal derivative, Legendre drifts, motion; betas in percent signal change. StudyForrest's left- and right-hemifield betas are combined as one complex map (left + i·right); ds002702's six ring conditions are six real maps.
    - ds002702 field-map correction (Jezzard & Balaban, 1995; FSL fugue, Jenkinson et al., 2012; the field map closest to run 1) was applied per session only where it improved that registration check.
    - The per-run GLMs run sequentially, and every session passes a per-run consistency check (Supplementary S9).

##### 3.3 Operators and emulation

- **Operator.** Kernels are tabulated per voxel axis from the acquisition parameters; columns are integrated at quadrature points between the white and pial surfaces.
    - Checked against brute-force integration of the PSF over each column: maximum relative error $\approx 10^{-7}$.
    - Column mass 0.99–1.01.
    - Effective PSF FWHM at HCP 1.6 mm: 2.56 / 2.70 / 2.55 mm (x / y / z). In voxels at MNI scale 1.0: the sinc of the voxel alone 1.21; with the Tukey-0.3 taper 1.41; with the T2* decay over the echo train 1.56 along the phase-encoding axis; with the 0.4 mm interpolation blur 1.46 / 1.62 / 1.40; the slice profile 1.32 before the blur. The taper and the blur

are modelling assumptions, the T2* term uses the protocol's echo spacing (Supplementary S15.1).

- **Emulation.** Coarser acquisitions are emulated by cropping k-space and applying a Tukey window ($\alpha = 0.5$) that is zero at the coarse Nyquist frequency, applied frame by frame to the FIX-cleaned time series as released (voxels with data at 1.6 mm). This is exact per axis, including on oblique grids, and keeps the noise and physiology of the real data. It is an emulation, not a coarse acquisition: the window is a modelling choice (a sharp crop would leave sinc ringing that no compact operator represents), the emulated PSF is 1.6–1.7 voxels wide at every grid, the noise is band-limited together with the signal, and the signal-to-noise gain is at most the square root of the voxel-volume ratio, less than a real coarse acquisition would give. The 2.4 and 3.2 mm leak figures and maps are for this emulated PSF; Section 4.2 gives them for native-acquisition kernels at those voxel sizes as well.
    - Emulated effective PSF: 3.8–4.1 mm at 2.4 mm, 5.1–5.4 mm at 3.2 mm.
    - The operator built with the emulated kernel agrees with the emulation applied to synthetic fields to 0.2–1.2% (relative L2).
- **Within-session designs.**
    - StudyForrest and ds002702: odd and even runs form two halves. The source half is coarsened; the target is the other half at native resolution; both directions are averaged.
    - HCP: each wedge run is coarsened and predicts the other run's 1.6 mm map.

##### 3.4 Competitors

All competitors received the same data and the same setting-selection rule as the model:

- standard mapping of the coarse data, ± surface smoothing (implicit heat-equation smoothing, FWHM 2–16 mm);
- band-limited (k-space) upsampling to the target grid, then the target's mapping, ± smoothing;
- across sessions, resampling to the target grid (cubic B-spline or trilinear), ± smoothing.

In the misregistration test the competitors received the same registration error as the model, as a band-limited shift .

##### 3.5 Scoring

The score is the residual energy fraction of the target map after the best single scaling of the prediction (one complex scalar for wedge maps, one real scalar otherwise, fitted within each vertex set or bin that is scored):

- complex maps: phase-conjugated across wedge directions in HCP, real and imaginary parts in StudyForrest;
- ds002702: each ring map scaled separately.

Lower is better. Vertex sets:

- covered (both sessions sample the vertex);
- visual (occipital labels of the Desikan–Killiany atlas; Desikan et al., 2006) or coherent (HCP: response coherence > 0.3 in the 1.6 mm standard maps);
- strata by far leak or CWL at the coarse grid, restricted to vertices clean at the native grid (far leak < 2%).

##### 3.6 Choosing settings without target leakage

Three rules were used, always the same for every method:

- fixed in advance on separate tuning subjects (HCP: 5 subjects; the same rule chose the model's range and amplitude and the competitors' smoothing widths);
- leave-one-subject-out;
- for real second acquisitions, chosen on the other subject of the same pair type.

For ds002702 within session the search grids were widened once, before any subject's results were summarised, after the first fits had their optimum at the edge of the initial grid (Supplementary S3). A rule tuned in one regime was not reused in another without saying so. Section 4.6 shows why.

##### 3.7 Leakage maps

Per-subject far-leak maps were computed at 1.6 mm and at emulated 2.4 / 3.2 mm. They were resampled to fs_LR 32k with each subject's MSMAll sphere (Robinson et al., 2014, 2018; area-corrected barycentric), averaged across 19 subjects (one lacked an MSMAll sphere), and summarised by Desikan region (area-weighted).

##### 3.8 Statistics

The unit of analysis is the subject. The primary dose-response contrast is D = gain in the highest contamination bin (≥ 30 vertices) − gain in the 0–2% bin, where gain = residual(model) − residual(best competitor). It is tested with a one-sided Wilcoxon signed-rank test across subjects, with a bootstrap 95% interval of the mean.

The tests, bins, vertex sets and settings were fixed in plans written before the analyses were run (2026-09-30 and 2026-10-01), with their SHA-1 sums recorded at the time; the plans are released with the code. This is pre-specification attested by the authors, not a registration with a third party: no external timestamp exists. `STATS_PLAN.md` covers the first pre-declared analysis (contamination-weighted leak) and `STATS_PLAN_w9.md` the second (a denoised response estimate). Both were written after the development-set analyses of Sections 4.1–4.3 and the far-leak dose-response of Section 4.4 were known and before the contamination measure was computed, so the development-set tests are pre-specified within an explored dataset; the bin edges, the 2% clean criterion and the 30-vertex rule were set on the development set and carried into every later plan. Analyses run before these plans are reported with subject counts and are labelled descriptive. The summary table across datasets (Section 4.8) was specified before it was computed (`TABLE_SPEC.md`), and the confirmatory cohort's hypotheses, subject set and tests before any of its subjects was fitted (`COHORT_PLAN.md`, 2026-10-02; Section 3.9). No StudyForrest or ds002702 result is confirmatory in the sense of Section 4.9.

##### 3.9 Confirmatory cohort

After the analyses above, every other subject of the HCP 7T retinotopy release was processed with the same pipeline and the settings fixed in Section 3.6, under an analysis plan written on 2026-10-02, before the first of them was fitted (`COHORT_PLAN.md`, Supplementary S11.4; pre-specification attested by the authors' recorded hashes, not registered with a third party):

- **Subjects.** 158 subjects in 86 families: the 164 7T retinotopy subjects outside the 20 development subjects, minus six — five relatives of the tuning subjects (one of whom also lacks the MSMAll sphere) and one without a concatenated retinotopy run. Twins and siblings are not independent, so the primary tests use one subject per family (the lowest subject ID; 86 subjects); all 158 are reported as a secondary analysis with a family-level bootstrap. The five relatives were added afterwards as a labelled sensitivity set (Amendment 1, hashed separately; Supplementary S11.5).
- **Pipeline per subject.** Operator, leakage and wedge maps as in Sections 3.2–3.3; fits at emulated 2.4 and 3.2 mm with the PSF as modelled and stretched by ±15%; fits from the first and from the second half of each wedge run (4 of 8 cycles; both halves fitted and averaged), scored against the same full-run reference; the same chain for the eccentricity runs (expanding and contracting rings, which have the wedges' period); contamination maps as in Sections 2.3 and 4.4.
- **Hypotheses (eight primary one-sided tests; no correction across hypotheses was planned; Holm within each family of secondary tests).** H1 and H4 were set before any cohort-type data were seen; H2 and H3 were derived from the data of the 15 development subjects, which were run through the batch pipeline first (below), and then tested in the cohort. H1: the contamination dose-response of Section 4.4 replicates ($D < 0$ at both grids, primary; the coarse-data-only measure, the plain-leak contrast and the overall gain, secondary). H2: for rings, the plain-leak dose-response is shallower than for wedges at 3.2 mm (H2a); contamination predicts the ring gain (H2b, both grids), and more sharply than the plain leak does (H2c, secondary). H3: the gain where no far unmixing is needed (the 0–2% far-leak bin) decreases with half-run sources (H3a, both grids), the leak-dependent gain does not change (H3b, an interval), and the model still beats the competitor with half the data (H3c, secondary). H4: with the PSF assumed 15% too wide, the overall gain at 2.4 mm stays negative (primary; the other three PSF cases secondary). In H3 neither the model's prior nor the

competitor's smoothing was re-tuned for the noisier half-run sources; both stay at their full-run settings. A check with both re-tuned on the tuning subjects' half runs was run afterwards, under its own hashed plan, in the 15 development subjects (Section 4.9).

- **Statistics.** As in Section 3.8; a subject enters a test when every stratum the test needs has ≥ 30 vertices.
- **Interim look.** After 8 of the 16 batches had been scored, we computed every quantity of the table in Section 4.9 on those 80 subjects. No test, subject set or setting was changed afterwards; the only consequence was Amendment 1.

The exploratory look at the 15 development subjects did not bear out two prior expectations, that rings would show no far-leak dose-response and that a denoising gain would grow with source noise; H2 and H3 were given their present form before the plan was hashed, and the look is recorded in the plan. The development subjects are not part of the confirmatory tests.

##### 3.10 External replication: NYU Retinotopy Dataset

A replication of the emulation result in an independent dataset, specified and hashed before any fit (`PLAN_NYU.md`, Supplementary S11.10):

- **Data.** OpenNeuro ds003787 (Himmelberg et al., 2021): 44 subjects, Siemens Prisma 3T, 2.0 mm isotropic, multiband 6, sweeping-bar stimulus in 4–12 runs of 192 s, spin-echo field maps, FreeSurfer reconstructions. Split fixed with a seed: 8 tuning and 36 test subjects.
- **Preprocessing.** Field estimation from the spin-echo pair (topup; Andersson et al., 2003); motion correction to the first run's single-band reference; distortion correction (applytopup); boundary-based registration to the FreeSurfer anatomy. Runs displaced by more than 4 mm from the reference are excluded; one test subject kept a single run and was excluded.
- **Maps.** Per run, a GLM with the bar aperture's coverage of the visual field weighted by 1, cos θ and sin θ (HRF-convolved; the three regressors are uncorrelated in the aperture sequence), drifts and motion. The complex map β_cos + i·β_sin is the first angular harmonic of each voxel's response; its phase estimates polar angle, as the wedge's Fourier phase does in HCP. Odd and even runs form two halves.
- **Emulation, methods, score** as in HCP: each half's series coarsened in k-space to 3.0 and 4.0 mm (×1.5, ×2); the other half's native 2.0 mm map as the target; standard mapping, smoothing, upsampling, upsampling + smoothing and the model; the residual of Section 3.5 over responsive vertices (run-averaged native $R^2 \geq 0.2$ for the stimulus regressors).
- **Selection and tests.** Settings chosen on the tuning subjects by the week-3 rule; on the test subjects, N1: the model beats the chosen competitor over responsive vertices; N2: D_far < 0; N3: D_CWL-M < 0 (Section 3.8), one-sided Wilcoxon, Holm across the six tests. One deviation, recorded before any test fit: the model's prior grid was extended to weaker priors and shorter ranges because the first tuning subject's optimum lay at the planned grid's edge (Supplementary S16).

#### 4. Results

##### 4.1 The operator captures how the maps change under emulated coarsening

Standard ribbon mapping on real HCP geometry:

| grid | dilution of a unit field | mean far leak | cortex with far leak > 5% / > 10% | pericalcarine far leak |
|---|---|---|---|---|
| 1.6 mm | 0.80 | 2.1% | 13% / 4.7% | 3.8% |
| 2.4 mm (emulated) | 0.73 | 2.6% | 18% / 9.8% | 5.0% |
| 3.2 mm (emulated) | 0.69 | 3.6% | 26% / 16.8% | 6.5% |

Leak rises steeply as the opposite bank comes close (Fig. 1B). At 1.6 mm it is 12.8% where the opposite bank is within 2.25 mm, and 0.3% beyond 6 mm.

The operator predicts how the real HCP wedge maps change under emulated coarsening of the same data. Predicting the emulated 2.4 / 3.2 mm maps from the 1.6 mm field estimate, adding the operator's far part to its near part lowered the residual in **all 20 subjects (40 of 40 hemispheres) at both grids**. Where the operator predicts a leak above 10%, the residual fell from 0.095 to 0.059 at 2.4 mm and from 0.124 to 0.059 at 3.2 mm. A far part shuffled across vertices made the prediction worse in all 20 subjects (0.150 and 0.204). The predicted mixing is therefore geometrically right, not just right on average (Fig. 2). Two limits of this test: the emulated maps and the prediction share the emulation window, so it checks the geometry of the far part, not the native PSF model; and the near-only predictor also lacks the far part's mass, so part of its deficit could be normalisation rather than geometry. Both limits are met by the real-data test that follows.

**Real coarser acquisitions.** StudyForrest has 7T sessions of the same subjects at 0.8, 1.4, 2.0 and 3.0 mm (Section 3.1). With the week-4 registrations and the native kernel of each session, the finer session's standard map divided by its dilution predicts the coarser session's standard map through the coarser session's operator, as in the emulation test. Pooled over the six session pairs of each of the 7 subjects, in visual cortex where the predicted far leak exceeds 10%, adding the far part lowers the residual from 0.378 (near part alone) to 0.345 (7 of 7 subjects); a far part shuffled across vertices makes it worse (0.426, 7/7); a near part renormalised to the full row mass does not help (0.373, 7/7); and where the predicted leak is below 2% the four predictors agree (0.320). The primary pair, 1.4 → 3.0 mm, gives 0.344 → 0.303 with the far part, 0.417 shuffled, 0.337 renormalised. Cross-session differences raise every residual, but the paired comparison stands: the operator's far part predicts real coarser acquisitions where the leak is high, with the right geometry (Fig. 3B; Supplementary S14.3).

##### 4.2 Leakage maps

The group-mean far leak is 2.0–2.1%, 2.3–2.4% and 3.1–3.3% at 1.6, 2.4 and 3.2 mm. Leak above 5% covers 4%, 10% and 22% of cortex; above 10%, 0.3%, 1% and 4%. Group means are lower than individual maps (Section 4.1) because hot spots do not align perfectly across subjects.

Regions with the highest leak at 3.2 mm, as area-weighted mean far leak (share of the region above 10%):

| region | mean far leak | share above 10% |
|---|---|---|
| posterior cingulate | 8.7% | 40% |
| cuneus | 8.4% | 35% |
| banks of the STS | 7.0% | 33% |
| transverse temporal gyrus | 6.5% | 32% |
| pericalcarine | 6.3% | 27% |
| lingual | 6.2% | 27% |
| postcentral | 5.9% | 25% |

The central sulcus appears as a band on both banks. Frontal and temporal poles and orbitofrontal cortex are lowest (< 1.5%) (Fig. 4).

**Realistic kernels.** The 2.4 and 3.2 mm figures above are for the emulated kernel. Run with the kernel that our PSF model gives a native acquisition at those voxel sizes (no emulation window), the same operator machinery yields, in the same 20 subjects, a lower net far leak (mean 1.3% and 2.4% against 2.6% and 3.6%; cortex above 5%: 14.9% and 24.3% against 17.9% and 25.8%), and with partial Fourier 7/8 along the phase-encoding axis one as high or higher (2.0% and 3.8%; 18.0% and 29.8% of cortex above 5%). The gross far weight is higher for both native kernels (13.3–14.1% and 24.9–26.0% against 11.8% and 21.1%), because a sinc has side lobes that the emulation window removes. The emulated kernel therefore sits within the range of realistic kernels, and the leak figures should be read with that range in mind (Fig. 3C; Supplementary S14.2).

**Kernel width.** The figures scale with the kernel's width: with the kernels 15% narrower (the width the inversion preferred in a scan of the PSF model, Section 4.3), the net far leak in the same 20 subjects is 2.0% and 2.8% instead of 2.6% and 3.6% at 2.4 and 3.2 mm, and cortex above 5% is 14% and 20% instead of 18% and 26% (Supplementary S15.2). The PSF model's width is thus the leading uncertainty in the leak figures, in the direction of lower leak.

**Far criterion.** With the threshold at 10 or 20 mm instead of 15, the figures scale smoothly: in the same 20 subjects (individual-map means), the net far leak is 4.4 / 2.6 / 1.6% at 10 / 15 / 20 mm at 2.4 mm and 5.1 / 3.6 / 2.3% at 3.2 mm, and the share of cortex above 5% is 30 / 18 / 11% and 36 / 26 / 17%. The other hemisphere alone contributes 0.2–0.35%: the far leak of these acquisitions is same-hemisphere mixing across sulci and between neighbouring gyri, and the 15 mm convention sets the scale of the figures, not their pattern. Measured with true geodesic distances along the midthickness surface instead of on the sphere (five subjects; mesh-graph distances calibrated against exact geodesics), the maps agree with the sphere version (per-vertex r = 0.86–0.90; the > 5% label changes on 4–9% of cortex), and the sphere slightly overstates the net far leak at the coarse grids: 2.8% against 2.6% at 2.4 mm and 4.0% against 3.6% at 3.2 mm, in 5 of 5 subjects (Fig. S1B, C; Supplementary S15.4, S15.5).

##### 4.3 Emulation in HCP: the model helps everywhere, most where mixing is strong

The 15 development subjects, settings fixed on the five tuning subjects; cross-run residual against the other run's 1.6 mm map (Fig. 5A; the competitor here is the one best on average, in Section 4.8 the per-subject best, hence 13/15 here against 12/15 there):

| grid | standard mapping | standard + smoothing | upsampling + 1.6 mm mapping + smoothing | model | model better than the best competitor |
|---|---|---|---|---|---|
| 2.4 mm | 0.422 | 0.341 | 0.321 | **0.313** | 13/15 |
| 3.2 mm | 0.458 | 0.352 | 0.339 | **0.325** | 14/15 |

- **Noise ceiling.** The other 1.6 mm run itself, as the predictor, scores 0.362 raw and 0.305 after 6 mm surface smoothing over all coherent vertices (0.344 / 0.331 where leak > 10% at 2.4 / 3.2 mm): the target is a single run, and no prediction can do much better than that. Against this ceiling the model closes 51% (2.4 mm) and 41% (3.2 mm) of the gap left by upsampling + smoothing over all coherent vertices; where leak > 10% it reaches the ceiling at 2.4 mm (0.343 against 0.344) and closes 68% at 3.2 mm (Supplementary S5.8).
- **Strong mixing.** Where unmixing is needed (clean at 1.6 mm, leak > 10%), the model's lead grows to −0.024 (2.4 mm) and −0.027 (3.2 mm), 14/15 subjects each.
- **Bank contrast.** The model keeps more of the polar-angle difference across sulcal banks: dissimilarity 0.75 vs 0.61 for standard mapping and 0.68 for upsampling at 2.4 mm.
- **Averaging kernels.** At 360 sample vertices per grid (24 per subject) the model's kernels have no net far leak (standard mapping: 11–13%), but their gross far weight falls only from 19% to 14% at 2.4 mm and from 26% to 23% at 3.2 mm: the estimator cancels the mean of the far field, it does not stop averaging over far cortex. The kernels are 0.9 mm wider along the sheet.
- **Robustness.** A PSF assumed 15% too narrow fitted slightly better than the modelled one (0.311 against 0.313 at 2.4 mm, 0.322 against 0.325 at 3.2 mm; in the cohort −0.0109 against −0.0087 and −0.0158 against −0.0128), which suggests that the modelled native PSF is, if anything, slightly too wide. A scan of the kernel width on the five tuning subjects (stretch α = 0.7–1.3 on all axes, and 0.85 / 1.15 on the phase-encoding or the slice axis alone, prior settings fixed) puts the optimum on the narrow side: α = 0.85 lowers the residual in 5 of 5 subjects at both grids (0.3103 against 0.3126; 0.3182 against 0.3214), α = 0.7 gives the same residual as 0.85, and wider kernels cost more than narrower ones gain (α = 1.15: +0.004 and +0.007; α = 1.3: +0.010 and +0.017; 0 of 5 better); the phase-encoding axis alone recovers most of the improvement, the slice axis alone a smaller part. For scale, the model's lead over upsampling + 8 mm smoothing at these settings is 0.0047 and 0.0063. The optimum of a cross-run residual at fixed prior strength is a bias–variance trade-off, so a preference for α < 1 is compatible with a correct PSF as well as with a 15–30% over-estimate; the leak-dependent gain is the same at every α (D +0.020 against +0.021 at 2.4 mm), and α = 1 is kept throughout (Fig. S1A; Supplementary S15.1). A PSF assumed 15% too wide shrinks the lead, most at 2.4 mm, where it stays clear only where mixing is strong (Section 4.8); with 86 families the overall lead at 2.4 mm remains (Section 4.9). The lead also survives a translational misregis-

tration of up to 1 mm (a band-limited shift) given to every method: −0.008 / −0.009, 14/15 subjects.

- **Depth.** A depth-gradient component is superficial in every subject (b1/b0 = 0.24 at 2.4 mm, 0.34 at 3.2 mm) and lowers the residual slightly (−0.005 / −0.006, 15/15). Its size changes with the voxel size and the forward model has no compartment for pial veins or CSF, so it is a nuisance term that absorbs superficial signal, not a depth profile.
- **Correlated noise.** The likelihood treats the noise as spatially white. A likelihood with stationary, spatially correlated noise — its radial spectrum estimated from the eight neighbouring frequency bins of both runs, with 41–133 times the white power at wavelengths above 20 mm — made the estimates worse in every development subject: +0.054 at 2.4 mm and +0.026 at 3.2 mm over all coherent vertices, at the white likelihood's settings. Its maps have nearly the same pattern as the white fit's (complex correlation 0.95 and 0.97) at 73% and 84% of the amplitude: the likelihood discounts the large scales over which the retinotopic map varies, and the estimate shrinks. With its own prior chosen on the tuning subjects by the week-3 rule (r = 24 mm, s = 0.25 at 2.4 mm and r = 48 mm, s = 0.5 at 3.2 mm, against 12 / 0.125 and 24 / 0.25 for the white likelihood), under a plan hashed before those fits, it is still worse in every subject (+0.047 and +0.025), the best setting chosen on the development subjects themselves is the same one, and at 2.4 mm it loses the dose-response (D_far +0.002 against −0.024). The white likelihood therefore does not cost accuracy here, and we keep it (Fig. S1E; Supplementary S15.6, S15.7).
- **Known-truth simulation.** With the truth taken as the two runs' 1.6 mm maps averaged, smoothed 3 mm and undiluted, rendered through the coarse operator with noise of the empirical per-voxel power, and every method at its real-data settings, the model beats k-space upsampling clearly at 3.2 mm (0.037 against 0.057) but loses to it at 2.4 mm (0.029 against 0.028 over all coherent vertices; 0.039 against 0.032 where leak > 10%): with an exact operator and little noise, band-limited upsampling is itself near-exact at 2.4 mm, and the prior's smoothing costs more than the unmixing gains (Supplementary S5.3).
- **In degrees of polar angle.** Per coherent vertex, the mean absolute difference between the prediction from one coarsened run and the other run's 1.6 mm map is 29.7° for the model, 30.3° for upsampling + smoothing and 31.5° for standard mapping at 2.4 mm (30.3°, 30.8° and 32.7° at 3.2 mm); the two 1.6 mm runs themselves disagree by 31.6°, so the vertex-level error is dominated by single-run noise, and a smoothed or model-based estimate can agree with the other run better than a raw run does. The model's advantage over upsampling + smoothing is 0.5–0.6° over all coherent vertices (13–14 of 15 subjects) and 1.9–2.1° where mixing is strong (> 20% far leak; 11 of 14 and 14 of 15 subjects), against a noise floor of about 30° (23° at strongly coherent vertices) (Supplementary S5.7).
- **Far-part ablation.** Inverting with the far entries of the operator removed — per voxel, the columns farther than 15 mm from the voxel's main unknown or in the other hemisphere: 18% and 25% of the voxel mass on average at 2.4 and 3.2 mm — leaves the result almost un-

changed: overall residual 0.3126 against 0.3131 (2.4 mm) and 0.3250 against 0.3248 (3.2 mm); gain in the

20% bin −0.033 against −0.035 and −0.039 against −0.039; far-leak contrast −0.022 against −0.024 and −0.023 against −0.023 (D_far(near) − D_far(full) +0.0025, 10 of 15 subjects, p = 0.06; 0.0000, 7 of 15). A near part renormalised to the full row mass behaves the same (Fig. 3A; Supplementary S14.1). The inversion's gain and its dose-response do not come from the far part of the operator (Section 5).

##### 4.4 In HCP the gain grows with predicted mixing of different responses; in resolution-limited data this is not established

**Far leak (descriptive)** (Fig. 6A). In the HCP development set (cohort: Section 4.9) the model's gain over the best competitor rises steadily with the far leak of the coarse map:

- 2.4 mm: from −0.007 (clean, 13/15 subjects) to −0.035 above 20% (12/14; one subject had fewer than 30 such vertices);
- 3.2 mm: from −0.016 to −0.038 (14/15).

(These bins use the far leak saved with the operator. The pre-declared analysis below recomputed it from a rebuilt operator (Supplementary S9), so its far-leak bins differ by up to 0.002 per bin and give the plain-leak contrasts quoted next.)

**Contamination-weighted leak (first pre-declared analysis)** (Fig. 6B). The plan named ds002702 as its key prediction, HCP as the replication of a pattern already seen and StudyForrest as a third test. In HCP the primary contrast D is negative at both grids; the plain-leak contrast is −0.024 and −0.023 (12/15 and 13/15). The key prediction was not confirmed at the planned level (ds002702: p = 0.094; the smallest attainable p with five subjects is 0.031), and the StudyForrest primary test was null.

| dataset | n | D (CWL, source's own native map) | subjects with D < 0 | p (one-sided) | D with CWL from the coarse data alone |
|---|---|---|---|---|---|
| HCP 2.4 mm | 15 | −0.023 | 11/15 | 0.008 | −0.041 (14/15) |
| HCP 3.2 mm | 15 | −0.037 | 15/15 | < 0.001 | −0.037 (15/15) |
| ds002702 2.4 mm | 5 | −0.015 | 4/5 | 0.094 | −0.013 (4/5) |
| StudyForrest | 7 | −0.002 | 3/7 | 0.53 | −0.009 (5/7) |

At 3.2 mm, contamination separates the vertices where the model helps more sharply than the plain leak does: the gain goes from −0.011 (0–2%) to −0.047 (> 20%), against −0.017 to −0.039 for the plain leak. In the confirmatory cohort the same holds with smaller numbers: −0.011 to −0.030 by contamination, against −0.016 to −0.020 by the plain leak (all 158 subjects; Section 4.9, Fig. 7).

The measure computed from the coarse data alone predicts the gain as well as the one computed from fine data. A practical "where does correction pay" map therefore needs nothing beyond the coarse acquisition and the anatomy.

In ds002702 (eccentricity rings) at 2.4 mm, the plain leak predicted no gain at all (contrast +0.008, 2/5): rings drive the two banks of the calcarine alike. At the dataset's finer emulated grids, however, the same subjects show far-leak contrasts in the predicted direction (−0.024, 3/5, at 1.6 mm; −0.038, 5/5, at 1.05 mm; descriptive, Supplementary S8.1), and in HCP rings show a plain-leak dose-response of their own at 2.4 mm (Section 4.9): the ring stimulus is a negative control for the plain leak only where the far cortex within reach carries the same eccentricity. Contamination points the predicted way (4/5) but is not significant with five subjects. In StudyForrest the primary test is null. Binning by the share of the other hemisphere instead, which is where hemifield stimuli differ, shows the effect: −0.019, 6/7 ($p = 0.023$; exploratory, added after the plan; Supplementary S8.1).

**Denoised response estimate (second pre-declared analysis).** We recomputed the contamination after smoothing the source's native map on the surface (4 mm FWHM, primary), and with the model's own estimate from the coarse data. Both resolution-limited datasets moved in the predicted direction, but neither primary test reached significance:

| dataset | n | D, smoothed 4 mm (raw, first analysis) | subjects with D < 0 | p (one-sided) | gain in the > 20% bin | gain in the 0–2% bin |
|---|---|---|---|---|---|---|
| StudyForrest | 7 | −0.010 (−0.002) | 4/7 | 0.19 | −0.002 | +0.010 |
| ds002702 2.4 mm | 5 | −0.021 (−0.015) | 4/5 | 0.062 | −0.011 | +0.010 |

- In both, the model beats upsampling only in the highest-contamination bin and trails it below.
- Secondary: with 8 mm smoothing, ds002702 is negative in 5 of 5 subjects ($p = 0.031$; Holm 0.19).
- In HCP the denoised estimate reproduces the result of the first analysis: −0.028 and −0.033 at 2.4 and 3.2 mm (12/15 and 15/15 subjects); the model-based estimate gives −0.031 and −0.028 (14/15 and 15/15); Holm $p \leq 0.007$ throughout.

The evidence that contamination predicts the gain in resolution-limited 7T data therefore stays suggestive.

**Gating.** Using the model only where the predicted leak or contamination exceeds a threshold (chosen leave-one-subject-out) never beat the better of its two parts:

- HCP: the chosen threshold was always the lowest. The gated estimator then falls short of the model everywhere (leak gate: −0.004 vs −0.008 at 2.4 mm) or matches it (contamination gate: −0.008 vs −0.008).
- StudyForrest and ds002702: the gated estimator equals the competitor.

The vertices with strong harmful mixing are too few to move a region-wide score.

##### 4.5 Real 7T data within one session: resolution-limited targets

In StudyForrest the two halves of a session share registration, distortion and physiology; only resolution differs (Fig. 9A). Across 28 subject-cells (subject × session × coarsening factor: 1.4 and 2.0 mm sessions coarsened ×1.5 and ×2), the model trails the best competitor, band-limited upsampling with little or no smoothing. With a correlated-noise likelihood it is +0.014 in visual cortex (better in 9/28). It wins only where the coarse map draws more than 20% from the other hemisphere: −0.026, 16/20. Run-to-run noise is dominated by large spatial scales: power at wavelengths above 20 mm is tens to hundreds of times the white-noise level (26–69× in the subject shown in Supplementary S6.2; 35–500× across subjects and sessions). Modelling that correlation helps a little at 3–4 mm (−0.010 against white noise).

In ds002702 the model loses at small resolution gaps (in every one of the five subjects; one-sided p = 0.031 against it) and ties at a large one (Fig. 9B):

| coarse grid (from 0.8 mm) | model – best competitor | subjects better |
|---|---|---|
| 1.05 mm | +0.033 | 0/5 |
| 1.6 mm | +0.029 | 0/5 |
| 2.4 mm | +0.003 | 2/5 |

In both datasets the targets are averages of many runs, so they are resolution-limited rather than noise-limited. This is the regime in which a well-tuned band-limited competitor is hardest to beat.

##### 4.6 Real second acquisitions

- **StudyForrest, settings fixed in advance from HCP.**
  - The model predicted the target session's map worse than smoothing: +0.036 (better in 11 of 102 subject-pairs). Re-tuned leave-one-subject-out it was +0.016 (27/102).
  - A 3T session predicts the 7T session of the same voxel size about as badly as a finer one. Between-session differences (field, distortion, registration, physiology), not resolution, dominate these residuals.
- **ds002702, settings pre-specified from the within-session emulation.** The comparison was uninformative, although the model's residual is the lower one in 6 of 8 session pairs (Fig. 10, left): both methods carried within-session settings, where the best competitor needs no smoothing, whereas across sessions it needs 6–8 mm of smoothing and the model a stronger prior, so neither was at its best.
- **ds002702, settings chosen on the other subject of the pair type (a second analysis, added afterwards; Fig. 10).**
  - The model beat the best competitor for **3T 2.4 mm → 7T 0.8 mm after field-map correction in both subjects** (−0.021, −0.028; two subjects, each with settings chosen on the other, under a rule adopted after the primary analysis: suggestive, not confirmatory).
  - It lost clearly for **SE 1.05 mm → GE 0.8 mm** (+0.08 to +0.09). SE and GE BOLD differ at fine scales, and the forward model has no term for that.

- Field-map correction improved the best cross-session predictions by up to 0.12, more than the choice of estimator.

- **StudyForrest, the same rule.** It did not reproduce the ds002702 gain: 3T → 7T at a voxel ratio of at least 2.5 gave +0.014 (2/5 subjects better, p = 0.84; pre-declared). The model's deficit shrinks as the resolution gap grows, but it never becomes a clear gain. No field maps exist for these 3T sessions.

##### 4.7 Uncertainty (Supplementary)

The posterior is not calibrated as is.

- **Noise part.** The model's noise variance was underestimated by a factor of 4–5. One factor, estimated on the other hemisphere, restores nominal tail rates (6.0–6.1% beyond the 95% bound).
- **Prior part.** In the known-truth simulation the 95% intervals covered only 26–28% of responsive vertices: shrinkage bias from a prior tuned for point prediction is not in the posterior s.d.

##### 4.8 One set of statistics across datasets

The table restates the main comparisons with one set of statistics:

- per subject, model − best competitor, where the best competitor is the better competitor for that subject (stricter than Section 4.3, which compares with the competitor that is best on average); cells and directions are averaged within subjects;
- a bootstrap 95% interval of the mean, the subjects with the model better, and a one-sided Wilcoxon signed-rank p, uncorrected;
- high mixing: far leak > 10% at the coarse grid and < 2% at the native grid (StudyForrest within session: > 10% of the coarse map from the other hemisphere).

The table was specified, and hashed, before it was computed (Supplementary S11.3). It is descriptive: only the high-mixing values of ds002702 within session are new. Its competitor is the per-subject best, whereas Sections 4.3–4.6 compare with the competitor best on average, hence small differences (13/15 there against 12/15 here). A one-sided p near 1 means that the model is significantly worse; the intervals for n = 2 are not meaningful.

| comparison | settings chosen | all vertices: model – best competitor [95% CI] | better | p | high mixing: model – best competitor [95% CI] | better | p |
|---|---|---|---|---|---|---|---|
| **Emulation within one acquisition** | | | | | | | |
| HCP 7T, emulated 2.4 mm | fixed on 5 other subjects | −0.007 [−0.011, −0.002] | 12/15 | 0.006 | −0.022 [−0.031, −0.012] | 13/15 | < 0.001 |
| HCP 7T, emulated 3.2 mm | fixed on 5 other subjects | −0.013 [−0.017, −0.007] | 14/15 | 0.002 | −0.026 [−0.033, −0.017] | 14/15 | < 0.001 |
| StudyForrest 7T, emulated 2.1–4.0 mm | leave-one-subject-out | +0.014 [+0.001, +0.029] | 2/7 | 0.961 | −0.010 [−0.023, +0.005] | 5/7 | 0.148 |
| ds002702 7T, emulated 1.05 mm | leave-one-subject-out | +0.033 [+0.028, +0.037] | 0/5 | 1.000 | – | – | – |

| comparison | settings chosen | all vertices: model – best competitor [95% CI] | better | p | high mixing: model – best competitor [95% CI] | better | p |
|---|---|---|---|---|---|---|---|
| ds002702 7T, emulated 1.6 mm | leave-one-subject-out | +0.029 [+0.023, +0.034] | 0/5 | 1.000 | −0.002 [−0.022, +0.016] | 3/5 | 0.500 |
| ds002702 7T, emulated 2.4 mm | leave-one-subject-out | +0.003 [−0.004, +0.009] | 2/5 | 0.781 | +0.009 [−0.008, +0.026] | 2/5 | 0.781 |
| **Robustness of the HCP result** | | | | | | | |
| HCP 2.4 mm, 1 mm misregistration (all methods) | fixed on 5 other subjects | −0.006 [−0.011, −0.002] | 13/15 | 0.005 | −0.014 [−0.024, −0.003] | 12/15 | 0.013 |
| HCP 3.2 mm, 1 mm misregistration (all methods) | fixed on 5 other subjects | −0.008 [−0.013, −0.003] | 14/15 | 0.002 | −0.018 [−0.025, −0.010] | 14/15 | < 0.001 |
| HCP 2.4 mm, model PSF 15% too narrow | fixed on 5 other subjects | −0.009 [−0.013, −0.004] | 14/15 | 0.002 | −0.024 [−0.033, −0.014] | 13/15 | < 0.001 |
| HCP 2.4 mm, model PSF 15% too wide | fixed on 5 other subjects | −0.002 [−0.007, +0.002] | 10/15 | 0.227 | −0.018 [−0.027, −0.009] | 13/15 | 0.001 |
| HCP 3.2 mm, model PSF 15% too narrow | fixed on 5 other subjects | −0.016 [−0.020, −0.010] | 14/15 | < 0.001 | −0.029 [−0.036, −0.021] | 14/15 | < 0.001 |
| HCP 3.2 mm, model PSF 15% too wide | fixed on 5 other subjects | −0.005 [−0.009, +0.001] | 11/15 | 0.028 | −0.018 [−0.027, −0.009] | 13/15 | 0.002 |
| **Second acquisitions** | | | | | | | |
| StudyForrest 3T → 7T, voxel ratio ≥ 2.5 | other subjects of the pair type | +0.014 [−0.004, +0.040] | 2/5 | 0.844 | – | – | – |
| StudyForrest 7T → 7T, all pairs | other subjects of the pair type | +0.010 [−0.001, +0.022] | 2/7 | 0.945 | – | – | – |
| ds002702 3T 2.4 mm → 7T 0.8 mm | the other subject of the pair type | −0.024 [−0.028, −0.021] | 2/2 | 0.250 | −0.029 [−0.049, −0.008] | 2/2 | 0.250 |
| ds002702 7T SE 1.05 mm → GE 0.8 mm | the other subject of the pair type | +0.086 [+0.081, +0.090] | 0/2 | 1.000 | – | – | – |

Dashes: fewer than 30 strongly mixed vertices in some subject (ds002702 at 1.05 mm, within a session and as the SE source), or no such vertex set in that analysis (StudyForrest across sessions).

- **Only HCP emulation shows a consistent lead.** It holds in 12–14 of 15 subjects, also under a 1 mm misregistration and with a PSF assumed 15% too narrow. A PSF assumed 15% too wide removes the overall lead at 2.4 mm (−0.002, 10/15 subjects, p = 0.23), but not the lead where mixing is strong (−0.018, 13/15). In the confirmatory cohort the overall lead at 2.4 mm survives this PSF error (−0.004, 63 of 86 families, p < 0.001; Section 4.9).
- **Resolution-limited 7T data within a session.** The model trails overall. Where mixing is strong the difference is close to zero: −0.010 in StudyForrest (5/7, p = 0.15); −0.002 and +0.009 in ds002702 at 1.6 and 2.4 mm.
- **Second acquisitions.** Only ds002702 3T → 7T after field-map correction favours the model (2/2 subjects, too few for a test).

##### 4.9 Confirmatory cohort: 158 further HCP subjects

The plan of Section 3.9 was run on every remaining subject of the HCP 7T retinotopy release. All 158 passed the pipeline's checks. The table gives the tests on one subject per family (n = 86) and, in the last column, all 158 subjects with a family-level bootstrap; D and Δ are differences in residual (model − competitor; negative: model better), as in Section 4.4. The full table, the sensitivity analyses and the development-set look are in Supplementary S13.

| hyp. | test | grid | role | n | mean | in direction | bootstrap 95% | p | all subjects: n, mean [family bootstrap 95%] |
|---|---|---|---|---|---|---|---|---|---|
| H1 | D, contamination (native-map estimate) | 2.4 mm | primary | 86 | −0.0264 | 64/86 | [−0.0332, −0.0197] | < 0.001 | 158, −0.0250 [−0.0310, −0.0191] |
| H1 | D, contamination (native-map estimate) | 3.2 mm | primary | 86 | −0.0219 | 77/86 | [−0.0262, −0.0177] | < 0.001 | 158, −0.0188 [−0.0224, −0.0152] |
| H1 | D, contamination (coarse data only) | 2.4 mm | secondary | 86 | −0.0339 | 70/86 | [−0.0418, −0.0264] | < 0.001 | 158, −0.0302 [−0.0365, −0.0242] |
| H1 | D, contamination (coarse data only) | 3.2 mm | secondary | 86 | −0.0211 | 71/86 | [−0.0259, −0.0163] | < 0.001 | 158, −0.0182 [−0.0221, −0.0143] |
| H1 | D, plain far leak | 2.4 mm | secondary | 86 | −0.0201 | 57/86 | [−0.0271, −0.0133] | < 0.001 | 158, −0.0189 [−0.0246, −0.0135] |
| H1 | D, plain far leak | 3.2 mm | secondary | 86 | −0.0049 | 48/86 | [−0.0092, −0.0008] | 0.015 | 158, −0.0038 [−0.0072, −0.0004] |
| H1 | overall gain, all coherent vertices | 2.4 mm | secondary | 86 | −0.0087 | 82/86 | [−0.0099, −0.0075] | < 0.001 | 158, −0.0089 [−0.0099, −0.0079] |
| H1 | overall gain, all coherent vertices | 3.2 mm | secondary | 86 | −0.0128 | 85/86 | [−0.0141, −0.0114] | < 0.001 | 158, −0.0126 [−0.0136, −0.0115] |
| H2 | D_far(wedges) − D_far(rings) | 3.2 mm | primary (H2a) | 86 | −0.0047 | 46/86 | [−0.0087, −0.0006] | 0.010 | 158, −0.0041 [−0.0065, −0.0016] |
| H2 | D_far(wedges) − D_far(rings) | 2.4 mm | interval | 86 | −0.0076 | 48/86 | [−0.0142, −0.0010] | – | 158, −0.0077 [−0.0126, −0.0028] |
| H2 | D, contamination (rings) | 2.4 mm | primary (H2b) | 86 | −0.0263 | 71/86 | [−0.0327, −0.0199] | < 0.001 | 158, −0.0268 [−0.0326, −0.0209] |
| H2 | D, contamination (rings) | 3.2 mm | primary (H2b) | 86 | −0.0209 | 79/86 | [−0.0243, −0.0176] | < 0.001 | 158, −0.0210 [−0.0239, −0.0179] |
| H2 | D_CWL(rings) − D_far(rings) | 3.2 mm | secondary (H2c) | 86 | −0.0207 | 78/86 | [−0.0243, −0.0170] | < 0.001 | 158, −0.0213 [−0.0243, −0.0182] |
| H2 | D, plain far leak (rings) | 2.4 mm | descriptive | 86 | −0.0125 | 53/86 | [−0.0195, −0.0060] | – | 158, −0.0112 [−0.0166, −0.0058] |
| H2 | D, plain far leak (rings) | 3.2 mm | descriptive | 86 | −0.0002 | 40/86 | [−0.0036, +0.0031] | – | 158, +0.0003 [−0.0027, +0.0032] |
| H3 | Δ_clean(half) − Δ_clean(full), > 0 predicted | 2.4 mm | primary (H3a) | 86 | +0.0032 | 71/86 | [+0.0023, +0.0041] | < 0.001 | 158, +0.0032 [+0.0026, +0.0039] |
| H3 | Δ_clean(half) − Δ_clean(full), > 0 predicted | 3.2 mm | primary (H3a) | 86 | +0.0010 | 53/86 | [+0.0002, +0.0018] | 0.007 | 158, +0.0010 [+0.0005, +0.0016] |
| H3 | D_far(half) − D_far(full) | 2.4 mm | interval (H3b) | 86 | −0.0002 | 42/86 | [−0.0020, +0.0016] | – | 158, +0.0004 [−0.0009, +0.0017] |
| H3 | D_far(half) − D_far(full) | 3.2 mm | interval (H3b) | 86 | +0.0004 | 39/86 | [−0.0007, +0.0015] | – | 158, −0.0002 [−0.0010, +0.0008] |
| H3 | Δ_clean(half) | 2.4 mm | secondary (H3c) | 86 | −0.0063 | 71/86 | [−0.0082, −0.0046] | < 0.001 | 158, −0.0070 [−0.0084, −0.0056] |
| H3 | Δ_clean(half) | 3.2 mm | secondary (H3c) | 86 | −0.0152 | 85/86 | [−0.0168, −0.0136] | < 0.001 | 158, −0.0150 [−0.0162, −0.0138] |
| H4 | overall gain, PSF 15% too wide | 2.4 mm | primary | 86 | −0.0038 | 63/86 | [−0.0051, −0.0025] | < 0.001 | 158, −0.0041 [−0.0052, −0.0030] |
| H4 | overall gain, PSF 15% too wide | 3.2 mm | secondary | 86 | −0.0050 | 64/86 | [−0.0065, −0.0036] | < 0.001 | 158, −0.0048 [−0.0059, −0.0036] |
| H4 | overall gain, PSF 15% too narrow | 2.4 mm | secondary | 86 | −0.0109 | 85/86 | [−0.0121, −0.0097] | < 0.001 | 158, −0.0111 [−0.0121, −0.0101] |
| H4 | overall gain, PSF 15% too narrow | 3.2 mm | secondary | 86 | −0.0158 | 85/86 | [−0.0171, −0.0145] | < 0.001 | 158, −0.0156 [−0.0167, −0.0146] |

Secondary tests are Holm-corrected within H1–H4: all corrected $p < 0.001$ except the plain-leak contrast at 3.2 mm (0.015). There are eight primary tests and no correction across them was planned; three results are small and would not survive one (Bonferroni 0.05/8 = 0.006): H2a ($p = 0.010$, 46 of 86 in the predicted direction), H3a at 3.2 mm ($p = 0.007$, 53 of 86) and the plain-leak contrast at 3.2 mm ($p = 0.015$, 48 of 86). The curves of Fig. 7 use all 158 subjects; the tests use the 86.

- **The contamination dose-response replicates (H1).** D is negative at both grids (64 and 77 of 86 families, $p < 0.001$), as is the version from the coarse data alone (70 and 71 of 86). The effect is smaller than in the development set at 3.2 mm (−0.022 against −0.037): the development set was optimistic in the highest-contamination bin. The overall gain over the better smoothing competitor holds in 82 and 85 of 86 families. The plain far-leak contrast is clear at 2.4 mm (−0.020, 57/86) and weak at 3.2 mm (−0.005, 48/86, $p = 0.015$). The contamination results do not depend on the bin conventions (the 2–5% bin as reference, the 10–20% bin as top bin, a slope over all bins or Spearman's ρ all give $p < 0.001$; Supplementary S13.6); the plain-leak contrast at 3.2 mm does (−0.009 with the 2–5% reference, +0.000 with the top bin fixed at 10–20%, slope −0.0013 per bin, $p = 0.006$). Two of 86 subjects contributed a 10–20% rather than a > 20% top bin at 2.4 mm, none at 3.2 mm.
- **Rings (H2).** Contamination predicts the model's gain for eccentricity maps as for polar angle (H2b: 71 and 79 of 86, $p < 0.001$), and at 3.2 mm it does so where the plain leak predicts nothing ($D_{far}$ for rings −0.0002 [−0.0036, +0.0031]; H2c: −0.021, 78/86). At 2.4 mm rings carry a far-leak dose-response of their own (−0.0125, 53/86), smaller than the wedges' (the difference −0.008 [−0.014, −0.001]); at 3.2 mm the difference is small (H2a: −0.005, 46/86, $p = 0.010$; the development set's 15/15 was optimistic, as for H1, and the effect stays small under every bin convention, $p = 0.004$–$0.013$). Rings are therefore not a clean negative control in HCP; contamination is the better predictor for both stimuli (Fig. 7).
- **Source noise (H3).** With half the source data and the settings held at their full-run values, the gain in the clean bin shrinks (+0.003 at 2.4 mm, 71/86, $p < 0.001$; +0.001 at 3.2 mm, 53/86, $p = 0.007$), the leak-dependent gain does not change (the 95% intervals [−0.002, +0.002] and [−0.001, +0.002] lie inside an equivalence margin of ±0.003, a tenth of the top-bin gain, chosen after the fact), and the model still beats the competitor (71/86 and 85/86). Re-tuned for half-run sources on the tuning subjects, the rule doubles both prior parameters at both grids and leaves the competitor's smoothing nearly unchanged; with both methods re-tuned, the clean-bin gain in the 15 development subjects still shrinks with half the data (+0.0026, 14 of 15, $p = 0.003$, at 2.4 mm; +0.0022, 13 of 15, $p = 0.006$, at 3.2 mm; with fixed settings +0.0041 and +0.0024 in the same subjects), and the half-run model still beats the re-tuned competitor (−0.005 and −0.011 over all coherent vertices) (Fig. S1D). The gain where no far unmixing is needed is therefore not denoising, and the far-part ablation of Section 4.3 shows that neither it nor the dose-dependent part is far unmixing: both survive without the far part of the operator (Section 5).
- **PSF (H4).** With the PSF assumed 15% too wide the overall lead at 2.4 mm remains (−0.004, 63/86, $p < 0.001$); the development set's $p = 0.23$ (Section 4.8) was a matter of power, not of sign. The other PSF cases hold in 64–85 of 86.

- **Sensitivity.** Adding the five relatives of the tuning subjects (163 subjects, 91 families) changes no conclusion (D −0.027 and −0.022; H2a −0.005, p = 0.004; H3a +0.003 and +0.001; H4 −0.004; Supplementary S13.3). Alone, the five give D = −0.043 and −0.018 (5/5 each; descriptive).

##### 4.10 External replication: 35 subjects at 3T

The rule of Section 3.10 chose a model range of 12 mm and amplitude 2 at 3.0 mm and of 6 mm and 1 at 4.0 mm, much weaker priors than in HCP, and upsampling with 2 mm smoothing as the competitor at both grids: smoothing barely helped any method, because each half-map averages two to six runs. On the 35 included test subjects (Fig. 8; Supplementary S16):

| test | 3.0 mm (×1.5) | 4.0 mm (×2.0) |
|---|---|---|
| N1: overall gain, responsive vertices | +0.005 [+0.001, +0.008], 13/35 (not replicated) | −0.007 [−0.011, −0.004], 27/35, Holm p < 0.001 |
| N2: D_far | −0.002 [−0.011, +0.006], 19/35, Holm p = 0.932 | +0.003 [−0.004, +0.011], 18/35, Holm p = 1.000 |
| N3: D_CWL-M | −0.015 [−0.026, −0.005], 25/35, Holm p = 0.018 | −0.011 [−0.019, −0.003], 25/35, Holm p = 0.029 |

- **Overall gain.** The model's advantage replicated at the stronger coarsening (27 of 35 subjects; in cortex clean at 2.0 mm with leak > 10%, −0.014, 28 of 35) but not at the milder one, where upsampling with 2 mm smoothing was better in 22 of 35 subjects and plain upsampling better still (+0.009).
- **Contamination, not leak.** The gain grew with contamination at both grids — at 3.0 mm from +0.010 in the lowest bin, where the model loses, to −0.005 in the highest, where it wins — but not with the plain far leak. This is the dissociation of Section 4.4 (leak is necessary, not sufficient) in an independent dataset.
- **Where it failed.** In the pericalcarine cortex the model was worse at both grids (+0.013 and +0.009), and its polar-angle error was slightly larger (+1.7° and +0.7°). A tuning subject with a poor registration (boundary-based cost 0.68, twice the usual distortion field) lost at every grid: the model depends on geometry in a way that upsampling does not.
- **Leak of a 3T acquisition.** For the 2.0 mm acquisition itself, the mean far leak of standard mapping is 3.2% and 19% of cortex lies above 5% (43 subjects, individual maps); at the 3.0 and 4.0 mm emulations 4.2% and 27%, 5.0% and 33% — a typical 3T acquisition at 2 mm leaks about as much as HCP's 7T data emulated at 2.4 mm.

#### Supplementary figures

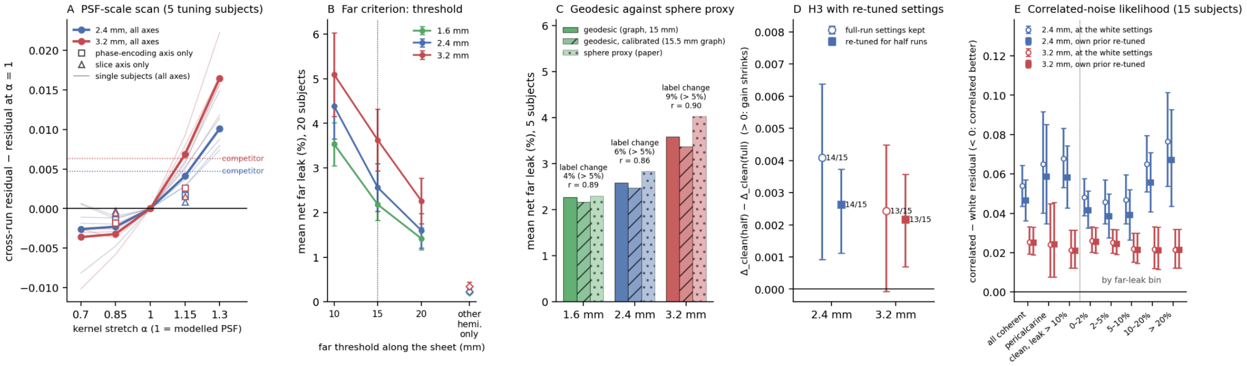


**Figure S1. Four further analyses (main-text Figure 10 of the long version).** (A) Kernel-width scan: cross-run residual relative to the modelled kernel ($\alpha = 1$) against the stretch factor $\alpha$ applied to the inversion's kernels on all axes (thin lines: the five tuning subjects; bold: their mean) and on the phase-encoding or the slice axis alone (open symbols, mean), prior settings fixed; dotted lines mark the level of the best competitor (upsampling + 8 mm smoothing). (B) Far criterion: mean net far leak of standard mapping (± SD over 20 development subjects) with the far threshold at 10, 15 and 20 mm along the sheet, and for the other hemisphere alone. (C) True geodesic against the sphere proxy (5 tuning subjects): mean net far leak with the far label from the graph geodesic on the midthickness mesh at 15 mm, from the same geodesic at the threshold calibrated against Workbench's exact geodesic, and from the area-scaled sphere used throughout; numbers: the share of cortex whose > 5% label differs between the geodesic and the sphere criteria, and the per-vertex correlation of the two maps. (D) H3 with settings re-tuned for half-run sources: $\Delta\_clean(half) - \Delta\_clean(full)$ (15 development subjects; mean and bootstrap 95% interval; numbers: subjects above zero) with the full-run settings kept and with the model and the competitor re-tuned on the tuning subjects' half runs. (E) Correlated-noise likelihood on HCP: residual of the correlated-noise fit minus the white-noise fit, by stratum (15 development subjects; mean and bootstrap 95% interval); negative favours the correlated-noise likelihood.

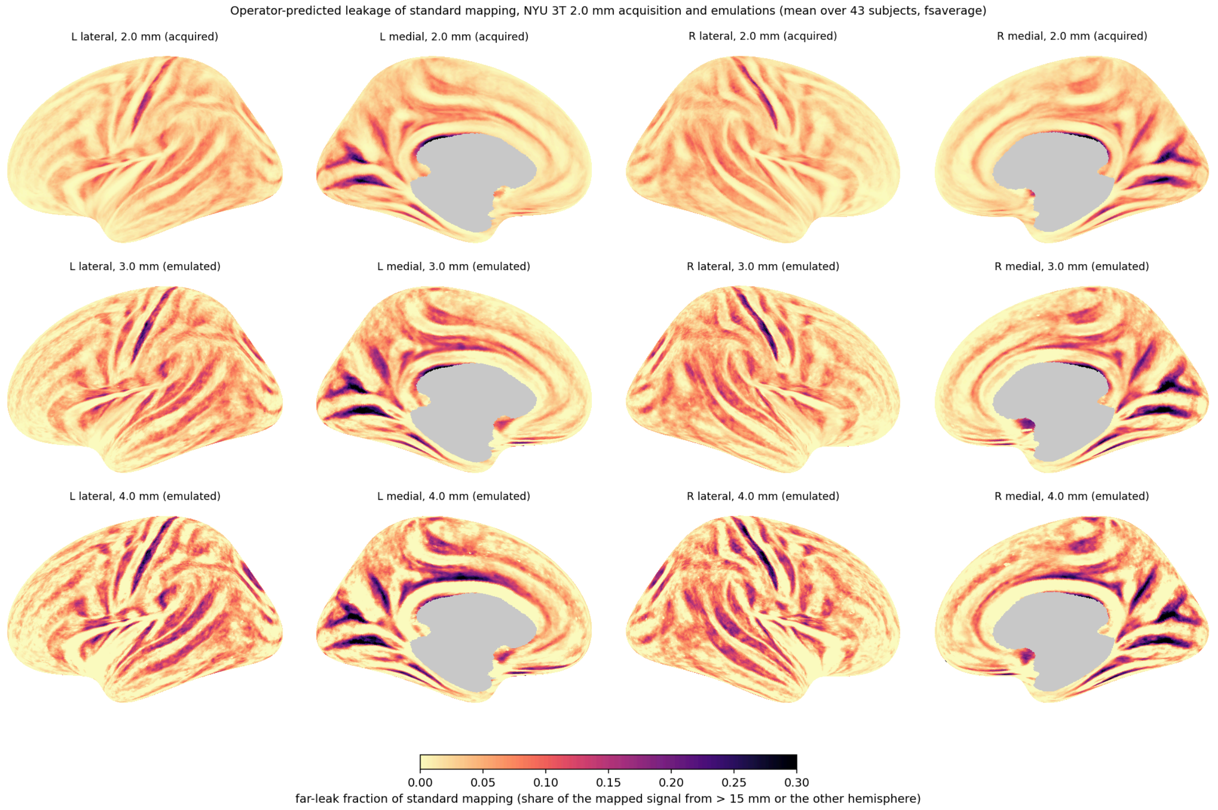


**Figure S2. Operator-predicted far leak of standard mapping for the NYU 3T 2.0 mm acquisition and its 3.0 and 4.0 mm emulations, mean over the included subjects on fsaverage.** Lateral and medial views of the inflated surfaces; per subject the far-leak map was carried from the subject's registration sphere to fsaverage (three nearest vertices, inverse-distance weights) and averaged; grey: medial wall and vertices covered in fewer than half of the subjects. Colour scale as Figure 4 (0–0.3). GIFTI files of the mean and of the share of subjects above 5% are released with the code (S16.5).
